# Phylogeny, morphology and the role of hybridization as driving force of evolution in grass tribes Aveneae and Poeae (Poaceae)

**DOI:** 10.1101/707588

**Authors:** Natalia Tkach, Julia Schneider, Elke Döring, Alexandra Wölk, Anne Hochbach, Jana Nissen, Grit Winterfeld, Solveig Meyer, Jennifer Gabriel, Matthias H. Hoffmann, Martin Röser

## Abstract

To investigate the evolutionary diversification and morphological evolution of grass supertribe Poodae (subfam. Pooideae, Poaceae) we conducted a comprehensive molecular phylogenetic analysis including representatives from most of their accepted genera. We focused on generating a DNA sequence dataset of plastid *matK* gene–3’*trnK* exon and *trnL– trnF* regions and nuclear ribosomal ITS1–5.8S gene–ITS2 and ETS that was taxonomically overlapping as completely as possible (altogether 257 species). The idea was to infer whether phylogenetic trees or certain clades based on plastid and nuclear DNA data correspond with each other or discord, revealing signatures of past hybridization. The datasets were analysed using maximum parsimony, maximum likelihood and Bayesian approaches. Instances of severe conflicts between the phylogenetic trees derived from both datasets, some of which have been noted earlier, unambiguously point to hybrid origin of several lineages (subtribes, groups of genera, sometimes genera) such as Phalaridinae, Scolochloinae, Sesleriinae, Torreyochloinae; *Arctopoa*, *Castellia*, *Graphephorum*, *Hyalopodium*, *Lagurus*, *Macrobriza*, *Puccinellia* plus *Sclerochloa*, *Sesleria*, *Tricholemma*, American *Trisetum*, etc. and presumably Airinae, Holcinae and Phleinae. ‘*Calamagrostis*’ *flavens* appears to be an intergeneric hybrid between *Agrostis* and *Calamagrostis*. Most frequently there is good agreement of other regions of the trees, apart from intrinsic different phylogenetic resolution of the respective DNA markers. To explore the to date rather unclear morphological evolution of our study group a data matrix encompassing finally 188 characters was analysed for ancestral state reconstructions (ASR) using the tree from the combined molecular dataset as presumably best approximation to the species phylogeny. For 74 characters ASRs were feasible and yielded partly surprising results for the study group as a whole but also for some of their subdivisions. Considering taxonomy and classification it became evident, that many morphological characters show a very high degree of homoplasy and are seemingly able to change within comparatively short timespans in the evolution of our grasses. Most of the taxonomic units distinguished within our study group, e.g. as subtribes, are defined less by consistent morphological characters or character combinations and should be rather understood as clades revealed by molecular phylogenetic analysis. One reason for this extreme homoplasy concerning traditionally highly rated characters of inflorescences or spikelets and their components might be that they have little to do with pollination (always wind) or adaptation to pollinators as in other angiosperms but rather with dispersal and diaspores. Easily changing structure of spikelet disarticulation, of glume, lemma or awn characters might be advantageous in the rapid adaptation to different habitats and micro-habitats, which was evidently most successfully accomplished by these grasses. A partly revised classification of Poodae is presented, including a re-instatement of tribes Aveneae and Poeae s.str. Following a comparatively narrow delineation of preferably monophyletic subtribes, Antinoriinae, Avenulinae, Brizochloinae, Helictochloinae, Hypseochloinae are described as new. New genera are *Arctohyalopoa* and *Hyalopodium*. New combinations are *Arctohyalopoa lanatiflora*, *A. lanatiflora* subsp. *ivanoviae*, *A. lanatiflora* subsp. *momica*, *Colpodium biebersteinianum*, *C. kochii*, *C. trichopodum*, *C. verticillatum*, *Deschampsia micrathera*, *Dupontia fulva*, *Festuca masafuerana*, *Hyalopodium araraticum*, *Paracolpodium baltistanicum*, *Parapholis cylindrica*, *P*. ×*pauneroi*. *Festuca masatierrae* is a new name.

**Supporting Information** may be found online in the Supporting Information section at the end of the article.

## INTRODUCTION

The grass supertribe Poodae with Poeae sensu lato (s.l.) as sole tribe (i.e., including Aveneae) encompasses 106–121 genera, depending on the respective width of their delineation, and 2562–2578 species (Kellogg, 2015; Soreng & al., 2017). It is a characteristic group of C_3_ grasses proliferating in the northern temperate and boreal regions and represented by many annuals especially in the Mediterranean/Near East, a region that was also the cradle of *Avena* with cultivated oat(s). Economically enormously significant are also the pasture and forage grasses. Poodae are scarce in the subtropics and tropics but bridge them on the top of high mountains and have a second centre of diversity in the temperate and cool zones of the southern hemisphere. Concepts of relationship in this group of grasses based on morphological characters were summarized by *Genera graminum* (Clayton & Renvoize, 1986) that served as an important basis for later molecular phylogenetic studies. Due to the sheer size of the group, usually representative genera were selected since then for comparative studies to gain an overview on the whole Poodae and their major groupings using traditional Sanger and, more recently, plastid genome sequencing (Soreng & Davis, 2000; Davis & Soreng, 2007; Döring & al., 2007; Quintanar & al., 2007; Soreng & al., 2007; Schneider & al., 2009; Saarela, & al., 2015, 2018; Pimentel & al., 2017; Orton & al., 2019). Other studies focused on special groups using an in-depth sampling of taxa, for example, within Aveneae (Grebenstein & al., 1998; Döring, 2009; Saarela & al., 2010, 2017; Wölk & Röser, 2014, 2017; Barberá & al., 2019) and Poeae sensu stricto (s.str.; Schneider & al., 2012; Birch & al., 2014, 2017), in which especially the subtribes Poinae (Hunter & al., 2004; Gillespie & Soreng, 2005; Gillespie & al., 2007, 2008, 2009, 2010, 2018; Refulio-Rodríguez & al., 2012; Hoffmann & al., 2013; Soreng & al., 2010, 2015a; Nosov & al., 2015, 2019), Loliinae (Torrecilla & Catalán, 2002, Catalán & al., 2004; Torrecilla & al., 2004; Catalán & al., 2007; Inda & al., 2008; Cheng & al., 2016; Minaya & al., 2017), Brizinae and Calothecinae (Essi & al., 2008; Persson & Rydin, 2016) or Sesleriinae (Kuzmanović & al., 2017) were studied.

Recent classifications of Poodae took up the progress made by molecular phylogenetic studies and numerous changes in classification proposed relative to *Genera graminum*, which was superseded by the comprehensive Poaceae treatment for *The families and genera of vascular plants* (Kellogg, 2015). The recent taxonomic accounts on grasses of Soreng & al. (2015, 2017) and Kellogg (2015) abandoned the traditional distinction of tribes Aveneae and Poeae s.str. since molecular phylogenetic data did not corroborate their separation according to their previous circumscription based on morphology. Nevertheless, the occurrence of two different plastid DNA sequences (“Aveneae type” and “Poeae type”) led to the nomenclaturally informal recognition of two lineages (Soreng & Davis, 2000), each of which was divided in a number of subtribes (Soreng & al., 2007, 2017; Kellogg, 2015) as followed also in the molecular phylogenetic account on the Aveneae type lineage of Saarela & al. (2017).

Hybridization between species is a widespread process that acts in almost any group of grasses. It is well-known to be especially frequent in connection with polyploidy and within polyploid complexes, as documented in may grass groups including the economically highly important Triticeae, Andropogoneae and Paniceae (Hunziker & Stebbins, 1987; Kellogg & Watson, 1993; Kellogg, 2015). Hybridization was also considered a potential reason for the discrepancies between traditional morphology-and molecular phylogeny-based classification for Poodae (Soreng & Davis, 2000).

To address the role of hybridization as suspected factor in the evolution of several of its lineages (Soreng & Davis, 2000; Quintanar & al., 2007; Soreng & al., 2007) and to contribute to an improved classification we aimed at a comparative sequencing study of representatives of most genera of Poodae, except for some lineages that were already shown to be clearly monophyletic (e.g., Calothecinae, Loliinae, Poinae), in which we sampled only a small selection of taxa. Due to the different inheritance of the plastid and the nuclear genome we tried to generate a taxonomically overlapping dataset for both genomes, since incongruent placement of taxa in phylogenetic trees derived from both individual datasets is a reliable indicator of past hybridization events.

Secondly, we attempted to clarify the phylogenetic position of several genera previously not sampled in molecular phylogenetic investigations, to address questionable data in DNA sequence repositories and to correct a few problem cases we have created ourselves in previous publications from our lab.

Finally, we wanted to compare the molecular phylogenetic information with a morphological matrix for the molecularly studied taxa and performed an extensive analysis of available and newly collected morphological data. For characters sufficiently densely scored for our taxa in question we conducted an ancestral state reconstruction using the molecular phylogenetic information.

## MATERIAL AND METHODS

### Classification employed

We follow in this study, as far as possible, the classification of grass subfamily Pooideae displayed by Soreng & al. (2017). This classification utilizes a comparatively narrow delineation of subtribes and the rather infrequently used taxonomic ranks of supersubtribes and supertribes. It is easy to compare with the classification used by Kellogg (2015) for her account on Poaceae in *The families and genera of vascular plants*. We also follow the treatment of genera and synonyms presented by Soreng & al. (2017) unless otherwise stated. Genus names occasionally misapplied in the literature are enclosed in the following by single quotation marks.

### Plant material and choice of study taxa

For the molecular phylogenetic study we tried to sample as complete as possible all currently acknowledged genera and important segregate genera of Poodae except for subtribes Calothecinae, Loliinae and Poinae (see Introduction). The types of the genera were preferably included. For information retrieval on nomenclatural types we consulted the *Index nominum genericorum* (ING; botany.si.edu/ing/), *Tropicos* (tropicos.org), Clayton & Renvoize (1986), Clayton & al. (2002 onwards), *Catalogue of New World grasses* (Soreng & al., 2000 onwards) and other taxonomic sources (see References). In non-monospecific genera we tried to investigate two or more species. Sometimes, we used more than one accession for the same taxon. In total, 117 accepted genera and 257species were treated in this study. No plant material has been obtained in the genera *Agropyropsis* A.Camus, *Agrostopoa* Davidse, Soreng & P.M.Peterson, and *Pseudophleum* Doğan. Taxa selected from the lineages next to Poodae, namely *Hordelymus europaeus* (L.) O.E.Harz, *Hordeum marinum* Huds. subsp. *gussoneanum* (Parl.) Thell. and *Secale sylvestre* Host from Triticeae subtribe Hordeinae in the sense of Schneider & al. (2009), *Boissiera squarrosa* (Sol.) Nevski and *Bromus erectus* Huds. from Triticeae subtribe Brominae, *Littledalea tibetica* Hemsl. from Triticeae subtribe Littledaleinae as well as *Brachypodium distachyon* (L.) P.Beauv. from Brachypodieae were selected as suitable outgroup taxa based on previous studies (Davis & Soreng, 1993; Catalán & al., 1997; Hilu & al., 1999; Schneider & al., 2009, 2011; GPWG, 2012; Blaner & al., 2014; Hochbach & al., 2015). The molecular phylogenetic studies were conducted using silica gel-dried leaf material collected in the field from living plants or leaves from specimens of the following herbaria: AD, ALTB, B, BBG, C, CAN, CHR, COL, FI, HAL, HO, ICN, JACA, K, LE, LISU, MEXU, MO, NS, NSK, NSW, NU, NY, PRE, RO, RSA, SGO, TROM, UPS, US (abbreviations according to *Index herbariorum;* http://sweetgum.nybg.org/science/ih/). Information on origin, collectors, collection details and ENA/GenBank sequence accession numbers of the analysed taxa is given in Appendices 1, 2.

### Molecular methods and sequence alignment

FastPrep FP120 cell disrupter (Qbiogene, Heidelberg, Germany) was used to homogenize 20–45 mg leaf tissue per sample. Extraction of total genomic DNA was conducted with the NucleoSpin Plant Kit in accordance to the manufacturer’s protocol (Macherey-Nagel, Düren, Germany). The concentration of the DNA samples was checked with a NanoDrop spectrophotometer (Thermo Fisher Scientific, Waltham, USA). The entire internal transcribed spacer region (ITS) of the nuclear ribosomal (nr) DNA (ITS1–5.8S rRNA gene–ITS2) and the *matK* gene–3’*trnK* exon of the plastid DNA were PCR-amplified following the protocols described by Schneider & al. (2009) and Wölk & Röser (2014). The 3’ end of the external transcribed spacer region (ETS) of the nrDNA was amplified with primers 18S-Rcyper (Starr & al., 2003), RETS4-F (Gillespie & al., 2010) and RETS-B4F (Alonso & al., 2014) under conditions following Tkach & al. (2008). For amplification of the plastid non-coding region of the *trnL*–*trnF*, including *trnL*(UAA) intron and adjacent intergenic spacer between the *trnL*(UAA) 3′exon and *trnF*(GAA) gene, were used primers c, d, e and f and the PCR protocol of Taberlet & al. (1991). Additional new primers created for this region (cps ACGGACTTGATTGTATTGAGCC; dps CTCTCTCTTTGTCCTCGTCCG; eps CGGACGAGGACAAAGAGAGAG; fps AACTGAGCATCCTGACCTTTTCTTG) were used in combination with the primers cited. PCR was carried out on a thermocycler manufactured by Eppendorf (Hamburg, Germany). Purification and sequencing of all PCR products were performed in our lab or by StarSEQ (Mainz, Germany), Eurofins MWG Operon (Ebersberg, Germany) and LGC Genomics (Berlin, Germany) with the same primers as used for amplifications. PCR products of the ITS region with ambiguous sequence peaks were cloned. Cloning was performed using the pGEM-T Easy Vector System (Promega Mannheim, Germany). Ligation and transformation of the relevant purified amplicons were carried out according to the technical manual. The plasmid DNA was isolated using the GeneJET Plasmid Miniprep Kit (Fermentas, St. Leon-Rot, Germany) according to the manufacturer’s protocol. The PCR products were quantified spectrophotometrically. Highly similar ITS clone sequences were combined to one consensus sequences to reduce the number of singletons in the alignment. All sequences were edited by eye in Sequencher 5.0 (Gene Codes Corporation, Ann Arbor, USA). The automatically performed alignments by using ClustalW2 (Larkin & al., 2007) were manually adjusted in Geneious 9.1.6 (https://www.geneious.com; Kearse & al., 2012).

### Phylogenetic analysis

Sequences generated in this or previous studies of our lab could be used in many taxa (Appendix 1). For comparison with our own data and to complete our datasets we included publicly available sequences for the taxa and sequence regions in question in the alignments (Appendix 2). The nuclear (including ITS and ETS sequences) and plastid (including *matK* gene–3’*trnK* exon and *trnL–trnF*) DNA sequence datasets were first analysed separately using the phylogenetic approaches of Maximum Likelihood (ML), Maximum Parsimony (MP) and Bayesian Inference (BI) following Tkach & al. (2019). All trees were visualized with FigTree 1.4.3 (http://tree.bio.ed.ac.uk/software/figtree/). Support values are cited in the text in the following sequence: ML bootstrap support/MP bootstrap support/Bayesian posterior probability (PP).

To avoid redundancy and to improve the clarity of the phylogenetic trees we finally omitted unnecessary duplicate sequences for the same taxon from the alignments. The final DNA sequence matrices for Poodae and outgroups are provided as fasta files in the supporting information (suppl. Appendix S1). The tree topologies obtained from the individual nuclear and plastid DNA sequence alignments were examined visually for incongruity. The node bootstrap support of ≥ 70 in ML analysis was chosen as value for supported incongruence (Wiens, 1998; Schneider & al., 2009; Baker & al., 2011; Pirie, 2015; Tkach & al., 2015, 2019). Since significantly conflicting relationships occurred as localized incongruences caused by specific taxa or clades in the individual phylogenetic trees, we combined the sequences of the nuclear and plastid markers in a second round of analyses to a concatenated dataset. It was analysed as described above to obtain a molecular total evidence tree that served as guideline for taxonomic classification.

### Morphological analysis and character mapping

For morphological analysis we compiled for our study group a data matrix of 188 characters in total. Supplementary Appendix S2 is the character list with coding of character states used to assess character evolution. Supplementary Fig. S2 graphically displays the characters states for the taxa studied. Morphological characteristics were gathered from various published resources, especially the genus and species data of *GrassBase* (Clayton & al., 2006 onwards), data from *Grass genera of the world* (Watson & al., 1992 onwards; Watson & Dallwitz, 1994) and from own observations using an incident light microscope Stemi 508doc equipped with a digital camera AxioCam ERc (Zeiss, Oberkochen, Germany).

We commonly used the genus data in the morphological character matrix. Exceptions were made when genus boundaries did not match our genus circumscriptions or when no data were available for a genus. Then we used species data and checked them for their applicability for the whole genus and eventually modified them accordingly. Character mapping on the molecular phylogenetic tree derived from the concatenated plastid and nuclear DNA sequence dataset was conducted in R version 3.5.0 with the package Phytools (Revell, 2012, 2013).

### Ancestral state reconstruction

For ancestral state reconstruction (ASR; suppl. Appendix S3) we selected 74 sufficiently variable of the 188 morphological characters studied. To avoid exceedingly large uncertainties, we considered for ASR only characters that were known in more than 145 (67%) of the 218 taxa scored for morphological data. ASR was also conducted in R version 3.5.0 using package phytools (Revell, 2012) by stochastic character mapping (Huelsenbeck & al., 2003). Therefore, we generated 1.000 stochastic character maps from the dataset using the ER (equal rate) model and obtained posterior probabilities for the nodes by averaging the state frequencies across all maps. In the case of unknown states, we used a prior probability distribution on the tips that is flat across all possible states. According to Revell (see http://blog.phytools.org/2016/10/stochastic-mapping-discrete-character.html) this leaves the posterior probabilities at internal nodes largely unaffected compared to just dropping tips with unknown states from the tree.

### Scanning electron microscopes (SEM) observation

Lemmas and awns of selected representatives of major lineages were investigated using scanning electron microscopy. The lemmas of the samples (legends to Figs. 9–11) were mounted under an incident light microscope on aluminium stubs covered by double stick carbon conductive tabs (Plano, Wetzlar, Germany). Samples were gold-coated using sputter coater MED010 (Balzers Union, Balzers, Liechtenstein). Images were taken on the tabletop scanning electron microscope TM-3030Plus (Hitachi Europe, Maidenhead, UK) with 5 kV acceleration voltage and the secondary electron detector.

## RESULTS

### Plastid DNA analysis

The plastid *matK* gene–3’*trnK* exon DNA sequence dataset for 208 taxa included 2,530, the *trnL–trnF* DNA dataset for 199 taxa included 1,414 aligned positions, respectively. The combined data matrix of the two plastid DNA markers for 214 taxa included a total of 3,927 aligned positions, of which 1,470 were variable (*matK* gene– 3’*trnK* exon: 984, *trnL–trnF*: 488) and 907 parsimony-informative (*matK* gene–3’*trnK* exon: 622, *trnL–trnF*: 286).

Poodae were corroborated a monophyletic lineage (100/100/1.00), using our extended set of outgroup taxa. It split in two main clades, which were supported by 100/98/1.00 and 100/99/1.00, respectively (Fig. 1). One of the lineages agreed with the “Poeae chloroplast group 1 (Aveneae type)” as termed by Soreng & al. (2007) or tribe Aveneae as suggested here to use a taxonomic rank.

**Fig. 1.**
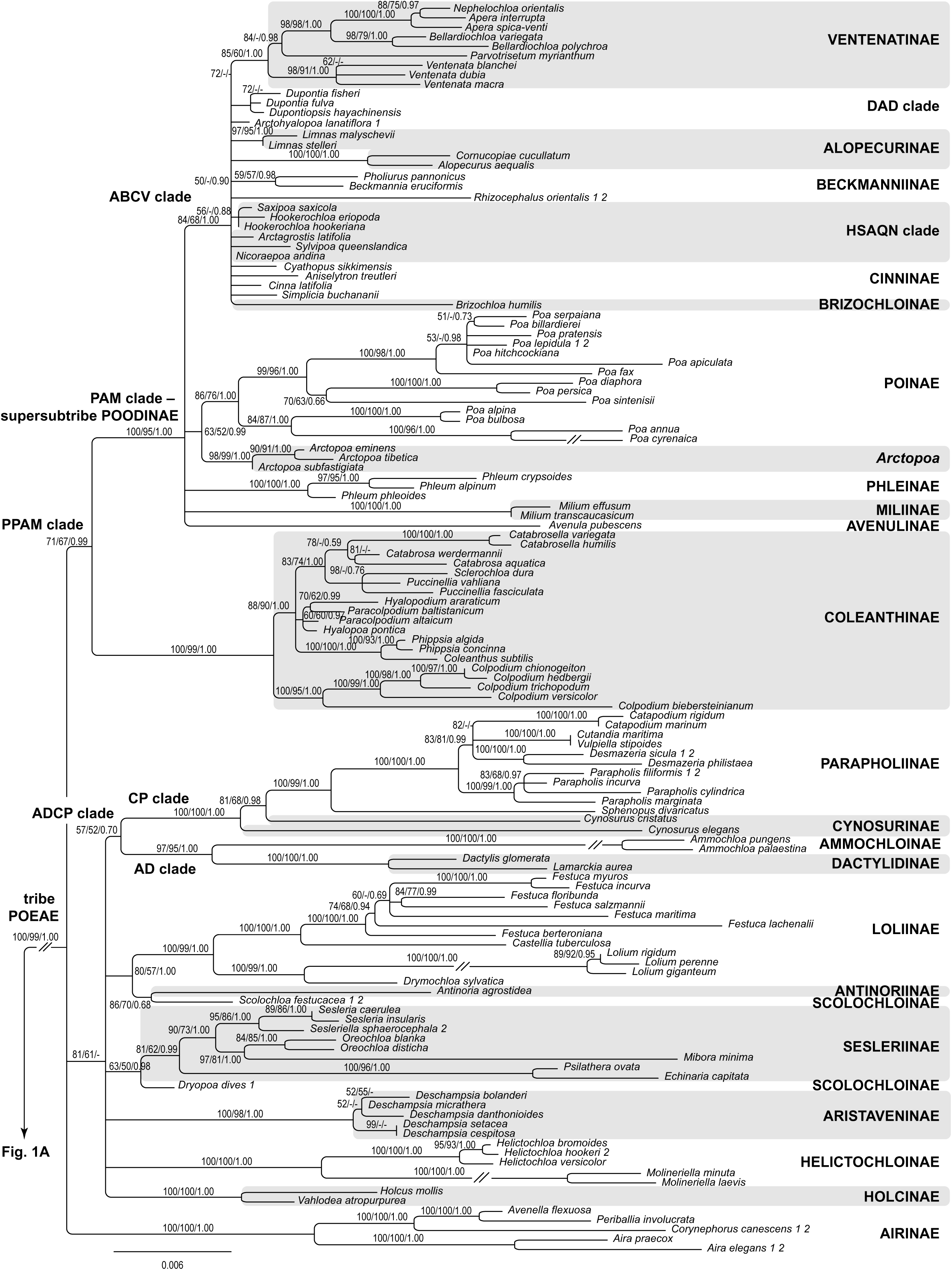
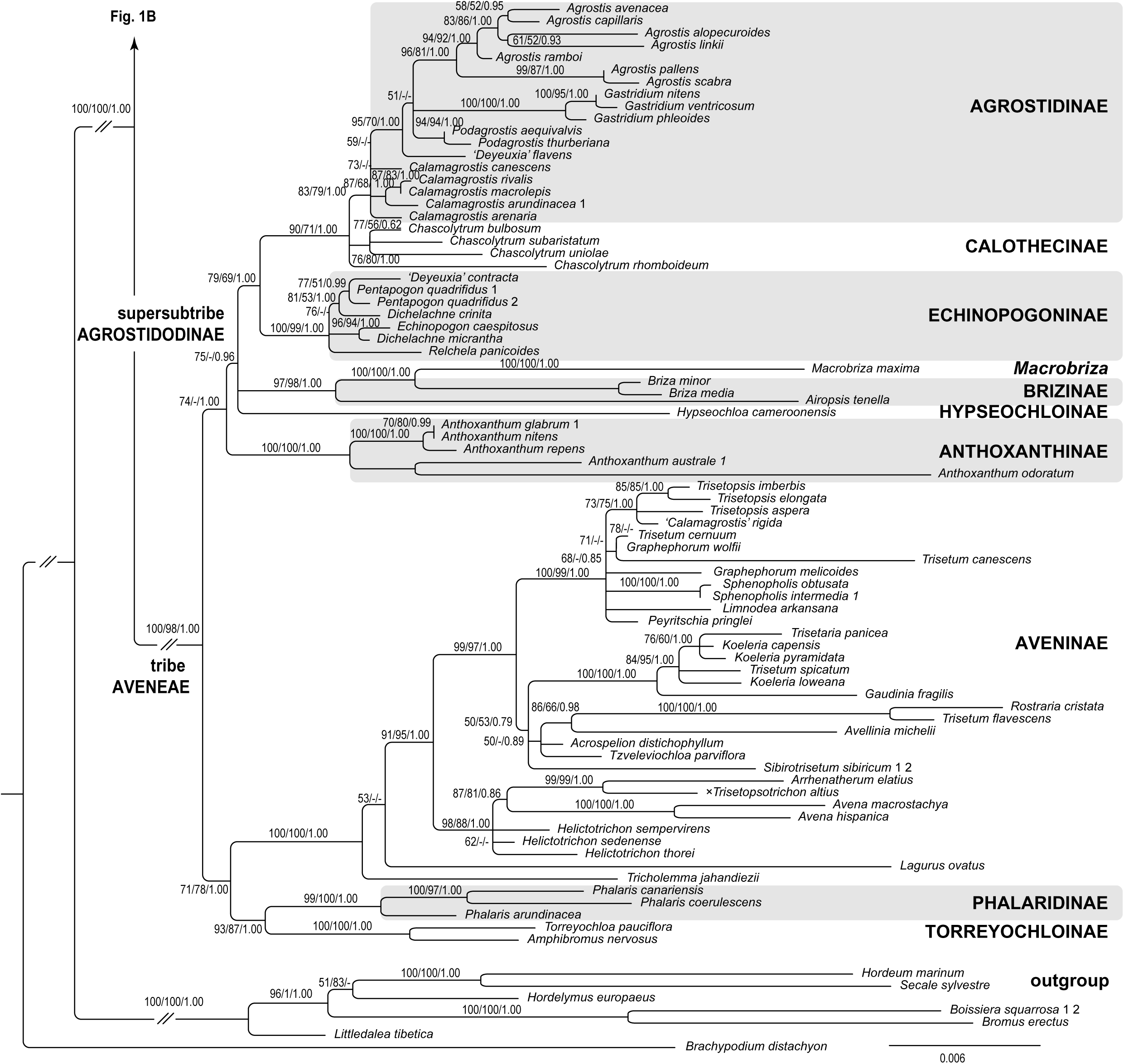
Maximum Likelihood phylogram of Poodae (Aveneae and Poeae) inferred from plastid DNA sequences (*matK* gene–3’*trnK* exon, *trnL–trnF*) with species of Triticodae and Brachypodieae used as outgroup. Maximum Likelihood and Maximum Parsimony bootstrap support values >50% as well as Bayesian posterior probabilities >0.5 are indicated on the branches. Clades with Maximum Likelihood support <50% are collapsed. The subtribes mentioned in the text are labelled on the right-hand side.

Aveneae showed more or less a polytomy of six lineages. Following the recent classification of Soreng & al. (2017) and new names coined in this study (see below *New names and combinations*) these lineages corresponded to subtribes Torreyochloinae and Phalaridinae unified in a common clade (93/87/1.00), Aveninae (100/100/1.00), Anthoxanthinae (100/100/1.00), new subtribe Hypseochloinae (only *Hypseochloa* C.E.Hubb.), Brizinae with *Macrobriza* (Tzvelev) Tzvelev (97/98/1.00), Echinopogoninae (100/99/1.00) and a well-supported common clade (90/71/1.00) of Calothecinae and Agrostidinae.

Within Aveninae, *Lagurus ovatus* L., assigned to monogeneric subtribe Lagurinae by Saarela & al. (2017), and *Tricholemma jahandiezii* (Litard. ex Jahand. & Maire) Röser were opposed to a larger clade (91/95/1.00) formed of supported Aveninae s.str. (98/88/1.00) and a lineage (99/97/1.00) sometimes referred to as separate subtribe Koeleriinae (Quintanar & al., 2007; Saarela & al., 2017; Barberá & al., 2019). ‘*Calamagrostis*’ *rigida* (Kunth) Trin. ex Steud., member of the Central to South American ‘*Calamagrostis*’ Adans. or ‘*Deyeuxia*’ Clarion ex P.Beauv. species, was nested among Aveninae and not Agrostidinae as *Calamagrostis* s.str. and *Deyeuxia* s.str. It was placed in Koeleriinae in agreement with the previous findings (Saarela & al., 2010, 2017; Wölk & Röser, 2014). Monospecific *Limnodea* L.H.Dewey [*L. arkansana* (Nutt.) L.H.Dewey], so far considered either Poinae or Agrostidinae (Kellogg, 2015; Soreng & al., 2017), was nested within Aveninae and likewise in its Koeleriinae lineage. Strongly supported Echinopogoninae encompassed among others (*Dichelachne* Endl., *Echinopogon* P.Beauv., *Pentapogon* R.Br., *Relchela* Steud.) also ‘*Deyeuxia*’ *contracta* (F.Muell. ex Hook.f.) Vickery as representative of Australasian ‘*Deyeuxia’* or ‘*Calamagrostis’*. Altogether, supersubtribe Agrostidodinae (Soreng & al., 2017) encompassing Agrostidinae, Brizinae, Calothecinae, Echinopogoninae and Hypseochloinae received some support (75/-/0.96).

The second main lineage of the plastid DNA tree was the “Poeae chloroplast group 2 (Poeae type)” or tribe Poeae s.str. as suggested in this study. It had a major basal polytomy consisting of Airinae, which received maximum support (100/100/1.00), a large lineage with 81/61/- support and the PPAM clade, an acronym derived from the subtribe names Puccinelliinae (= Coleanthinae), Poinae, Alopecurinae and Miliinae (Gillespie & al., 2008, 2010), supported by 71/67/0.99.

The large lineage with 81/61/- support encompassed Holcinae (100/100/1.00), Aristaveninae (100/98/1.00), Sesleriinae (81/62/0.99), Loliinae (100/99/1.00), Ammochloinae (only *Ammochloa*), Dactylidinae (100/100/1.00), Cynosurinae, Parapholiinae (100/99/1.00) and the new subtribe Helictochloinae (100/100/1.00) with *Helictochloa* Romero Zarco and *Molineriella* Rouy, two genera previously accommodated in Airinae. *Antinoria* Parl., (new monogeneric subtribe Antinoriinae), was closer to *Scolochloa* Link (86/70/0.68). *Dryopoa* Vickery, the second Scolochloinae genus, was placed separate from *Scolochloa* with Sesleriinae (63/50/0.98). Ammochloinae and Dactylidinae were sister clades (97/95/1.00). Cynosurinae with *Cynosurus* species forming a grade and Parapholiinae were placed in a common clade (100/100/1.00). Ammochloinae/Dactylidinae and Cynosurinae/Parapholiinae are acknowledged here as AD and CP clades, altogether as ADCP clade, which had weak support (57/52/0.70).

The PPAM clade split into subtribe Coleanthinae and supersubtribe Poodinae (Soreng & al., 2017), both of which gained strong support (100/99/1.00 and 100/95/1.00, respectively). The latter agrees with the PAM clade, an acronym derived from the subtribe names Poinae, Alopecurinae and Miliinae (Gillespie & al., 2008), or with subtribe Poinae as broadly delineated by Kellogg (2015). It included the subtribes Miliinae (only *Milium*), Phleinae (only *Phleum*) and Poinae (only *Poa*; 86/76/1.00), *Arctopoa* (Griseb.) Prob., monospecific *Avenula* (Dumort.) Dumort. assigned to the monogeneric new subtribe Avenulinae and a considerably supported lineage (84/68/1.00) termed here ABCV clade. It encompassed a large polytomy Alopecurinae, Beckmanniinae, Cinninae, Ventenatinae (85/60/1.00), along with *Brizochloa* V.Jirásek & Chrtek (monogeneric new subtribe Brizochloinae) and a number of genera not classed as to subtribe, including *Arctohyalopoa* Röser & Tkach, a new monospecific genus harboring former *Hyalopoa lanatiflora* (Roshev.) Tzvelev. Moreover, Alopecurinae, Beckmanniinae and Cinninae did not resolve as monophyletic. Only a sister relation of Alopecurinae genera *Alopecurus* L. and *Cornucopiae* L. was strongly supported (100/100/1.00), whereas *Beckmannia* Host. and *Pholiurus* Trin. (Beckmanniinae) obtained weak support as sister (59/57/0.98). The DAD clade, an acronym originally derived from *Dupontia*, *Arctophila* and *Dupontiopsis* (Soreng & al., 2015a), in this study encompassing *Dupontia* including *Arctophila*, *Dupontiopsis* and new *Arctohyalopoa*, was obvious within the ABCV clade but with low support (72/-/-), whereas the HSAQN clade (Soreng & al., 2015b; Gillespie & al., 2010; Kellogg, 2015) was unresolved.

### Nuclear DNA analysis

The nuclear ITS DNA sequence dataset for 215 taxa included 673 and the ETS DNA dataset for 200 taxa included 1,135 aligned positions, respectively. The combined data matrix of two nuclear DNA markers for 218 taxa included a total of 1,808 aligned positions, of which 1,093 were variable (ITS: 383, ETS: 710) and 863 parsimony-informative (ITS: 320, ETS: 543).

Poodae was supported by the nr ITS and ETS sequence data as monophyletic (100/88/1.00). The tree backbone consisted of a polytomy of six clades (Fig. 2) comprising Antinoriinae, Helictochloinae (100/100/1.00), Aristaveninae (100/96/1.00) and a supported lineage (100/79/1.00) harboring Loliinae (70/-/1.00) and the ADCP clade (69/-/1.00). This well-supported lineage of the latter two elements, unresolved in the plastid DNA tree, represented supersubtribe Loliodinae (Soreng & al., 2017). The remaining two clades of the backbone polytomy were the PPAM clade (95/74/1.00), and a large clade with 85/-/1.00 support. The PPAM clade consisted of Miliinae, a common lineage of Phleinae with Poinae (84/-/0.99) that had not been resolved in the plastid DNA tree, Coleanthinae (68/-/0.98), Avenulinae and a lineage termed ABCV+A clade (83/67/1.00), which corresponded to the ABCV clade in the plastid tree complemented by *Arctopoa*. Supersubtribe Poodinae (∼PAM clade) was not resolved within the PPAM clade. The HSAQN clade was well-defined (97/97/1.00) within the ABCV+A clade. Sister relations of *Alopecurus* and *Cornucopiae* (Alopecurinae) and of *Beckmannia* and *Pholiurus* (Beckmanniinae) were supported (95/97/1.00 and 100/100/1.00, respectively).

**Fig. 2.**
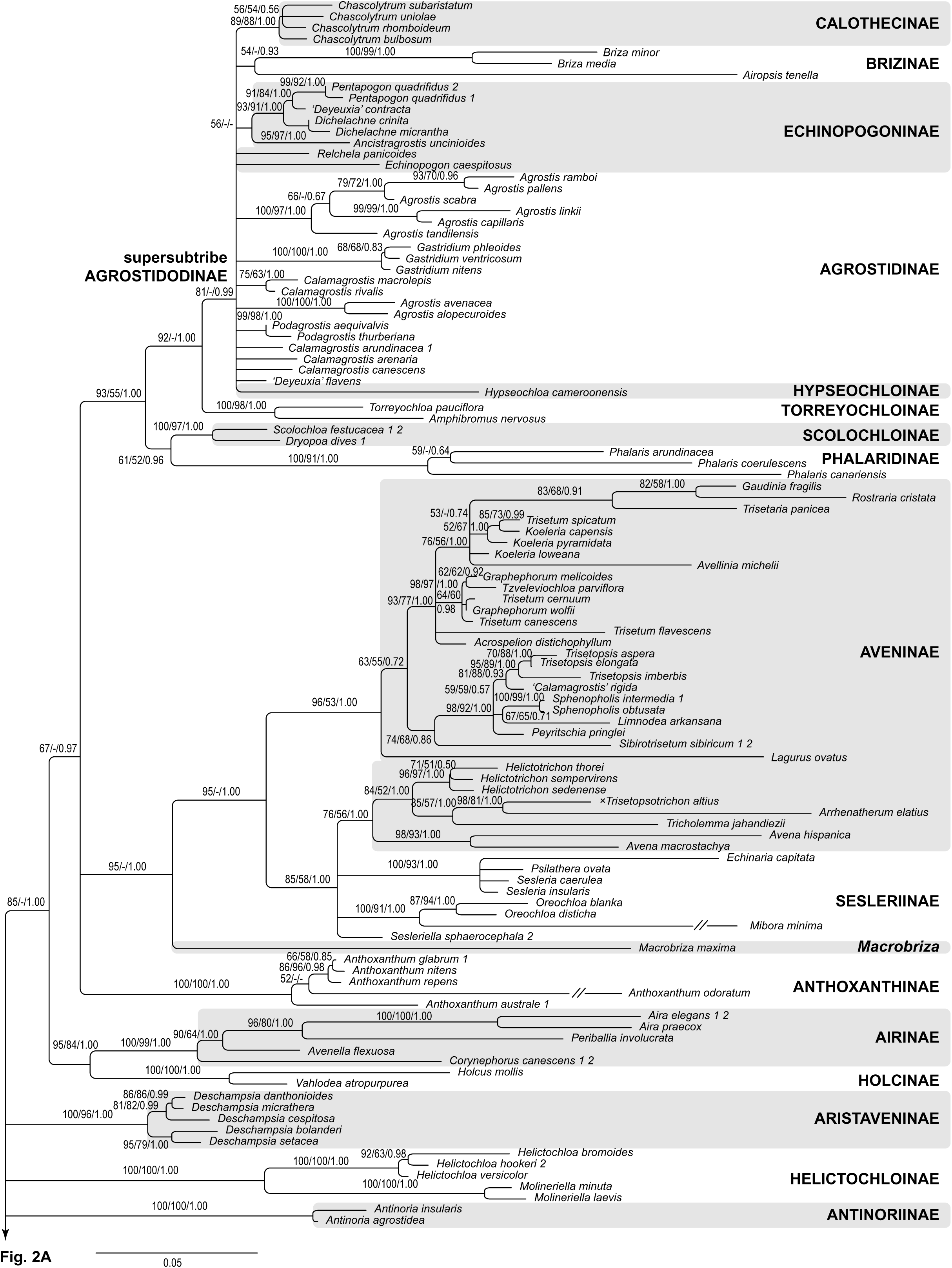
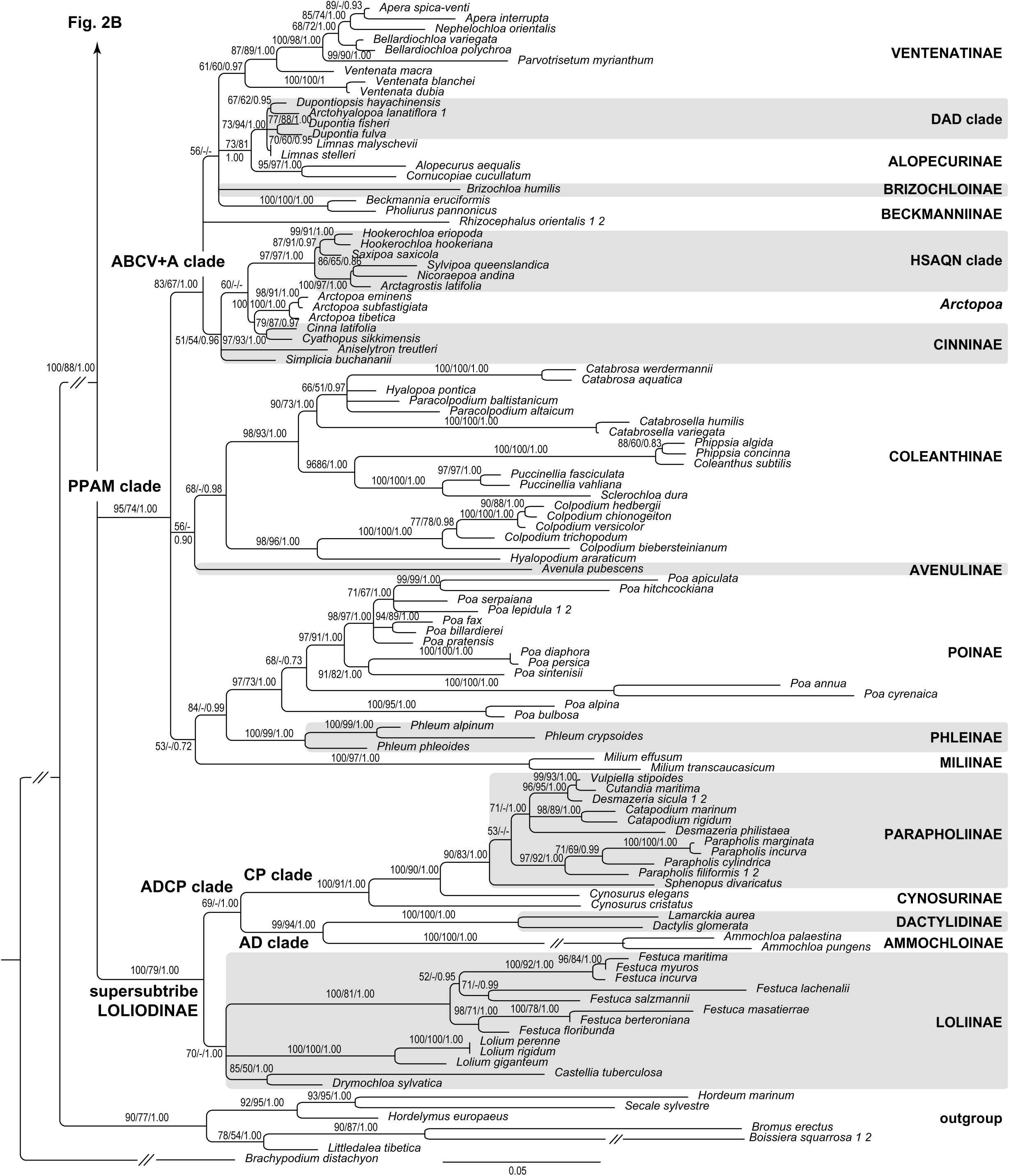
Maximum Likelihood phylogram of Poodae (Aveneae and Poeae) inferred from nr (ITS, ETS) DNA sequences with species of Triticodae and Brachypodieae as outgroup. Maximum Likelihood and Maximum Parsimony bootstrap support values >50% as well as Bayesian posterior probabilities >0.5 are indicated on the branches. Clades with Maximum Likelihood support <50% are collapsed. The subtribes mentioned in the text are labelled on the right-hand side.

The large clade with 85/-/1.00 support of the backbone polytomy showed a partly well-supported internal structure. It encompassed Holcinae (100/100/1.00) and Airinae (100/99/1.00) as supported sister clades (95/84/1.00), Anthoxanthinae, a lineage (95/-/1.00) of *Macrobriza*, Sesleriinae and Aveninae, which had not been encountered in the plastid DNA tree, and a clade supported by 93/55/1.00. It was formed by Scolochloinae with *Dryopoa* and *Scolochloa* (100/97/1.00), Phalaridinae, Torreyochloinae (100/98/1.00) and supersubtribe Agrostidodinae (81/-/0.99), which contained Hypseochloinae, Brizinae (54/-/0.93), Calothecinae (89/88/1.00) as well as species and small clades of Agrostidinae and Echinopogoninae in a polytomy. *Ancistragrostis* S.T.Blake (available only ITS) was placed with low support along with Echinopogoninae, which encompassed also ‘*Deyeuxia*’ *contracta* (93/91/1.00).

Aveninae segregated into two different lineages similar to the ones encountered in the plastid DNA tree, except for the position of *Tricholemma* (Röser) Röser and *Lagurus* L. One of the lineages, Aveninae s.str. (76/56/1.00), assembled with non-monophyletic Sesleriinae in a common lineage (85/58/1.00), whereas the other corresponded to the Koeleriinae lineage (96/53/1.00). It had *Lagurus* (Lagurinae) as early branching genus and encompassed ‘*Calamagrostis*’ *rigida*.

### Analysis of the combined DNA dataset

Following the rationale outlined in Material and Methods we analyzed also a concatenated dataset of plastid and nuclear DNA sequence data to evaluate which of the clades retrieved by the individual analyses kept stable or eventually became even better supported and which clades became less supported or collapsed.

The combined data matrix of all plastid and nuclear DNA sequences for 218 taxa included a total of 5,736 aligned positions of which 2,564 were variable and 1,770 parsimony-informative.

The backbone of the Poodae tree showed the same deep dichotomy as the plastid DNA tree, which reflected the Aveneae and Poeae s.str., whereas further tree resolution was overall low. Fig. 3 gives a simplified overview of the tree as a cladogram, the detailed phylogram is shown in suppl. Fig. S1. Within the clade of Aveneae (100/67/1.00)), a series of clades arranged in a polytomy was found. Anthoxanthinae (100/100/1.00), Aveninae (100/92/1.00), Torreyochloinae (100/100/1.00) and Phalaridinae (100/99/1.00) were well-supported, to a lesser extent Brizinae excluding *Macrobriza* (75/-/1.00) and Calothecinae (82/90/1.00). Agrostidinae and Echinopogoninae did not resolve as monophyletic, respectively, but were part of a polytomy. Supersubtribe Agrostidodinae under exclusion of *Macrobriza* was slightly supported (62/-/1.00). Aveninae showed an internal structure of two main clades (Aveninae s.str., Koeleriinae) with *Tricholemma* sister to one of these (93/90/1.00) and *Lagurus* to the other (69/61/1.00).

**Fig. 3.**
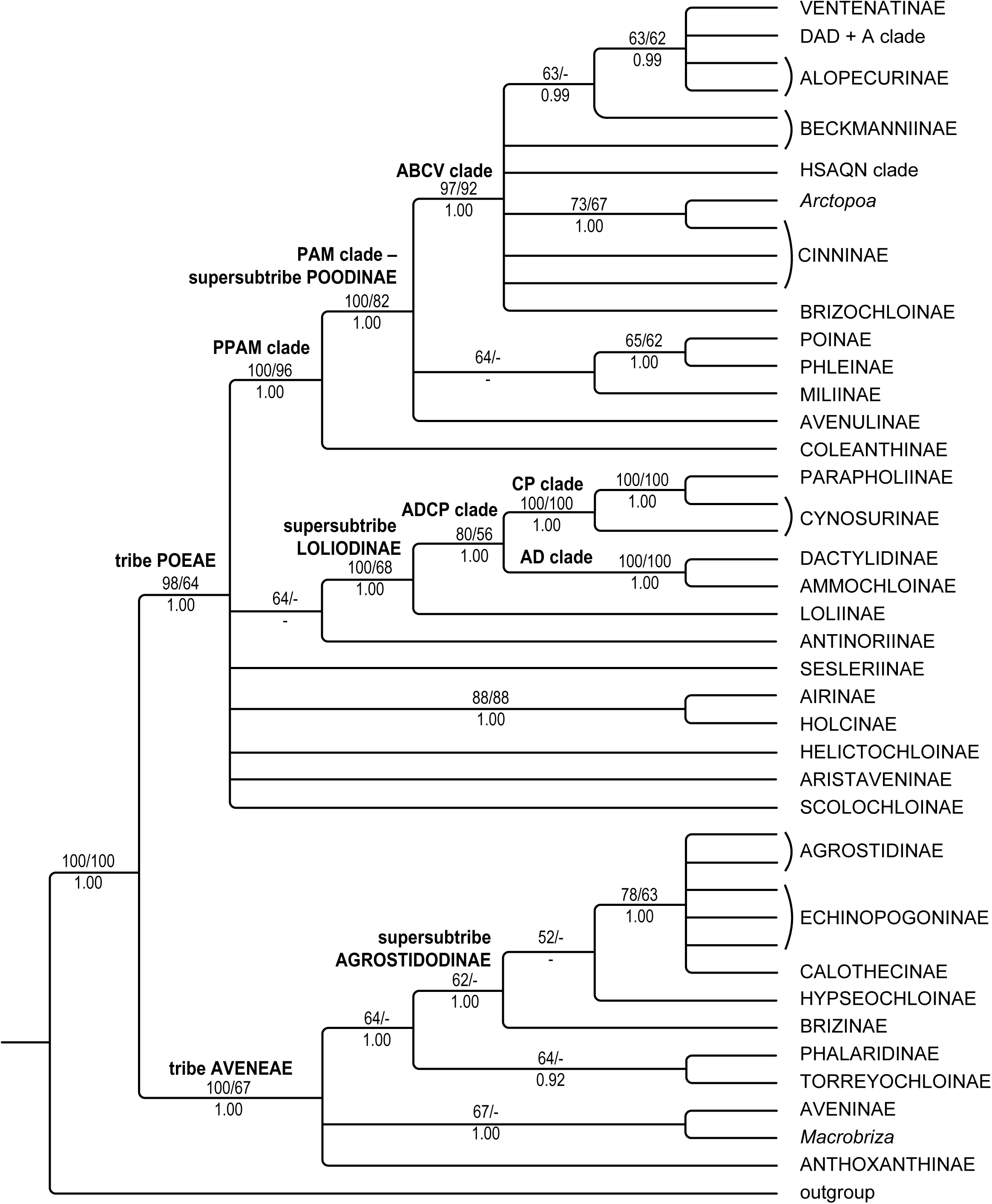
Overview of the Maximum Likelihood cladogram of Poodae (Aveneae and Poeae) inferred from the concatenated matrix of plastid (*matK* gene–3’*trnK* exon, *trnL–trnF*) and nr (ITS, ETS) DNA sequences with species of Triticodae and Brachypodieae as outgroup. Maximum Likelihood and Maximum Parsimony bootstrap support values >50% as well as Bayesian posterior probabilities >0.5 are indicated on the branches. Clades with Maximum Likelihood support <50% are collapsed. The expanded tree is displayed in Supplementary Fig. S1.

Within the clade of Poeae (98/64/1.00), several highly supported lineages were resolved but were part of a polytomy: Scolochloinae, Aristaveninae, Helictochloinae (100/100/1.00 each), Sesleriinae (100/99/1.00), Holcinae (100/100/1.00) unified with Airinae (100/100/1.00) in a common clade (88/88/1.00), a low-support clade of Antinoriinae with supersubtribe Loliodinae (100/68/1.00) containing Loliinae (99/67/1.00) and the ADCP clade (80/56/1.00) and, finally, the PPAM clade (100/96/1.00). The latter encompassed Coleanthinae (100/99/1.00) and supersubtribe Poodinae (∼PAM clade; 100/82/1.00), in which also Avenulinae was placed. Within Poodinae, the ABCV+A (97/97/92/1.00), the HSAQN (99/98/1.00) and the DAD clade (96/91/1.00) were supported.

## DISCUSSION

### Molecular phylogenetics

#### Comparison of the plastid and nuclear DNA trees

Both phylogenetic trees agreed widely in the resolution of minor clades, whose support values were frequently comparatively high (Figs. 1, 2, 4). The larger clades, by contrast, corresponded only partly and disagreed strikingly in some instances.

Concordant groupings were (1) supersubtribe Agrostidodinae, which was resolved in both individual analyses although excluding *Macrobriza* in the nuclear tree, in which it was sister to a clade of Aveninae and Sesleriinae; (2) the PPAM clade; (3) the ADCP clade. There were (4) many congruent clades, which corresponded to subtribes, for example, Phalaridinae, Torreyochloinae, Anthoxanthinae, Holcinae, Aristaveninae, Loliinae, Ammochloinae, Dactylidinae, Parapholiinae, Coleanthinae, Phleinae, Miliinae, Poinae and Ventenatinae. (5) Former subtribe Airinae (Airinae s.l.) was non-monophyletic in both analyses but its subgroups were resolved and congruent (Airinae, Antinoriinae, Helictochloinae).

Several clades were monophyletic in one of the individual plastid and nuclear DNA analyses, whereas they were unresolved in the other, appearing as a polytomy or a grade. We consider this not as severe conflict. Supersubtribe Loliodinae was clearly monophyletic in the nuclear DNA tree but formed a polytomy with several other lineages in the plastid DNA tree. Calothecinae and the DAD clade were likewise monophyletic in the nuclear but paraphyletic in the plastid DNA tree. Conversely, supersubtribe Poodinae (∼PAM clade) including *Avenula* as well as subtribes Echinopogoninae, Agrostidinae and Sesleriinae were clearly monophyletic in the plastid DNA tree but form polytomies with other lineages in the nuclear DNA tree.

Discordant groupings occurred starting with the backbone of the trees since the bifurcation of the plastid DNA tree into Aveneae and Poeae s.str. was not reflected in the nuclear tree, which represented a polytomy. Sesleriinae from the Poeae lineage of the plastid were placed along with Aveninae in the nuclear DNA tree (Quintanar & al., 2007; Saarela & al., 2017). Subtribe Holcinae and Airinae as part of Poeae in the plastid DNA tree were placed in the nuclear DNA tree close to subtribes of Aveneae such as Aveninae, Agrostidinae, etc. The same pattern was encountered in Scolochloinae as belonging to Poeae (plastid) but nested (nuclear) in a common clade with subtribes of Aveneae such as Phalaridinae, Torreyochloinae, Echinopogoninae, Agrostidinae etc. A number of further subtribes showed different affiliations depending on the individual tree: Phalaridinae and Torreyochloinae were sister in the plastid but not in the nuclear DNA analyses (Saarela & al., 2017). Subtribe Aveninae was monophyletic in the plastid DNA tree but disintegrated in the nuclear tree into two lineages, one of which (Aveninae s.str. with *Tricholemma*) aggregated with the taxa of Sesleriinae, whereas the other (Koeleriinae with *Lagurus*) was separate.

*Macrobriza* as member of monophyletic Brizinae in the plastid DNA tree was nested along with Sesleriinae and Aveninae in the nuclear tree. Furthermore, *Arctopoa* was placed in a clade along with Poinae in the plastid but within the ABCV+A clade in the nuclear tree (Gillespie & al., 2008, 2010; Nosov & al., 2015, 2019). Many further genera showed switching positions within their respective subtribes, for example, within Aveninae, Coleanthinae, Loliinae and Sesleriinae (see below *Reticulations within major lineages*).

#### Tree of the combined plastid and nuclear DNA dataset

This tree obtained from the concatenated dataset all in all combined features of the individual plastid and nuclear DNA trees. Lineages that were retrieved in both individual analyses were present also in the combined tree, for example, the PPAM clade, supersubtribe Agrostidodinae (except for *Macrobriza*), the ADCP, AD and CP clades (Fig. 4, suppl. Fig. S1). Also many subtribes were recovered in the combined analyses such as Anthoxanthinae, Torreyochloinae and Phalaridinae, Aristaveninae, Holcinae, Loliinae, Dactylidinae, Ammochloinae, Parapholiinae, Coleanthinae, Miliinae, Phleinae, Poinae, Ventenatinae and the monophyletic subdivisions of former Airinae s.l., namely Airinae, Antinoriinae and Helictochloinae. Occasionally, the clade support values of concordant lineages were higher in the analysis of the combined dataset than in the individual analyses, for example, for the PPAM clade (combined 100/96/1.00, plastid 71/67/0.99, nuclear 95/74/1.00), the ABCV(+A) clade (combined 97/92/1.00, plastid 84/68/1.00, nuclear 83/67/1.00) or the ADCP clade (combined 80/56/1.00, plastid 57/52/0.70, nuclear 69/-/1.00).

**Fig. 4.**
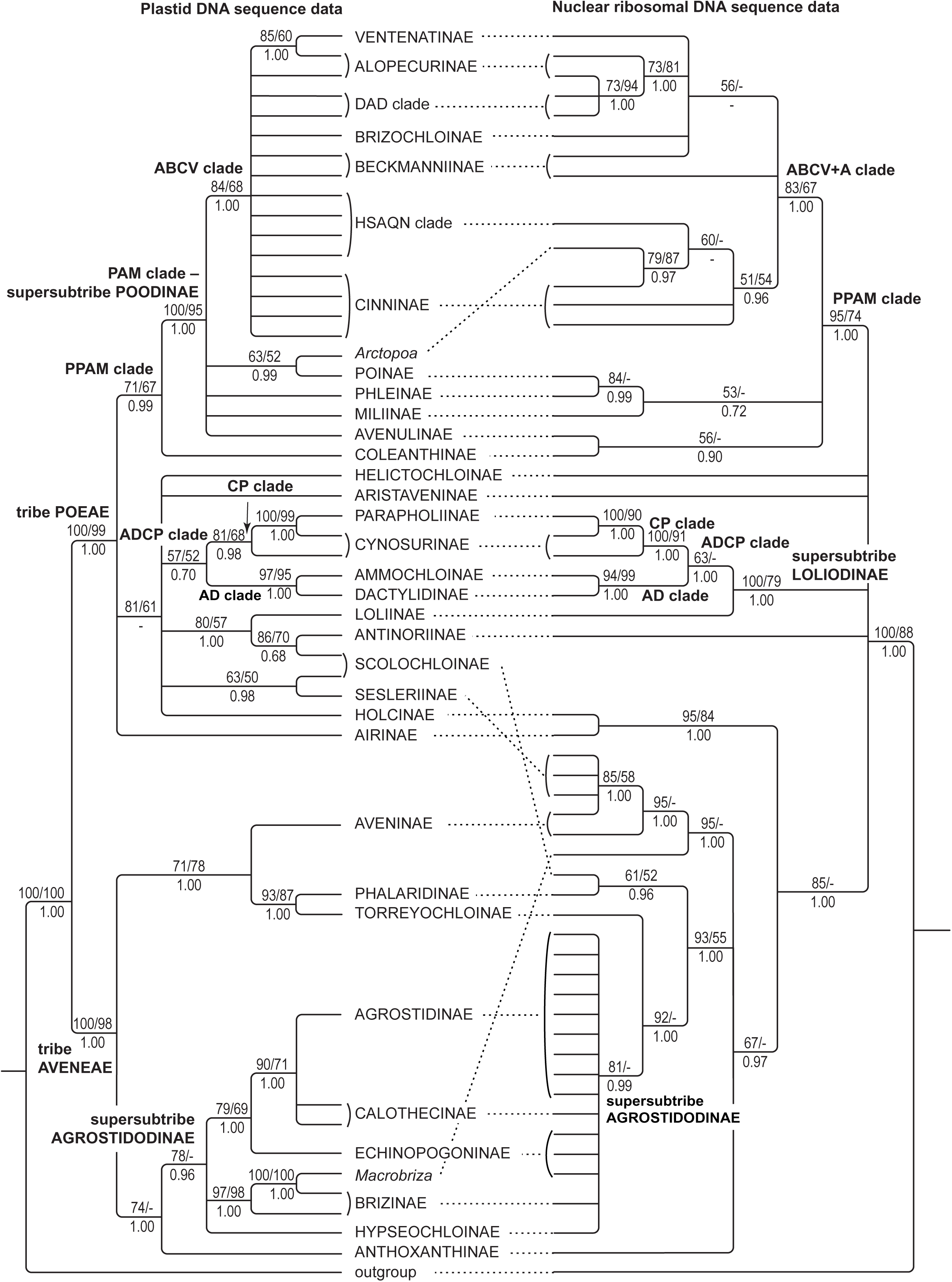
Comparison of simplified Maximum Likelihood cladograms of Poodae (Aveneae and Poeae) inferred from plastid (*matK* gene–3’*trnK* exon, *trnL–trnF*) and nr (ITS, ETS) DNA sequences with species of Triticodae and Brachypodieae as outgroup. Maximum Likelihood and Maximum Parsimony bootstrap support values >50% as well as Bayesian posterior probabilities >0.5 are indicated on the branches. Clades with Maximum Likelihood support <50% are collapsed. The expanded trees are displayed in Figs. 1 and 2, respectively.

The combined tree contained most well-resolved groups from the individual trees, even if they were supported only in one of them. It followed this way the main dichotomy of Aveneae and Poeae s.str. observed in the plastid tree and resolved supersubtribe Poodinae and subtribe Agrostidinae, which were likewise supported only in the plastid DNA tree. Conversely, supersubtribe Loliodinae, the ABCV+A and the HSAQN clade, subtribes Calothecinae, Scolochloinae and the Koeleriinae lineage combined with *Lagurus* found within Aveninae were present in the combined tree although they were supported only in the nuclear but not the plastid DNA tree.

Lineages with a discordant grouping in the individual analyses followed one of these placements in the combined tree. Sesleriinae were nested within the Poeae clade as in plastid DNA tree and not together with Aveninae as in the nuclear DNA tree. Torreyochloinae and Phalaridinae built a clade sister to supersubtribe Agrostidodinae as similarly encountered in the nuclear DNA tree, whereas they were sister to Aveninae in the plastid DNA tree. Subtribe Aveninae resolved monophyletic as in the plastid DNA tree, however, the placement of *Lagurus* and *Tricholemma* was different. *Lagurus* was sister to the Koeleriinae lineage as in the nuclear DNA tree. *Tricholemma* belonged to the Aveninae s.str., although its position was new relative to the nuclear DNA tree. Subtribe Scolochloinae grouped within Poeae as in the plastid DNA tree, whereas it was affiliated with supersubtribe Agrostidodinae, namely subtribes Torreyochloinae and Phalaridinae, in the nuclear DNA tree. Finally, *Macrobriza* was sister to the lineage of Aveninae and Sesleriinae in the nuclear but was placed within Brizinae in the plastid DNA tree.

#### Hybrid origin of major lineages or subtribes and genera derived from hybridization between them

The examples of discordant grouping are best explained by ‘chloroplast capture’, which means hybridization (Rieseberg & Soltis, 1991). Some lineages and genera have seemingly reticulate origin documented by the incongruent placement in the plastid and nuclear trees (Figs. 1, 2, 4).

1. The whole lineage of Sesleriinae had one ancestor with Poeae s.str. plastid DNA (Figs. 1, 4). Due to the usually maternal inheritance of plastids in angiosperms, this ancestor was supposedly the maternal parent. The paternal parent inherited the Aveninae-like rDNA (Figs. 2, 4). This can be stated even more precisely because it was an Aveninae s.str.- and not Koeleriinae-like parent. Maternal rDNA seemingly is no longer present in Sesleriinae or at least was not detected by our approach using direct sequencing of PCR products, presumably due to sequence homogenization of this repetitive rDNA in favor of one parental copy type, a well-documented process of unidirectional loss (Winterfeld & al., 2009, 2012; Kotseruba & al., 2010; Wölk & al., 2015; Tkach & al., 2019).
2. Subtribes Phalaridinae and Torreyochloinae had a maternal ancestor with Aveninae-like plastid DNA, whereas the paternal parent was close to supersubtribe Agrostidodinae (Figs. 1, 2, 4).
3. Scolochloinae had a maternal parent inheriting the plastid DNA from Poeae s.str., most likely from outside the PPAM clade, namely from relatives of Loliinae and Sesleriinae (Figs. 1, 4). The paternal parent as indicated by the nuclear rDNA was distantly related and was close to supersubtribe Agrostidodinae such as seen in Phalaridinae and Torreyochloinae (Figs. 2, 4).
4. The position of Phleinae is intriguing, because this subtribe is sister lineage to Poinae in the nuclear DNA analyses (84/-/0.99), whereas in the plastid DNA analyses it is part of a polytomy with Miliinae, Poinae, the ABCV clade, *Avenula* and *Arctopoa* within the strongly supported supersubtribe Poodinae (∼PAM clade; Figs. 1, 2, 4). The placements of Phleinae might actually bear witness of a further instance of reticulation.
5. *Macrobriza* and *Arctopoa* are examples of genera with hybrid origin. Monospecific *Macrobriza* had a maternal parent with Brizinae plastid DNA and a paternal parent with Aveninae-/Sesleriinae-like rDNA (Figs. 1, 2, 4). *Arctopoa* had a maternal parent (plastid donor) related with, or from, Poinae, whereas its paternal parent inheriting its rDNA belonged to the ABCV clade in accordance with Gillespie & al. (2008, 2010), while the maternal rDNA is no longer detectable, at least by our direct sequencing of PCR products. Further examples of genera originating from hybridization across major lineages were discussed also in the instances of *Avenula*, *Helictochloa* and *Aniselytron* Merr. In the latter, a strongly divergent, *Poa* L.-like ITS copy was found in addition to the regular type, pointing to either hybrid origin of *Aniselytron* or recent hybridization with *Poa* (Soreng & Davis, 2000; Gillespie & al., 2008, 2010; Soreng & al., 2017). Our results, however, indicate a largely concordant placement of these genera in the plastid and nuclear DNA trees, respectively. Only in monospecific *Avenula*, which was consistently placed in all our analyses within the PPAM clade, there are slight differences but seem to be too small to corroborate hybrid origin of *Avenula* (see below *PAM clade…*).

#### Reticulations within major lineages

1. Within Aveninae, *Tricholemma* and *Lagurus*, with a plastid DNA seemingly characteristic of early-branching Aveninae as a whole (Figs. 1, 5; Wölk & Röser, 2017), have rDNA sequences with characteristics of either Aveninae s.str. or the lineage of Koeleriinae. In the nuclear DNA tree, *Lagurus* was sister to the remaining genera of the latter and represents an early-branching offspring (Figs. 2, 5). *Tricholemma* was nested amidst the taxa of Aveninae s.str. Within the Koeleriinae lineage, there were several further instances of non-concordant placements of taxa, the most remarkable being that of American *Trisetum* Pers. species and *Graphephorum* Desv. (Wölk & Röser, 2017; see below *Aveninae*).
2. *Sesleria* Scop. (Sesleriinae) was sister to *Sesleriella* Deyl according to the plastid DNA data (Figs. 1, 6), whereas the nuclear rDNA of *Sesleria* points to a close relation with *Psilathera* Link and *Echinaria* Desf. (Figs. 2, 6; see below *Sesleriinae and Scolochloinae*). As suggested by Kuzmanović & al. (2017), *Sesleria* originated most likely from hybridization between a maternal *Sesleriella*- and a paternal *Psilathera*-like ancestor. The monospecific genus *Echinaria* was unlikely to be involved in the origin of *Sesleria*, because it is a short-lived annual of the Mediterranean lowlands in contrast to the other genera in question, which are characteristic perennials of mountainous habitats.
3. The new Coleanthinae genus *Hyalopodium* Röser & Tkach, gen. nov., comprises only *H. araraticum* (Lipsky) Röser & Tkach, comb. nov. [≡ *Colpodium araraticum* (Lipsky) Woronow ex Grossh.]. With respect to the nuclear rDNA, *Hyalopodium* largely agreed with *Colpodium* Trin. (Figs. 1, 7; Rodionov & al., 2008; Kim & al., 2009), whereas it shared plastid DNA characteristics with *Hyalopoa* (Tzvelev) Tzvelev [*H. pontica* (Balansa) Tzvelev] and *Paracolpodium* (Tzvelev) Tzvelev [*P. altaicum* (Trin.) Tzvelev, *P. baltistanicum* Dickoré; Figs. 2, 7], indicative of hybrid background. The incongruent tree position of the sister genera *Puccinellia* Parl. and *Sclerochloa* P.Beauv. also points to hybrid origin because they clustered with *Catabrosa* P.Beauv. and *Catabrosella* (Tzvelev) Tzvelev in the plastid but with *Coleanthus* Seidel ex Roem. & Schult. and *Phippsia* (Trin.) R.Br. in the nuclear DNA tree (Figs. 1, 2, 7; Schneider & al., 2009).
4. The monospecific genus *Castellia* Tineo [*C. tuberculosa* (Moris) Bor] of subtribe Loliinae presumably originated from a *Festuca* Tourn. ex L.-like maternal ancestor providing the plastid and a paternal ancestor related to *Drymochloa* Holub (Figs. 1, 2; see below *Loliinae*).

**Fig. 5.**
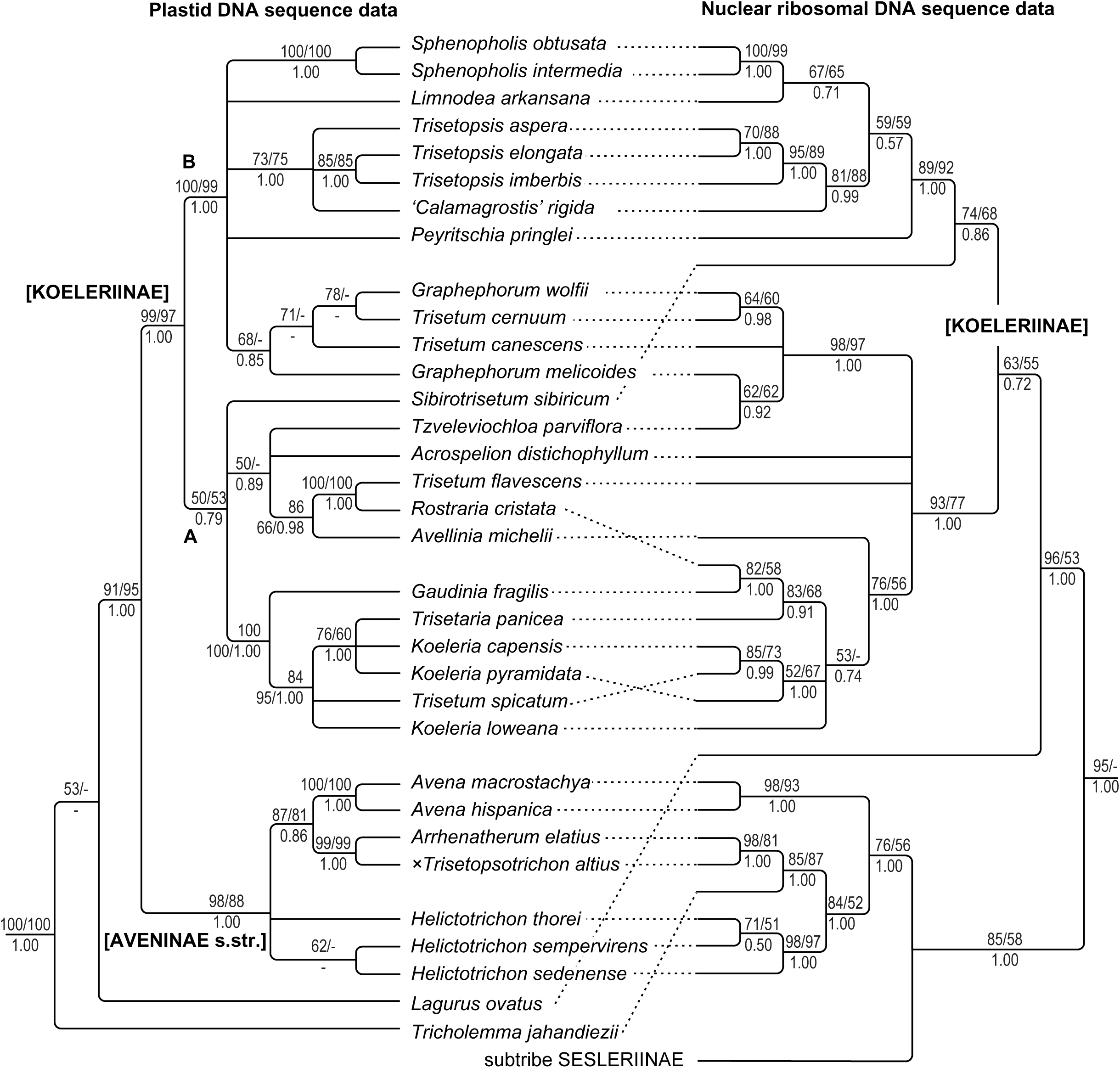
Comparison of Maximum Likelihood cladograms for the genera of subtribe Aveninae inferred from plastid (*matK* gene–3’*trnK* exon, *trnL–trnF*) and nuclear (ITS, ETS) DNA sequences. Maximum Likelihood and Maximum Parsimony bootstrap support values >50% as well as Bayesian posterior probabilities >0.5 are indicated on the branches. Clades with Maximum Likelihood support <50% are collapsed. Aveninae is sometimes split into Koeleriinae, clades A and B, and Aveninae s.str. (square brackets).

**Fig. 6.**
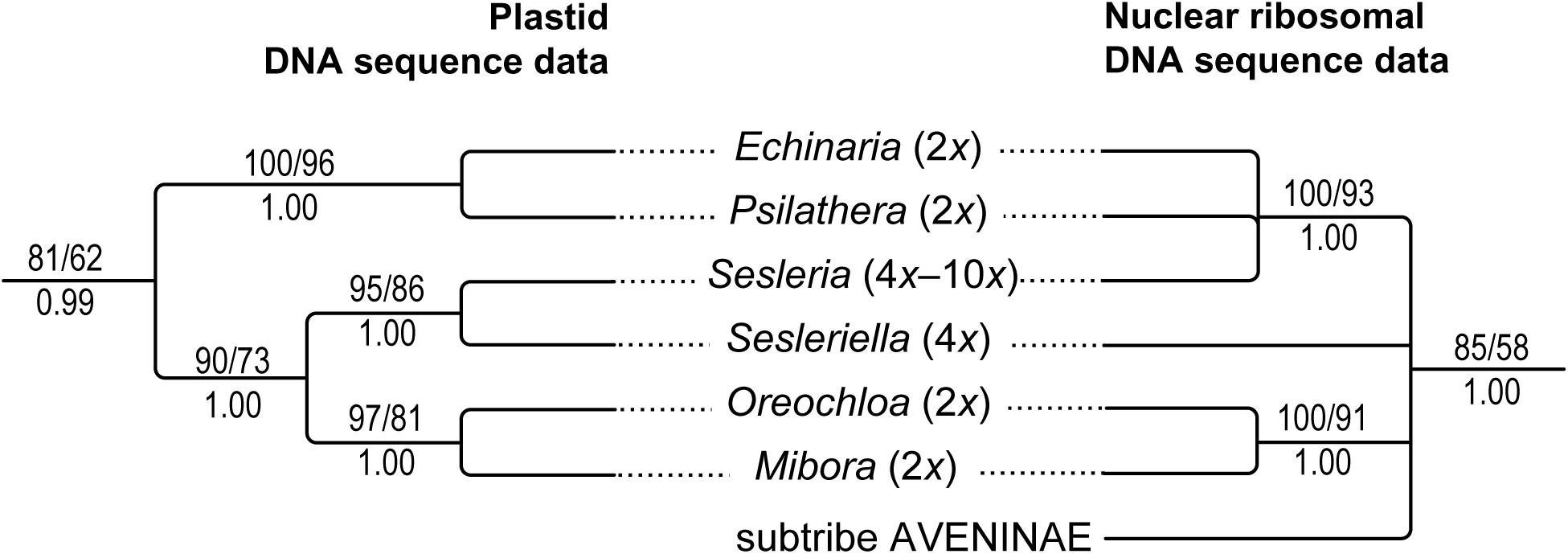
Comparison of Maximum Likelihood cladograms for the genera of subtribe Sesleriinae inferred from plastid (*matK* gene–3’*trnK* exon, *trnL–trnF*) and nr (ITS, ETS) DNA sequences. Maximum Likelihood and Maximum Parsimony bootstrap support values >50% as well as Bayesian posterior probabilities >0.5 are indicated on the branches. Clades with Maximum Likelihood support <50% are collapsed.

**Fig. 7.**
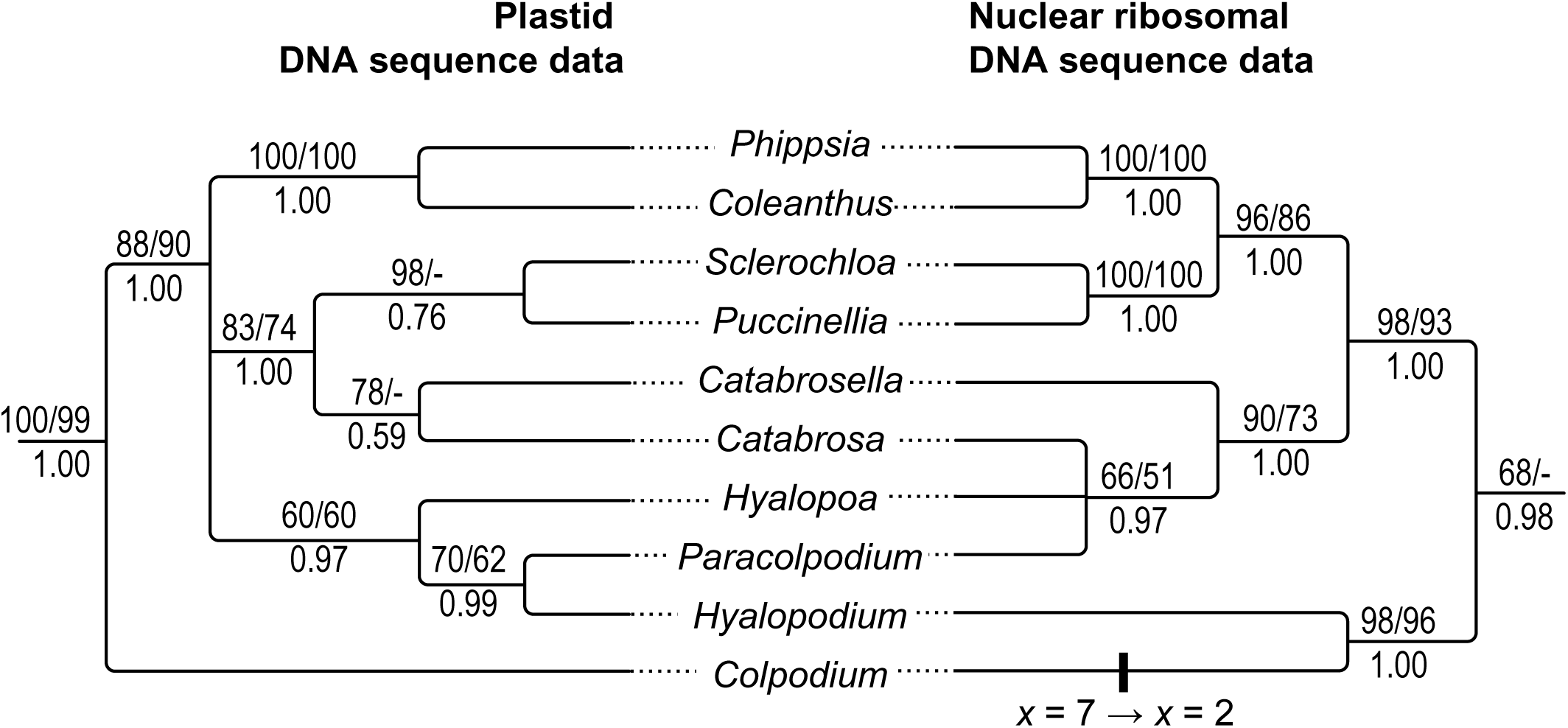
Comparison of Maximum Likelihood cladograms for the genera of subtribe Coleanthinae inferred from plastid (*matK* gene–3’*trnK* exon, *trnL–trnF*) and nr (ITS, ETS) DNA sequences. Maximum Likelihood and Maximum Parsimony bootstrap support values >50% Bayesian posterior probabilities >0.5 are indicated on the branches. Clades with Maximum Likelihood support <50% are collapsed.

### Circumscription of lineages or genera

Our molecular phylogenetic data support the circumscription of many lineages that have been recognized already previously. However, due to the inclusion of several taxa that have not been sampled before and the sampling strategy including a comparatively broad representative set of taxa used for both plastid and nuclear DNA sequence data some re-arrangements and emendations are required.

#### Anthoxanthinae, Torreyochloinae and Phalaridinae

Cumarin-scented subtribe Anthoxanthinae was corroborated as clearly monophyletic and distinct from scentless Phalaridinae (Figs. 1, 2, 4) as suggested by several previous molecular phylogenetic studies (Döring & al., 2007; Quintanar & al., 2007; Döring, 2009; Saarela & al., 2015, 2017; Rodionov & al., 2017; Orton & al., 2019). Species traditionally assigned to *Hierochloe* R.Br. [including the type *H. odorata* (L.) P.Beauv. = *Anthoxanthum nitens* (Weber) Y.Schouten & Veldkamp], namely *H. australis* (Schrad.) Roem. & Schult. [≡ *A. australe* (Schrad.) Veldkamp], *H. glabra* Trin. [≡ *A. glabrum* (Trin.) Veldkamp], *H. pauciflora* R.Br. (= *A. arcticum* Veldkamp), *H. redolens* (Vahl) Roem. & Schult. [≡ *A. redolens* (Vahl) P.Royen] and *H. repens* [≡ *A. repens* (Host) Veldkamp; not all shown in Figs. 1, 2], were not consistently separated from *Anthoxanthum* L.: *A. odoratum* (type of *Anthoxanthum*) was sister to traditional *H. australis* (plastid DNA tree) or was placed between species of traditional *Hierochloe* (nuclear DNA tree). The peculiar tree position of *A. australe*, a species not sampled by Pimentel & al. (2013), agreed with that obtained by Rodionov & al. (2017) on ITS. All in all, the findings support to assign preliminarily the species of *Hierochloe* to *Anthoxanthum* (Schouten & Veldkamp, 1985) as suggested also by Pimentel & al. (2013: 1025) in view of the intermediate floral characters of *A.* sect. *Ataxia* (R.Br.) Stapf between typical *Anthoxanthum* and *Hierochloe* (Connor, 2012).

Subtribe Torreyochloinae consists of south hemispheric *Amphibromus* Nees and North American/East Asian *Torreyochloa* Church. They shared a plastid DNA type with *Phalaris* L. (Döring, 2009; Saarela & al., 2015; Orton & al., 2019), the only genus of holarctic Phalaridinae, but were more distant to each other in nuclear ITS since Torreyochloinae were supported sister to supersubtribe Agrostidodinae, whereas Phalaridinae were closer to Scolochloinae (Figs. 1, 2, 4). Interestingly, the three subtribes share eco-morphological characteristics because *Amphibromus, Torreyochloa*, *Phalaris*, *Scolochloa* and *Dryopoa* have rather tall, sometimes reed-like perennial species, which prefer aquatic habitats or wet mountain forests (*Dryopoa*), except for the annuals of *Phalaris*, some of which are adapted to seasonally dry Mediterranean-type climate (Baldini, 1995).

#### Aveninae

This large lineage encompasses two main subgroups, namely Aveninae s.str. and the Koeleriinae lineage. Additionally, there are two somewhat isolated genera with hybrid background (see above *Reticulations within major lineages*), namely monospecific *Lagurus* (type *L. ovatus*) and *Tricholemma* (two species; type *T. jahandiezii*; Röser, 1989, 1996; Röser & al., 2009; Gabriel & al., 2019), which makes a separation of Aveninae s.str. and Koeleriinae as distinct subtribes not straightforward (Figs. 1, 2, 5), even if *Lagurus* is accommodated under a monogeneric subtribe Lagurinae (Saarela & al., 2017). For this reason we argue for summarising all of them under a single subtribe, i.e., Aveneae, in the classification below.

The taxa of Aveninae have several flowers per spikelet, but there are exceptions with only a single flower such as *Lagurus*, *Limnodea* or Mexican to South American members of ‘*Calamagrostis*’ or ‘*Deyeuxia*’, for which the genus name *Cinnagrostis* Grieseb. recently has been suggested (Soreng & al., 2017; see Saarela & al., 2017; Barberá & al., 2019). The phylogenetic trees of the Aveninae s.str. showed a rather narrowly delineated genus *Helictotrichon* Besser after exclusion of *Tricholemma*, Subsaharo-African to Southeast Asian *Trisetopsis* Röser & A.Wölk [type *T. elongata* (Hochst. ex A.Rich.) Röser & A.Wölk] and East Asian *Tzveleviochloa* Röser & A.Wölk [type *T. parviflora* (Hook.f.) Röser & A.Wölk], which are members of the Aveninae but belong to the Koeleriinae lineage. Excluded from *Helictotrichon* were also *Helictochloa* and *Avenula*, which were placed even more distantly in the molecular phylogenetic analyses in the clade of Poeae. Further excluded was the nothogenus ×*Trisetopsotrichon* Röser & A.Wölk. The redefined genus *Helictotrichon* [type *H. sempervirens* (Vill.) Pilg.; studied also by Wölk & Röser, 2014, 2017; Wölk & al. 2015] encompasses the former genus *Pseudarrhenatherum* Rouy [type *H. thorei* Röser = *P. longifolium* (Thore) Rouy] and was corroborated in this circumscription as monophyletic (data not shown; Appendix 1; Schneider & al., 2009). *Avena* L., a genus with consistently annual species except for perennial *A. macrostachya* Balansa ex Coss. & Durieu, was close to *Arrhenatherum* P.Beauv. [type *A. elatius* (L.) P.Beauv. ex J.Presl & C.Presl] according to the plastid DNA but not the nuclear DNA trees, in which *Arrhenatherum* clustered with *Tricholemma* and *Helictotrichon*.

The delineation of genera within the Koeleriinae lineage is still an insufficiently solved problem as there are seemingly intermediates between traditionally acknowledged genera and a considerable degree of hybrid speciation and allopolyploid evolution (Quintanar & al., 2010; Saarela & al., 2010, 2017; Wölk & Röser, 2014, 2017; Wölk & al., 2015; Soreng & al., 2017; Barberá & al., 2019). As a possible consequence, all genera of Koeleriinae widely accepted at the present time and additionally *Leptophyllochloa* Calderón ex Nicora were unified by Kellogg (2015) under a single genus, *Trisetaria* Forssk. More detailed investigations using a broad phylogenetic sampling of taxa are evidently still necessary to delineate well-defined genera within Koeleriinae or, alternatively, infrageneric entities in broadly delineated *Trisetaria*.

The backbone of the plastid DNA tree showed largely a polytomy for Koeleriinae (excluding *Lagurus*) if not considering the unsupported placement of *Sibirotrisetum* Barberá, Soreng, Romasch., Quintanar & P.M.Peterson (Figs. 1, 5). One maximally supported clade (100/99/1.00) contained the sampled species of *Graphephorum* [type *G. melicoides* (Michx.) Desv.], *Limnodea* (type *L. arkansana*), *Peyritschia* E.Fourn. [the type *P. koelerioides* (Peyr.) E.Fourn. together with *P. pringlei* (Scribn.) S.D.Koch used in this study and *P. deyeuxioides* (Kunth) Finot studied by Wölk & Röser (2017) showing their monophyly], *Sphenopholis Scribn.* [type *S. obtusata* (Michx.) Scribn.], *Trisetopsis* as well as *Trisetum canescens* Buckley, *T. cernuum* Trin. and ‘*Calamagrostis*’ *rigida* as representative of the Mexican to South American taxa of ‘*Calamagrostis’*/‘*Deyeuxia’*. This clade agreed largely with Koeleriinae clade B of Saarela & al. (2017) and Barberá & al. (2019). This lineage was present in principle also in the nuclear DNA analyses (98/92/1.00; Figs. 2, 5) but without *Graphephorum* spp., *Trisetum canescens* and *T. cernuum*. They assembled in a strongly supported clade (ML 98/97/1.00) with *Avellinia* Parl. [type *A. michelii* (Savi) Parl.], *Gaudinia* J.Gay [type *G. fragilis* (L.) P.Beauv.], *Rostraria* Trin. [type *R. pubescens* Trin. = *R. cristata* (L.) Tzvelev], *Trisetaria*, *Koeleria* Pers. [type *K. pyramidata* (Lam.) P.Beauv.] including the former genus *Parafestuca* E.B.Alexeev [type *P. albida* (Lowe) E.B.Alexeev ≡ *K. loweana* Quintanar, Catalán & Castrov.], *Trisetum* [type *T. flavescens* (L.) P.Beauv.], *Acrospelion* Besser [type *A. distichophyllum* (Vill.) Barberá] and *Tzveleviochloa* This clade agreed with Koeleriinae clade A (Saarela & al., 2017; Barberá & al., 2019) despite different sampling. The changing position of American *Graphephorum* species, *Trisetum cernuum* and *T. canescens* in plastid and nuclear analyses (Figs. 1, 2, 5; Wölk & Röser, 2014, 2017; Saarela & al., 2017) points to their likely hybrid origin.

*Sibirotrisetum sibiricum* (Rupr.) Barberá, type of *Sibirotrisetum*, segregated from the species currently ascribed to genera *Trisetum* and *Acrospelion* in both the plastid and the nuclear DNA analysis. It was part of the backbone polytomy or placed in an unsupported clade with clade A genera (Figs. 1, 5) in the former analysis and sister to the Koeleriinae clade B (74/68/0.86) in the latter (Figs. 2, 5). Most likely due to the different taxon sampling, *S. sibiricum* stood in the study of Barberá & al. (2019) in a polytomy with clades A and B according to the nuclear DNA, whereas it was sister to clade B according to their plastid DNA data.

#### Brizinae and Macrobriza

Monospecific *Airopsis* Desv. (*A. tenella*) was a supported member of subtribe Brizinae. It segregated from the representatives of the genus *Briza* L. (type *B. minor* L.) in the molecular phylogenetic trees and is also morphologically distinct enough to be acknowledged as separate genus (Figs. 1, 2, suppl. Fig. S1). Remarkable is its long branch in the trees, comparable to that of other annual taxa in this study such as *Echinaria* Desf., *Mibora* Adans. (Sesleriinae), *Ammochloa* Boiss. (Ammochloinae), *Rhizocephalus* Boiss. (Beckmanniinae), *Brizochloa* (Brizochloinae) and annual species of *Poa* (Figs. 1, 2, suppl. Fig. S1). *Briza media* L., *B. minor* L. and *Macrobriza maxima* (L.) Tzvelev were placed in a common lineage according to the plastid DNA data, whereas *Macrobriza* deviated clearly in the nuclear DNA tree (see above *Hybrid origin of major lineages…*). This discordant placement is implicitly obvious also in the study of Persson & Rydin (2016), which showed *M. maxima* together with *B. marcowiczii* Woronow, *B. media* L. and *B. minor* L. placed in a common clade according to the plastid DNA data, whereas *M. maxima* clustered with taxa of Aveninae according to the nuclear ITS/GBSSI data. The sample studied by Essi & al. (2008) encompassed *Briza*, *Macrobriza* and species nowadays assigned to *Chascolytrum* Desv. but no taxa of Aveninae. *Macrobriza* clustered with *Briza minor* in the ITS/GBSSI tree of their Fig. 2, whereas *B. media* clustered with taxa of *Chascolytrum*, which may be due to insufficient taxon sampling as noted by Saarela & al. (2017).

Considering morphology, monospecific *Macrobriza* by and large resembles *Briza* but differs by its overall tall size, comparatively few-flowered spikelets and a linear hilum of the caryopsis, which induced Tzvelev to treat it firstly as *Briza* subsect. *Macrobriza* Tzvelev and later as a genus (Tzvelev, 1970, 1993). Its hybrid origin between ancestors from Brizinae (plastid donor) and Aveninae/Sesleriinae as discussed above suggests its exclusion from Brizinae.

#### Echinopogoninae and Calothecinae

These are subtribes of the southern hemisphere. Australasian Echinopogoninae were strongly supported (100/99/1.00) as monophyletic by the plastid DNA data (not available for New Guinea to Queensland *Ancistragrostis*; Fig. 1) but formed to a large extent a polytomy with other subtribes of supersubtribe Agrostidodinae in the nuclear and combined data (Figs. 2, 3, suppl. Fig. S1). Echinopogoninae were represented in this study by monospecific *Ancistragrostis*, two species of *Dichelachne*, *Echinopogon*, *Relchela* (monospecific), two accessions of *Pentapogon* (monospecific) and ‘*Deyeuxia’ contracta*. A placement within Echinopogoninae was reported also for the species of ‘*Deyeuxia*’ from Australia and New Zealand studied by Saarela & al. (2017). They belong to the ∼40 species of this region unified under the genus name ‘*Deyeuxia’* (Vickery, 1940; Weiller & al., 2009; Edgar & Connor, 2000) as a presumed segregate of *Calamagrostis*. ‘*Deyeuxia’ contracta* was closer to *Pentapogon* than to *Dichelachne* according to the plastid and nuclear DNA data (Figs. 1, 2). Merging *Dichelachne* with ‘*Deyeuxia’* (Kellogg, 2015) therefore was not supported, unless *Pentapogon* would likewise be abandoned as a genus. For nomenclature reasons, the genus name ‘*Deyeuxia’* is not applicable anyway (see following chapter).

Calothecinae (Mexico to South America) encompasses only *Chascolytrum* Desv. [type *C. subaristatum* (Lam.) Desv.] after inclusion of several segregate genera such as *Erianthecium* (type *E. bulbosum* Parodi), *Rhombolytrum* Link (type *R. rhomboideum* Link), *Poidium* Nees [*P. uniolae* (Nees) Matthei sampled] and others (Essi & al., 2017). *Chascolytrum* proved monophyletic in this study according to the nuclear DNA data (89/88/1.00; Fig. 2) such as found by Persson & Rydin (2016) for their set of taxa, whereas *Briza media* was nested among New World *Chascolytrum* taxa in the ITS/GBSSI tree of Essi & al. (2008: fig. 2) although without support. As addressed above, this was possibly due to insufficient taxon sampling.

*Agrostidinae and Hypseochloinae;* Calamagrostis *and* Deyeuxia. – Subtribe Agrostidinae, characterized by single-flowered spikelets, is well-supported as monophyletic by the plastid DNA (83/79/1.00) but not the nuclear and combined data analyses, due to the polytomy mentioned. African *Hypseochloa* deviating from Agrostidinae in all analyses (Figs. 1, 2, 4) belongs to supersubtribe Agrostidodinae but it may be best to assign it to a new monogeneric subtribe, Hypseochloinae, which is morphologically supported by peculiar lemma characters not found elsewhere in supersubtribe Agrostidodinae (Hubbard, 1936, 1981; Clayton & Renvoize, 1986; Kellogg, 1995; see below *New names and combinations*).

Agrostidinae comprised three well-supported lineages in the plastid DNA tree, which were arranged largely in a polytomy with the remainder of this subtribe and, in the nuclear DNA tree, even in an even more extended polytomy with taxa of Brizinae, Calothecinae and Echinopogoninae.

One of these well-supported Agrostidinae lineages in both analyses (plastid and nuclear DNA; 100/100/1.00 and 100/0.99/1.00, respectively) was composed of *Gastridium phleoides* (Nees & Meyen) C.E.Hubb., *G. ventricosum* (Gouan) Schinz & Thell. (type of *Gastridium* P.Beauv.) and *G. nitens* (Guss.) Coss. & Durieu (type of *Triplachne* Link) as similarly found in several previous studies (Davis & Soreng, 2007; Quintanar & al., 2007; Soreng & al., 2007; Döring, 2009; Saarela & al., 2010, 2017; Orton & al., 2019). Species of *Gastridium* and former *Triplachne* even were intermingled in the plastid DNA tree. This supports to assign them to a single genus, which is emphasized also by their strong morphological similarity (Clayton & Renvoize, 1986).

The second supported lineage in the plastid DNA tree (96/81/1.00; Fig. 1) comprised *Agrostis alopecuroides* Lam. [= *Polypogon monspeliensis* (L.) Desf., type of *Polypogon* Desf.], *A. avenacea* J.F.Gmel. [= *Lachnagrostis filiformis* (G.Forst.) Trin., type of *Lachnagrostis* Trin.], *A. capillaris* L., *A. linkii* Banfi, Galasso & Bartolucci [= *Chaetopogon fasciculatus* (Link) Hayek, type of *Chaetopogon* Janch.], *A. pallens* Trin., *A. ramboi* Parodi [≡ *Bromidium ramboi* (Parodi) Rúgolo] and *A. scabra* Willd. In the nuclear DNA tree, the lineage disintegrated into a polytomy with strongly supported *A. avenacea* and *A. alopecuroides* as sister (100/100/1.00) and a monophyletic lineage of the remaining species (100/97/1.00), which was complemented by a further species of former genus *Bromidium* Nees & Meyen [*A. tandilensis* (Kuntze) Parodi; only ITS]. *Agrostis linkii* was sister to *A. capillaris* and former *Bromidium* was non-monophyletic. This makes it reasonable to include all taxa in a broadly circumscribed genus *Agrostis* L. as suggested already for *Chaetopogon* (Kellogg, 2015; Soreng & al., 2017; Banfi & al., 2018). Former *Chaetopogon* was also nested within *Agrostis* considering the ITS data investigated by Quintanar & al. (2007) and Saarela & al. (2010, 2017). The same applies to former *Polypogon*, which shares spikelets falling entire and other characters with former *Chaetopogon*. Barely separable are also *Lachnagrostis*, a richly evolved group in temperate Australasia encompassing ∼38 species (Jacobs & Brown, 2009), and *Bromidium*, which encompasses 5 species in South America (Rúgolo de Agrasar, 1982).

The third well-supported lineage of Agrostidinae (plastid 94/94/1.00, nuclear 99/98/1.00) was represented by two species sampled of *Podagrostis* (Griseb.) Scribn. & Merr., *P. aequivalvis* (Trin.) Scribn. & Merr. (type of *Podagrostis*) and *P. thurberiana* (Hitchc.) Hultén. It was separate from *Agrostis* in both analyses, which supports to maintain *Podagrostis* as a distinct genus (Figs. 1, 2).

The holarctic, temperate species of *Calamagrostis* sampled in this study (Old and New World) belonged to Agrostidinae, whereas the Mexican to South American and Australasian taxa sampled were nested in the lineages of Aveninae and Echinopogoninae, respectively (Figs. 1, 2; Saarela & al., 2010, 2017; Wölk & Röser, 2014, 2017). The latter were usually treated under either *Calamagrostis* or more frequently *Deyeuxia* (for example, Bor, 1960; Nicora & Rúgolo de Agrasar, 1987; Villavicencio, 1995; Renvoize, 1998; Edgar & Connor, 2000; Sharp & Simon, 2002; Rúgolo de Agrasar, 2006, 2012a; Weiller & al., 2009). None of both genus names can be used for them because the type of *Calamagrostis* is *Arundo calamagrostis* L., a synonym of *C. canescens* (Weber) Roth, which was nested within Agrostidinae (Figs. 1, 2). The type of *Deyeuxia* Clar. ex P.Beauv. is *D. montana* (Gaud.) P.Beauv., a synonym of *C. arundinacea* (L.) Roth, which likewise belongs to Agrostidinae. Moreover, *Deyeuxia* is synonymous with *Calamagrostis* (Wölk & Röser, 2014; Saarela & al., 2017), if *C. canescens* and *C. arundinacea* belong to a single genus, which is more than likely considering the plastid and nuclear DNA analyses, in which both species were placed with all other species of *Calamagrostis* sampled from Eurasia and North America in a polytomy (Figs. 1, 2).

An exception was Tibetan ‘*Calamagrostis*’ *flavens* (Keng) S.L.Lu & Z.L.Wu. This species clustered in the plastid DNA tree together with *Podagrostis*, *Gastridium* and *Agrostis* in a considerably supported clade (95/70/1.00), whereas it was part of the polytomy of Agrostidinae/Brizinae/Calothecinae/Echinopogoninae in the nuclear DNA tree (Figs. 1, 2). Morphologically, this species has an unusual combination of characters otherwise found in *Agrostis* and *Calamagrostis* as noted by Lu & al. (2006), which seems to fit its ambiguous placement in the trees and points to a possible intergeneric hybrid origin.

*Calamagrostis arenaria* (L.) Roth, type of the former genus *Ammophila* Host, fell within *Calamagrostis* in the nuclear DNA data tree (Fig. 2) in agreement with Saarela & al. (2017). Our plastid DNA tree was not decisive (Fig. 1), whereas that of Saarela & al. (2017) clearly showed *C. arenaria* nested within traditional *Calamagrostis* species. Inclusion of this awnless or shortly awned species in *Calamagrostis* agrees also with morphological data, although previous literature mostly kept this ecologically notable species of coastal dune sands separate (Tutin, 1980; Conert, 1979–1998, 2007). Species of the former genus *Ammophila*, viz. *C. arenaria* in Europe and *C. breviligulata* (Fernald) Saarela in North America, are known to hybridize with *C. epigejos* and *C. canadensis* (Michx.) P.Beauv., respectively. Hybrids between the former species were named *C.* ×*baltica* (Flüggé ex Schrad.) Trin. [≡ ×*Calammophila baltica* (Flüggé ex Schrad.) Brand ≡ ×*Ammocalamagrostis baltica* (Flüggé ex Schrad.) P.Fourn. = *Calamagrostis ×calammophila* Saarela]. They form amazingly extensive stands along the coasts of the North and the Baltic Sea (Conert, 1979–1998; Tutin, 1980), where they are locally more abundant than the parental species.

#### Sesleriinae and Scolochloinae

The largely European subtribe Sesleriinae encompasses species with capitate or spiciform inflorescences, among them two small genera of short-lived annuals, *Mibora* with two species [type *M. minima* (L.) Desv.] and monospecific *Echinaria* [*E. capitata* (L.) Desf.]. *Mibora* and *Oreochloa* [type *O. disticha* (Wulfen) Link] were sister in all analysis.

They formed a sister clade to *Sesleria* [type *S. caerulea* (L.) Ard.] and monospecific *Sesleriella* [type *S. sphaerocephala* (Ard.) Deyl] in the plastid DNA tree (Figs. 1, 6), in which *Echinaria* clustered with monospecific *Psilathera* [type *P. ovata* (Hoppe) Deyl]. In the nuclear tree, *Mibora*/*Oreochloa* stood in a polytomy with *Sesleriella* and supported clade of *Echinaria*, *Psilathera* and *Sesleria* (Figs. 2, 6). The origin of *Sesleria* through hybridization between a *Sesleriella*-like maternal ancestor and a *Psilathera*-like paternal ancestor (see Kuzmanović & al., 2017 and above) represents also a good example of allopolyploidy because *Sesleria* comprising consistently polyploid species (4*x*–12*x*), whereas *Sesleriella* (most likely monospecific) and *Psilathera* (monospecific) are diploid. Also for *Oreochloa*, a genus with four species occurring in the European Alpine mountain system, only diploids are known so far. The relationships resolved by the plastid and the nuclear DNA analysis were congruent and *Oreochloa* is not involved in the origin of *Sesleria* (Figs. 1, 2, 6).

Scolochloinae encompass *Scolochloa* [two species; type *S. festucacea* (Willd.) Link] from the temperate regions of the Holarctic and monospecific Australian *Dryopoa* [*D. dives* (F.Muell.) Vickery]. Both resemble one another morphologically (Clayton & Renvoize, 1986) and represent a remarkable example of bipolar distribution. The genera were not supported to be closely related by the plastid DNA analysis, in which *Scolochloa* aggregated with *Antinoria* (86/70/0.68) and *Dryopoa* with Sesleriinae, although without strong support (Fig. 1). The nuclear and the combined DNA data analyses showed *Scolochloa* and *Dryopoa* as strongly supported sister (100/97/1.00 and 100/100/1.00, respectively; Figs. 2, 3, suppl. Fig. S1). The nuclear DNA revealed Scolochloinae in a common and considerably supported clade comprising Phalaridinae, Torreychloinae and supersubtribe Agrostidodinae, which corroborates the findings of Birch & al. (2014) who noted a relationship of *Dryopoa* to Brizinae and Agrostidinae.

#### Aristaveninae

Segregation of this subtribe from Holcinae and Airinae as addressed earlier (Schneider & al., 2012; Saarela & al., 2017; Soreng & al., 2017) was supported by all analyses of this study. Aristaveninae encompass only *Deschampsia* P.Beauv. [type *D. cespitosa* (L.) P.Beauv.], in which the former monospecific genus *Scribneria* Hack. [*S. bolanderi* (Thurb.) Hack.], whose relationship with *Deschampsia* was established by Schneider & al. (2012), was included (Saarela & al., 2017). *Deschampsia* encompasses seemingly also the former monospecific south Andean genus *Leptophyllochloa* as supported by all analyses of this study (Figs. 1, 2, suppl. Fig. S1; Wölk & Röser, 2017). The accessions of *Leptophyllochloa* studied by Saarela & al. (2017) and Barberá & al. (2019), however, resolved within the Koeleriinae lineage of Aveninae. This induced us to re-examine the voucher specimen we used. It had been collected and identified by Z. Rúgolo (*Rúgolo* 1245; see Appendix 1) and we were able to ascertain the correct identification, such as for a second accession examined (*Rúgolo* 1250, B 10 0448862). Transferring *L. micrathera* (É.Desv.) Calderón ex Nicora to *Deschampsia* and placing *Leptophyllochloa* under synonymy of *Deschampsia* (see below *New names and combinations*) also is in good agreement with their morphology (pers. observ.; Nicora, 1978; Rúgolo, 2012b).

#### Helictochloinae, Antinoriinae, Airinae and Holcinae

Helictochloinae is newly established as subtribe to accommodate *Helictochloa* Romero Zarco [type *H. bromoides* (Gouan) Romero Zarco], a widespread Eurasian/Mediterranean perennial, and *Molineriella* [type *M. minuta* (L.) Rouy], a Mediterranean annual genus. Both genera had maximum support as sisters in all analyses and segregated consistently from Airinae s.l., which disintegrated further into Antinoriinae and Airinae (Figs. 1, 2, 3, suppl. Fig. S1). The subtribe Helictochloinae is morphologically hard to define, because its genera differ substantially. However, the spikelets disarticulate below each floret, the rhachilla is glabrous or sparsely hairy, the lemma has a hairy callus and a dorsal awn though not consistently in *Molineriella*. Lodicules have a lateral tooth (pers. observ.; Cebrino Cruz & Romero Zarco, 2017)

Subtribe Antinoriinae encompassing only *Antinoria* [type *A. agrostidea* (DC.) Parl.] was close to Loliinae in all analyses (Figs. 1, 2, suppl. Fig. S1). In the nuclear and combined DNA trees, it was weakly supported sister to supersubtribe Loliodinae, viz. the lineage with subtribes Loliinae, Ammochloinae, Dactylidinae, Cynosurinae and Parapholiinae, a placement agreeing with previous ITS studies (Quintanar & al., 2007; Inda & al., 2008).

Airinae as defined in this study were clearly monophyletic in all analyses (Figs. 1, 2, suppl. Fig. S1). They encompass *Aira* L. (type *A. praecox* L.), *Avenella* Bluff ex Drejer [type *A. flexuosa* (L.) Drejer], *Corynephorus* P.Beauv. [type *C. canescens* (L.) P.Beauv.] and *Periballia* Trin. [type *P. involucrata* (Cav.) Janka].

Monophyletic Holcinae with *Holcus* and *Vahlodea* Fr. [type *V. atropurpurea* (Wahlenb.) Fr.] were sister to Airinae as supported by the nuclear DNA data (95/84/1.00; Fig. 2) and the combined data analysis (88/88/1.00; suppl. Fig. S1) as similarly found by Quintanar & al. (2007), who sampled only *Holcus*, and by Schneider & al. (2009) for *Holcus* and *Vahlodea*. Plastid DNA data showed Holcinae in a polytomy with Aristaveninae and Helictochloinae (Figs. 1, 4; similarly found by Quintanar & al., 2007; Schneider & al., 2009), as well as with Loliinae, Scolochloinae, Sesleriinae and the ADCP clade, whereas Airinae were more distant. This means that Holcinae share plastid DNA characters with a larger set of subtribes but to a lesser extent with Airinae, whereas nuclear DNA connects Holcinae in particular with Airinae. This indicates that Holcinae might have ancient hybrid origin slightly different from that of Airinae although both tribes share an overall similar pattern of conflicting placements with respect to the plastid and nuclear DNA trees (see above *Comparison of the plastid and nuclear DNA trees*).

#### Loliinae

The large and worldwide distributed subtribe Loliinae was represented in this study by a small sample of taxa. It has been investigated and shown to be monophyletic by several previous studies (see Introduction). Affiliation of former *Megalachne* Steud. and *Podophorus* Phil., endemics of the Juan Fernándes Islands (Chile), with Loliinae was established by Schneider & al. (2011, 2012). Subtribe Loliinae was characterized in this study by two supported main lineages in the plastid and combined data analyses (Fig. 1, suppl. Fig. S1), one of which was formed by *Drymochloa sylvatica* (Pollich) Holub (= *Festuca altissima* All., type of *Drymochloa*) as clear sister to *Lolium* L. species, namely *L. perenne* L. (type of *Lolium*), *L. rigidum* Gaudin and *L. giganteum* (L.) Darbysh. [≡ *F. gigantea* (L.) Vill. ≡ *Schedonorus giganteus* (L.) Holub].

The second lineage was formed by *Castellia tuberculosa* (type of *Castellia*) as sister to a monophyletic lineage, which corresponds to a narrowly defined genus *Festuca* that is equivalent to the “fine-leaved fescues” (Torrecilla & Catalán, 2002) as suggested (Kellogg, 2015; Soreng & al., 2015b, 2017). This lineage of *Festuca* s.str. was represented in our sample by species from several morphologically partly well-defined segregate genera, namely *F. berteroniana* Steud. (type of *Megalachne*), *F. floribunda* (Pilg.) P.M.Peterson, Soreng & Romasch. (type of *Dielsiochloa* Pilg.), *F. incurva* (Gouan) Gutermann (type of *Psilurus* Trin.), *F. lachenalii* (C.C.Gmel.) Spenn. [type of *Micropyrum* (Gaudin) Link], *F. masatierrae* Röser & Tkach, nom. nov. (type of *Podophorus*; no plastid DNA data available), *F. maritima* L. [= *Vulpia unilateralis* (L.) Stace], *F. myuros* L. (type of *Vulpia* C.C.Gmelin) and *F. salzmannii* (Boiss.) Boiss. ex Coss. (type of *Narduroides* Rouy).

The nuclear DNA results agreed widely with the trees of the plastid data analyses, however, *Castellia* was differently placed, namely together with *Drymochloa* (Fig. 2) and not with the lineage of *Festuca* s.str. as just mentioned. This points to a hybrid origin of this odd monotypic, Mediterranean to mid-East genus (see above *Reticulations within major lineages*).

The South American representatives sampled of *Festuca* s.str., namely *F. floribunda* from the Andes and the endemics of Chilean Juan Fernández Islands, *F. berteroniana* and *F. masatierrae*, formed a monophyletic cluster in the nuclear DNA analysis. *Festuca floribunda* belongs to the group “American II” of fine-leaved *Festuca* in the study on the historical biogeography of Loliinae by Minaya & al. (2017). Group “American II” has colonised South America in the Miocene, a time frame that makes sense also for the establishment of *F. berteroniana* and *F. masatierrae*. The islands started to originate in the Upper Miocene 5.8 million years ago (Stuessy & al., 1984).

#### ADCP clade

The species of the small sister subtribes Ammochloinae and Dactylidinae forming the AD clade have spikelets arranged in dense clusters. The close relationship of both tribes (Figs. 1, 2) was revealed already by the plastid DNA data of Quintanar & al. (2007) and Orton & al. (2019), whereas the ITS data of *Ammochloa palaestina* Boiss. of the former study used by Saarela & al. (2010) were wrong and belonged to *Helictochloa* (Appendix 3). Ammochloinae are monogeneric (*Ammochloa*; type *A. palaestina*), whereas Dactylidinae encompass Eurasian *Dactylis* (type *D. glomerata* L.) und Mediterranean to mid-East monospecific *Lamarckia* Moench [type *L. aurea* (L.) Moench]. The sister relation of the AD to the CP clade was found in all analyses of this study (Figs. 1, 2, 4, suppl. Fig. S1) although without strong support, which agrees with the plastid DNA results of several previous studies (Davis & Soreng, 2007; Quintanar & al., 2007; Bouchenak-Khelladi & al., 2008).

The Cynosurinae species *C. cristatus* L. (type of *Cynosurus* L.) and *C. elegans* Desf. were not resolved as monophyletic but formed a grade basal to Parapholiinae.

Parapholiinae were strongly supported as monophyletic in all analyses (Figs. 1, 2, suppl. Fig. S1), in agreement with Davis & Soreng (2007), Quintanar & al. (2007) and Bouchenak-Khelladi & al. (2008). Its species are distributed from the Mediterranean to the Middle East and frequently grow on saline soil. Parapholiinae encompass six genera of annuals if, firstly, the former monotypic genus *Hainardia* Greuter [type *H. cylindrica* (Willd.) Greuter] is reduced to synonymy of *Parapholis* C.E.Hubb. [type *P. incurva* (L.) C.E.Hubb.] as concordantly suggested by our plastid and nuclear DNA analyses (Figs. 1, 2) and, secondly, the endemic Algerian monotypic and perennial genus *Agropyropsis*, which was not molecularly studied to date, belongs to Loliinae as suggested by morphological characters (Schneider at al., 2012). The remaining Parapholiinae genera in addition to *Parapholis* are *Catapodium* Link [type *C. marinum* (L.) C.E.Hubb.], *Cutandia* Willk., *Desmazeria* Dumort. [type *D. sicula* (Jacq.) Dumort.], *Sphenopus* Trin. [type *S. divaricatus* (Gouan) Rchb.] and *Vulpiella* (Batt. & Trab.) Burollet [type *V. stipoides* (L.) Maire].

*Desmazeria philistaea* and *D. sicula* were sister taxa and monophyletic (100/100/1.0) in the plastid (Fig. 1) but not in the nuclear DNA tree, in which *D. sicula* clustered with *Vulpiella* and *Cutandia* (96/95/1.00; Fig. 2; see also Schneider & al., 2012). *Desmazeria sicula* is likely to be a hybrid, which may lead to a name change for this genus, pending further investigation.

#### PPAM clade and Coleanthinae

The PPAM clade was resolved in all analyses of this study. It was more strongly supported in the nuclear than the plastid DNA analysis but obtained maximum support in the combined data tree (see above *Tree of the combined plastid and nuclear DNA dataset*; Figs. 1, 2, suppl. Fig. S1).

Within the monophyletic subtribe Coleanthinae (= Puccinelliinae), several species were repeatedly transferred from one genus to another and genus limits are still in dispute. Our results support to recognize ten genera, which partly have a new delineation. A close relationship of perennial *Colpodium* and annual species that usually were treated under *Zingeria* P.A.Smirn. was suggested by all our trees and had already been noted by Tzvelev & Bolkhovskikh (1965) and Soreng & al. (2017). The tree from the plastid DNA data showed the sampled representatives of *Colpodium* and former *Zingeria* intermingled (Fig. 1), namely *C. biebersteinianum* (Claus) Röser & Tkach, comb. nov. (type of *Zingeria*), *C. versicolor* (Steven) Schmalh. (type of *Colpodium*), *C. trichopodum* (Boiss.) Röser & Tkach, comb. nov. [≡ *Z. trichopoda* (Boiss.) P.A.Smirn.], *C. hedbergii* (Melderis) Tzvelev and *C. chionogeiton* (Pilg.) Tzvelev, both of which occur in Africa and had sometimes been accommodated also under *Keniochloa* Melderis [type *K. chionogeiton* (Pilg.) Melderis]. The nuclear DNA tree showed the former *Zingeria* species in a grade with the other *Colpodium* species sampled (Fig. 2) as similarly encountered by Kim & al. (2009). If their different life form is left aside there are no striking differences between *Colpodium* and former *Zingeria*, both of which have small spikelets with a single bisexual flower, and we suggest unifying them under a single genus. *Colpodium* is the genus with the lowest monoploid chromosome number known in grasses of *x* = 2. There are known diploids with 2*n* = 4 (*C. biebersteinianum*, *C. versicolor*) and several polyploids, namely *C. trichopodum* and *C. pisidicum* (Boiss.) Röser & Tkach, comb. nov., with 2*n* = 8 and *C. kochii* (Mez) Röser & Tkach, comb. nov., with 2*n* = 12. *Colpodium versicolor* (2*x*) was shown to be the donor of one genome in allohexaploid *C. kochii* (Kotseruba & al., 2010), whereas it is not represented in allotetraploid *C. trichopodum* (Kotseruba & al., 2005).

The nuclear tree revealed *Hyalopodium* (*H. araraticum*) as sister to the clade with the species of *Colpodium*, which was similarly encountered in the ITS studies of Rodionov & al. (2008) and Kim & al. (2008, 2009). However, the plastid tree supported a deviant relationship of *Hyalopodium* (Figs. 2, 7), namely to *Paracolpodium* and *Hyalopoa* (*H. pontica*). These differences between the plastid and nuclear DNA analyses suggest an origin of *Hyalopodium* as a hybrid between two different lineages of Coleanthinae (Fig. 7). *Hyalopodium araraticum* has formerly been treated under *Catabrosa*, *Colpodium* or *Catabrosella*. It is long-known as remarkable species because of its odd combination of morphological characters and had been placed in a monospecific section of *Catabrosella*, namely *C.* sect. *Nevskia* (Tzvelev) Tzvelev (see Tzvelev, 1976), which seemingly had never been validly raised to genus rank although that was stated by Kim & al. (2008). *Hyalopodium araraticum* has spikelets with several flowers such as found in *Catabrosella* and *Hyalopoa* s.str., has creeping underground shoots like *Paracolpodium* and *Hyalopoa*, whereas *Catabrosella* and *Colpodium* are not creeping (Tzvelev, 1964a). A conspicuous character of *Hyalopodium* among Coleanthinae are its aerial shoots with reticulate-fibrous sheaths of dead leaves at the base, which, however, resemble the filamentously, though not reticulately decaying basal leaf sheaths of *Hyalopoa pontica* (pers. observ.; Mill, 1985). Chromosomally, it has a monoploid number of *x* = 7 like *Paracolpodium* and *Hyalopoa* (CCBD, 2019), not *x* = 2 as in *Colpodium*. This makes it likely that the paternal parent of *Hyalopodium* did not come from present-day *Colpodium*, but was an ancestor still having the plesiomorphic monoploid chromosome number of *x* = 7 (Fig. 7).

*Paracolpodium altaicum* (type of *Paracolpodium*) and *P. baltistanicum* clustered together with *Hyalopoa pontica* (type of *Hyalopoa*) in all analysis of this study although with weak support (Figs. 1, 2, 7). Both genera consistently encompass species with creeping underground shoots in contrast to tufted *Colpodium* and *Catabrosella*. *Paracolpodium* and *Hyalopoa* also share further morphological characters such as comparatively long glumes, large lodicules, a caryopsis with a rostrate tip and ay long hilum and the margins of leaf sheaths fused for more than 1/3 from the base (Tzvelev, 1976, Cope, 1982). The main difference are the number of florets in the spikelets, which usually have a single but sometimes an additional sterile floret in *Paracolpodium* or the spikelet is two-flowered with the lower floret sterile (*P. baltistanicum*; Dickoré, 1995), whereas *Hyalopoa* has 3–4 flowered spikelets.

*Catabrosa* [type *C. aquatica* (L.) P.Beauv.] and *Catabrosella* [type *C. humilis* (M.Bieb.) Tzvelev] were well-supported separate genera (see Appendices 1 and 2 for further species molecularly sampled). In the plastid DNA analyses, both genera formed a sister clade to *Puccinellia*/*Sclerochloa.* In the nuclear DNA tree, they were together with *Hyalopoa*/*Paracolpodium* sister to a clade of *Puccinellia*/*Sclerochloa* and *Coleanthus*/*Phippsia* as similarly found in other studies (Fig. 7; Rodionov & al., 2008; Schneider & al., 2009; Soreng & al., 2015b; Nosov & al., 2019). This suggests a reticulation process within Coleanthinae in way that the *Puccinellia*/*Sclerochloa* lineage has hybrid background, namely *Coleanthus*/*Phippsia*-like rDNA from its paternal ancestor while its maternal rDNA from *Catabrosa*/*Catabrosella* was lost.

Monospecific holarctic annual *Coleanthus* [*C. subtilis* (Tratt.) Seidel ex Roem. & Schult.]) was clear sister to perennial Arctic (two species) and high Andean (one species) *Phippsia* [type *P. algida* (Sol.) R.Br.]. Both genera share conspicuous morphological characters such as missing or obsolescent, small glumes and a caryopsis protruding from the floret at maturity (Nicora & Rúgolo de Agrasar, 1981; Clayton & Renvoize, 1986; Rúgolo, 2012b).

The sister relation of the small genus *Sclerochloa* [2–3 species; type *S. dura* (L.) P.Beauv.] and the large genus *Puccinellia* (110 species) was likewise firmly supported (Figs. 1, 2, 7), even after inclusion of more species of the latter genus (data not shown; Appendix 2; Hoffmann & al., 2013; Soreng & al., 2015b).

#### PAM clade, Avenulinae, Miliinae and Phleinae

The small subtribes Avenulinae (*Avenula*), Miliinae (*Milium* L.) and Phleinae (*Phleum* L.) formed together with Poinae and the elements of the ABCV(+A) clade the PAM clade (∼supersubtribe Poodinae). It was resolved in the plastid DNA and combined data analysis of this study and encompassed also *Avenula pubescens* (Huds.) Dumort., type of the monospecific genus *Avenula* (Fig. 1, suppl. Fig. S1). The PAM clade was unresolved in the nuclear DNA tree since its subtribes did not join together in a common clade but stood in a polytomy with Coleanthinae (Figs. 2, 4).

In the plastid and nuclear DNA analyses, monogeneric Miliinae and Avenulinae were more or less in a polytomy with the remainder of the PAM clade. Considering the plastid DNA tree (Fig. 1), this applies also to monogeneric Phleinae, in which three species of *Phleum* including *P. crypsoides* (d’Urv.) Hack., the type of *Maillea* Parl., were sampled. Phleinae, however, were sister to Poinae with considerable support according to the nuclear DNA analysis (Figs. 2, 4), which underpins a possible hybrid origin of this lineage. For *Avenula*, a suspected intergeneric hybrid (Soreng & Davis, 2000) between *Helictotrichon* (Aveninae) and *Helictochloa* (Helictochloinae), there was no supported incongruence between the placements in the plastid and nuclear DNA trees. *Avenula* is unsupported sister to Coleanthinae in the nuclear DNA tree (Figs. 2, 4), whereas it was part of the PAM clade resolved only in the plastid but unresolved in the nuclear DNA tree (Figs. 1, 4). The PAM clade was sister to Coleanthinae, which makes the conflicting placements rather negligible and does not give evidence on hybrid origin of this taxon. In- or exclusion of this taxon did not fundamentally change the tree structure of the nuclear phylogram for the PPAM clade such as described by Gillespie & al. (2008) for their ITS analysis. Morphological characteristics of *Avenula* also speak against the hybrid hypothesis (Gabriel & al., 2019).

#### Poinae

The monogeneric subtribe Poinae was sampled in this study using a small selection of species of traditional *Poa* s.str., namely *P. annua* L., *P. bulbosa* L. and the type of the genus, *P. pratensis* L. This set of taxa was complemented by a several species of previous segregate genera that meanwhile were shown to belong to an enlarged but subsequently monophyletic genus *Poa* (see Introduction for references). Our results corroborate monophyly of *Poa* encompassing *P. apiculata* Refulio (type of *Tovarochloa* T.D.Macfarl. & P.But), *P. labillardierei* Steud. [type of *Austrofestuca* (Tzvelev) E.B.Alexeev], *P. cyrenaica* E.A.Durand & Barratte (type of *Libyella* Pamp.), *P. fax* J.H.Willis & Court (type of *Neuropoa* Clayton), *P. hitchcockiana* Soreng & P.M.Peterson [type of *Aphanelytrum* Hack.], *P. lepidula* (Nees & Meyen) Soreng & L.J.Gillespie (type of *Anthochloa* Nees & Meyen), *P. persica* Trin. (type of *Eremopoa* Roshev.), *P. serpaiana* Refulio (type of *Dissanthelium* Trin.), *P. sintenisii* H.Lindb. (type of *Lindbergella* Bor) and *P. diaphora* Trin., a second species of former *Eremopoa*.

The plastid and nuclear DNA trees were widely congruent and showed sister relations of *P. alpina* and *P. bulbosa*, of *P. annua* and *P. cyrenaica*, of *P. diaphora* and *P. persica*, respectively, and the latter two species together with *P. sintenisii* (corresponding to *Poa* clade E in Gillespie & al., 2018) as sister to the remaining species of *Poa* included in our study (Figs. 1, 2). Nevertheless, there were some differences between the plastid and nuclear analyses. The nuclear DNA tree (Fig. 2), for example, revealed a supported sister relation between *P. labillardierei* and *P. fax* or between *P. apiculata* and *P. hitchcockiana*, respectively, whereas the plastid DNA tree placed them, along with others, in a polytomy (Fig. 1).

#### ABCV(+A) clade

This clade contains many monospecific or species-poor genera, *Alopecurus* with ∼40 species being the largest genus. *Arctopoa* joined this clade only in the nuclear and combined analyses, whereas it was placed outside of it in the plastid DNA tree and close to subtribe Poinae (see above *Comparison of the plastid and nuclear DNA trees*).

Relationships within the ABCV(+A) clade were overall weekly resolved, except for well-supported monophyletic subtribe Ventenatinae, which was retrieved in all analyses. Our results support to abandon *Gaudinopsis* (Boiss.) Eig as monospecific genus [*G. macra* (Steven ex M.Bieb.) Eig] distinct from *Ventenata* Koeler [type *V. dubia* (Leers) Coss. & Durieu]. Monospecific *Parvotrisetum* Chrtek [*P. myrianthum* (Bertol.) Chrtek] was clearly excluded from *Trisetaria*, a member of distantly related subtribe Aveninae. The monospecific genus *Nephelochloa* Boiss. (*N. orientalis* Boiss.) was sister to *Apera* Adans. [type *A. spica-venti* (L.) P.Beauv.] in the nuclear analyses (Fig. 2; Hoffmann & al., 2013). With regards to the plastid DNA (Fig. 1), *N. orientalis* was even nested within the two species sampled of *Apera* (altogether ∼5 species). Also morphologically, both genera share certain characters (usually richly branched inflorescences with numerous primary branches in whorls, similar shape of glumes and lemmas). The main difference is the number of flowers in the spikelets, one in *Apera* and three to six in *Nephelochloa*, which supports to maintain them as separate genera.

The HSAQN clade was well-supported only in the trees of the nuclear and combined DNA data, such as the DAD clade (Fig. 2, suppl. Fig. S1). The former is biogeographically characterized by bipolar distribution. *Arctagrostis* Griseb. [two species; type *A. latifolia* (R.Br.) Griseb.] is distributed in the boreal and arctic regions of the northern hemisphere, whereas the remaining taxa of the HSAQN clade occur in Australasia and southern South America.

The taxa of DAD clade also occur in the boreal and the arctic regions of the northern hemisphere. The new genus *Arctohyalopoa* was nested in the nuclear and combined analyses within this clade (Fig. 2, suppl. Fig. S1), whereas the plastid DNA tree placed it along with many other taxa in the large polytomy of the ABCV clade (Fig. 1). *Arctohyalopoa* comprises only *A. lanatiflora* (Roshev.) Röser & Tkach, comb. nov., which previously has been accommodated in the genus *Hyalopoa* together with *H. pontica* (Balansa) Tzvelev, the type of *Hyalopoa*, and few other species. *Arctohyalopoa lanatiflora* was not nested in the clade of Coleanthinae but in the ABCV or ABCV+A clade, respectively, in which it was part of a polytomy with many other taxa according to the plastid DNA data. It belongs to the DAD clade according to the nuclear and, with strong support, according to the combined DNA data along with *Dupontia* R.Br. (type *D. fisheri* R.Br.), which includes *Arctophila* (Rupr.) Andersson [type *A. fulva* (Trin.) Andersson] (see below *New names and combinations*), and with monospecific *Dupontiopsis* Soreng, L.J.Gillespie & Koba [*D. hayachinensis* (Koidz.) Soreng, L.J. Gillespie & Koba; Figs. 2, 3, suppl. Fig. S1]. This placement of *Arctohyalopoa lanatiflora* distant to Coleanthinae was verified in this study also by analyzing a second accession (data not shown; Appendix 1). In should be noted that our previously published sequence of *Hyalopoa lanatiflora* (Döring & al., 2007; Döring, 2009) is wrong such as seemingly a sequence of Rodionov & al. (2008), which was also used by Hoffmann & al. (2013; for details see Appendix 3).

The molecular phylogenetic results on *Arctohyalopoa* were supported also by morphological data because as pointed out by Tzvelev (1964a: 8) and (1964b: 14–15), *A. lanatiflora* [≡ *Colpodium lanatiflorum* (Roshev.) Tzvelev] differs from both the other species of *Colpodium* subg. *Hyalopoa* Tzvelev (≡ *Hyalopoa*) as well as *Poa* by “lemmas … on basal half especially on nerves with rather copious and long pubescence, with distal part of callus (including that adjoining internerves) also copiously covered with rather long crinkly hairs, … paleas bare and smooth on keels…” (cited from Tzvelev, 1995a: 94–95). The epithet *lanatiflora* refers to the conspicuous indumentum of the lemmas. Tzvelev (1964c, 1995b) also addressed that *A. lanatiflora* otherwise strikingly resembles *Dupontia fulva* (≡ *Arctophila fulva*). Moreover, *Arctohyalopoa lanatiflora* is geographically separated as an eastern Siberian endemic from the species of *Hyalopoa*, which are Caucasian (5 species) and West Himalayan [only *H. nutans* (Stapf) E.B.Alexeev ex T.A.Cope] in distribution. It seems to be also ecologically different due to its preference of non-carbonatic bedrock (Tzvelev, 1964c, 1995b).

Merging the small genera *Dupontia* und *Arctophila* as already suggested by Kellogg (2015) was supported also by Hoffmann & al. (2013), who showed that the nuclear ITS sequences of both were intermingled in the molecular phylogenetic tree. It further agrees with their overall morphological similarity except for rather small difference in the shape of their lemmas (Clayton & Renvoize, 1986; Cayouette & Darbyshire, 2007a,b; see also Brysting & al., 2004) and the occurrence of hybrids between *D. fisheri* and *D. fulva* (Trin.) Röser & Tkach, comb. nov. that were formerly regarded as intergeneric hybrid and treated under the nothogenus ×*Arctodupontia* Tzvelev (Tzvelev, 1973; Brysting & al., 2003; Darbyshire & Cayouette, 2007).

Tribes Alopecurinae, Beckmanniinae and Cinninae in each case did not resolve as monophyletic. The species of *Limnas* Trin. sampled (type *L. stelleri* Trin. and *L. malyschevii* O.D.Nikif.) formed in none of the analysis (plastid, nuclear, combined data) a clade with the other species of Alopecurinae (*Alopecurus aequalis* Sobol., *Cornucopiae cucullatum* L., type of *Cornucopiae*). In the nuclear tree they even were closer to the DAD clade (73/94/1.00) than to *Alopecurus* and *Cornucopiae* (Fig. 2). The latter genera were always sister, which agrees with their common spikelet structure.

*Beckmannia* [type *B. eruciformis* (L.) Host] and monospecific *Pholiurus* [*P. pannonicus* (Host) Trin.] were well-supported sister in the nuclear DNA tree in agreement with Hoffmann & al. (2013) but less supported in the plastid DNA tree (Figs. 1, 2). *Rhizocephalus orientalis*, type of monospecific genus *Rhizocephalus* Boiss. and the third taxon of the tribe Beckmanniinae as delineated by Soreng & al. (2017), was placed in all analyses remotely from *Beckmannia* and *Pholiurus* in the main polytomy of ABCV(+A) clade. Spikelets in *Alopecurus, Cornucopiae*, *Limnas* and *Rhizocephalus* are single-flowered, in *Beckmannia* (the upper staminate) and *Pholiurus* two-flowered (Schneider & al., 2012).

Also the Cinninae genera sampled [*Aniselytron* with type *A. treutleri* (Kuntze) Soják, *Cinna* L., monospecific *Cyathopus* Stapf with *C. sikkimensis* Stapf, *Simplicia* Kirk], appeared in the main polytomy of ABCV(+A) clade, except for *Cinna* and *Cyathopus*, which were supported sister in the nuclear and combined trees. Both share spikelets that are falling entire, whereas *Aniselytron* and *Simplicia* have spikelets disarticulating above the glumes. In all Cinninae genera, the spikelets are single-flowered, with occasional occurrence of a second floret reported for *Simplicia* (Watson & al., 1992 onwards; Edgar & Connor, 2000). Notwithstanding the established sister relations of each *Beckmannia*/*Pholiurus* and *Alopecurus*/*Cornucopiae*, the phylogenetic relationships of all genera of Alopecurinae, Beckmanniinae and Cinninae and the delineation of these tribes certainly warrant future work.

Monospecific *Limnodea* (*L. arkansana*), sometimes placed near or included within *Cinna* (Clayton & Renvoize, 1986; Tucker, 1996), was placed very distant to this genus in the molecular trees, namely within Aveninae and close to *Sphenopholis* (Figs. 1, 2) within the Koeleriinae lineage (see also Döring, 2009; Hochbach & al., 2015; Saarela & al., 2017: suppl. 7).

*Brizochloa humilis* with lemmas that are not cordate as in *Briza* or *Macrobriza* and upright pedicels of the spikelets is a morphologically most striking species of the ABCV(+A) clade. Monospecific genus *Brizochloa* cannot be accommodated under any of the subtribes yet described and we assign it to a new monogeneric subtribe, Brizochloinae. The exclusion of *B. humilis,* an annual distributed from the Eastern Mediterranean to Iran, from *Briza* had been suggested already by previous morphological and molecular studies (Jirásek & Chrtek, 1967; Tzvelev, 1968, 1976; Hoffmann & al., 2013; Persson & Rydin, 2016; Essi & al., 2017).

### Morphological characteristics

The sequence of the following examples of morphological characteristics is character no., character state in brackets, whereby character states with an underline mean the occurrence of different character states in a single taxon. This instance appears in suppl. Fig. S2 as a pie chart. If more than one character was found within a lineage, the character states are given in alphabetical order. If character state (b) is more frequent than (a), it is mentioned first (for example, b, a).

Several morphological characters listed in suppl. Appendix S2 displayed states that were almost consistently found in most lineages retrieved in the molecular phylogenetic analyses: 002 (a), 027 (d), 028 (a), 029 (a), 038 (d), 039 (a), 040 (b), 041 (b), 043 (b), 045 (a), 046 (c), 047 (b), 053 (b), 054 (a), 055 (b), 058 (c), 060 (a), 061 (a), 062 (a), 063 (a), 064 (a), 066 (a), 069 (e), 071 (a), 072 (a), 083 (a), 084 (a), 092 (a), 103 (c), 112 (a), 115 (a), 116 (a), 117 (a), 118 (a), 122 (a, b), 123 (a, b), 124 (a), 126 (a, b), 127 (c), 139 (a), 140 (a), 142 (a), 145 (a), 151 (a), 156 (b), 159 (b), 160 (b, a), 161 (a, a_b) and 164 (a), 169 (d), 176 (a), 181 (c), 182 (b, a), 183 (b, a), 184 (a), 185 (a), 187 (a, b; see suppl. Fig. S2).

Some lineages were characterized by particular character states such as Anthoxanthinae: 57 (b_c), 126 (a_b), 138 (d); Aveninae: 165 (a_e); Phalaridinae: 45 (f); Calothecinae 165 (f), 167 (f); Airinae: 146 (a_d); Dactylidinae: 36 (e); Cynosurinae: 27 (d_g), 45 (b_c), Parapholiinae 28 (c), 57 (b_c), 59 (d), 73 (c), 74 (c), 102 (b_c); Coleanthinae 136 (a_g); Miliinae 16 (c_d), 167 (g); Phleinae 31 (c), 115 (c), (136 (a_g), 127 (b_c); Poinae 119 (b); Beckmanninae 28 (c) and Alopecurinae 50-52 (a), 119 (b_c), 122 (b). Antinoriinae differed from Airinae and Helictochloinae in characters 2, 20, and 119 but also 14 and 74 through which they resembled rather Loliinae. Helictochloinae differed from Airinae and Antinoriinae in characters 41, 73, 125, 149 and 188. Agrostidinae differed from Hypseochloinae in characters 85, 102, 125, 138, 139, 171. Within the ABCV+A clade the placement of *Rhizocephalus* was not unambiguously ascertainable. Its inclusion within Beckmanniinae was supported by characters 17, 18, 24, 26, 48, 50, 52, 124 and possibly 56, whereas affinities to Brizochloinae were suggested by characters 28, 58, 70, 74, 89 and 122. Characters 29 and 30 underscored the unique inflorescence shape of *Rhizocephalus*.

On a whole the number of clear-cut synapomorphic characters that could be used to characterize the retrieved clades in terms of phylogenetic systematics was rather low, which points to a high degree of homoplasy in most morphological characters that were scored. None of the major lineages such as tribes Aveneae and Poeae, supersubtribes Agrostidodinae, Loliodinae and Pooidinae (∼PAM clade), the PPAM clade and also larger subtribes such as Aveninae, Loliinae or Coleanthinae were morphologically reliably identifiable. Some characters revealed identifiable clades corresponding to subtribes but frequently the number of suitable characters was rather low: Agrostidinae: 50 (a), 52 (a), 74 (mostly c); Airinae: 148 (d); Alopecurinae: 50 (a), 51 (a), 52 (a), 58 (b), 148 (d), 169 (b_d); Anthoxanthinae: 50 (a), 51 (a), 52 (a), 115 (c), 138 (d), 139 (b); Calothecinae: 1 (b), 118 (b), 165 (f), 167 (f), 182 (a); Coleanthinae: 9 (a); Dactylidinae: 36 (e); Helictochloinae: 24 (c), 49 (b), 125 (c), 150 (c), 186 (c); Holcinae: 96 (b); Phalaridinae: 45 (a_f), 59 (d), 84 (a_b), 101 (a_b), 115 (c), 165 (c), 169 (a) and Scolochloinae: 72 (b).

Smaller groups of genera sharing common characters could be discerned within several subtribes, for example, *Coleanthus* and *Phippsia* within Coleanthinae: 182 (a); *Parvotrisetum* and *Ventenata*: 148 (d), 150 (c), 155 (c) or *Alopecurus* and *Cornucopiae* within Alopecurinae: 137 (a_b, b), 181 (a).

Single genera representing monogeneric subtribes were frequently identifiable based on (aut-)apomorphic attributes: Ammochloinae (*Ammochloa*): 31 (e), 59 (d), 84 (a_b), 136 (a_f) and 184 (a_c); Antinoriinae (*Antinoria*): 2 (b), 24 (c), 186 (a); Cynosurinae (*Cynosurus*): 45 (b_c); Hypseochloinae (*Hypseochloa*): 76 (d), 83 (b), 100 (b), 153 (b), 171 (b); Miliinae (*Milium*): 54 (c), 90 (b), 167 (g), 169 (a) and Phleinae (*Phleum*): 31 (c), 84 (a_b), 124 (b), 127 (b_c), 136 (a_g), 175 (c).

*Macrobriza maxima* shared a number of characters with Brizinae, the subtribe which provided one of its ancestors: 54 (a_b), 118 (a_b_c), 136 (a_f), 138 (a_c), 140 (a_b). In other characters *Macrobriza* deviated from Brizinae, for example, 24 (c), 142 (d_e), 171 (a_b), 175 (a_c), which supports its hybrid origin as disclosed by molecular phylogenetics.

### Ancestral state reconstruction

Ancestral state reconstructions applied for 74 non-molecular characters (described in suppl. Appendix S2) of the taxa of Poodae studied are visualised in suppl. Appendix S3 and, exemplarily for character 50, in Fig. 8. The ancestral character states of supertribe Poodae (tribes Aveneae and Poeae) were:

**Fig. 8.**
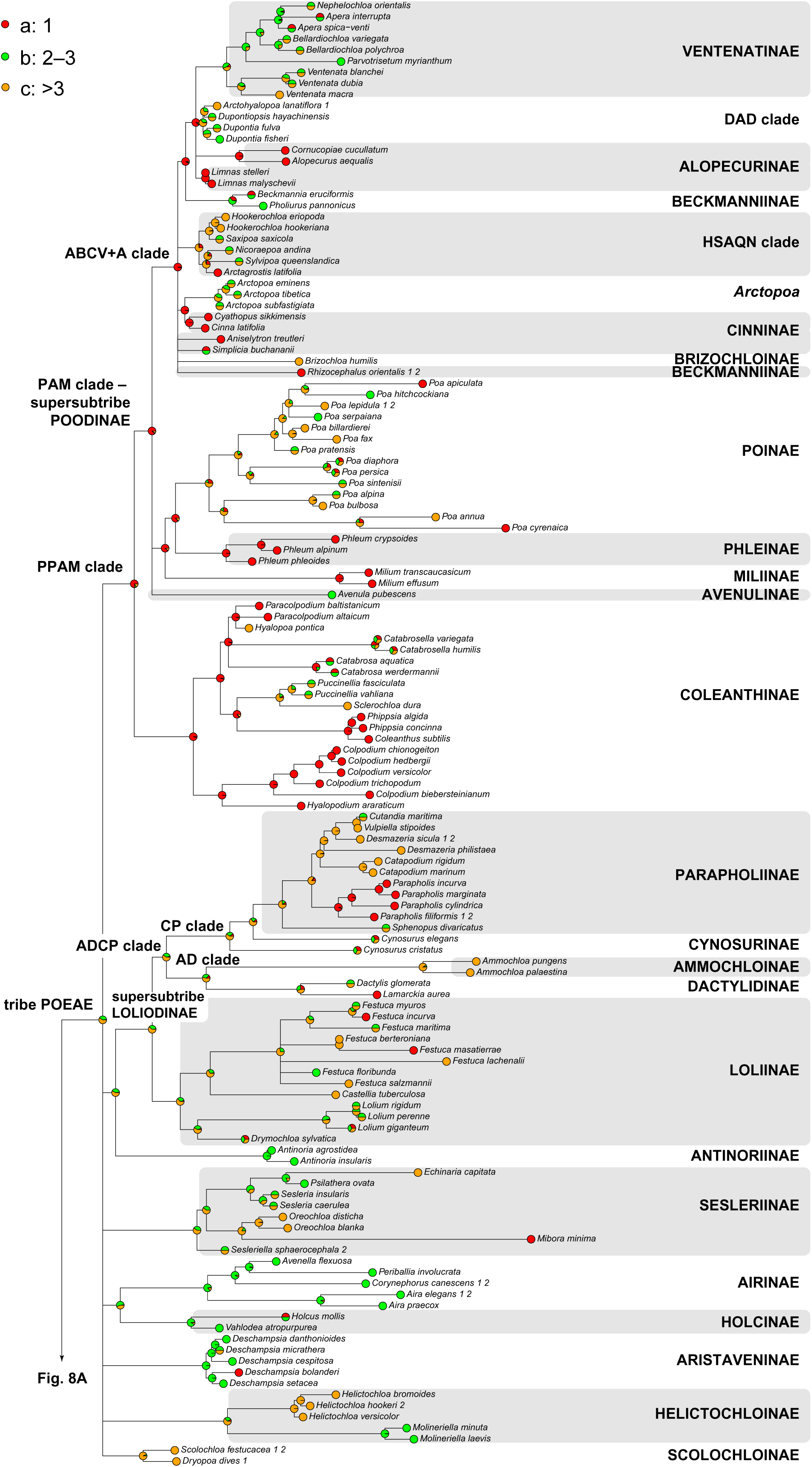
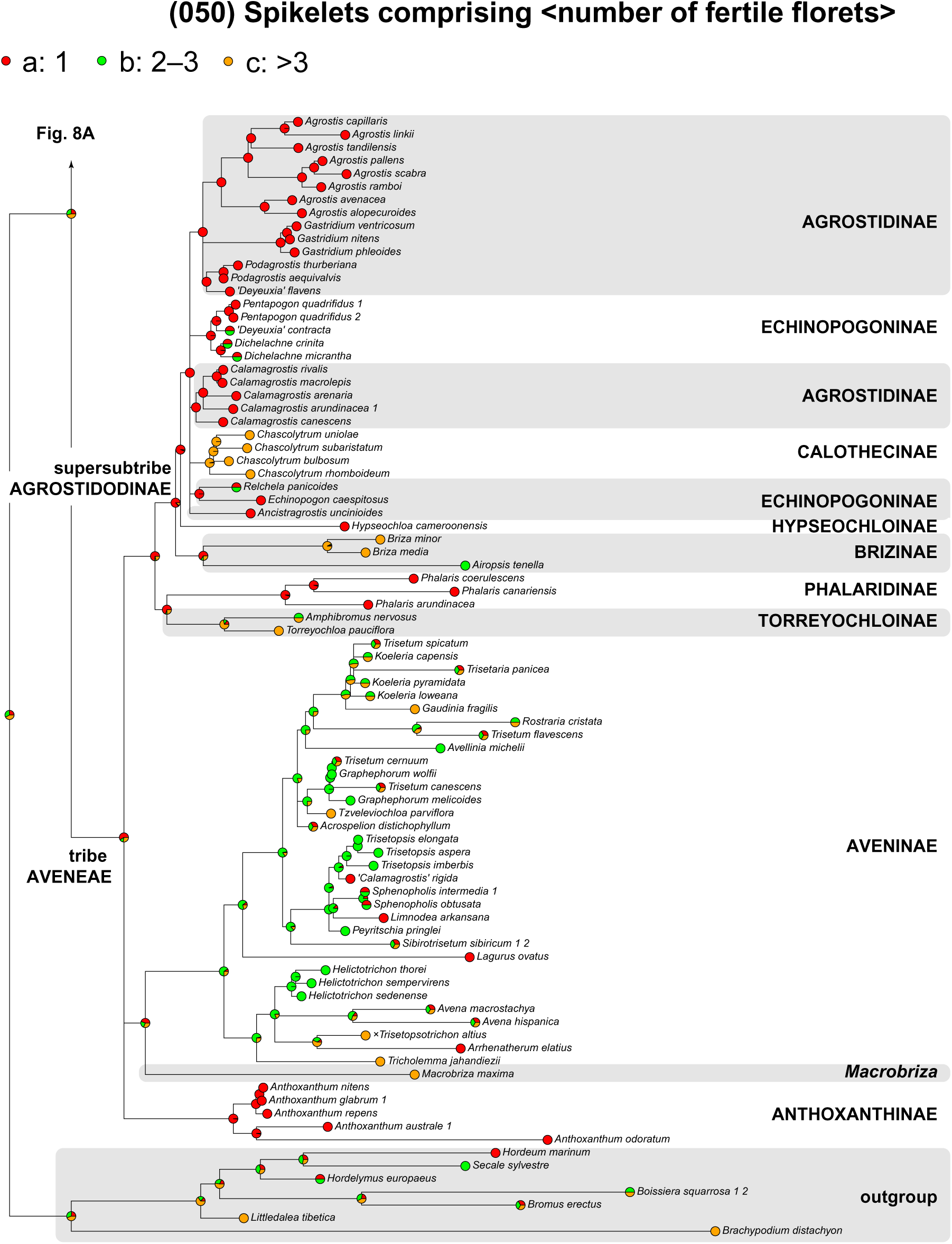
Example of an ancestral state reconstruction (ASR) in Poodae (Aveneae and Poeae) for the number of florets in the spikelets (character 50). See text (Discussion – *Ancestral state reconstruction*) and Appendix 5 for further explanation. See suppl. Appendix S3 for all ASRs conducted for 74 characters.

perennial life form (character 1) with a conspicuous transition to annual life cycle within Aveneae especially in Agrostidinae and within Poeae in Parapholiinae, most Ventenatinae, some Airinae, Loliinae and Poinae;

absence of rhizomes (character 4) with secondary development of rhizomatous species in many lineages;

ligule an eciliate membrane (character 12);

flat leaves (character 17) with secondary development of conduplicate, involute and convolute vernation in many lineages;

inflorescence an open panicle (characters 28 and 31) with a transition to contracted panicle or even more condensed, spike-like or glomerate inflorescence within Aveneae especially in Aveninae (prevalently the lineage of Koeleriinae), Phalaridinae and parts of Agrostidinae, within Poeae in Aristaveninae, Dactylidinae and Phleinae. Capitate inflorescences originated within some Poeae, namely in Sesleriinae, Ammochloinae and some members of the ABCV+A clade;

panicles with many spikelets (character 35) and reduction to few spikelets in many lineages, within Aveneae especially in some Aveninae and Agrostidinae, within Poeae in some Loliinae, many Parapholiinae and Coleanthinae;

panicle branches at most moderately divided (character 38); flexible (character 41). Stiff branches originated infrequently (e.g., in Parapholiinae), whereas capillary branches were frequent in Airinae and Coleanthinae. In Airinae, such capillary branches were likely to be the original character state, whereas in Coleanthinae flexible branches appeared to be plesiomorphic;

panicle branches smooth (character 42), whereas scaberulous or scabrous branches developed multiple times in parallel;

spikelets pedicelled (characters 46 and 47), whereas sessile spikelets developed multiple times and less frequently in Aveneae (some Aveninae, especially *Koeleria*, and Phalaridinae) than within Poeae (in Loliinae two times in parallel, CP clade, parts of Coleanthinae);

spikelets with >3 fertile florets, which had higher probability for Poodae than spikelets with 2–3 florets or 1 floret (character 50; Fig. 8). Spikelets with 1 floret had the highest probability to be ancestral in Aveneae as a whole and were most likely ancestral within Agrostidodinae and Agrostidinae, whereas spikelets with 2–3 florets were seemingly ancestral in Aveninae. Within 1-flowered Agrostidodinae, Calothecinae have seemingly secondarily developed spikelets with >3 florets. In Poeae, spikelets with >3 florets, followed by spikelets with 2–3 spikelets appeared to be the most probable ancestral state. Spikelets with 1 floret have developed secondarily within parts of Parapholiinae but, significantly, seem to be plesiomorphic in the entire PPAM clade, in which a transition to 2–3 or 3-flowered spikelets occurred secondarily in Poinae and most parts of the ABCV+A clade, except for Alopecurinae, Cinninae and some others;

spikelets with a rhachilla extension bearing a sterile florets at the apex were slightly more likely than a barren rhachilla extension or a missing rhachilla extension (character 51). For Aveneae, a barren rhachilla extension was the most likely ancestral state, which applies also for Aveninae and Agrostidinae but parts of Echinopogoninae and Agrostidinae and *Airopsis* of Brizinae showed an obviously secondary loss of the rhachilla extension, whereas absence of a rhachilla extension was the plesiomorphic character state in the anyway strongly modified spikelets of Anthoxanthinae and Phalaridinae. For Poeae, a rhachilla extension bearing a sterile floret at the apex was the most likely ancestral state and was present also in most Loliinae, Poinae and the ABVC clade. The reconstruction was ambiguous for the large PPAM clade, because this character state was equally likely to be ancestral than the absence of a rhachilla extension as found in Antinoriinae, most Coleanthinae (except *Puccinellia*/*Sclerochloa*), Miliinae, Alopecurinae, most Beckmanniinae and Cinninae;

spikelets with 2 or more fertile florets, which was likewise ancestral for both Aveneae and Poeae (character 52). Within Aveneae, spikelets with only 1 fertile floret originated in Anthoxanthinae and Phalaridinae and might be the plesiomorphic state in supersubtribe Agrostidodinae, which means that Brizinae and Calothecinae were characterized by a reversal to the ancestral state of Aveneae or Poodae as a whole with 2 or more fertile florets. Within Poeae, the transition to spikelets with only 1 fertile floret occurred infrequently in some Loliinae, some Coleanthinae (especially *Colpodium* and the lineage of *Coleanthus*/*Phippsia*), Miliinae and Phleinae. The situation was less clear in the ABCV+A clade, in which 2 or more fertile florets versus 1 fertile floret were almost equally likely as ancestral states. In the former instance, the presence of only 1 fertile floret in Alopecurinae, Beckmanniinae, Cinninae and some Ventenatinae (*Apera*) would be a derived character state;

the lowermost flower in the spikelet not male or barren (character 53);

spikelets laterally and at most moderately compressed (characters 54 and 55); terete or dorsally compressed spikelets in only few lineages (Brizinae, *Holcus*, some Loliinae, *Colpodium* p.p., Miliinae);

comparatively large spikelets of either 4–6 mm or >6 mm in length with almost equal probability. This applied also to be the ancestral state in Aveneae, whereas a length of 4–6 mm is more likely in Poeae (character 57). For Aveninae, the larger size of >6mm was most likely ancestral, with a seemingly secondary diminution in the American lineage of *Limnodea*/*Peyritschia*/*Sphenopholis* and the supersubtribe Agrostidodinae, in which especially Agrostidinae underwent a reduction to <3 mm. Secondary downsizing was likely also for Airinae, Antinoriinae, a part of Coleanthinae (*Coleanthus*, *Colpodium*, *Phippsia*) and some representatives of the ABCV+A clade, namely within Cinninae and Ventenatinae (especially *Apera* and *Parvotrisetum*);

spikelets breaking up at maturity (character 58). Spikelets falling entire appeared occasionally and in various lineages, namely Aveninae (*Gaudinia*, *Limnodea*, *Sphenopholis*), Holcinae (*Holcus*), Loliinae, Parapholiinae (*Parapholis*), and especially Cinninae, Beckmanniinae and Alopecurinae;

spikelets, which disarticulate below each floret (character 61), but with few exception in Aveninae, Parapholiinae, Alopecurinae and Ventenatinae; rhachilla internodes that are not thickened (character 63) but with sporadic exceptions in various groups; glumes persistent on branch, if spikelet breaks up (character 71) but with sporadic exceptions in Aveninae, Agrostidinae, Holcinae, Parapholiinae, Coleanthinae and the ABCV+A clade; similar glumes but with sporadic exceptions in Aveninae and Agrostidinae and especially Scolochloinae (character 72);

glumes shorter than spikelet, which applied also for tribes Aveneae and Poeae as a whole, respectively, but not for Aveninae, in which the longer glumes reaches or exceeds the length of the spikelet (character 73). The latter applied also to Agrostidinae, whereas Torreyochloinae, most Brizinae and Calothecinae have kept the ancestral state of shorter glumes. Within Poeae, there was a comparatively rare trend to longer glumes discernable, namely in Airinae, some Parapholiinae and Coleanthinae, Miliinae and few members of the ABCV+A clade (Alopecurinae, Cinninae);

glumes in consistency thinner than the lemma or similar (character 74). Both states were equally probable for Poodae, but thinner glumes were ancestral for Aveneae, whereas similarly firm glumes and lemmas are ancestral in Poeae. Within Agrostidodinae, there was a transition to firmer glumes in Calothecinae and especially Agrostidinae. Within Poeae, a similar consistency of glumes and lemmas prevailed by far, but there were firmer lemmas in Scolochloinae, Avenulinae and Miliinae, *Helictochloa* (Helictochloinae), a part of Parapholiinae, Poinae and Ventenatinae, whereas the opposite, i.e., glumes firmer than lemmas, also occurred, namely in *Parapholis*, some Loliinae and Beckmanniinae;

lower glume 3–6 mm long (character 78), with a sporadic trend to diminution, namely within Aveneae in Aveninae (especially Koeleriinae clade A; Fig. 5) and Calothecinae, within Poeae in *Molineriella* (Helictochloinae), Aristaveninae, parts of Airinae, Sesleriinae and Ventenatinae (*Apera*, *Bellardiochloa* Chiov., *Nephelochloa*), in Antinoriinae and Coleanthinae. The opposite, namely enlargement of glumes, was infrequently found. Examples were some Aveninae and Agrostidinae (*Calamagrostis* s.str.), South American members of Loliinae and a few members of the ABCV+A clade;

lower glume 0.6-fold to as long as the upper, with shortening as well as enlargement sporadically encountered in various groups (character 79); of similar consistency on margins or margins much thinner in several groups (character 81); 1-keeled, with sporadic transition to unkeeled shape in various groups (character 82); keeled all along, with sporadic transition to keeled only above or below in various groups (character 83); 1-veined, with rare transition to either veinless (few Coleanthinae) and infrequent to ≥3-veined, namely in some Aveninae, Phalaridinae, Brizinae, Calothecinae, *Helictochloa* (Helictochloinae), Antinoriinae, most Parapholiinae, Miliinae, Phleinae and the majority of Ventenatinae, in which this character state might have even been plesiomorphic (character 85); primary vein eciliate (character 87); without lateral veins and infrequent presence of distinct lateral veins in Anthoxanthinae, some Aveninae, Calothecinae, *Helictochloa* (Helictochloinae), some Loliinae and the majority of Ventenatinae, in which this character state might have been plesiomorphic (character 88), are the ancestral states, respectively;

upper glume 3–6 mm long (character 95), with occasional diminution comparable to the lower glume (character 78) occurring within Aveneae in some Aveninae, Calothecinae and Agrostidinae, within Poeae in *Molineriella* (Helictochloinae), parts of Airinae, Sesleriinae and Ventenatinae (*Apera*, *Bellardiochloa*, *Nephelochloa*) as well as in Antinoriinae and Coleanthinae. Enlargement of glumes occurred in most Aveninae, in which this character appeared to be the plesiomorphic state, within Poeae in *Helictochloa* (Helictochloinae), South American members of Loliinae (*Festuca berteroniana*, *F. floribunda*, *F. masatierrae*) and some members of the ABCV+A clade (*Arctopoa*, *Hookerochloa* E.B.Alexeev, *Pholiurus*, etc.).

uncertain considering the length relation of upper glume and adjacent lemma (character 96). The upper glume shorter than the lemma was the most likely ancestral state in Aveneae and Aveninae, in which most of the Koeleriinae lineage, few members of the Aveninae s.str. lineage, Calothecinae and some Echinopogoninae showed a secondary transition to shorter glumes. The upper glume shorter than the adjacent lemma was, by contrast, the most likely ancestral state in Poeae, with a secondary change to longer glumes in Holcinae, Airinae, Antinoriinae, *Parapholis* (Parapholiinae), Avenulinae, Miliinae, some Beckmanniinae and Ventenatinae;

upper glume with undifferentiated margins (character 98) and sporadic transition to hyaline, membranous or scarious margins in various lineages of both Aveneae as well as Poeae; 1-keeled, with sporadic transition to unkeeled shape in various groups (character 99); keeled all along, with sporadic transition in various lineages to keeled only below (some Coleanthinae) or above (character 100); 3-veined, with transition to 1-veined within Aveneae in most Agrostidinae (probably plesiomorphic in this subtribe), within Poeae in Sesleriinae, Ammochloinae, some Coleanthinae; with transition to ≥5-veined within Aveneae in some Aveninae, Phalaridinae, Brizinae, within Poeae in some Helictochloinae (partly in *Helictochloa*), Loliinae and sporadically within the ABCV+A clade (character 102); primary vein distinct (character 103); primary vein smooth but in several lineages transition to scaberulous or scabrous, especially within Aveneae in a part of Aveninae, namely Koeleriinae clade A (Fig. 5), in Phalaridinae, Brizinae, Hypseochloinae, Echinopogoninae, Agrostidinae, and within Poeae only in Antinoriinae, Ammochloinae, Poinae and most of the ABCV+A clade except for the DAD clade (character 104); lateral veins distinct but absent in some Echinopogoninae, Agrostidinae, Sesleriinae, sporadically in Loliinae and Coleanthinae (character 106); upper glume muticous (absence of awns), but seemingly secondarily mucronate or awned in several lineages, namely in Aveninae, Echinopogoninae (*Pentapogon*), Sesleriinae, some Loliinae, Parapholiinae and Phleinae (character 114);

spikelets without basal sterile florets (character 115) but present in Anthoxanthinae and Phalaridinae and, as a rare exception, in Aveninae (*Arrhenatherum*); fertile florets (if more than 1) all alike, with occasional exceptions, especially Holcinae and *Ventenata* (character 116);

lemma 1.6–4 mm, which was slightly more likely than >4mm long (character 119), which applied also for both Aveneae and Poeae. Nevertheless, the latter is the ancestral state in Aveninae (Figs. 9E,M,N, 10I, 11I,K,L) most likely also in Echinopogoninae (Fig. 11C; all Aveneae) and, within Poeae in Scolochloinae, probably in Helictochloinae (Fig. 10H), Aristaveninae (Fig. 9H), Loliinae (Fig. 11A), and some lineages within the ABCV+A clade (*Arctopoa*, HSAQN clade and *Ventenata*; Figs. 9K,11H). A diminution to 1.5 mm was found within Helictochloinae, in which it occurred apparently secondarily in *Molineriella* and within Coleanthinae, namely in *Coleanthus* and *Phippsia*;

**Fig. 9.**
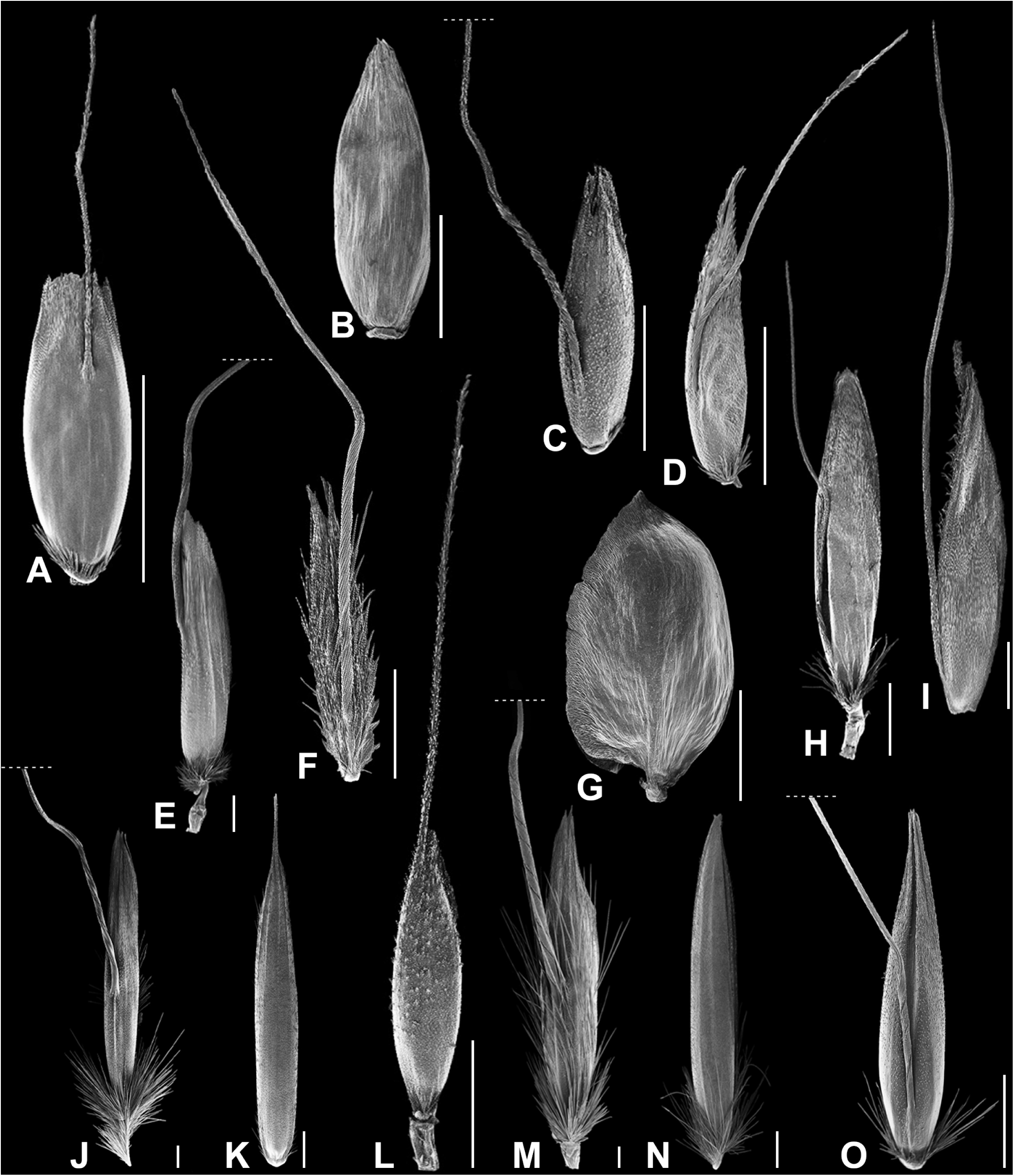
Lemmas and awns (partly trimmed) in species of Aveneae and Poeae. Scanning electron microphotographs: **A**, *Agrostis avenacea* (*M. Röser 10762*, HAL); **B**, *A. capillaris* (*M. Röser 11296 & N. Tkach*, HAL0140613); **C,** *A. rupestris* (*M. Röser 11312* & *N. Tkach*, HAL0144916); **D,** *Aira praecox* (*M. Röser 11041*, HAL); **E,** *Amphibromus nervosus* (*M. Röser 10770*, HAL); **F,** *Anthoxanthum odoratum* (*M. Röser 11006*, HAL); **G,** *Briza media* (*M. Röser 11072*, HAL); **H,** *Avenella flexuosa* (*M. Röser 11202 & N. Tkach*, HAL0141248); **I,** *Alopecurus pratensis* (*M. Röser 11222 & N. Tkach*, HAL0141246); **J,** *Avenula pubescens* (*M. Röser 6528*, HAL); **K,** *Beckmannia eruciformis* (*s.coll. R382*, HAL); **L,** *Apera spica-venti* (*M. Röser 10699*, HAL0140288); **M,** *Avena fatua* (N. *Röser 11267 & N. Tkach*, HAL0140638); **N,** *Calamagrostis arenaria* (*M. Röser 11291*, HAL0144749); **O,** *Calamagrostis arundinacea* (*M. Röser 11274 & N. Tkach*, HAL0141306). **A, B, F, G, J, K, L, M & O,** Dorsal view; **C, D, E, H, I, N,** Lateral view. — Scale bars = 1 mm.

**Fig. 10.**
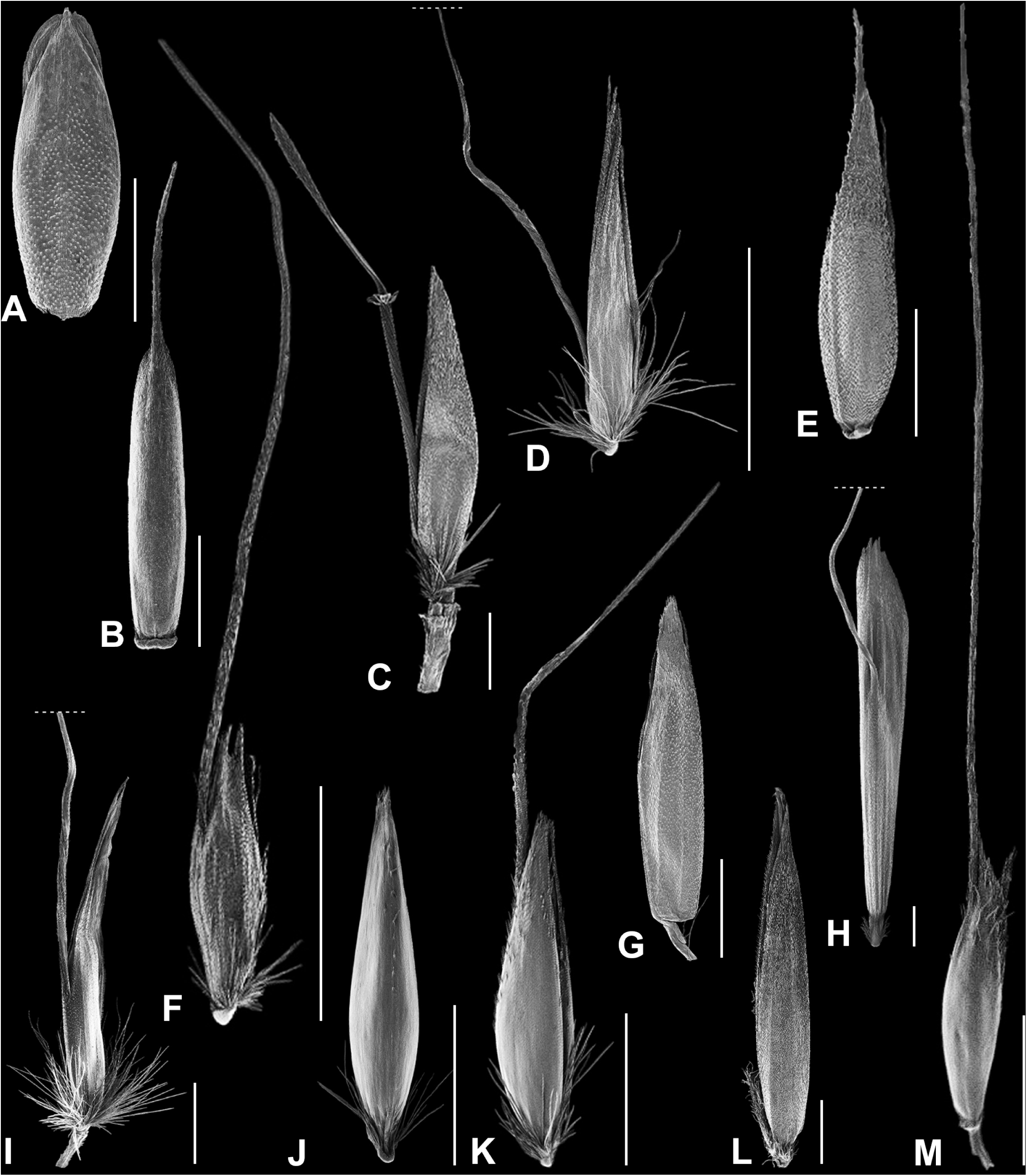
Lemmas and awns (partly trimmed) in species of Aveneae and Poeae. Scanning electron microphotographs: **A,** *Catapodium marinum* (*M. Röser 4352*, HAL); **B,** *Festuca lachenalii* (*M. Röser 5470*, HAL); **C,** *Corynephorus canescens* (*E. Willing 25.870D*, HAL0108617); **D,** *‘Calamagrostis’ flavens* (*I. Hensen*, HAL); **E,** *Cynosurus cristatus* (*M. Röser 622*, HAL); **F,** *Gastridium nitens* (*G. van Buggenhout 11991*, ROM); **G,** *Graphephorum wolfii* (*R.J. Soreng*, NY); **H,** *Helictochloa bromoides* subsp. *bromoides* (*M. Röser 10519*; HAL); **I,** *Helictotrichon petzense* subsp. *petzense* (*M. Röser 10646*, HAL); **J & K,** *Holcus mollis* (*M. Röser 10658,* HAL) with lower (**J**) and upper lemma (**K**); **L,** *Hookerochloa eriopoda* (*R. Pullen 4003*, AD96435171); **M,** *Lamarckia aurea* (*M. Röser 311*, HAL). **A, B, H, J,** Dorsal view; **C, D, E, F, G, I, K, L, M,** Lateral view. — Scale bars = 1 mm.

**Fig. 11.**
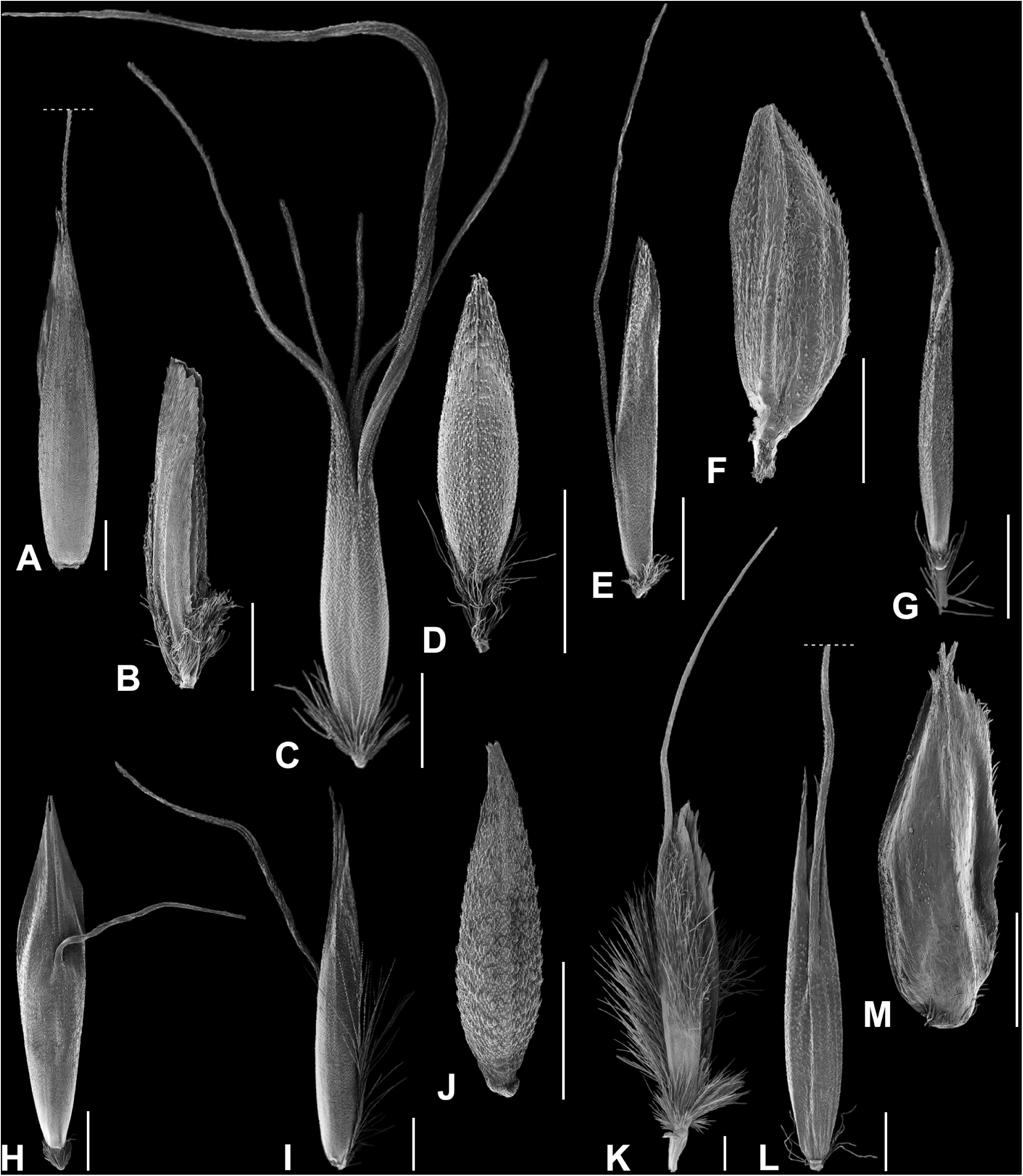
Lemmas and awns (partly trimmed) in species of Aveneae and Poeae. Scanning electron microphotographs: **A,** *Lolium giganteum* (*M. Röser 11275 & N. Tkach*, HAL0141305); **B,** *Poa fax* (*D.E. Murfet 1278*, AD99151120); **C,** *Pentapogon quadrifidus* (*A. Moscal 11543*, HO 95925); **D,** *Periballia involucrata* (*s. coll., R58/R319*, HAL); **E,** *Peyritschia pringlei* (*P. Tenorio 15095*, MEXU542571); **F,** *Phleum crypsoides* (*F. Skovgaard*, C); **G,** *Trisetum flavescens* (*M. Röser 11245 & N. Tkach*, HAL0141270); **H,** *Ventenata macra* (*M. Röser 10688*, HAL); **I,** *Tzveleviochloa parviflora* (*S. & G. Miehe & K. Koch 01-073-24*, Institute of Geography, University Marburg, Germany); **J,** *Simplicia buchananii* (*A.P. Druce*, CHR 394262); **K,** *Tricholemma jahandiezii* (*M. Röser 10297*, HAL); **L,** *Trisetopsis elongata* (*S. Wagner R118b*, HAL0144713); **M,** *Sesleria caerulea* (*M. Röser 11239 & N. Tkach*, HAL0141254). **A, D, G, H, J, L, M**, Dorsal view; **B, C, E, F, I, K,** Lateral view. — Scale bars = 1 mm.

lemma without keel (character 122), which was the ancestral state also for both Aveneae and Poeae but transition to keeled occurred within Aveneae in Koeleriinae clade A (Figs. 5, 10G), Phalaridinae and Calothecinae, within Poeae in parts of Loliinae and Parapholiinae, in the PPAM and, even more, in the PAM clade (∼supersubtribe Poodinae), in which keeled lemmas (Fig. 11B,F,L) were seemingly the ancestral state. This implies that the unkeeled lemmas in parts of Beckmanniinae and Ventenatinae (Figs. 9K,11H) were most likely a reversal within the PAM clade to the original character state of the whole Poeae and Poodae;

lemma with 4–5 veins (characters 125, 126), which was the ancestral state also of both Aveneae and Poeae but transition to >5 veins occurred within Aveneae in Aveninae (in Aveninae s.str. and parts of Koeleriinae), Hypseochloinae and parts of Brizinae, within Poeae in Helictochloinae, Ammochloinae, Phleinae and parts of Loliinae. Diminution to 1–3 veins such as found in some Aveninae (parts of Koeleriinae), Agrostidinae, Parapholiinae, Coleanthinae and the DAD clade was encountered more rarely;

lemma surface generally rough (character 133) with sporadic exceptions, such as in Sesleriinae (Fig. 11M) and others (Figs. 9C,L, 10A, 11D,F,J,L); general extent of hairiness all along (character 134; Figs. 9F,M, 11K) with occasional secondary restriction to hairiness above (Fig. 10E,L) or in the middle to below (frequent in many lineages);

lemma apex erose or dentate (character 143). Entire lemma apices occurred occasionally in various groups but were frequent in Loliinae and especially in supersubtribe Poodinae (except for *Avenula*; Fig. 9J), in which it seemed to be the plesiomorphic state;

lemma apex mucronate to awned (character 146), which was the ancestral state also of both Aveneae and Poeae. Within Aveneae, there were occasional transitions to muticous apices (for example, Anthoxanthinae, few Aveninae, Agrostidinae; Fig. 9A, 10G). Within Poeae, muticous apices seemed to be plesiomorphic for the PPAM clade (Fig. 11B) with a reversal to mucronate/awned lemma apices in some Poinae and members of the ABCV+A clade such as Alopecurinae (Fig. 9I) and especially Ventenatinae (Figs. 9L, 11H);

apical awns were the ancestral state of both Aveneae and Poeae (character 148). Within Aveneae, the characteristic dorsal awns of Aveninae (Figs. 9M, 10I, 11I,K,L) and Agrostidinae (Figs. 9A,C,O, 10D,F) as well as of Hypseochloinae (*Hypseochloa*) and *Amphibromus* (Torreyochloinae; Fig. 9E) seemingly were derived secondarily and in parallel, moreover, this seemed to be the plesiomorphic state of Aveninae and Agrostidinae. Within Poeae, the dorsal awns occurred secondarily in Helictochloinae (Fig. 10H), Aristaveninae (Fig. 9H), Holcinae (Fig. 10K), Airinae (Figs. 9D, 10C), Alopecurinae (Fig. 9I) and some Ventenatinae (Fig. 11H). Due to the unresolved relationships of the former four subtribes, it remains unclear, if this transition occurred only once in their putative common ancestor or multiple times in parallel. Considering Ventenatinae, the occurrence of the apical awns present in *Apera* (Fig. 9L), *Bellardiochloa* and *Nephelochloa* might have been a character reversal to the ancestral state in Poeae and Poodae, if dorsal awns had been the plesiomorphic character state for Ventenatinae;

principal lemma awn (if present) straight, which was the ancestral state also for both Aveneae and Poeae (character 150), and secondary transition to geniculately bent shape within Aveneae in Aveninae (Figs. 9M, 10D,F,I, 11E,G,I,K,L; note a reversal to straight awns in parts of the Koeleriinae lineage), in Torreyochloinae (*Amphibromus*; Fig. 9E), Hypseochloinae, Echinopogoninae (Fig. 11C) and Agrostidinae (Fig. 9A,C,O), within Poeae in Helictochloinae (Fig. 10H), Airinae (Figs. 9D,H, 10C; with an apparent reversal in awned specimens of *Periballia* Trin.), Avenulinae (Fig. 9J) and some members of Ventenatinae (Fig. 11H); geniculate awns mostly clearly exserted from spikelets, whereas straight awns were not or scarcely exserted (character 154);

principal lemma awn not coloured, whereas coloured awns seemed to occur sporadically and secondarily, namely within Aveneae in parts of Aveninae, Echinopogoninae and Agrostidinae, within Poeae in Helictochloinae, Airinae, more rarely in Loliinae and Poinae (character 156);

absence of a distinct column of the lemma awn, which was the ancestral state also for Aveneae and Poeae; acquisition of distinct columns (character 155) was seemingly linked with the origin of dorsal awns (as described before). Distinctness of columns was secondarily lost especially in parts of the Aveninae, namely some members of the Koeleriinae lineage, and Agrostidinae;

column of lemma awn not twisted, which was the ancestral also for both Aveneae and Poeae (character 160), whereas twisted columns originated within Aveneae in Aveninae (Figs. 9M,O, 10D,F,I, 11E,G,I,K,L) for which they were seemingly plesiomorphic, including a reversal to untwisted columns in some members of the Koeleriinae lineage and some Agrostidinae, in which twisted columns (Figs. 9A,C) were most likely plesiomorphic. Twisted columns evolved within Poeae also repeatedly in parallel, namely within Helictochloinae (Fig. 10H), Aristaveninae, Airinae (Fig. 9D,H), *Avenulinae* (Fig. 9J), some Alopecurinae (Fig. 9L) and Ventenatinae (*Ventenata*, *Parvotrisetum*; Fig. 11H);

column of lemma awn (if present) transversally not flattened (character 159), whereas flattened columns occurred sporadically, namely in some Aveninae, Agrostidinae (Fig. 9C), Helictochloinae (*Helictochloa*; Fig. 10H), Alopecurinae (*Alopecurus*; Fig. 9I) and Ventenatinae (*Ventenata*; Fig. 11H);

lateral lemma awns absent (character 161). Lateral lemma awns originated occasionally but not consistently in various lineages, e.g., Aveninae (Figs. 10I, 11I), Echinopogoninae (Fig. 11C), Sesleriinae (Fig. 11M), Coleanthinae, Poinae and the ABCV+A clade;

palea ≥0.7-fold longer than the lemma (character 166) and with smooth keels (character 172). Shorter or missing paleas as well as scaberulous or scabrous palea keels originated sporadically and inconsistently in many lineages;

eciliate palea keels, which was the ancestral state also for both Aveneae and Poeae individually (character 173). Puberulous, pubescent, ciliolate or ciliate palea keels originated in many lineages, within Aveneae especially in Aveninae, in which this might be the plesiomorphic character state of Aveninae s.str., within Poeae especially in Sesleriinae, Poinae and some Coleanthinae;

apical sterile florets (if present) resembling fertile though underdeveloped (character 179). Sterile florets variously modified and distinct from fertile were characteristic of Anthoxanthinae, in which they represented the plesiomorphic state, but originated secondarily within various other lineages of Aveneae, namely sporadically in Aveninae, Agrostidinae and Echinopogoninae, within Poeae in Helictochloinae, Aristaveninae, Airinae, Coleanthinae, in which this character represented most likely the plesiomorphic state, and the ABCV+A clade;

stamens 3, with reduction in number sporadically encountered in various lineages of both Aveneae and Poeae (character 182); anthers ≥1 mm with diminution of size likewise in various lineages of both tribes and presumably frequently related with self-pollination (character 183);

caryopsis ≥1.6 mm long, which was the ancestral state also for both Aveneae and Poeae (character 187), whereas shorter caryopses originated within Aveneae especially in Agrostidinae, for which they were seemingly plesiomorphic, but also in some Torreyochloinae, Phalaridinae, Brizinae, Hypseochloinae and Calothecinae and thus might be plesiomorphic for the common lineage comprising the latter subtribes. Shorter caryopses originated more sporadically also in Aveninae. Within Poeae, they originated infrequently, especially in Sesleriinae, Coleanthinae, Phleinae, Poinae, and a part of the ABCV+A clade, but sporadically also in Loliinae, Parapholiinae, etc.

hilum linear and straight, which was the ancestral state for also for both Aveneae and Poeae individually (character 188). A transition to short, elliptic or punctiform hila occurred within Aveneae especially in the Koeleriinae lineage of Aveninae, in Calothecinae and some Agrostidinae. Within Poeae it seemed to be quite characteristic of Airinae, Sesleriinae, Ammochloinae, Coleanthinae but occurred more sporadically also in parts of Parapholiinae and the ABCV+A clade.

### Classification

Using the molecular phylogenetic data of both DNA analyses (plastid, nuclear) it can be concluded that the main bifurcation of the plastid DNA tree backbone is not reflected in the nuclear tree, which, however, does not provide a supported alternative topology. In this regard, the nuclear tree is un-informative and does not contribute to answer the question why classification should not use the supported plastid DNA lineages to re-instate tribes Aveneae and Poeae instead of acknowledging an enlarged Poeae (Poeae s.l.) as done in most recent classifications (Kellogg, 2015; Soreng & al., 2015, 2017; Saarela & al., 2017). All of these classifications actually make use of an informal arrangement of the subtribes in two groups within Poodae or Poeae s.l. according to the chloroplast DNA types (Soreng & Davis, 2000; Davis & Soreng, 2007; Döring & al., 2007; Quintanar & al., 2007; Soreng & al., 2007; Döring, 2009; Schneider & al., 2009, 2012; Saarela & al., 2010, 2017, 2018; Pimentel & al., 2017; Orton & al. 2019).

We suppose (1) that the consistent occurrence of two clearly differentiated chloroplast DNA lineages without intermediates reflects a major evolutionary differentiation.

The absence (2) of equivalent differentiation in the nuclear DNA of these plants does make it impossible to use the plastid DNA results for classification. It should be noted in this context that supported backbone structure was absent also in several nuclear single copy gene trees we have studied (J. Schneider, unpub. data) and not only in the repetitive rDNA tree used in this study.

The occurrence of hybridization (3) between the two different plastid DNA lineages as documented by several instances does not make a classificatory recognition of two tribes impossible. Morphologically, the two tribes are not clearly defined, which holds true, however, also for many of their subtribes. All in all, there is obviously a high degree of homoplasy in many morphological characters as seen, for example, in traditionally highly ranked characters for classification such as the presence of a dorsal lemma awn or long glumes in relation to the entire spikelet as presumably typical of Aveneae. Comparable difficulties in underlining classification by morphology are encountered also in many other grass groups. Examples would be the delineation of Stipeae within subfamily Pooideae, whether or not including *Ampelodesmos* Link as sole genus with several-flowered spikelets (see Schneider & al., 2011; Kellogg, 2015; Soreng & al., 2017), or the vague morphological circumscription of subfamily Micrairoideae (Sánchez-Ken & al., 2007; Kellogg, 2015).

The proposed modified classification uses narrowly delineated and preferably monophyletic subtribes as applied in most recent treatments of the study group (e.g., Soreng & al. (2007, 2015, 2017); Kellogg, 2015; Saarela & al. 2017):

Supertribe Poodae L.Liu: (1) tribe Aveneae Dumort.: subtribes Aveninae J.Presl, Anthoxanthinae A.Gray, Torreyochloinae Soreng & J.I.Davis, Phalaridinae Fr., Brizinae Tzvelev, Hypseochloinae Röser & Tkach, Echinopogoninae Soreng, Calothecinae Soreng, Agrostidinae Fr.; (2) tribe Poeae R.Br.: subtribes Scolochloinae Tzvelev, Aristaveninae F.Albers & Butzin, Helictochloinae Röser & Tkach, Holcinae Dumort., Airinae Fr., Sesleriinae Parl., Antinoriinae Röser & Tkach, Loliinae Dumort., Ammochloinae Tzvelev, Dactylidinae Stapf, Cynosurinae Fr., Parapholiinae Caro, Coleanthinae Rouy, Avenulinae Röser & Tkach, Miliinae Dumort., Phleinae Dumort., Poinae Dumort., Brizochloinae Röser & Tkach, Cinninae Caruel, Beckmanniinae Nevski, Alopecurinae Dumort., Ventenatinae Holub ex L.J.Gillespie, Cabi & Soreng.

## NEW NAMES AND COMBINATIONS

**Antinoriinae** Röser & Tkach, **subtribus nov.** – Type: *Antinoria* Parl., Fl. Palerm. 1: 92. 1845.

*Description.* – Annual or rarely (in *A. agrostidea*) perennial, caespitose or decumbent; leaf sheath margins free, leaf blades flat; ligule an unfringed membrane, 1–3 mm long; inflorescence paniculate; spikelets pedicellate, laterally compressed, 1–2 mm long, with 2 florets, disarticulating above the glumes and between the florets, with distinctly elongated rhachilla internode between the florets, glabrous, terminated by a female-fertile floret; glumes relatively large, more or less equal, exceeding the spikelets, awnless, carinate, 3-nerved; lemmas elliptic, widest near the tip, membranous, incised or blunt, awnless, glabrous, 5-nerved; palea relatively long, tightly clasped by the lemma, 2-nerved, 2-keeled; anthers 0.5–1 mm long; ovary glabrous; caryopsis pyriform, compressed dorsiventrally, smooth; hilum short; embryo less than 1/3 as long as fruit.

*Included genus.* – *Antinoria*.

*Distribution.* – Mediterranean.

**Avenulinae** Röser & Tkach, **subtribus nov. –** Type: *Avenula* (Dumort.) Dumort., Bull. Soc. Roy. Bot. Belgique 7: 68. 1868.

*Description*. – Perennial, loosely caespitose, with creeping underground shoots; roots without sclerenchyma surrounding endodermis; culms with 1–3 visible nodes. Leaf sheaths closed over more than 1/2 their length from base; leaf blades flat or ± conduplicate, not furrowed, relatively soft but rigid, with long hairs; bulliform cells forming a row each side of the adaxial midrib; with abaxial midrib and margins scarcely evident; secondary nerves few; well-developed subepidermal sclerenchyma forming O-shaped girders at lateral nerves; inflorescence lax panicle; spikelets 14–20 mm long, with 3–4 developed bisexual florets, two upper floret not or scarcely exceeding the upper glume, apical floret reduced; glumes unequal, keeled on the back, somewhat scabrid on the central nerve at the base, the lower glume 1-3-nerved, the upper glume 3-nerved; rhachilla disarticulating above the glumes and between the florets; lemmas glabrous (except for the callus); dorsally awned, with a strongly twisted, rounded column, without pale margins; palea scarcely 2-keeled, with glabrous and smooth keels; lodicules as long or shorter than the ovary, ovate or obovate, 2–3-lobed or with a irregularly dentate apex; caryopsis furrowed; hilum linear; embryo with a truncated epiblast and obtuse scutellum.

*Included genus.* – *Avenula*.

*Distribution*. – Europe to eastern Siberia, Caucasus, northern Central Asia, Mongolia.

**Brizochloinae** Röser & Tkach, **subtribus nov. –** Type: *Brizochloa* Jirás. & Chrtek, Novit. Bot. Delect. Seminum Horti Bot. Univ. Carol. Prag. (1966). 40. 1966.

*Diagnosis.* – Differs from Brizinae and *Macrobriza* by upright pedicels of the spikelets, slightly scabrous rhachillas and non-cordate lemmas.

*Included genus.* – *Brizochloa*.

*Distribution.* – Eastern Mediterranean to Caucasus and Iran.

**Helictochloinae** Röser & Tkach, **subtribus nov.** – Type: *Helictochloa* Romero Zarco, Candollea 66: 96. 2011.

*Description.* – Perennial (*Helictochloa*) or annual (*Molineriella*); leaf sheaths split almost up to base, leaf blades flat, conduplicate or convolute; inflorescence lax panicle to (sometimes in *Helictochloa*) raceme-like; spikelets 10–36 mm (*Helictochloa*) or 1.5–2.5 mm (*Molineriella*), with (2–)3-9(–12) (*Helictochloa*) or 2 (*Molineriella*) developed, bisexual florets; glumes shorter than spikelets, the lower glume with (1-)3-5 (*Helictochloa*) or 1 (*Molineriella*) nerves, the upper glume with 3-5(-7) (*Helictochloa*) or 3 (*Molineriella*) nerves; rhachilla disarticulating above the glumes and between the florets; lemmas glabrous or sericeous towards the base, awned dorsally in the half (*Helictochloa*) or in upper 1/3 of the lemma or awnless (*Molineriella*); awn with a loosely twisted column and a long subula (*Helictochloa*) or straight, extending by more than 10 mm (*Helictochloa*) or by 0.3-0.6 mm (*Molineriella*) beyond the lemma apex; palea 2-keeled, keels minutely ciliate (*Helictochloa*) or almost smooth (*Molineriella*); lodicules lanceolate, with a lateral lobe.

*Included genera.* – *Helictochloa*, *Molineriella*.

*Distribution.* – Mediterranean, Eurasia, North America.

**Hypseochloinae** Röser & Tkach, **subtribus nov.** – Type: *Hypseochloa* C.E.Hubb., Bull. Misc. Inform. Kew 1936: 300, Fig. 1 (1936).

*Diagnosis.* – Differs from Airinae by 1-flowered spikelets, 5-nerved glumes (the upper rarely 3-nerved), an apically deeply bifid lemma (about 1/3 incised), which is crustaceously indurated at maturity, the awn arising from the apical sinus.

*Included genus.* – *Hypseochloa*.

*Distribution.* – Cameroon Mt. and Tanzania.

***Anthoxanthum glabrum*** (Trin.) Veldkamp subsp. ***sibiricum*** (Tzvelev) Röser & Tkach, **comb. nov.** ≡ *Hierochloe odorata* (L.) P.Beauv. subsp. *sibirica* Tzvelev, Novosti Sist. Vyssh. Rast. 1968: 21. 1968.

***Anthoxanthum nitens*** (Weber) Y.Schouten & Veldkamp subsp. ***kolymensis*** (Prob.) Röser & Tkach, **comb. nov.** ≡ *Hierochloe odorata* (L.) P.Beauv. subsp. *kolymensis* Prob., Novosti Sist. Vyssh. Rast. 15: 69. 1979.

***Arctohyalopoa*** Röser & Tkach, **gen. nov.** – Type: *Poa lanatiflora* Roshev., Izv. Bot. Sada Akad. Nauk S.S.S.R. 30: 303. 1932 ≡ *Arctohyalopoa lanatiflora* (Roshev.) Röser & Tkach

*Description*: Differs from *Hyalopoa* by lemmas with copious and long hairs on the basal half and especially on nerves, calli copiously covered with long crinkly hairs and glabrous paleas with rarely a few hairs along keels.

***Arctohyalopoa lanatiflora*** (Roshev.) Röser & Tkach, **comb. nov.** ≡ *Poa lanatiflora* Roshev., Izv. Bot. Sada Akad. Nauk S.S.S.R. 30: 303. 1932.

***Arctohyalopoa lanatiflora*** subsp. ***ivanoviae*** (Malyschev) Röser & Tkach, **comb. nov.** ≡ *Colpodium ivanoviae* Malyschev, Novosti Sist. Vyssh. Rast. 7: 295. 1971 [1970 publ.

1971].

***Arctohyalopoa lanatiflora*** subsp. ***momica*** (Tzvelev) Röser & Tkach, **comb. nov.** ≡ *Colpodium lanatiflorum* Tzvelev subsp. *momicum* Tzvelev, Fl. Arct. URSS 2: 172. 1964.

***Colpodium biebersteinianum*** (Claus) Röser & Tkach, **comb. nov.** ≡ *Agrostis biebersteiniana* Claus, Beitr. Pflanzenk. Russ. Reiches 8: 264. 1851 ≡ *Zingeria biebersteiniana* (Claus) P.A.Smirn., Byull. Moskovsk. Obshch. Isp. Prir., Otd. Biol. 51: 67. 1946 ≡ *Zingeria trichopoda* subsp. *biebersteiniana* (Claus) Doğan, Notes Roy. Bot. Gard. Edinburgh 40: 86. 1982.

***Colpodium kochii*** (Mez) Röser & Tkach, **comb. nov.** ≡ *Milium kochii* Mez, Notes Roy. Bot. Gard. Edinburgh 17: 211. 1921 ≡ *Zingeria kochii* (Mez) Tzvelev, Bot. Zhurn. (Moscow & Leningrad) 50: 1318. 1965.

***Colpodium pisidicum*** (Boiss.) Röser & Tkach, **comb. nov.** ≡ *Agrostis pisidica* Boiss., Ann. Sci. Nat., Bot., sér. 4, 2: 255. 1854 ≡ *Zingeria pisidica* (Boiss.) Tutin, Bot. J. Linn. Soc. 76: 365. 1978.

***Colpodium trichopodum*** (Boiss.) Röser & Tkach, **comb. nov.** ≡ *Zingeria trichopoda* (Boiss.) P.A.Smirn., Byull. Moskovsk. Obshch. Isp. Prir., Otd. Biol. 51: 67. 1946 ≡ *Zingeria biebersteiniana* subsp. *trichopoda* (Boiss.) R.R.Mill, Fl. Turkey 9: 365. 1985.

***Colpodium verticillatum*** (Boiss. & Balansa) Röser & Tkach, **comb. nov.** ≡ *Milium verticillatum* Boiss. & Balansa, Bull. Soc. Bot. France 5: 169. 1858 ≡ *Zingeria verticillata* (Boiss. & Balansa) Chrtek, Novit. Bot. Delect. Seminum Horti Bot. Univ. Carol. Prag. 1963: 3. 1963 ≡ *Zingeriopsis verticillata* (Boiss. & Balansa) Prob., Novosti Sist. Vyssh. Rast. 14: 12. 1977.

***Deschampsia micrathera*** (É.Desv.) Röser & Tkach, **comb. nov.** ≡ *Trisetum micratherum* É.Desv., Flora Chilena [Gay] 6: 352. 1854 ≡ *Leptophyllochloa micrathera* (É.Desv.) C.E.Calderón ex Nicora, Fl. Patagonica 3: 70. 1978.

***Dupontia fulva*** (Trin.) Röser & Tkach, **comb. nov.** ≡ *Poa fulva* Trin., Mém. Acad. Imp. Sci. St.-Pétersbourg, Sér. 6, Sci. Math. 1: 378. 1830.

***Festuca masafuerana*** (Skottsb. & Pilg. ex Pilg.) Röser & Tkach, **comb. nov.** ≡ *Bromus masafueranus* Skottsb. & Pilg. ex Pilg., Repert. Spec. Nov. Regni Veg. 16: 385. 1920 ≡ *Megalachne masafuerana* (Skottsb. & Pilg. ex Pilg.) Matthei

*Festuca masatierrae* Röser & Tkach, **nom. nov.**

*Replaced synonym*. – *Podophorus bromoides* Phil., Bot. Zeitung (Berlin) 14: 649. 1856.

*Blocking name*. – *Festuca bromoides* L., Sp. Pl. 1: 75. 1753.

***Hyalopodium*** Röser & Tkach, **gen. nov.** – Type: *Catabrosa araratica* Lipsky, Trudy Imp. S.-Peterburgsk. Bot. Sada 13: 358. 1894 ≡ *Hyalopodium araraticum* (Lipsky) Röser & Tkach

*Description*: Perennial, caespitose, with creeping underground shoots; aerial shoots enclosed at the base by reticulately fibrous sheaths of dead leaves; culms erect, 20–55 cm long; ligule an eciliate membrane, 3–5 mm long, acute; leaf blades 4–11 cm long, 1–3 mm wide, midrib prominent beneath, surface glabrous, margins cartilaginous; inflorescence a panicle, contracted, linear, interrupted, 4–11 cm long, 0.5–1.5 cm wide; primary panicle branches short, 0.2–0.6 cm long; spikelets solitary, pedicelled, comprising 2(–3) fertile florets, without rhachilla extension, cuneate, laterally compressed, 6–7 mm long, disarticulating below each fertile floret; glumes persistent, similar, shorter than spikelet, similar to fertile lemma in texture, gaping; lower glume oblong, 4.5 mm long, 3/4 to as long as upper glume, membranous, much thinner above and on margins, purple, 1-keeled, 1-veined, lateral veins absent, apex acute; upper glume elliptic, 4.5–6 mm long, as long as adjacent fertile lemma, membranous, much thinner above, with hyaline margins, purple, 1-keeled, 3-veined, apex acute; lemma elliptic, 4–6 mm long, membranous, much thinner above, purple and yellow, tipped with yellow, keeled, 5-veined; lateral veins less than 2/3 length of lemma; lemma surface pubescent, hairy below; lemma apex erose, obtuse; callus very short, pilose; palea keels smooth, eciliate; anthers 3.3–4.5 mm long, yellow or purple; caryopsis about 3 mm long; hilum elliptic, 1/3–1/2 of the grain.

***Hyalopodium araraticum*** (Lipsky) Röser & Tkach, **comb. nov.** ≡ *Catabrosa araratica* Lipsky, Trudy Imp. S.-Peterburgsk. Bot. Sada 13: 358. 1894.

***Paracolpodium baltistanicum*** (Dickoré) Röser & Tkach, **comb. nov.** ≡ *Colpodium baltistanicum* Dickoré, Stapfia 39: 114. 1995.

***Parapholis cylindrica*** (Willd.) Röser & Tkach, **comb. nov.** ≡ *Hainardia cylindrica* (Willd.) Greuter, Boissiera 13: 177. 1967.

***Parapholis* ×*pauneroi*** (Castrov.) Röser & Tkach, **comb. nov.** ≡ ×*Hainardiopholis pauneroi* Castrov., Anales Jard. Bot. Madrid 36: 238. 1980 [1979 publ. 1980].

## CONCLUSIONS

Our survey of the molecular phylogenetic differentiation of supertribe Poodae, including most of its genera and based on nuclear and plastid DNA sequence markers investigated in an almost overlapping set of taxa, provides a robust and well-resolved topology for most regions of the phylogenetic trees. Some major polytomies remain and should be resolved in future studies. Notably, the nuclear and plastid DNA trees agree in wide portions and show congruent branching patterns, making it likely that they reflect the actual phylogenetic relation of the taxa in these tree portions. Severe conflict between the trees, however, occurs but is confined to several clearly defined and localized, though sometimes larger stretches of the trees and is interpreted to be indicative of past hybridization (Figs. 1, 2, 4). Taxonomic groups with hybrid origin are subtribes Scolochloinae, Sesleriinae, Torreyochloinae, Phalaridinae, Airinae, Holcinae and Phleinae. Major reticulation processes across subtribes include *Macrobriza* and *Arctopoa*. Well-identifiable infra-subtribe hybrid origins, which partly encompass lineages with several genera, were found, for example, within Aveninae, Coleanthinae, Loliinae, Puccinelliinae and Sesleriinae (Figs. 1, 2, 4, 5–7) but may be more frequent if denser sampling of taxa will be implemented and tree resolution will be improved by future studies. We found no evidence on a hybrid origin of *Avenula* and *Helictochloa*, whereas ‘*Calamagrostis*’ *flavens* is likely an intergeneric hybrid between *Agrostis* and *Calamagrostis* that warrants further study.

An analysis of morphological and other characteristics based on a final data matrix of 188 mainly morphological characters and utilizing a phylogenetic tree based on all plastid and nuclear DNA markers studied was performed to reconstruct the evolutionary ancestral states of our study group Poodae and its major lineages. Altogether 74 characters could be analysed in detail this way.

The phylogenetically ancestral character states (suppl. Appendix S3) of Poodae include perennial life form; absence of rhizomes; ligule an eciliate membrane; flat leaves; inflorescence an open panicle with many spikelets; panicle branches at most moderately divided, smooth; spikelets pedicelled, with >3 fertile florets, with a rhachilla extension bearing a sterile floret; spikelets without basal sterile florets, fertile florets (if more than 1) all alike, laterally and at most moderately compressed, comparatively large, >4 mm in length, breaking up at maturity, disarticulating below each floret; glumes shorter than spikelet, in consistency thinner than the lemma or similar; lower glume 3–6 mm long, 0.6-fold to as long as the upper, of similar consistency on margins, 1-keeled, keeled all along, 1-veined, primary vein eciliate, without lateral veins; upper glume 3–6 mm long, with undifferentiated margins, muticous, 1-keeled, 3-veined, primary vein distinct, smooth, lateral veins distinct; lemma 1.6– 4 mm, without keel, with 4–5 veins, surface generally rough, lemma apex erose or dentate, mucronate to awned, principal lemma awn straight, not coloured, without distinct column; column of lemma awn (if present) not twisted and not flattened, lateral lemma awns absent; palea ≥0.7-fold longer than the lemma, with smooth and eciliate keels; apical sterile florets (if present) resembling fertile though underdeveloped; stamens 3; caryopsis ≥1.6 mm long; hilum linear and straight.

Interestingly, the phylogenetically ancestral character states are sometimes different for Aveneae and Poeae, for example, the number of florets in the spikelets or spikelet size. A repeated switch of states during evolution including reversals is likely for many characters. The analysis revealed an overall high degree of homoplasy of spikelet characters. It includes, for example, the parallel, independent evolution of elaborate, geniculately bent awns multiple times in several evolutionarily separated lineages (Figs. 9‒11). This character, once assumed by taxonomists to be characteristic of Aveneae, originated at least six times and also could become secondarily lost again (see above *Ancestral state reconstruction*: characters 146, 146, 150; suppl. Fig. S2, suppl. Appendix S2). This parallels the findings in the PACMAD clade of grass subfamilies, in which similarly shaped twisted geniculate awns have originated at least five times independently (Teisher & al., 2017).

The overall high degree of homoplasy in many spikelet characters, we assume, relates to high degree of selective pressure acting on these structures. They have little to do with pollination as an important factor for the floral structures in many other angiosperms because all grass taxa in question are wind-pollinated. We suppose they have much more to do with efficient dispersal of diaspores, which is highly varied in grasses (Davidse, 1987). It can be supposed that the variety of dispersal mechanisms caused by spikelet structures (spikelet disarticulate or fall entire, different types of disarticulation, types of awns, animal dispersal, hygroscopic movement, bristles and hairs, lemma and palea structure, release of caryopses, etc.) are one of important evolutionary factors, which enabled Aveneae and Poeae to colonise successfully almost any habitat type in the temperate and cold zones of the world.

## AUTHOR CONTRIBUTIONS

MR, JS and NT designed the study. MR guided the sampling, contributed taxonomic knowledge, contributed data for the ancestral state reconstructions and wrote the manuscript. JS, NT, ED, AW, AH, GW, JG contributed lab work. NT and JS contributed data by supervising students in the lab. NT and JS undertook the phylogenetic analyses. JN and MR performed the ancestral state reconstructions. NT and MHH contributed to write the manuscript. — GW, https://orcid.org/0000-0002-9866-335X; MR, https://orcid.org/0000-0001-5111-0945; NT, https://orcid.org/0000-0002-4627-0706.

## ACKNOWLEDGEMENTS

We are grateful to Liliana Essi (Universidade Federal de Santa Maria, Santa Maria, RS), Božo Frajman (University of Innsbruck), Paola Gaiero (University of the Republic Uruguay, Montevideo), Caroline Mashau (South African National Biodiversity Institute, Pretoria), Mélica Muñoz-Schick (Museo Nacional de Historia Natural de Chile, Santiago de Chile), Carlos Romero Zarco (Departamento de Biología Vegetal y Ecología, Sevilla), Robert J. Soreng (Smithsonian Institution, Washington, D.C.), the late Nikolai N. Tzvelev (Komarov Botanical Institute, St. Petersburg) and the directors of the herbaria listed in Appendix 1 for supplying plant material for our study. Further we thank Bärbel Hildebrandt and Laura Freisleben (Halle) for technical support in our lab. Parts of this study were conducted by Antonia Lisker, Kerstin Schmidt, Peter Süssmann, Anne Weigelt within the framework of their university degree theses. Maria Vorontsova and Sarah Ficinski (both Royal Botanic Gardens, Kew) kindly provided the GrassBase data used for morphological analyses and ASR. A research grant of the German Research Foundation to MR (DFG RO 865/8) is gratefully acknowledged.

---

**Supplementary Figure S1.**

Maximum Likelihood phylogram of Poodae (Aveneae and Poeae) inferred from a concatenated matrix of plastid (*matK* gene–3’*trnK* exon, *trnL–trnF*) and nr (ITS, ETS) DNA sequences with species of Triticodae and Brachypodieae as outgroup. Maximum Likelihood and Maximum Parsimony bootstrap support values >50% as well as Bayesian posterior probabilities >0.5 are indicated on the branches. Clades with Maximum Likelihood support <50% are collapsed. The subtribes mentioned in the text are labelled on the right-hand side.

**Supplementary Figure S2.**

States of the scored 188 mainly morphological characters mapped on the tips of an ultrametric phylogenetic tree. The characters are listed in suppl. Appendix S2. The tree is based on the Maximum Likelihood analysis of the concatenated plastid and nuclear DNA sequence matrix as detailed in suppl. Fig. S1. Differently colored wedges indicate the presence of different character states.

## Appendix 1

Taxa studied in our lab for DNA sequences with geographical origin, voucher information with collectors and herbarium code and ENA/GenBank accession numbers for plastid *matK* gene–3’*trnK* exon; plastid *trnL–trnF*; nuclear ribosomal ITS1–5.8S gene–ITS2 and nuclear ribosomal ETS. Sequences LR606315–LR607006, LR655821 and LR655822 were newly generated for this study. Missing sequence data are indicated by a dash. BG: Botanical Garden. MR: herbarium of G. & S. Miehe deposited at the Institute of Geography, University Marburg, Germany.

***Acrospelion distichophyllum*** (P.Beauv.) Barberá: Austria, High Tauern, Glockner Alps, Pasterzen Kees, 19.07.2000, *G. Winterfeld 26* (HAL); LR606806, LT159704; LR606607; LT159798; LR606315. ***Agrostis alopecuroides*** Lam.: Cultivated in BG Halle, Germany from seed obtained from BG Dijon, France (no. 2001-1096), s.d., *M. Röser 11078* (HAL); LR606807, AM234719; LR606608; LR606513; LR606316. ***A. avenacea*** J.F.Gmel.: Australia, New South Wales, Great Dividing Range, 13.09.1998, *M. Röser 10762* (HAL); LR606808; LR606609; LR606514; LR606317. ***A. capillaris*** L.: Germany, Saxony, Upper Lusatia, 30.07.1998, *M. Röser 10660/2* (HAL); LR606809, AM234560; LR606610; FM179384; LR606318. ***A. linkii*** Banfi, Galasso & Bartolucci: Cultivated in BG Halle, Germany from seeds obtained from BG Copenhagen, Denmark, s.d., *s.coll*.. (HAL0140383); LR606810; LR606611; LR606515; LR606319. ***A. pallens*** Trin.: USA, Oregon, Clackamas County, Mt. Hood, 26.08.2000, *R.J. Soreng 6361* (US); –; LR606612; LR606516; LR606320. ***A. ramboi*** Parodi: Brazil, Sta Catarina, Campo dos Padres, 22.01.1957, *B. Rambo 60074* (B 10 0448888); LR606811; –; –; LR606321. ***A. scabra*** Willd.: USA, Alaska, Kenai Peninsula, Resurrection River, 08.07.2000, *R.J. Soreng 6078* (US Catalog No.: 3682815, Barcode: 01259848); –; LR606613; LR606517; –. ***A. tandilensis*** (Kuntze) Parodi: Brazil, Rio Grande do Sul, Garibaldi, 13.10.1957, *O. Camargo 62575* (B 10 0448889); –; –; LR606518; LR606322. ***Aira elegans*** Willd. ex Roem. & Schult. (1): Cultivated in BG Halle, Germany from seed obtained from BG Munich-Nymphenburg, Germany, 02.07.2002, *s.coll*. (HAL0140286); –; –; LR606519; –; (2): Austria, Tyrol, Paznaun, Verwall Alps; cultivated in BG Halle, Germany from seed obtained from BG Berlin-Dahlem, Germany (no. 2001-3947), 14.10.2002, *Royl & Hempel* (HAL); LR606812; LR606614; –; LR606323. ***A. praecox*** L.: Germany, Mecklenburg-Vorpommern, Müritz Lake, 28.05.2003, *M. Röser 11009/1* (HAL); LR606813, AM234540; LR606615; FM179385; LR606324. ***Airopsis tenella*** (Cav.) Coss. & Durieu: France, Montpellier, Bellargues, 01.05.1956, *R. Schubert* (HAL0080969); LR606814; LR606616; LR606520; LR606325. ***Alopecurus aequalis*** Sobol.: Germany, Baden-Württemberg, near Tübingen, 28.06.1984, *M. Röser 1892* (HAL); LR606815; LR606617; LR606521; LR606326. ***Ammochloa palaestina*** Boiss.: Spain, Andalucía, Province Almería, Tabernas, 13.04.1965, *F. Bellot & S. Rivas Goday* (C); LR606816; LR606618; LR606522; LR606327. ***A. pungens*** Boiss.: Algeria, between Djelfa and Bou-Saâda, 06.04.1965, *V.P. Bochantsev 1238* (LE); LR606817; LR606619; LR606523; LR606328. ***Amphibromus nervosus*** (Hook.f.) Baill.: Australia, New South Wales, Great Dividing Range, 23.05.2002, *M. Röser 10770* (HAL); LR606818; LR606620; LR606524; LR606329. ***Ancistragrostis uncinioides*** S.T.Blake: New Guinea, Central District, Papua, Mount Victoria, 10.07.1974, *L.A. Craven 3006* (L0533422); –; –; LR606525; –. ***Aniselytron treutleri*** (Kuntze) Soják: China, Yunnan, Fugong Province, Bilou Mts., 08.09.1997, *R.J. Soreng 5229, P.M. Peterson & Sun Hang* (US); LR606819; LR606621; –; –. ***Anthoxanthum arcticum*** Veldkamp: Russia, Yakutia, lower reaches of the Kolyma River, 27.07.1975, *V.V. Petrovskiy & I.A. Mikhaylova* (LE); LR606820; –; LR606526; –. ***A. australe*** (Schrad.) Veldkamp (1): Austria, Burgenland, Bernstein, upper area of Steinstückel range, 13.05.1992, *M. Röser 9089* (HAL); LR606821; –; LR606527; –; (2): France, Hautes-Alpes, 05.08.1984, *M. Röser 2206* (HAL); –; LR606622; –; LR606330. ***A. glabrum*** (Trin.) Veldkamp (1): Russia, Khakassia, Ust-Abakan District, 08.06.1968, *I. Neyfeld* (LE); LR606822; LR606624; LR606529; LR606331; (2): Russia, Kemerovo Oblast, 23.05.2003, *s.coll.* (LE); –; LR606623; LR606528; –; (3) subsp. ***sibiricum*** (Tzvelev) Röser & Tkach: Russia, Tomsk Oblast, 03.06.1912, *L. Utkin* (LE); LR606823; LR606625; LR606530; –. ***A. monticola*** (Bigelow) Veldkamp: Russia, Sibiryakova Island, 19.06.2016, *M.B. Matveeva & I.I. Zanokha 2730* (LE); LR606824; LR606626; LR606531; LR606332. ***A. nitens*** (Weber) Y.Schouten & Veldkamp subsp. ***kolymensis*** (Prob.) Röser & Tkach: Russia, Yakutia, Nizhnekolymskiye Kresty, 30.06.1950, *G. Nemlin 340* (LE); LR606825; LR606627; LR606532; LR606333. ***A. odoratum*** L.: Russia, Irkutsk Oblast, Trehgolovyy Golez Mount, 04.07.1986, *K. Baykov 298* (NS/NSK); LR606826; LR606628; LR606533; LR606334. ***A. redolens*** (Vahl) P.Royen: Chile, Chiloe Island; cultivated in BG Halle, Germany from seed obtained from BG Olomouc, Czech Republic, s.d., *s.coll.* (HAL); LR606827; LR606629; LR606534; LR606335. ***A. repens*** (Host) Veldkamp: Russia, Tomsk Oblast, Barnaul District, 05.06.1890, *S. Korshinskiy* (LE); LR606828; LR606630; LR606535; LR606336. ***Antinoria agrostidea*** (DC.) Parl.: Portugal, Province Beira Alta, Serra da Estrela, Lagoa do Paixão, 15.08.1986, *Arriegas, Loureiro, Santos & Seleiro 192* (COI); LR606829; –; LR606536; LR606337. ***A. insularis*** Parl.: Greece, Crète, Nomos Hania, plateau d’Omalos, 23.05.1998, *A. Charpin 25346* (G86517); –; –; LR606537; LR606338. ***Apera spica-venti*** (L.) P.Beauv.: Germany, Mecklenburg-Vorpommern, Müritz Lake, 26.07.2001, *M. Röser 11005* (HAL); LR606830, AM234542; LR606631; LR606538; LR606339. ***Arctagrostis latifolia*** (R.Br.) Griseb.: Norway, Finnmark, Nesseby, 23.08.1997, *T. Alm & A. Often 563* (TROM 64713); LR606831; LR606632; HE802200; LR606340. ***Arctohyalopoa lanatiflora*** (Roshev.) Röser & Tkach (1): Russia, Yakutia, basin of Tompon River, 01.07.1956, *I.D. Kildyushevskiy 18/1* (LE); LR606833, AM234604; LR606633; LR606540; LR606341; (2): Russia, Yakutia, Verkhoyanskiy Range, 17.07.1985, *E. Rybinskaya 395* (NS/NSK); LR606832; –; LR606539; –. ***Arctopoa eminens*** (J.Presl) Prob.: Russia, Far East, Kuril Islands, Iturup, 25.07.1959, *E. Pobedimova & G. Konovalova 986* (LE); LR606834; LR606634; HE802201; LR606342. ***Arrhenatherum elatius*** (L.) P.Beauv. ex J.Presl & C.Presl: Germany, Saxony, Leipzig, s.d., *G. Winterfeld 77* (HAL); LR606835, AM234543, HG797415; LR606635; FM179388; LR606343. ***Avellinia michelii*** (Savi) Parl.: Spain, Valencia, Devesa de l’Albufera, s.d., *J.B. Peris & G. Stubing 1977* (RO); LR606836; LR606636; LT159736; LR606344. ***Avena hispanica*** Ard.: Cultivated in BG Halle, Germany from seed obtained from Agriculture Canada, Ottawa, Canada (no. CAV 6604); s.d., *s.coll.* (HAL); LR606837; LR606637; LR606541; LR606345. ***A. macrostachya*** Balansa ex Coss. & Durieu: Algeria; cultivated in BG Halle, Germany from seed obtained from M. Leggett, Institute of Grassland and Environmental Research, Aberystwyth, UK (no. CC7068); s.d., *s.coll.* (HAL); FM253118, FM957002, HG797416; LR606638; FM179443; LR606346. ***Avenella flexuosa*** (L.) Drejer: Germany, Mecklenburg-Vorpommern, Müritz Lake, 27.07.2001, *M. Röser 11008* (HAL); LR606838, AM234545; LR606639; FM179389; LR606347. ***Avenula pubescens*** (Huds.) Dumort.: Hungary, Vezsprem, between Csabrendek and Sümeg, 23.05.1999, *M. Röser 10928/2* (HAL); FM253118, FM957003, HG797417; LR606640; FM956100, HG797487; LR606348. ***Beckmannia eruciformis*** (L.) Host: Russia, Yakutia, Ordzhonikidzevskiy District, 22.08.1982, *Bolshakov & Vlasova 4377* (NS/NSK); LN554423; LR606641; HE802171; LR606349. ***Bellardiochloa polychroa*** (Trautv.) Roshev.: Armenia, Agaraz Mount, 10.08.1969, *V.E. Voskonjan* (LE); LR606839, FM253119; LR606642; FM179390; LR606350. ***B. variegata*** (Lam.) Kerguélen subsp. ***aetnensis*** (C.Presl) Giardina & Raimondo: Italy, Sicily, Catania Province, Mount Etna, 29.10.1987, *M. Röser 6032* (HAL); LR606840, AM234605; LR606643; FM179391; LR606351. ***Boissiera squarrosa*** (Sol.) Nevski (1): Iran, Gilan, Zandjan, 09.05.1969, *H. Eckerlein* (HAL0022065); LR606841, FM253120, LN554424; –; FM179392; –; (2): Israel; cultivated in BG Halle, Germany from seed obtained from Kew’s Millennium Seed Bank, UK (no. 537580), s.d., *s.coll.* (HAL); –; LR606644; LR606542; LR606352. ***Brachypodium distachyon*** (L.) P.Beauv.: Spain, Andalucía, Province Almería, Cabo de Gata, 11.04.1986, *M. Röser 4359* (HAL); LR606842, AM234568, LN554426; LR606645; –; LR606353. ***Briza media*** L.: Germany, Thuringia, NE Jena, Tautenburger Forest, 21.05.2005, *M. Röser 11072* (HAL); AM234610, HG797418, LN554427; –; FM179393; LR606354. ***B. minor*** L.: Italy, Abruzzo; cultivated in BG Halle, Germany from seed obtained from Kew’s Millennium Seed Bank, UK (no. 6150), 19.08.1977, *P. Newman, P.A. Thompson, E.A.M. Ormerod & R.H. Sanderson* (HAL); LR606843; LR606646; KJ598892; LR606355. ***Brizochloa humilis*** (M.Bieb.) Chrtek & Hadač: Russia, Krym, Peninsula Tarkhankut, 25.05.1984, *N.N. Tzvelev, D.V. Geltman, N.A. Medvedeva & G.V. Mustafina 1110* (LE); LR606844; –; HE802178; LR606356. ***Bromus erectus*** Huds.: France, Hérault, Causses du Larzac, 07.06.1984, *M. Röser 1721* (HAL); AM234570, FM956476; LR606647; FM179394, FM956470; –. ***Calamagrostis arenaria*** (L). Roth subsp. ***arundinacea*** (Husn.) Banfi, Galasso & Bartolucci: Portugal, Odemira, Vil Nova de Milfontes; cultivated in BG Halle, Germany from seed obtained from BG Lisbon, Portugal, s.d., *M. Röser 11055* (HAL); LR606845, AM234561; LR606648; LR606543; LR606357. ***C. arundinacea*** (L.) Roth: Germany, Lower Saxony, Harz Mts., Siebertal above Herzberg, 02.08.1983, *M. Röser 1232* (HAL); LR606846; LR606649; LR606544; LR606358. ***C. canescens*** (Weber) Roth: Germany, Saxony, Freiberger Mulde, 01.08.2005, *S. Schiebold & A. Golde* (HAL0004118); LR606847; LR606650; LR606545; LR606359. ***C. macrolepis*** Litv.: Mongolia, s.d., *K. Wesche 4279* (HAL); LR606848, AM234559; LR606651; LR606546; LR606360. ***C. neglecta*** (Ehrh.) G.Gaertn., B.Mey. & Scherb. subsp. ***borealis*** (C.Laest.) Selander: USA, Alaska, Barrow, Gas Well Road, 01.08.2000, *R.J. Soreng 6204* (US); –; LR606652; LR606547; LR606361. ***C. nutkaensis*** (J.Presl) Steud.: USA, Alaska, Kenai Peninsula, Seward, 08.07.2000, *R.J. Soreng 6062* (US); LR606849; LR606653; LR606548; LR606362. ***C. purpurascens*** R.Br. (1): Canada, Yukon, Marsh Lake, 14.06.2000, *R.J. Soreng 5996b* (US); LR606850, LT222486; LR606655; FM179395; LR606363; (2): Canada, Yukon, Kluane Lake, Duke River Bridge, 14.08.2000, *R.J. Soreng 6301* (US); –; LR606654; LR606549; –. ***‘C.’ rigida*** (Kunth) Trin. ex Steud.: Bolivia, Department La Paz, Province Murillo, 12.02.1989, *S.G. Beck 14738* (B 10 0448895); LR606851, HG797422; LR606656; HG797492; LR606364. ***C. rivalis*** H.Scholz: Germany, Saxony, Mulde River, 30.09.2002, *M. Röser 11054/D* (HAL); LR606852, AM234564; LR606657; LR606550; LR606365. ***Catabrosa aquatica*** (L.) P.Beauv.: Germany, Baden-Württemberg, Zollhausried near Blumberg, 18.07.1984, *M. Röser 2007* (HAL); LR606853, AM234589; –; FM179396; LR606366. ***Catabrosella humilis*** (M.Bieb.) Tzvelev: Kazakhstan, Ili River, 08.05.1934, *N.I. Rubtsov* (LE); LR606854; –; HE802182; LR606367. ***C. variegata*** (Boiss.) Tzvelev: Russia, Kabardino-Balkar Republic, Caucasus, Mount Elbrus’ foot, 24.07.1939, *E.V. Schiffers & T.A*. *Moreva* (LE); LR606855; –; HE802181; –. ***Catapodium marinum*** (L.) C.E.Hubb.: Spain, Valencia, Province Alicante, Cabo de Santa Pola, 10.04.1986, *M. Röser 4299* (HAL); LR606856, HE646574; LR606658; HE646600; LR606368. ***C. rigidum*** (L.) C.E.Hubb.: Greece, Macedonia, Thessaloniki, Chalcidice, 25.05.1985, *M. Röser 2571* (HAL); LR606857, AM234586; LR606659; FM179399; –. ***Chascolytrum bulbosum*** (Parodi) Essi, Longhi-Wagner & Souza-Chies: Brazil, Rio Grande do Sul, Pirationi, 16.11.2003, *L. Essi 50, J.F.M. Valls, A. Guglieri & S. Hefler* (ICN); LR606858; LR606660; LR606551; LR606369. ***C. rhomboideum*** (Link) Essi, Longhi-Wagner & Souza-Chies: Chile, Linares Province, Department Loncomilla, 12.10.1954, *R. Avendaño T.* (SGO071551); LR606859; LR606661; LR606552; LR606370. ***C. subaristatum*** (Lam.) Desv.: Argentina, Buenos Aires Province; cultivated in BG Halle, Germany from seed obtained from BG Berlin-Dahlem, Germany (no. 2001-3817), s.d., *M. Röser 11079* (HAL); LR606860, AM234608; LR606662; LR606553; LR606371. ***C. uniolae*** (Nees) Essi, Longhi-Wagner & Souza-Chies: Paraguay, Department Paraguarí, National Park Ybycui, 31.10.1989, *Zardini & Guard 14580* (MO3879842); LR606861; LR606663; LR606554; LR606372. ***Cinna latifolia*** (Trevir. ex Göpp.) Griseb.: Finland, South Savo, Rantasalmi, 13.08.1977, *M. Isoviita* (HAL0050605); LR606862; LR606664; HE802198; LR606373. ***Coleanthus subtilis*** (Tratt.) Seidel ex Roem. & Schult.: Austria, Lower Austria, 30.08.2006, *H. Rainer & M. Röser 11082* (HAL); LR606863; LR606665; HE802180; LR606374. ***Colpodium biebersteinianum*** (Claus) Röser & Tkach: Cultivated in BG Halle, Germany from seed obtained from Institute of Plant Genetics and Crop Plant Research, Gatersleben, Germany, 28.05.2003, *s.coll.* (HAL); AM234551, LN554457; LR606666; HE802184; LR606375. ***C. chionogeiton*** (Pilg.) Tzvelev: Tanzania, Kilimanjaro, above Mawengi hut, 25.11.1967, *D.G. King 6* (UPS:BOT:V-652825); LR606864; –; HE802185; LR606376. ***C. hedbergii*** (Melderis) Tzvelev: Ethiopia, Bale Province, Bale Mountains National Park, Saneti Plateau; cultivated in BG Uppsala, Sweden, 07.06.1905, *O. Hedberg 5618* (UPS:BOT:V-652843); LR606865; –; HE802186; LR606377. ***C. trichopodum*** (Boiss.) Röser & Tkach: Cultivated in BG Halle, Germany from seed obtained from Institute of Plant Genetics and Crop Plant Research, Gatersleben, Germany, 28.05.2003, *M. Röser 11074* (HAL); LR606866, AM234551; LR606667; FM179441; LR606378. ***C. versicolor*** (Steven) Schmalh.: Georgia, South Ossetia, Ermany., 23.08.1938, *E.A. & N.A. Bush* (LE); LR606867, FM253122; –; FM179397; LR606379. ***Cornucopiae cucullatum*** L.: Cultivated in BG Halle, Germany from seeds obtained from Botanical Garden Frankfurt, Germany (no. 2005-926), s.d., *E. Döring* (HAL0100582); LR606868; LR606668; HF564627; LR606380. ***Corynephorus canescens*** (L.) P.Beauv. (1): Portugal, Province Estremadura, Vieria, Pinhal de Leira, 14.07.1992, *M. Röser 9483* (HAL); LR606869; –; HE802179; –; (2): Germany, Saxony-Anhalt, Harz Mts., 14.05.2016, *M. Röser 11230 & N. Tkach* (HAL); –; LR606669; LR606555; LR606381. ***Cutandia maritima*** (L.) Barbey: France, Hérault, Etang d’Ingril, 30.05.1977, *A. Dubuis* (HAL0048831); LR606870, HE646572; LR606670; HE646601; LR606382. ***Cyathopus sikkimensis*** Stapf: China, Yunnan, Fugong Province, s.d., *R.J. Soreng 3224, P.M. Peterson, Sun Hang* (US); LR606871, AM234553; LR606671; HE802199; LR606383. ***Cynosurus cristatus*** L.: Germany, Baden-Württemberg, Ettenheim, 16.06.1989, *M. Röser 9965* (HAL); LR606872, HE646575; –; HE646602; LR606384. ***C. elegans*** Desf.: France, Corsica, Forêt d’Aitone, 28.06.1987, *M. Röser 5420* (HAL); LR606873, HG797427; LR606672; LR606556; LR606385. ***Dactylis glomerata*** L.: Greece, Macedonia, Serron, Vrondus range, 31.05.1985, *M. Röser 2948* (HAL); LR606874, AM234595; LR606673; LR606557; –. ***Deschampsia bolanderi*** (Thurb.) Saarela: USA, California, Monterey County, Hanging Valley, Santa Lucia Mts., 11.06.2003, *D.H. Wilken 16163 & E. Painter* (RSA 695253); LR606875, HE646588; –; HE646612; LR606386. ***D. cespitosa*** (L.) P.Beauv.: Germany, Brandenburg, Niederspree, s.d., *M. Röser 10737/1* (HAL); LR606876, AM234546; LR606674; AF532929; LR606387. ***D. danthonioides*** (Trin.) Munro: USA, California, Siskiyou, Klamath River, 03.06.2000, *R.J. Soreng 5965* (US); LR606877; LR606675; LR606558; LR606388. ***D. micrathera*** (É.Desv.) Röser & Tkach: Argentina, Neuquén Province, Los Lagos, Correntoso, 27.01.1990, *Zulma Rúgolo 1245* (B 10 0448863); LR606878, LT159689; –; LT159754; LR606389. ***Desmazeria philistaea*** (Boiss.) H.Scholz: Israel, Philistean Plain, 21.03.1989, *A. Danin et al. 03.074* (B 10 0240417); LR606879, HE646573; LR606676; HE646603; LR606390. ***D. sicula*** (Jacq.) Dumort. (1): Malta, Dwejra Point, 01.04.1975, *A. Hansen 490* (C); LR606880, HE646576; LR606677; HE646604; –; (2): Malta, Gozo; cultivated in BG Halle, Germany from seed obtained from Kew’s Millennium Seed Bank, UK (no. 17332), 02.08.1981, *J. Newmarch* (HAL); –; LR606678; LR606559; LR606391. ***‘Deyeuxia’ contracta*** (F.Muell. ex Hook.f.) Vickery: Australia, Tasmania; cultivated in BG Halle, Germany from seed obtained from Kew’s Millennium Seed Bank, UK (no. 391131), 21.02.2007, *E. Brüllhardt & M. Visoiu* (HAL); LR606881; LR606679; LR606560; LR606392. ***‘D.’ flavens*** Keng: China, Qinghai, surroundings of Maqen, 06.08.2004, *I. Hensen* (HAL); LR606882; LR606680; LR606561; LR606393. ***Dichelachne crinita*** (L.f.) Hook.f.: New Zealand, Canterbury, Banks Peninsula, Pigeon Bay, 03.12.1990, *J.R. Bulman* (CHR 477794); LR606883; LR606681; LR606562; LR606394. ***D. micrantha*** (Cav.) Domin: Australia, New South Wales, Thirlmer Lakes area, 04.10.1998, *R.J. Soreng 5901, P.M. Peterson, S.W.L. Jacobs* (US); LR606884, FM253124; LR606682; FM179401; LR606395. ***Drymochloa sylvatica*** (Pollich) Holub: Germany, Lower Saxony, Harz Mts., Siebertal above Herzberg, 02.08.1983, *M. Röser 1227* (HAL); LR606885, AM234585; LR606683; FM179404; LR606396. ***Dryopoa dives*** (F.Muell.) Vickery (1): Australia, Tasmania, Hobart District, 04.12.1980, *T. Walker* (AD98132291); LR606886; LR606684; LR606563; LR606397; (2): Australia, Victoria; cultivated in BG Halle, Germany from seed obtained from Kew’s Millennium Seed Bank, UK (no. 333531), 15.02.2006, *M.J. Hirst & S. Hodge* (HAL); –; LR606685; LR606564; –. ***Dupontia fisheri*** R.Br. subsp. ***psilosantha*** (Rupr.) Hultén: Russia, Yakutia, estuary of Yana River, near Nizhneyansk, 27.07.1988, *Doronkin & Bubnova 439* (NSK); LR606887, AM234601; LR606686; AY237848; LR606398. ***Dupontia fulva*** (Trin.) Röser & Tkach: Russia, Yakutia, estuary of Yana River, near Nizhneyansk, 21.07.1988, *Doronkin & Kulagin 81* (NSK); LR606888, AM234606; LR606687; FM179387; LR606399. ***Echinaria capitata*** (L.) Desf.: Spain, Andalusia, Province Granada, Sierra Nevada, 14.06.1985, *M. Röser 3336* (HAL); LR606889, AM234599, LN554434; LR606688; LR606565; LR606400. ***Echinopogon caespitosus*** C.E. Hubb.: Australia, New South Wales, Thirlmer Lakes area, 04.10.1998, *R.J. Soreng 5900, P.M. Peterson, S.W.L. Jacobs* (US); LR606890, AM234609; LR606689; FM179403; LR606401. ***Festuca berteroniana*** Steud.: Chile, Juan Fernandez, Masatierra, Corrales de Molina, 24.01.1990, *D. Wiens, P. Penailillo, R. Schiller, A. Andaur* (MO5259377); LR606891, HE646581; LR606690; FR692028; LR606402. ***F. floribunda*** (Pilg.) P.M.Peterson, Soreng & Romasch.: Peru, Department Moquegua, Province Mariscal Nieto, 01.03.1999, *P.M. Peterson 14566, N. Refulio Rodriguez & F. Salvador Perez* (NY); LR606892; LR606691; LR606566; LR606403. ***F. incurva*** (Gouan) Gutermann: Spain, Provincia of Salamanca, 31.05.1987, *F. Amich & J.A. Sánchez 19923* (RO); LR606893, HE646587; LR606692; HE646611; LR606404. ***F. lachenalii*** (C.C.Gmel.) Spenn.: France, Corsica, 29.06.1987, *M. Röser 5470* (HAL); LR606894; –; LR606567; –. ***F. maritima*** L.: France, Montpellier, Bois de Boscares, 01.04.1956, *R. Schubert* (HAL0081028); LR606895, HE646590; LR606693; AY118095; –. ***F. masatierrae*** Röser & Tkach: Chile, Valparaíso Region, Juan Fernándes, s.d., *R.A. Philippi* (HAL0052812); –; –; FR692035; –. ***F. myuros*** L.: Cultivated in BG Halle, Germany from seed obtained from BG Dijon, France (no. 1060), 02.07.2002, *s.coll.* (HAL); LR606896; LR606694; –; –. ***F. salzmannii*** (Boiss.) Boiss. ex Coss.: Spain, Andalucía, Province Malaga, Sierra de Mijas, Alhaurín el Grande, 08.05.1989, *S. Rivas-Martínez 17742* (BASBG); LR606897, HE646583; LR606695; HE646608; LR606405. ***Gastridium nitens*** (Guss.) Coss. & Durieu: Greece, Crete, Agios Nikolaos, 01.05.1983, *G. van Buggenhout* (ROM); LR606898; LR606696; LR606568; LR606406. ***G. phleoides*** (Nees & Meyen) C.E.Hubb.: Lebanon, North Lebanon; cultivated in BG Halle, Germany from seed obtained from Kew’s Millennium Seed Bank, UK (no. 241421), 28.07.2004, *M. van Slageren & Khairallah, S.* (HAL); LR606899; LR606697; LR606569; LR606407. ***G. ventricosum*** (Gouan) Schinz & Thell.: France, Corsica, 30.06.1987, *M. Röser 5491* (HAL); LR606900; LR606698; LR606570; LR606408. ***Gaudinia fragilis*** (L.) P.Beauv.: Spain, Andalucíia, Province Cádiz, NW Gibraltar, 20.06.1985, *M. Röser 11070* (HAL); LN554436; LR606699; LT159737; –. ***Graphephorum melicoides*** (Michx.) Desv.: Canada, New Brunswick, Madawaska County, 04.08.1990, *G. Flanders & R. Hinds 981* (CAN550357); LR606901, HG797428; LR606700; HG797505; LR606409. ***G. wolfii*** J.M.Coult.: USA, Colorado, San Juan County, 06.08.1982, *R.J. Soreng* (NY); LR606902, HG797429; LR606701; HG797506; LR606410. ***Helictochloa aetolica*** (Rech.f.) Romero Zarco: Greece, Epirus, Tomaros Mts., 23.02.2004, *M. Röser 10726/3* (HAL); FM957008; –; LR606571; LR606411. ***H. bromoides*** (Gouan) Romero Zarco subsp. ***bromoides***: France, Vaucluse, 19.08.1997, *M. Röser 10630/2* (HAL); LR606903, AM234721, FM956474, HG797430; LR606702; FM956463; LR606412. ***H. compressa*** (Heuff.) Romero Zarco: Greece, Macedonia, Drama, Orvilos region, 16.08.1998, *M. Röser 10707/8* (HAL); FM957009; LR606703; –; LR606413. ***H. hookeri*** (Scribn.) Romero Zarco (1) subsp. ***hookeri:*** Canada, Yukon, Kluane Lake, Duke River Bridge, 14.08.2000, *R.J. Soreng 6305* (US); LR606904; LR606704; LR606572; LR606414; (2) subsp. ***schelliana*** (Hack.) Romero Zarco: Mongolia, Chentej Aimag, 02.08.2002, *K. Wesche 4333* (HAL); LR606905, AM234550; LR606705; FM179409, FN984915; LR606415. ***H. levis*** (Hack.) Romero Zarco: Spain, Andalusia, Province Granada, Sierra Nevada, 23.04.2001, *G. Winterfeld 50* (HAL); FM958418; LR606706; LR606573; LR606416. ***H. marginata*** (Lowe) Romero Zarco: Portugal, Province Beira Alta, Serra da Estrela, between São Romão and Torre, 12.07.1992, *M. Röser 9421* (HAL); FM957007; –; LR606574; LR606417. ***H. versicolor*** (Vill.) Romero Zarco: France, Haute Garonne, Pyrenees, Pic de Cécire, 21.08.1985, *M. Röser 3937* (HAL); LR606906, FM957011; –; FM956467; LR606418. ***Helictotrichon convolutum*** (C.Presl) Henrard: Greece, Peloponnese, Arkadia, Menalon, 10.08.1998, *M. Röser 10697* (HAL); LR606907, AM234557, HG797431; LR606707; FM179406, FM956461; LR606419. ***H. mongolicum*** (Roshev.) Henrard: Russia, E Sayan Mts., Large Kishta River source, 14.08.1962, *L. Malyshev 795* (NS/NSK); LR606908, HG797439; LR606708; HG797516; LR606421. ***H. parlatorei*** (Woods) Pilg.: Austria, Carinthia, 16.10.2001, *B. Heuchert 11-08* (HAL); AM234566, FM957005, HG797442; LR606709; FM179408, LT159741; LR606422. ***H. sarracenorum*** (Gand.) Holub: Spain, Andalucía, Province Granada, between Guadix and Granada, 13.06.1985, *M. Röser 3266* (HAL); LR606909, FM956473, HG797443; LR606710; FM956462, HG797519; –. ***H. sedenense*** (DC.) Holub: France, Pyrénées-Orientales, Mount Canigou, 09.08.1997, *M. Röser 10545* (HAL); LR606910, FM957004, HG797444; LR606711; FM956104, HG797520; LR606423. ***H. sempervirens*** (Vill.) Pilg.: France, Drôme, 22.08.1984, *M. Röser 2429* (HAL); HG797445; LR606712; HG797521; –. ***H. setaceum*** (Vill.) Henrard subsp. ***petzense*** (H.Melzer) Röser: Austria, Carinthia, Karavankes near Bleiburg, Petzen, 09.07.1998, *M. Röser 10646* (HAL); LR606911, FM957010; LR606713; FM956468; –. ***H. thorei*** (Duby) Röser: Portugal, Province Minho, 02.07.2002, *M. Röser 9322/3A* (HAL); LR606912, AM234565, HG797448; LR606714; FM956102, FM179430; LR606424. ***H.* ×*krischae*** Melzer: Austria, Carinthia, Karavankes near Bleiburg, Petzen, 09.07.1998, *M. Röser 10648* (HAL); LR606913, FM958417, HG797451; LR606715; FM958415, HG797513; LR606420. ***Holcus mollis*** L.: Germany, Saxony, Upper Lusatia, 04.07.2002, *M. Röser 10658/2* (HAL); LR606914, AM234554; LR606716; FM179411; LR606425. ***Hookerochloa eriopoda*** (Vickery) S.W.L.Jacobs: Australia, Southern Tablelands, 30.01.1964, *R. Pullen 4003* (AD96435171); LR606915, HE646578; –; HE646605; LR606426. ***H. hookeriana*** (F.Muell. ex Hook.f.) E.B.Alexeev: Australia, Tasmania, Macquarie Rivulet, 01.02.2011, *A.M. Buchanan 15711* (HO 507299); LR606916, HE646579; LR606717; HE646606; LR606427. ***Hordelymus europaeus*** (L.) O.E.Harz: Germany, Baden-Württemberg, Suebian Alb, Urach, 01.08.1982, *M. Röser 708* (HAL); AM234596, LN554438; –; FM179412; LR606428. ***Hordeum marinum*** Huds. subsp. ***gussoneanum*** (Parl.) Thell.: Italy, Sardinia, Nuoro Province, Altipiano de Campeda, 09.05.1993, *M. Röser 10131* (HAL); FR694880, HG797452; LR606718; FR692026; –. ***Hyalopoa pontica*** (Balansa) Tzvelev: Russia, Balkaria, moraine of Karachiran glacier, 29.07.1925, *E. Bush & N. Bush* (LE); –; –; LR606575; –. ***Hyalopodium araraticum*** (Lipsky) Röser & Tkach: Armenia, Geghama Mts., Spitak-Syr, 21.08.1960, *Arverdyaev & Mirzaeva* (HAL0008785); LR606917; LR606719; HE802183; LR606429. ***Hypseochloa cameroonensis*** C.E.Hubb.: Cameroon, Cameroons Mountain, 01.12.1929, *T.S. Maitland 874* (B 10 0448883); LR606918; LR606720; LR606576; LR606430. ***Koeleria capensis*** Nees: Uganda, Mount Elgon, Sasa Trail, s.d., *K. Wesche 20026* (HAL); LR606919, AM234558, HG797453; LR606721; FM179413; –. ***K. loweana*** A.Quintanar, Catalán & Castrov.: Portugal, Madeira, 05.09.1983, *L. Dalgaard & V. Dalgaard 13276* (C); LR606920, HE646580; LR606722; HE646607; –. ***K. pyramidata*** (Lam.) P.Beauv.: Mongolia, Central Aimag, N to Ulan-Bator, 15.05.1944, *Ju.A. Yunatov 4381* (LE); LR606921, LT159683; LR606723; LT159743; LR606431. ***Lagurus ovatus*** L.: Portugal, Minho Province, coastal area at Eposende; cultivated in BG Halle, Germany from seed, 19.08.2002, *M. Röser 9271* (HAL); LR606922, AM234563, HG797455; LR606724; FM179414; –. ***Lamarckia aurea*** (L.) Moench: Spain, Murcia, between Murcia and Lorca, 11.04.1986, *M. Röser 4383* (HAL); LR606923; LR606725; LR606577; –. ***Limnas malyschevii*** O.D.Nikif.: Russia, Putorana plateau, Haya-Kuyol Lake, 10.08.1972, *S. Andrulajtis 1204* (NS/NSK); LR606924; LR606726; HE802176; LR606432. ***L. stelleri*** Trin.: Russia, Yakutia, Mirninskiy District, Mogdy River, 15.08.1975, *N. Vodopyanova, E. Ammosov, V. Strelkov 813* (NS/NSK); LR606925; LR606727; HE802175; –. ***Limnodea arkansana*** (Nutt.) L.H.Dewey: USA, Texas, Washington County, 01.05.1976, *T. F. Daniel 69* (NY); LR606926, LN554440; LR606728; LR606578; –. ***Littledalea tibetica*** Hemsl.: China, Qinghai, Kunlun Shan, 27.07.1994, *R.J. Soreng, P.M. Peterson, Sun Hang 5487-90-94* (US); LR606927, AM234572, LN554441; LR606729; FM179416; LR606433. ***Lolium giganteum*** (L.) Darbysh.: Germany, Lower Saxony, Harz Mts., Wolfshagen, 23.07.1987, *M. Röser 5719* (HAL); LR606928, AM234720; LR606730; HE646615; LR606434. ***Macrobriza maxima*** (L.) Tzvelev: France, Languedoc-Roussillon, Gard; cultivated in BG Halle, Germany from seed obtained from Kew’s Millennium Seed Bank, UK (no. 69618), 22.08.1988, *J. Feltwell* (HAL), LR606929; LR606731; LR606579; LR606435. ***Mibora minima*** (L.) Desf.: Cultivated in BG Halle, Germany from seeds, origin unknown, *s.coll.* (HAL0107426); LR606930, FR694894; LR606732; FR692030; LR606436. ***Milium effusum*** L.: France, Alpes-Maritimes, 21.07.1989, *M. Röser 6723* (HAL); LR606931, AM234598, HG797456; –; FM179419; LR606437. ***M. transcaucasicum*** Tzvelev: Armenia, Gukasyan District, Caucasus, Javakheti range’s foot, 27.06.1960, *N.N. Tzvelev & S. Czerepanov 425* (LE); LR606932; –; HE802197; –. ***Molineriella laevis*** (Brot.) Rouy: Spain, Province of Madrid, Manzanares el Real, 15.05.1984, *P. Montserrat* (C); LR606933; LR606733; LR606580; LR606438. ***M. minuta*** (L.) Rouy: Greece, Lesbos, 03.04.1994, *Nielsen & Skovgaard 9613* (C); LR606934; LR606734; LR606581; LR606439. ***Nephelochloa orientalis*** Boiss.: Turkey, between Denizli and Aydin, 22.06.1976, *C. Simon 76900* (BASBG); LR606935, HE646584; –; HE646609; LR606440. ***Oreochloa blanka*** Deyl: France, Pyrénées-Orientales, Massif du Puigmal d’Err, 10.07.1991, *J. Lambinon 91/205* (B 10 0448884); LR606936; LR606735; LR606582; LR606441. ***O. disticha*** (Wulfen) Link: Romania, Jud. Hunedoara, Retezat Mts., 31.07.1992, *M. Röser 9588* (HAL); LR606937, AM234592; –; FM179421; LR606442. ***Paracolpodium altaicum*** (Trin.) Tzvelev: Russia, Altai, Kosh-Agach, Saylyugem range, 12.08.1982, *V. Khanminchun & N. Friesen 8* (ALTB); LR606938; LR606736; HF564629; LR606443. ***P. baltistanicum*** (Dickoré) Röser & Tkach: Pakistan, Baltistan, E part of Deosai plateau, 15.07.1991, *G. Miehe & S. Miehe 5105* (MR); LR606939; LR606737; LR606583; LR606444. ***Parapholis cylindrica*** (Willd.) Röser & Tkach: Cultivated in BG Halle, Germany from seed obtained from BG Copenhagen, Denmark, 08.03.2010, *s.coll.* (HAL0140597); LR606940, HE646577; LR606738; LR606584; –. ***P. filiformis*** (Roth) C.E.Hubb. (1): France, Montpellier, 05.06.1957, *Streitberg & Stohr* (HAL0081242); LR606941, HE646585, LN554446; –; HE646610; LR606446; (2): France, Languedoc-Roussillon, Hérault; cultivated in BG Halle, Germany from seed obtained from Kew’s Millennium Seed Bank, UK (no. 63085), 06.08.1986, *J. Feltwell* (HAL); –; LR606739; LR606585; LR606445. ***P. incurva*** (L.) C.E.Hubb.: Greece, Macedonia, Thessaloniki, Chalcidice, 25.05.1985, *M. Röser 2517* (HAL); LR606942, AM234583; LR606740; FM179422; LR606447. ***P. marginata*** Runemark: Greece, Lasithiou, Eparchia Sitia, Xerocampos, Katsouria, 19.06.1905, *N. Böhling 5292b* (B 10 0199860); LR606943; LR606741; LR606586; LR606448. ***Parvotrisetum myrianthum*** (Bertol.) Chrtek: Greece, Macedonia, 21.06.1970, *A. Strid 221* (C); LR606944, LT159690; LR606742; HE802174; LR606449. ***Pentapogon quadrifidus*** (Labill.) Baill. (1) var. ***quadrifidus***: Australia, South Australia, Southern Tableland, 29.10.1998, *I. Crawford & N. Taws 4887* (NSW463696); LR606946; LR606744; LR606587; LR606451; (2) var. ***parviflorus*** (Benth.) D.I.Morris: Australia, Tasmania, South West Tasmania, Nye Bay, 09.01.1986, *A. Moscal 11543* (HO 95925); LR606945; LR606743; –; LR606450. ***Periballia involucrata*** (Cav.) Janka: Portugal, Minho, Portela do Homem, Cruz do Louro, 02.06.1990, *A.I.D. Correia & A. Fernandes* (LISU 160284); LR606947; LR606745; LR606588; LR606452. ***Peyritschia pringlei*** (Scribn.) S.D.Koch: Mexico, Puebla, Mun. S. Nícolas de los Ranchos Buenavista, 05.02.1988, *P. Tenorio 15095* (MEXU 542571); LR606948, HG797458; LR606746; HG797528; LR606453. ***Phalaris arundinacea*** L.: Russia, Yakutia, middle course of Kolyma, Lobuy, 30.07.1983, *Doronkin & Bubnova 2264* (NS/NSK); LR606949; LR606747; HF564628; LR606454. ***P. canariensis*** L.: Italy, Napoli Province, Campania; cultivated in BG Halle, Germany from seed obtained from BG Berlin-Dahlem, Germany (no. 2001-3939), 15.05.2003, *Royl 173* (HAL); LR606950; LR606748; HE802173; LR606455. ***P. coerulescens*** Desf.: Italy, Siena, cultivated in BG Halle, Germany from seed obtained from BG Berlin-Dahlem, Germany (no. 2001-3940); 07.06.2004, *s.coll.* (HAL); LR606951; LR606749; HE802172; –. ***Phippsia algida*** (Sol.) R.Br.: Russia, E Taymyr, Nyunkarakutari River, Poymennoe Lake, 05.08.1998, *I.N. Pospelov 98-158* (NS); LR606952, AM234603; –; FM179424; LR606456. ***P. concinna*** (Th.Fr.) Lindeb.: Russia, Taymyr, Syndasko River, 23.07.1979, *N. Vodopyanova, R. Krogulevich, N. Frisen, V. Nikolayeva & N. Shumik 224* (NS); LR606953, AM234582; LR606750; FM179425; LR606457. ***Phleum alpinum*** L.: Austria, Styria, near St. Oswald, 31.07.2001, *M. Röser 11023* (HAL); LN554448; LR606751; LR606589; LR606458. ***P. crypsoides*** (d’Urv.) Hack.: Cyprus, Cape Greco, 15.04.1992, *F. Skovgaard* (C); LR606954; –; HE802187; LR606459. ***P. phleoides*** (L.) Karsten: Norway, Oslo; cultivated in BG Halle, Germany from seed obtained from BG Oslo, Norway (no. 2003-669), 31.07.2003, *s.coll.* (HAL); LR606955, AM234552; LR606752; FM179426; LR606460. ***Pholiurus pannonicus*** (Host) Trin.: Hungary, Great Hungarian Plane (Alföld), Hortobágy Puszta, 13.06.1967, *W. Hilbig* (HAL0067272); LR606956, HE646586; LR606753; HE646616–HE646625 (clones consensus); LR606461. ***Poa annua*** L.: Germany, Saxony-Anhalt, 18.01.2005, *M. Röser 11065* (HAL); LR606957, AM234593; LR606754; FM179428; LR606462. ***P. bulbosa*** L.: Austria, Lower Austria, near Eggenburg, 28.04.1991, *M. Röser 7419* (HAL); LR606959, AM234594; LR606756; FM179429; LR606464. ***P. cyrenaica*** E.A.Durand & Barratte: Libya, Bengasi, 29.01.1924, *F. Lavara & L. Grande* (FI); LR606960; –; HE802196; LR606465. ***P. diaphora*** Trin.: Mongolia, Bajan Ölgi Aimak, 27.07.1977, *W. Hilbig* (HAL0044036);, LR606961; LR606757; HE802188; –. ***P. fax*** J.H.Willis & Court: Australia, South Australia, Coffin Bay Conservation Park, 08.10.1991, *D. E. Murfet 1278* (AD99151120); LR606962; LR606758; HE802191; LR606466. ***P. hitchcockiana*** Soreng & P.M.Peterson: Ecuador, Province Loja, Cajanuma, 05.03.1987, *I. Grignon* (MO5151808); LR606963; –; HE802195; –. ***P. labillardierei*** Steud.: Australia, Nora Creina, 11.10.1989, *P. C. Heyligers 89162* (AD99151199); LR606958; LR606755; HE802193; LR606463. ***P. lepidula*** (Nees & Meyen) Soreng & L.J.Gillespie (1): Peru, Department Moquegua, Provincia Mariscal Nieto, 01.03.1999, *P.M. Peterson* (MO5151809); LR606964, FR694884; LR606759; –; LR606467; (2): Chile, Tarapacá Region (Region I), Chungará, 04.04.2001, *P.M. Peterson 15759 & R.J. Soreng* (MO5698870); FR694884; –; FR692034; –. ***P. persica*** Trin.: Turkmenistan, Geok-Tepinskiy District, Central Kopet-Dag, 12.07.1969, *A.A. Mescheryakov* (LE); LR606965; LR606760; HE802189; LR606468. ***P. pratensis*** L.: Germany, Saxony-Anhalt, Dessau-Roßlau, 14.05.2009, *E. Willing 25.267 D* (HAL0109437); LR606966; LR606761; LR606590; LR606469. ***P. serpaiana*** Refulio: Chile, Tarapacá Region (Region I); Parinacota, 04.04.2001, *P.M. Peterson & R.J. Soreng* (MO5698869); LR606967; LR606762; HE802194; LR606470. ***P. sintenisii*** H.Lindb.: Cyprus, Ayios Nikolaos; Kew DNA Bank, London (no. 24200), 01.11.1988, *Meikle 2853* (K); LR606968; LR606763; HE802190; LR606471. ***Podagrostis aequivalvis*** (Trin.) Scribn. & Merr.: Canada, British Colombia, Queen Charlotte Islands, Moresby Island, 25.06.1957, *J.A. Calder 21762, D.B.O. Savile & R.L. Taylor* (B 10 0448891); LR606969; LR606764; LR606591; LR606472. ***P. thurberiana*** (Hitchc.) Hultén: USA, Washington, Kittitas County, Beverly Creek, 25.08.2000, *R.J. Soreng 6356* (US); –; LR606765; LR606592; LR606473. ***Psilathera ovata*** (Hoppe) Deyl: Austria, Tyrol, Grossglockner Mountain, Hochtor, 05.09.2017, *M. Röser 11318 & N. Tkach* (HAL); –; LR606766; –; LR606474. ***Puccinellia fasciculata*** (Torr.) E.P.Bicknell: Hungary, Hajdú-Bihar county, Hortobágy Puszta, 27.05.1991, *M. Röser 7633* (HAL); LR606970, AM234588, LN554450; LR606767; FM179431; LR606475. ***P. vahliana*** (Liebm.) Scribn. & Merr.: Denmark, W Greenland, Disko, Nodfjord, Stordal, 14.08.1975, *L. Andersen & B. Fredskild* (LE); LR606971; LR606768; LR606593; LR606476. ***Relchela panicoides*** Steud.: Chile, Andes, Malleco Province, Fundo Solano, Los Alpes, 13.01.1958, *W.J. Eyerdam 10152* (NY); LR606972, LT159692; LR606769; LT159756; LR606477. ***Rhizocephalus orientalis*** Boiss. (1): Turkmenistan, Geok-Tepinskiy District, Central Kopet-Dag, 03.06.1952, *V.V. Nikitin* (LE); LR606974; LR606770; LR606594; LR606478; (2): Turkmenistan, Geok-Tepinskiy District, Central Kopet-Dag, 04.06.1952, *V.V. Nikitin & A.A. Mescheryakov* (LE); LR606975; LR606771; LR606595; LR606479. ***Rostraria cristata*** (L.) Tzvelev: Cultivated in BG Halle, Germany from seed obtained from BG Dijon, France (no. 2001-1130), 19.08.2002, *M. Röser 11081* (HAL); LR606976, AM234670; LR606772; LT159757; LR606480. ***Sclerochloa dura*** (L.) P.Beauv. (1): Hungary, Veszprém, between Balatonakali and Balatonudvari, 25.05.1991, *M. Röser 7527* (HAL); LR606977, AM234587; LR606773; FM179433; LR606481; (2): Germany, Thuringia, Kyffhäuser, Gorsleben, 31.05.2016, *M. Röser 11255 & N. Tkach* (HAL); –; LR655822; LR655821; –. ***S. festucacea*** (Willd.) Link (1): Russia, Irkutsk Oblast, Kasachinskoye, 27.08.1982, *A. Kiseleva & T. Takmanova 403* (NS/NSK); LR606978, AM234600; –; LR606596; –; (2): Germany, Potsdam, 03.08.2016, *M. Röser 11281 & N*. *Tkach* (HAL); –; LR606774; –; LR606482. ***Secale sylvestre*** Host: Hungary, Bács-Kiskun, Bugac Puszta, 26.05.1999, *M. Röser 10954* (HAL); LR606979, AM234581, LN554452; LR606775; FM179434; LR606483. ***Sesleria argentea*** (Savi) Savi: Cultivated in BG Halle, Germany from seed obtained Museum National d’Histoire Naturelle Paris, France (no. 2008-44), no voucher; LR606980; LR606776; –; LR606484. ***S. caerulea*** (L.) Ard.: Germany, Thuringia, Harz Mts., 15.05.2016, *M. Röser 11239 & N. Tkach* (HAL); –; LR606777; LR606597; –. ***S. insularis*** Sommier: Italy, Sardinia, Nuoro Province, Golfo di Orosei, Mt. Tuttavista, 10.05.1993, *M. Röser 10166* (HAL); LR606981, AM234591; LR606778; FM179435; LR606485. ***S. varia*** (Jacq.) Wettst.: Austria, Tyrol, Grossglockner Mountain, Edelweissspitze, 05.09.2017, *M. Röser 11321 & N. Tkach* (HAL); –; LR606779; LR606598; LR606486. ***Sesleriella sphaerocephala*** (Ard.) Deyl (1): Slovenia, Gorenjska, Julian Alps, summit of Mt. Lanževica, s.d., *B. Frajman S024* (IB 12825); LR606983, LN554453; LR606781; LR606600; LR606488; (2): Austria, Carinthia, Karavankes, 16.06.1991, *M. Röser 7867* (HAL); LR606982, AM234590; LR606780; LR606599; LR606487. ***Sibirotrisetum sibiricum*** (Rupr.) Barberá (1): China, Qinghai, surroundings of Menyang, 29.07.2004, *I. Hensen* (HAL); –; LR606782; LT159800; –; (2): Russia, Lake Baikal, Olchon Island; cultivated in BG Halle, Germany from seed, 25.07.2006, *H. Heklau* (HAL); LR606984, LT159706; LR606783; –; LR606489. ***Simplicia buchananii*** (Zotov) Zotov: New Zealand, Nelson Land District, 13.03.1984, *A.P. Druce* (CHR 394262); LR606985; LR606784; HE802177; LR606490. ***Sphenopholis intermedia*** (Rydb.) Rydb. (1): Canada, Little Manitou Lake, 20.08.1992, *Hudson 5083* (CAN565509); LR606986, HG797460; –; HG797530; –; (2): USA, Illinois; cultivated in BG Halle, Germany from seed obtained from Kew’s Millennium Seed Bank, UK (no. 307008), 07.11.2005, *s.coll.* (HAL); –; LR606785; –; LR606491. ***S. obtusata*** (Michx.) Scribn.: USA, Kansas; cultivated in BG Halle, Germany from seed obtained from Kew’s Millennium Seed Bank, UK (no. 408330); 08.06.2007, *J. Hansen* (HAL); LR606987, HG797462, LN554455; LR606786; HG797532; LR606492. ***Sphenopus divaricatus*** (Gouan) Rchb.: Spain, Aragon, Province Huesca, 11.05.1980, *G. Montserrat 38080* (RO); LR606988, HE646589; LR606787; HE646613; LR606493. ***Torreyochloa pauciflora*** (J.Presl) Church: USA, Alaska, Haines, Chilkoot Lake Road, 17.08.2000, *R.J. Soreng 6327* (US Catalog No.: 3679690, Barcode: 01259790), LR606989; LR606788; LR606601; LR606494. ***Tricholemma jahandiezii*** (Litard. ex Jahandiez & Maire) Röser: Morocco, Moyen Atlas, 02.07.2002, *M. Röser 10297/1B* (HAL); LR606990, AM234556, HG797464; LR606789; FM179407, FM956101; LR606495. ***Trisetaria panicea*** (Lam.) Paunero: Portugal, Province Beira Alta, Serra da Estrela, Rio Zêzere-Tale, 12.07.1992, *M. Röser 9473* (HAL); LR606991, HG797465; LR606790; HG797534; LR606496. ***Trisetopsis aspera*** (Hook.f.) Röser & A.Wölk: Sri Lanka (Ceylon), Horton Plains, Badulla District, Province Uva, 27.01.1970, *D. Clayton 5505* (CANB); LR606992; LR606791; LR606602; LR606497. ***T. elongata*** (Hochst. ex A.Rich.) Röser & A.Wölk: Uganda, Mount Elgon, 23.02.2004, *K. Wesche* (HAL); LR606993, HG797469; LR606792; HG797566; LR606498. ***T. imberbis*** (Nees) Röser, A.Wölk & Veldkamp: South Africa, Western Cape, Betty’s Bay, corner Kreupel hout street and Lipkin road, 25.10.2010, *A.C. Mudau & L. Smook 452* (PRE); LR606994, HG797483; LR606793; HG797631; LR606499. ***T. longa*** (Stapf) Röser & A.Wölk: South Africa, Western Cape, Table Mountain National Park, Jonkersdam, 23.10.2010, *A.C. Mudau & L. Smook 450* (PRE); LR606995, HG797475; LR606794; HG797597; LR606500. ***T. turgidula*** (Stapf) Röser & A.Wölk: Lesotho, Ligholong, Mine, 02.01.1900, *T. Edwards 7141* (NU4-2005/15), –; –; –; LR606501. ***T. virescens*** (Nees ex Steud.) Röser & A.Wölk: Pakistan, Hazara, Himalaya foothills, Indus Kohistan, 28.08.1995, *B. Dickoré 12063* (MSB); –; LR606795; LT159791; LR606502. **×*Trisetopsotrichon altius*** (Hitchc.) Röser & A.Wölk: China, Sechuan, Nereku River, 26.07.1885, *G.N. Potanin* (LE); LR606996; LR606796; LT159792; LR606503. ***Trisetum canescens*** Buckley: USA, Oregon, Josephine, Cave Creek, 02.06.2000, *R.J. Soreng 5956* (US); LR606997, AM234611; LR606797; LR606603; LR606504. ***T. cernuum*** Trin.: USA, Montana, Glacier County, Alon Continental Divide, 19.07.2003, *P. Lesica 8714* (NY1819808); LR606998, LT159703; LR606798; LT159797; LR606505. ***T. flavescens*** (L.) P.Beauv.: Germany, Baden-Württemberg, near Tübingen, 26.06.1984, *M. Röser 1871* (HAL); LR606999; LR606799; LR606604; LR606506. ***T. spicatum*** (L.) K.Richt.: USA, Alaska, Dalton Hwy, Chandler Shelf, 05.08.2000, *R.J. Soreng 6221* (US Catalog No.: 3682816, Barcode: 01259847), LR607000, LT159707; LR606800; LT159801; LR606507. ***Tzveleviochloa parviflora*** (Hook.f.) Röser & A.Wölk: Bhutan, Thimphu, 18.07.2000, *G. Miehe & S. Miehe 00-223-32* (MR); LR607001, LT159708; LR606801; LT159802; LR606508. ***Vahlodea atropurpurea*** (Wahlenb.) Fr. ex Hartm.: Canada, British Columbia, Haines Hwy., Chilkat Pass, 15.08.2000, *R.J. Soreng 6316* (US); LR607002, AM234549; LR606802; FM179439; LR606509. ***Ventenata blanchei*** Boiss.: Syria, Djebel Ed Drouz, 09.05.1933, *G. Samuelsson* (C); LR607003; –; LR606605; LR606510. ***V. dubia*** (Leers) Coss.: Bulgaria, East Stara Planina Mts., 26.05.1999, *T. Raus, F. Pina Gata 21-1-5* (B 10 0417270); LR607004; LR606803; LR606606; –. ***V. macra*** (Steven ex M.Bieb.) Balansa ex Boiss.: Greece, Peloponnese, Achaia, 10.08.1998, *M. Röser 10688* (HAL); LR607005, AM234555; LR606804; FM179440; LR606511. ***Vulpiella stipoides*** (L.) Maire: Libya, Tripolitania, Jebel Nefoussa Zintan, 30.04.1965, *H. Eckerlein* (HAL0016576); LR607006, HE646591; LR606805; HE646614; LR606512.

## Appendix 2

Publicly available DNA sequences from ENA/GenBank used in this study. Sequences included in the final alignments (suppl. Appendix S1) for the phylogenetic reconstructions are marked by an asterisk (see Material and Methods). The taxon name is followed by ENA/GenBank accession numbers for plastid *matK* gene–3’*trnK* exon; plastid *trnL–trnF*; nuclear ribosomal ITS1–5.8S gene–ITS2; nuclear ribosomal ETS. A dash indicates unavailable or unused sequences.

***Agrostis alopecuroides*** Lam.: DQ786937; –; –; –. ***A. avenacea*** J.F.Gmel.: HE574415; –; –; –. ***A. capillaris*** L.: –; –; –; JX438119. ***A. linkii*** Banfi, Galasso & Bartolucci: –; DQ631457; –; –. ***A. scabra*** Willd.: DQ146807*; KX372376; –; –. ***Aira praecox*** L.: EF137480; EF137588; –; –. ***Airopsis tenella*** (Cav.) Coss. & Durieu: KJ529354; DQ631445; –; –. ***Alopecurus aequalis*** Sobol.: KM538789, KM523821; KM524037, EU639572; –; KM523673. ***Ammochloa palaestina*** Boiss.: –; DQ631451; –; –. ***Aniselytron treutleri*** (Kuntze) Soják: KM523839; EU792441; EU792373*; GQ324239*, GQ324240. ***Anthoxanthum arcticum*** Veldkamp: –; KC698978*; –; –. ***A. australe*** (Schrad.) Veldkamp: –; DQ631447; –; –. ***A. monticola*** (Bigelow) Veldkamp: –; DQ353953; –; GQ324241. ***A. nitens*** (Weber) Y.Schouten & Veldkamp: EF137503; –; –; KC898002. ***A. odoratum*** L.: DQ786884, EF137484; KC897747; –; –. ***A. redolens*** (Vahl) P.Royen: –; KC897757; –; KC898003. ***A. repens*** (Host) Veldkamp: –; KC698990; –; –. ***Antinoria agrostidea*** (DC.) Parl.: KJ529360; –; –; –. ***Apera interrupta*** (L.) P.Beauv.: EF137485*, KM523842; EU792439*; EU792364*; GQ324242*. ***Arctagrostis latifolia*** (R.Br.) Griseb.: DQ786885, KM523924, KM523844; DQ353969; –; GQ324243, GQ324244, GQ324245. ***Arctohyalopoa lanatiflora*** (Roshev.) Röser & Tkach: –; –; FJ178781; –. ***Arctopoa eminens*** (J.Presl) Prob.: KM523848; DQ353977; –; GQ324247, GQ324248, GQ324249. ***A. subfastigiata*** (Trin.) Prob.: KM523849*; EU792449*; EU792372*; GQ324250*. ***A. tibetica*** (Munro ex Stapf) Prob.: KM523850*; EU792444*; GQ324471*; GQ324252*. ***Arrhenatherum elatius*** (L.) P.Beauv. ex J.Presl & C.Presl: EU434292, EF137486, KJ529335; JF904748; –; –. ***Avellinia michelii*** (Savi) Parl.: KJ529340; DQ631465; –; –. ***Avena hispanica*** Ard.: GU367287, GU367288, EU833849; EU833874; –; –. ***A. macrostachya*** Balansa ex Coss. & Durieu: EU833852; EU833877; –; –. ***Avenella flexuosa*** (L.) Drejer: DQ786887; AY237913; –; –. ***Avenula pubescens*** (Huds.) Dumort.: EF137502; DQ631460; –; –. ***Bellardiochloa polychroa*** (Trautv.) Roshev.: –; –; –; GQ324256. ***B. variegata*** (Lam.) Kerguélen: DQ786890, KM523852; –; –; GQ324257. ***Boissiera squarrosa*** (Sol.) Nevski: EF137488; –; –; KJ632438, KP996869, KP996870. ***Brachypodium distachyon*** (L.) P.Beauv.: –; KU163229; AF303399; –. ***Briza media*** L.: –; EU395902*; –; –. ***B. minor*** L.: KJ599228, DQ786892; –; –; KJ599006. ***Bromus erectus*** Huds.: –; JX985261; –; –. ***Calamagrostis arenaria*** (L). Roth subsp. ***arundinacea*** (Husn.) Banfi, Galasso & Bartolucci: KJ529326; DQ631456; –; JX438118. ***C. arundinacea*** (L.) Roth: DQ786895; KX372396; GQ266675; –. ***C. canadensis*** (Michx.) P.Beauv.: –; –; FJ377628*; –. ***C. purpurascens*** R.Br.: –; FJ394570, FJ394568; –; –. ***Castellia tuberculosa*** (Moris) Bor: EF137492*; EF137596*; AF532954*; –. ***Catabrosa aquatica*** (L.) P.Beauv.: DQ786898, KM523853; DQ353958*; –; KM523697, GQ324258. ***C. werdermannii*** (Pilg.) Nicora & Rúgolo: –; EU792431*; EU792333*; GQ324259*. ***Catabrosella variegata*** (Boiss.) Tzvelev: KM523854; –; –; KM523698*. ***Catapodium marinum*** (L.) C.E.Hubb.: KJ529348; –; –; –. ***C. rigidum*** (L.) C.E.Hubb.: EF137491, KJ599274; AF533034; –; –. ***Chascolytrum bulbosum*** (Parodi) Essi, Longhi-Wagner & Souza-Chies: –; EU395894; –; –. ***C. subaristatum*** (Lam.) Desv.: DQ786899, KJ599293; –; –; KJ599067. ***C. uniolae*** (Nees) Essi, Longhi-Wagner & Souza-Chies: –; EU395874; –; –. ***Cinna latifolia*** (Trevir. ex Göpp.) Griseb.: KM523855; GQ324396; –; GQ324261. ***Colpodium versicolor*** (Steven) Schmalh.: KM523856; KM524063*; –; KM523699. ***Corynephorus canescens*** (L.) P.Beauv.: KJ529351; DQ631440; –; –. ***Cutandia maritima*** (L.) Barbey: KJ529370; AF487618; –; –. ***Cynosurus cristatus*** L.: DQ786901, HM453075, KJ599277; KF876179*; –; –. ***Dactylis glomerata*** L.: EF137494, KJ599276; AF533028; –; KJ599050*. ***Deschampsia cespitosa*** (L.) P.Beauv.: KM523858, DQ786903, EF137495; AY237912; –; –. ***D. setacea*** (Huds.) Hack.: –; DQ631479*; DQ539615*; –. ***Desmazeria sicula*** (Jacq.) Dumort.: DQ786904; EF592948; –; –. ***Dichelachne crinita*** (L.f.) Hook.f.: HE574411; –; –; –. ***D. micrantha*** (Cav.) Domin: DQ786906; –; –; –. ***Drymochloa sylvatica*** (Pollich) Holub: HM453070, KJ529372; AF478505; –; –. ***Dryopoa dives*** (F.Muell.) Vickery: KJ599286, KJ599326; KJ599438; –; –. ***Dupontia fisheri*** R.Br. subsp. ***psilosantha*** (Rupr.) Hultén: DQ786908, KM523859, KM523860, KM523925, KM523926; –; –; KM523702, KM523701, GQ324267, GQ324266. ***D. fulva*** (Trin.) Röser & Tkach: KM523845, KM523846, KM523847; KM524058; –; KM523694, KM523695, GQ324246. ***Dupontiopsis hayachinensis*** (Koidz.) Soreng, L.J.Gillespie & Koba: KM523861*; KM524066*; KM523779*; KM523703*. ***Echinaria capitata*** (L.) Desf.: KJ529361; DQ631453; –; –. ***Echinopogon caespitosus*** C.E. Hubb.: DQ786909, HE574414; –; –; –. ***Festuca floribunda*** (Pilg.) P.M.Peterson, Soreng & Romasch.: DQ786907, JF697821; JF904750; –; –. ***F. incurva*** (Gouan) Gutermann: KJ599280; AF478533; –; KJ599053. ***F. lachenalii*** (C.C.Gmel.) Spenn.: KJ529387; AF478534*; –; –. ***F. maritima*** L.: KJ529388; AY118107; –; –. ***F. myuros*** L.: KJ599273, AF164403; AY118103; KJ598937*; KJ599048*. ***F. salzmannii*** (Boiss.) Boiss. ex Coss.: –; AF478535; –; –. ***Gastridium nitens*** (Guss.) Coss. & Durieu: DQ786945, KJ529331; DQ336836; –; –. ***G. ventricosum*** (Gouan) Schinz & Thell.: FN908056, DQ786914; DQ336837; –; HE575740. ***Gaudinia fragilis*** (L.) P.Beauv.: DQ786915, EF137499; DQ631478; –; –. ***Graphephorum wolfii*** J.M.Coult.: DQ786917; DQ336843; –; –. ***Helictochloa aetolica*** (Rech.f.) Romero Zarco: –; EU792437; –; KM523706. ***H. bromoides*** (Gouan) Romero Zarco subsp. bromoides: KJ529356; DQ631459; –; –. ***H. hookeri*** (Scribn.) Romero Zarco: DQ786888; HM590299; –; –. ***Helictotrichon convolutum*** (C.Presl) Henrard: DQ786919, KM523865; DQ353954; –; KM523707. ***H. sedenense*** (DC.) Holub: –; –; –; KC899016. ***H. sempervirens*** (Vill.) Pilg.: –; DQ353955; –; GQ324269*. ***Hookerochloa eriopoda*** (Vickery) S.W.L.Jacobs: DQ786913, KJ599294, KM523866; GQ324397*; –; GQ324270, GQ324271. ***H. hookeriana*** (F.Muell. ex Hook.f.) E.B.Alexeev: KJ599295, DQ786922, KM523867; EU792435; –; KJ599068, GQ324272, KM523708. ***Hordelymus europaeus*** (L.) O.E.Harz: –; EU119368*; –; –. ***Hordeum marinum*** Huds.: –; AB732935; –; KJ632437*. ***Hyalopoa pontica*** (Balansa) Tzvelev: KM523868*; KM524070*; EU792365, FJ196302, FJ196303; KM523709*. ***Koeleria pyramidata*** (Lam.) P.Beauv.: EF137505; EU119370; –; –. ***Lagurus ovatus*** L.: –; DQ631464; –; –. ***Lamarckia aurea*** (L.) Moench: KJ599279; KJ599392; –; KJ599052*. ***Limnas stelleri*** Trin.: –; –; –; KM523710*. ***Littledalea tibetica*** Hemsl.: DQ786924; –; –; –. ***Lolium giganteum*** (L.) Darbysh.: HM453058; AF533043; –; –. ***L. perenne*** L.: DQ786925*; EF378973*; KJ598999*; KJ599109*. ***L. rigidum*** Gaudin: DQ786926*, KJ599336; EF378980*; KJ599000*; KJ599110*. ***Macrobriza maxima*** (L.) Tzvelev: FN908048; EU395901; –; –. ***Mibora minima*** (L.) Desf.: DQ786927, KJ529357; DQ631454; –; –. ***Milium effusum*** L.: KM523869, KM523870; KM524072*; –; KM523711, GQ324273. ***Molineriella laevis*** (Brot.) Rouy: DQ786929; KJ529413; –; –. ***Nephelochloa orientalis*** Boiss.: KM523873; KM524075*; –; KM523714. ***Nicoraepoa andina*** (Trin.) Soreng & L.J.Gillespie: DQ786934*, KM523874; DQ353971*; EU792354*; GQ324275*. ***Oreochloa disticha*** (Wulfen) Link: –; DQ631452*; –; –. ***Paracolpodium altaicum*** (Trin.) Tzvelev: KM523878; KM524076; –; KM523715. ***Parapholis cylindrica*** (Willd.) Röser & Tkach: EF137501, KJ599283, KJ529366; KJ599395; –; KJ599056*. ***P. filiformis*** (Roth) C.E.Hubb.: KJ529365; KJ529415; –; –. ***P. incurva*** (L.) C.E.Hubb.: DQ786931, EF137508, KJ599281; –; –; KJ599054. ***Periballia involucrata*** (Cav.) Janka: KJ529353; DQ631438; –; –. ***Peyritschia pringlei*** (Scribn.) S.D.Koch: –; FJ394581; –; –. ***Phalaris arundinacea*** L.: AF164396; JF951096; –; –. ***P. canariensis*** L.: –; DQ631443; –; –. ***P. coerulescens*** Desf.: KJ529325; JF951116; –; –. ***Phippsia algida*** (Sol.) R.Br.: KM523879, KM523880; KM524078*; –; KM523716, KM523717, GQ283228, GQ283229. ***Phleum alpinum*** L.: KM523881; KM524079; –; KM523718. ***P. phleoides*** (L.) Karsten: KM523884; KM524082; –; KM523718. ***Poa alpina*** L.: DQ786933*, KM523888; DQ353986*; EU792390*; –. ***P. annua*** L.: KJ599339, KJ599340; EU792452; –; KJ599113, KJ599114. ***P. apiculata*** Refulio: –; EU792469*; EU792428*; KU763389*. ***P. bulbosa*** L.: KJ529342, KJ599341; AH015559; –; KJ599115, GQ324297, GQ324298. ***P. diaphora*** Trin.: –; KJ746808; –; GQ324311*. ***P. fax*** J.H.Willis & Court: KJ599238; EU792460; –; KJ599016, KJ599065, GQ324318. ***P. hitchcockiana*** Soreng & P.M.Peterson: –; –; –; KU763378 *. ***P. labillardierei*** Steud.: DQ786935, KJ599324; AH015564; –; KJ599097, GQ324296. ***P. lepidula*** (Nees & Meyen) Soreng & L.J.Gillespie: –; AH015563*, EU792471; –; GQ324343, GQ324344. ***P. pratensis*** L.: KJ599260, KJ599261; JF904790; –; GQ324369, KJ599036. ***P. serpaiana*** Refulio: –; AH015566; –; GQ324265, KU763451. ***Podagrostis thurberiana*** (Hitchc.) Hultén: DQ786936*; –; –; –. ***Puccinellia arctica*** (Hook.) Fernald & Weath.: –; –; GQ283100 –. ***P. borealis*** Swallen: –; –; GQ283160 –. ***P. ciliata*** Bor: –; –; KJ598984 –. ***P. distans*** (Jacq.) Parl.: –; –; KP711085 –. ***P. fasciculata*** (Torr.) E.P.Bicknell: KJ599321; –; KJ598985; KJ599094. ***P. frigida*** (Phil.) I.M.Johnst.: –; –; JF904809 –. ***P. glaucescens*** (Phil.) Parodi: –; –; EU792338 –. ***P. interior*** T.J.Sørensen ex Hultén: –; –; KM523808 –. ***P. longior*** A.R.Williams: –; –; KJ598961 –. ***P. magellanica*** (Hook.f.) Parodi: –; –; KM523810 –. ***P. parishii*** Hitchc.: –; –; GQ283123 –. ***P. perlaxa*** (N.G.Walsh) N.G.Walsh & A.R.Williams: –; –; KJ598986 –. ***P. phryganodes*** (Trin.) Scribn. & Merr.: –; –; GQ283157 –. ***P. pumila*** (Macoun ex Vasey) Hitchc.: –; –; GQ283158 –. ***P. stricta*** (Hook.f.) C.H.Blom: –; –; EU792339 –. ***P. tenella*** (Lange) Holmb.: –; –; GQ283110 –. ***P. tenuiflora*** (Griseb.) Scribn. & Merr.: –; –; KP711084 –. ***P. vahliana*** (Liebm.) Scribn. & Merr.: KM523915; –; –; GQ283185, GQ283186, GQ283187, GQ283188, GQ324285. ***P. vassica*** A.R.Williams: –; –; KJ598963 –. ***P. walkeri*** (Kirk) Allan subsp. ***chathamica*** (Cheeseman) Edgar: –; –; EU331103 –. ***Relchela panicoides*** Steud.: –; JF904801; –; –. ***Rostraria cristata*** (L.) Tzvelev: –; DQ336853, GQ324465; –; –. ***Saxipoa saxicola*** (R.Br.) Soreng, L.J.Gillespie & S.W.L.Jacobs: KJ599265, KM523917*; GQ324465*; GQ324558*; GQ324392*. ***Sclerochloa dura*** (L.) P.Beauv.: DQ786941, KJ599275, KM523918; KM524102; –; KM523745, KJ599049, KJ632435. ***Scolochloa festucacea*** (Willd.) Link: KM523919; KM524103; –; KM523746*. ***Sesleria argentea*** (Savi) Savi: –; AF533030; –; –. ***S. insularis*** Sommier: KM523920; DQ353957; –; KM523747. ***Sibirotrisetum sibiricum*** (Rupr.) Barberá: –; KX372500; –; –. ***Simplicia buchananii*** (Zotov) Zotov: –; HM191465; –; HM191451, HM191452, HM191453. ***Sphenopholis intermedia*** (Rydb.) Rydb.: –; DQ631466; –; –. ***S. obtusata*** (Michx.) Scribn.: –; EU119377; –; –. ***Sphenopus divaricatus*** (Gouan) Rchb.: DQ786943; AF533033; –; –. ***Sylvipoa queenslandica*** (C.E.Hubb.) Soreng, L.J.Gillespie, & S.W.L.Jacobs: KJ599262*, KM523921; GQ324466*; GQ324559*; GQ324393*. ***Torreyochloa pauciflora*** (J.Presl) Church: DQ786944; –; –; –. ***Trisetaria panicea*** (Lam.) Paunero: –; DQ631474; –; –. ***Trisetum canescens*** Buckley: DQ786946; –; –; –. ***T. cernuum*** Trin.: DQ786946; –; –; –. ***T. flavescens*** (L.) P.Beauv.: –; JQ041860; –; –. ***T. spicatum*** (L.) K.Richt.: –; FJ394585; –; –. ***Vahlodea atropurpurea*** (Wahlenb.) Fr. ex Hartm.: DQ786947; AM041251; –; –. ***Ventenata dubia*** (Leers) Coss.: KM523922; KM524104; –; KM523748*. ***V. macra*** (Steven ex M.Bieb.) Balansa ex Boiss.: KM523863; KM524068; –; KM523705.

## Appendix 3

Questionable or wrong DNA sequences in repositories ENA/GenBank.

**A.** In the course of this study we came across some errors that we made in previous publications of our lab. The errata et corrigenda are as follows:

***Hierochloe occidentale* Buckley. –** The earlier published *matK* sequence (AM234562; Döring & al., 2007; Döring, 2009; Schneider & al., 2009) does not belong to *Hierochloe* or *Anthoxanthum* as evident from comparison with the DNA sequences of other species. A sample switching error in our lab or in the field seems likely.

***Hyalopoa* (*Arctohyalopoa*) *lanatiflora* (Roshev.) Tzvelev. –** Our *matK* gene sequence AM234604 (Döring & al., 2007; Döring, 2009) obtained from leaves taken from the herbarium specimen “Russia, Yakutskaya SSSR, Ordzhonikidzevskiy rayon, surroundings of the village Kytyl-Dyura, 22.07.1988, *Zuev & Agaltsev* 434, det. O. Nikiforova” (NSK) belongs to a species of *Poa* and not to *Hyalopoa* (*Arctohyalopoa*). Although we do not have the voucher specimen at hand to verify the identification, a re-examination of a photograph taken clearly corroborates that the inflorescences belong to this taxon but it cannot be ruled out that the very dense tufts of leaves, from which the sample for DNA study was gathered, is a mixture of different grasses. In this study, two other DNA extractions from herbarium specimens unambiguously representing *Arctohyalopoa lanatiflora* were used. They yielded ITS/ETS and chloroplast DNA sequences that were identical, respectively (see Appendix 1).

***Dryopoa dives* (F.Muell.) Vickery. –** Our ITS sequence HE802192 submitted as *Poa dives* F.Muell. (Hoffmann & al., 2013) belongs to a species of *Poa* and not to *Dryopoa*.

**B.** DNA sequences taken from ENA/GenBank that turned out to be questionable or wrong according to the results of this study are as follows:

***Hyalopoa* (*Arctohyalopoa*) *lanatiflora*. –** The ITS sequence FJ178781 (Rodionov & al., 2008) clusters with the sequences of *Catabrosa aquatica*, *C. werdermannii* (EU792333) and further ENA/GenBank entries for *Catabrosa* (not shown) and disagrees with our sequences for true *H.* (*Arctohyalopoa*) *lanatiflora*.

***Hyalopoa pontica* (Bal.) Tzvelev. –** The ITS sequence of *H.* correspond to sequencesFJ196303 (Rodionov & al., 2008) and EU792365 (Gillespie & al., 2008), all of which are nested within Coleanthinae. A deviant ITS sequence reported for *H. pontica* (FJ196302; see Rodionov & al., 2008, Nosov & al., 2015, 2019) clusters among the sequences of *Poa*. The presumed occurrence of different ITS copies in *H. pontica* was discussed to rest on genetic introgression of *Poa* into *Hyalopoa* or allopolyploidy with subsequent loss of one of the parental *Hyalopoa* rDNAs (Nosov & al., 2015; Rodionov & al. 2017). The issue warrants further investigation.

***Ammochloa palaestina* Boiss. –** The ITS sequence DQ539587 (Quintanar & al., 2005: Fig. 5) belongs to a species of the genus *Helictochloa* and not to *Ammochloa*.

***Macrobriza maxima* (L.) Tzvelev**. – The ETS sequence KJ599007 submitted as *Briza maxima* L. (Birch & al., 2014) belongs to a species of *Agrostis*.

