## Supplementary Figure S1 for "Phylogeny, morphology and the role of hybridization as driving force of evolution in grass tribes Aveneae and Poeae (Poaceae)"

**Supplementary Figure S1.**
Maximum Likelihood phylogram of Poodae (Aveneae and Poeae) inferred from a concatenated matrix of plastid (matK gene–3'trnK exon, trnL–trnF) and nr (ITS, ETS) DNA sequences with species of Triticodae and Brachypodieae as outgroup.
Maximum Likelihood and Maximum Parsimony bootstrap support values >50% as well as Bayesian posterior probabilities >0.5 are indicated on the branches.
Clades with Maximum Likelihood support <50% are collapsed.
The subtribes mentioned in the text are labelled on the right-hand side.

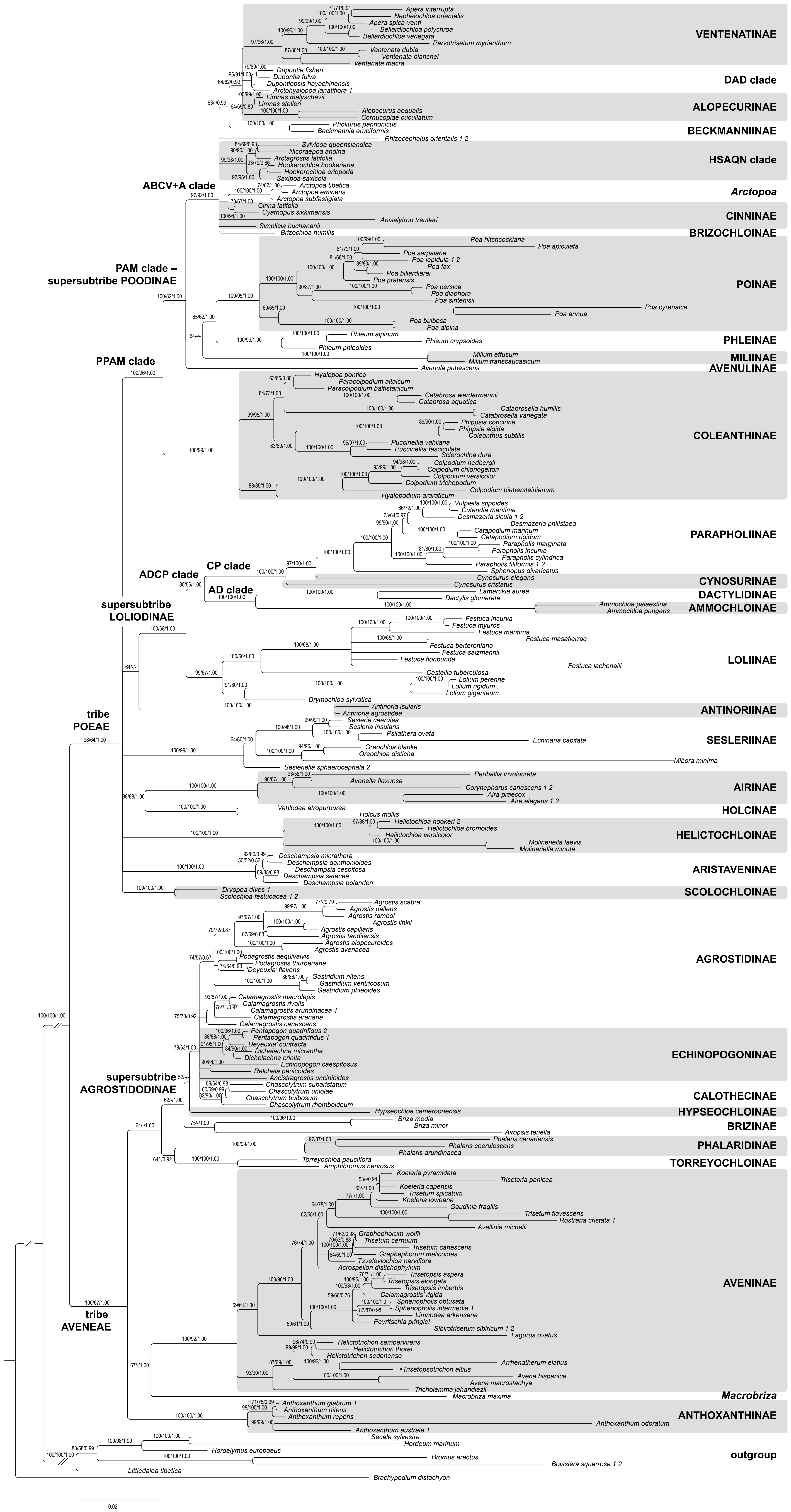
