## Supplementary figures and images for "Phylogeny, morphology and the role of hybridization as driving force of evolution in grass tribes Aveneae and Poeae (Poaceae)"

### Supplementary Figure S2

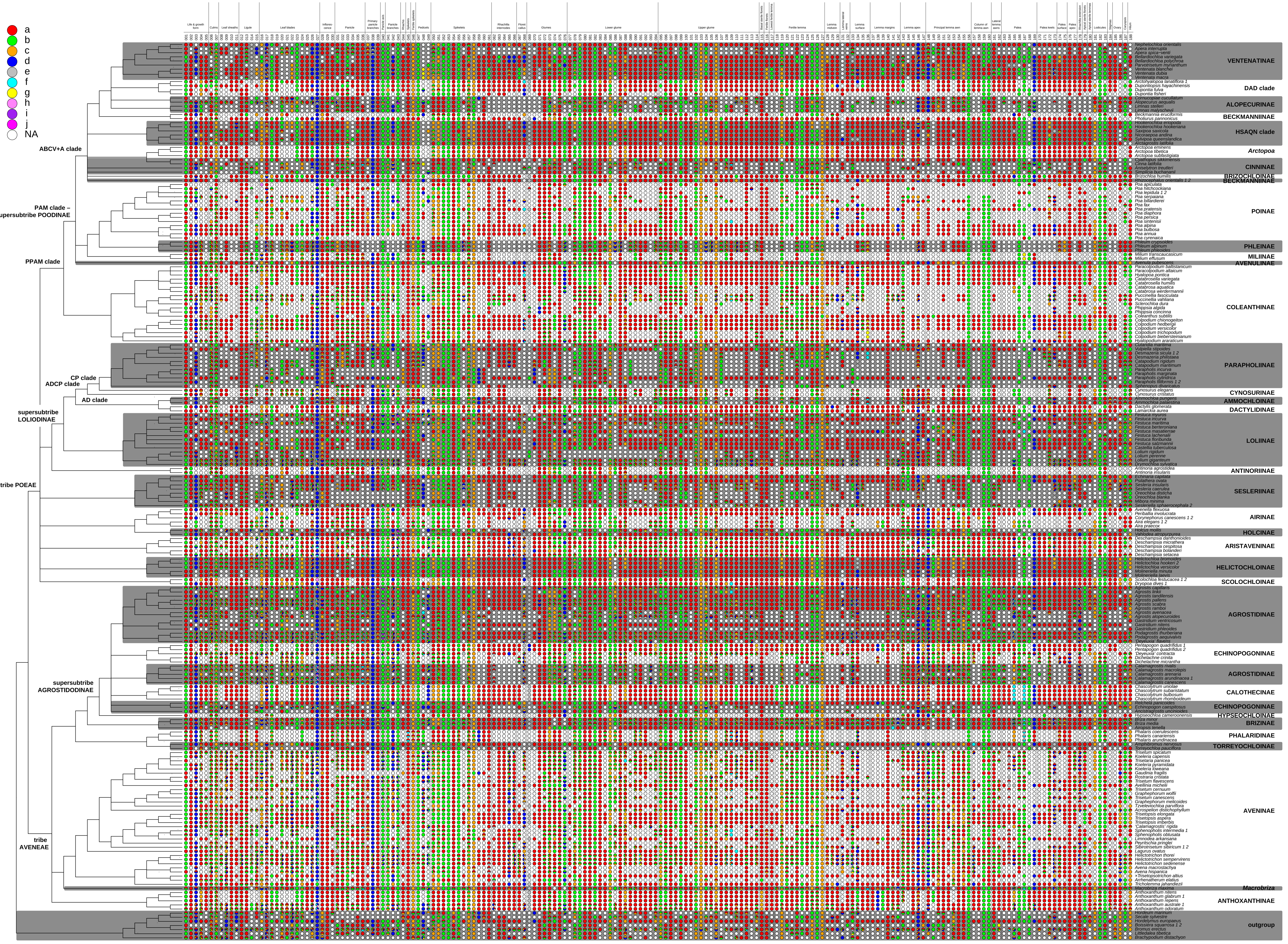
