## Supplementary Appendix S2 for "Phylogeny, morphology and the role of hybridization as driving force of evolution in grass tribes Aveneae and Poeae (Poaceae)"

**Supplementary Appendix S2.** 188 mainly morphological characters scored for the 218 taxa studied, character states mapped on the tips of the molecular phylogenetic tree (suppl. Fig. S1) and used for the ancestral state reconstructions (ASR) in suppl. Appendix S3. The characters are arranged according to the different parts of the plants (plant organs) or types of characters and listed with character number and character states (a–j).

**Life and growth form:** 001, Plant duration, a: annual, b: perennial. 002, Degree of perennation, a: persisting, b: short-lived <perennating organs weakly expressed and difficult to observe>. 003, Plant density, a: culms solitary, b: cushion forming, c: mat forming, d: <uni> caespitose <forming a single clump>. 004, Rhizomes <presence>, a: absent, b: short <often obliquely ascending>, c: elongated <running>. 005, Stolons <presence>, a: absent, b: present. **Culms:** 006, <habit>, a: erect, b: geniculately ascending, c: decumbent <touching ground at knees>, d: prostrate <lying on ground>, e: rambling <on low vegetation>, 007, <height in cm>, a: 0–10 cm, b: 11–40 cm, c: > 40 cm. **Leaf sheaths:** 008, <roughness>, a: smooth, b: scaberulous, c: antrorsely scabrous, d: retrorsely, scabrous, e: papillose, g: transversely wrinkled. 009, <surface indumentum>, a: glabrous on surface <excepting mouth & margin>, b: puberulous, c: pubescent, d: pilose, e: hirsute, f: hispid, g: woolly. 010, with <hair type>, a: simple hairs, b: tubercle-based hairs, c: reflexed hairs. 011, <presence of auricles>, a: absent, b: erect <usually adnate to ligule>, c: erect and connate forming a tooth opposite blade <when sheath tubular>, d: falcate. **Ligule:** 012, <structure - at midculm node when cauline foliage present>, a: an eciliate membrane <apical hairs absent>, b: a ciliolate membrane <apical hairs shorter than membrane>, c: a ciliate membrane <apical hairs as long as, or longer than, membrane>, d: a fringe of hairs <membrane absent or obscure>, e: absent. 013, Ligule <length on culm leaves in mm>, a: 0–5, b: 6–10, c: >10. 014, <apex incision>, a: entire, b: erose, c: bilobed, d: trilobed, e: lacerate. 015, <shape of apex> a: truncate, b: obtuse, c: acute, d: acuminate. **Leaf blades:** 016, <outline>, a: aciculate, b: filiform, c: linear, d: lanceolate, e: elliptic, h: triangular. 017, <vernation>, a: flat, b: plicate <pleated>, c: conduplicate <folded>, d: involute, e: convolute. 018, <stiffness>, a: stiff, b: firm, c: flaccid. 019, <presence of scent>, a: without scent, b: aromatic. 020, surface <roughness>, a: smooth, b: scaberulous, c: scabrous, d: papillose. 021, surface <roughness on which side>, a: adaxially <upper>, b: abaxially <lower>, c: on both sides. 022, surface < indumentum>, a: glabrous, b: puberulous, c: pubescent, d: pilose, e: hirsute, f: hispid. 023, surface <hair density>, a: sparsely hairy, b: moderately hairy, c: densely hairy. 024, margins <roughness>, a: smooth, b: scaberulous, c: scabrous. 025, margins < hairy where>, a: all along, b: at base <only>. 026, apex <shape>, a: obtuse, b: abruptly acute, c: acute, d: acuminate, e: attenuate. 027, apex <termination>, b: hooded, d: simple, e: apiculate, f: hardened, g: callose, h: filiform. **Inflorescence:** 028, <type>, a: a panicle <including spiciform panicles with dwarf laterals more or less accrescent to a central axis>, b: a panicle with branches tipped by a raceme, c: composed of <spikes or> racemes, d: comprising only a few spikelets <and type undefinable>. 029, <whether overtopping basal leaves>, a: aerial, b: shorter than basal leaves. 030, subtended by <type of subtending leaf>, a: an unspecialized leaf-sheath <and blade>, b: an inflated leaf-sheath <lightly inflated, with blade>, c: a spatheole <or spathe, much modified with rudimentary blade>. **Panicle:** 031, <type>, a: open, b: contracted, c: spiciform, d: glomerate, e: capitate. 032, Panicle <one-sided>, a: equilateral, b: nodding <to one side>, c: second. 033, <length in cm>, a: 0–5, b: 6–20, c: >20. 034, <width in cm>, a: 0–5, b: >5. 035, bearing <sparse>, a: many spikelets, b: few spikelets <few enough to count>. 036, <spikelet distribution>, a: evenly furnished <with spikelets>, b: contracted about primary branches, c: contracted about secondary branches, e: gathered into fascicles, f: with spikelets clustered towards branch tips. **Primary panicle branches:** 037, <whorled>, a: not whorled, b: whorled at lower nodes <only>, c: whorled at most nodes. 038, <amount of branching>, a: indistinct the panicle almost racemose, b: simple, c: sparsely divided, d: moderately divided, e:

profusely divided. 039, branching <type of branching>, a: laterally, b: dichotomously <dividing into equal parts>, c: divaricately <widely divergent>. **Panicle axis:** 040, <indumentum>, a: with scattered hairs, b: glabrous, c: puberulous, d: pubescent, e: pilose, f: hirsute, g: villous, i: hispidulous. **Panicle branches:** 041, <stiffness>, a: stiff, b: flexible, c: capillary. 042, roughness>, a: smooth, b: with occasional prickles, c: scaberulous, d: antrorsely scabrous, e: retrorsely scabrous. 043, <surface indumentum>, a: with scattered hairs, b: glabrous, c: puberulous, d: pubescent, e: pilose, f: hirsute, g: villous, i: hispidulous, j: hispid. **Rhachis:** 044, <fragility>, a: tough <or obsolete>, c: fragile at the nodes. **Spikelets:** 045, <kinds of cluster. The clusters may include sterile and monoecious male spikelets. A deciduous inflorescence does not count as a single cluster, but may contain clusters.>, a: solitary, b: in pairs <may include barren pedicels>, c: in threes, f: subtended by an involucre <may be of only a few bristles or sterile spikelets; not counting basal sterile spikelets>. **Fertile spikelets:** 046, <presence of pedicels, which may be connate>, a: sessile <or subsessile with pedicel represented by a brief stump>, b: sessile and pedicelled, c: pedicelled. **Pedicels:** 047, <presence>, a: absent <or square and inconspicuous>, b: present <or square and ornamented>. 048, <shape>, a: filiform, b: linear, d: oblong, f: clavate <linear below, widening above>, g: cuneate. 049, <indumentum>, a: bearing a few <long> hairs, b: glabrous, c: puberulous, d: pubescent, e: ciliate <or pilose>, f: villous. **Spikelets:** 050, comprising <number of fertile florets>, a: 1, b: 2–3, c: >3. 051, <presence of apical florets>, a: without rhachilla extension <beyond uppermost fertile floret>, b: with a barren rhachilla extension, c: with diminished <sterile or smaller> florets at the apex. 052, <key couplet for 1-flowered groups>, a: of 1 fertile floret with or without additional sterile florets, b: of 2 or more fertile florets. 053, a: two-flowered - the lower floret male or barren, the upper fertile <rarely with rhachilla extension or rudiment>, b: one–many-flowered - if two-flowered then both fertile or the upper sterile. 054, <compression of fertile>, a: laterally compressed, b: subterete, c: dorsally compressed. 055, compressed <how much>, a: slightly <spikelets plump>, b: moderately, c: strongly. 056, <shape of apex>, a: truncate, b: obtuse, c: subacute, d: acute, e: acuminate. 057, <length in mm>, a: 0–3, b: 4–6, c: >6. 058, <abscission>, a: persistent on plant <cereals>, b: falling entire, c: breaking up at maturity. 059, deciduous <site of abscission - if falling entire>, a: from the base, b: with the <whole> pedicel <but no other organ; excluding partial attachment (some *Polypogon*) which is treated as a callus.>, c: in a cluster with fused pedicels <and sometimes a basal extension, but no other organ>, d: with accessory branch structures. 060, <secondary abscission - if falling entire>, a: without secondary abscission, c: readily shedding fertile florets. 061, disarticulating <where - if rhachilla deciduous>, a: below each fertile floret <or below sterile if attached>, b: between fertile florets but the lowest falling with glumes attached, e: above glumes but not between florets <when 2 or more fertile>. **Rhachilla internodes:** 062, <elongation between glumes and lowest fertile floret>, a: brief up to lowest fertile floret, b: elongated between glumes, d: elongated between basal sterile florets <when more than 1>, e: elongated below proximal fertile floret. 063, <thickening>, a: narrow, b: clavate. 064, <alignment>, a: straight, b: curved, c: zig-zag. 065, <indumentum>, a: glabrous, b: sparsely hairy, c: pubescent, d: pilose, e: villous. 066, hairy <extent>, a: all along, b: all along but hairs longer above, c: at tip <only>, d: above, e: below. **Floret callus:** 067, <presence>, a: brief, b: evident <not much longer than wide>, c: elongated. 068, <indumentum>, a: glabrous, b: sparsely hairy, c: pubescent, d: pilose, e: bearded, f: woolly. **Glumes:** 069, <presence>, a: both absent or obscure, b: one the lower absent or obscure, d: one to two the lower present in some spikelets <often the sessile of a pair>, e: two. 070, <whether lateral>, a: distichous, b: lateral <opposite, but rotated with respect to lemma>, c: oblique <not strictly opposite>, d: collateral <side by side>. 071, <persistence - if spikelets break up>, a: persistent <on branch>, d: <both> deciduous, e: deciduous <together> with pedicel attached. 072, <similarity, apart from some difference in size>, a: similar <in shape, texture, etc>, b: dissimilar. 073, <length of longer

relative to rest of spikelet, excluding awns>, a: shorter than spikelet, b: reaching apex of florets, c: exceeding apex of florets. 074, <consistency relative to fertile lemma>, a: thinner than fertile lemma, b: similar to fertile lemma in texture, c: firmer than fertile lemma. 075, <sheen>, a: dull, b: shiny. 076, <divergence>, b: parallel to lemmas, c: recurved at apex, d: gaping. **Lower glume:** 077, <gibbosity>, a: not gibbous, b: gibbous, c: saccate. 078, <length in mm>, a: 0–2, b: 3–6, c: >6. 079, <length as fraction of upper>, length of upper glume, a: 0–0.5, b: 0.6–1, c: >1. 080, <apical consistency>, a: of similar consistency above, b: much thinner above. 081, <marginal consistency> a: of similar consistency on margins, b: much thinner on margins, c: firmer on margins. 082, <presence of keels>, a: without keels, b: 1-keeled, c: 2-keeled <or inflexed>. 083, keeled <extent>, a: all along, b: above, c: below. 084, <presence of wings>, a: wingless, b: winged on keel. 085, <vein number>/-veined, a: 0, b: 1, c: ≥3. 086, primary vein < midvein or keels - roughness. Knobs and spines on 2-keeled glumes are treated as surface (flank) roughness>, a: smooth, b: scaberulous, c: scabrous, d: spinulose. 087, primary vein <presence of hairs (except tufts) or spines>, a: eciliate, b: ciliolate, c: ciliate, d: pectinately ciliate <stiff and evenly spaced>. 088, lateral veins <clarity>, a: absent, b: obscure, c: distinct, d: prominent, f: unequally thickened. 089, lateral veins <presence of ribs>, a: without ribs, b: ribbed. 090, surface <relief>, a: smooth, b: asperulous, c: scabrous. 091, surface rough <where, excluding keels>, a: generally, b: at apex, c: above, e: below, f: on flanks <only>, g: on veins. 092, surface hairy <where> a: generally, b: at apex, c: above, e: below, g: on veins. 093, apex <incision>, a: entire, b: erose, d: dentate, e: lobed. 094, apex <presence of awn>, a: muticous, b: mucronate, c: awned. **Upper glume:** 095, <length in mm>, a: 0–2, b: 3–6, c: >6. 096, <length as fraction of lemma>/ length of adjacent fertile lemma, a: 0–1, b: >1. 097, <apical consistency>, a: of similar consistency above, b: much thinner above. 098, with <marginal consistency>, a: undifferentiated margins, b: hyaline margins, c: membranous margins, e: scarious margins. 099, <keels>, a: without keels, b: 1-keeled. 100, keeled <extent>, a: all along, b: above, c: below. 101, <presence of wings>, a: wingless, b: winged on keel. 102, <vein number>/-veined/, a: 1, b: 3, c: ≥5. 103, primary vein <clarity of mid or keel vein>, a: absent, b: obscure, c: distinct, d: conspicuous. 104, primary vein <roughness>, a: smooth, b: scaberulous, c: scabrous, d: spinulose. 105, primary vein <presence of hairs (except tufts) or spines>, a: eciliate, b: ciliolate, c: ciliate, d: pectinately ciliate <stiff and evenly spaced>. 106, lateral veins <clarity>, a: absent, b: obscure, c: distinct, d: prominent, e: <equally> thickened. 107, lateral veins <presence of ribs>, a: without ribs, b: ribbed. 108, surface <roughness>, a: smooth, b: asperulous, c: scabrous, f: papillose. 109, surface rough <where>, a: generally, b: at apex, c: above, e: below, f: on veins. 110, surface <general indumentum>, a: glabrous <except for keel, margins & special indumentum>, b: puberulous, c: pubescent, d: pilose, e: hirsute, f: villous, g: hispidulous, h: hispid. 111, surface hairy <where>, a: generally, b: at apex, c: above, e: below, g: on veins. 112, margins <hairiness>, a: eciliate, b: ciliolate, c: ciliate, f: pubescent. 113, apex <incision>, a: entire, b: erose, d: dentate, e: lobed. 114, <presence of awns>, a: muticous, b: mucronate, c: awned. **Basal sterile florets:** 115, <presence, including vestiges>, a: absent, b: 1, c: 2 or more. **Fertile florets:** 116, <similarity if more than 1>, a: all alike, c: with the lowest dissimilar. **Lowest fertile lemma:** 117, <relative consistency>, a: of similar consistency to adjacent lemma, b: thinner than adjacent lemma. **Fertile lemma:** 118, <presence of basal auricles or thickening>, a: without auricles, b: auriculate at base, c: thickened on margins at base. 119, <length in mm>, a: 0–1.5, b: 1.6–4, c: >4. 120, <apical consistency>, a: of similar consistency above, b: much thinner above. 121, <marginal consistency>, a: of similar consistency on margins, b: much thinner on margins. 122, <presence of keel>, a: without keel, b: keeled. 123, <clarity of keel>, a: lightly keeled, b: distinctly keeled. 124, keeled <extent>, a: all along, b: above, c: below. 125, <vein number>/-veined, a: 1–3, b: 4–5, c: >5. 126, a: 0–3-veined, b: more than 3-veined. 127, <presence of veins>, a: without veins, b: one-veined, c: several-veined. **Lemma midvein:** 128, <roughness,

especially when keeled>, a: without distinctive roughness, b: scaberulous, c: scabrous. 129, <hairiness - except tufts>, a: eciliate <not distinct from general indumentum>, b: ciliolate, c: ciliate, d: pectinately ciliate <stiff and evenly spaced>, g: pubescent. 130, hairy <where>, a: all along, d: below. **Lemma lateral veins:** 131, <clarity>, a: obscure, b: with distinct primaries but obscure intermediates, c: distinct, d: prominent. 132, <presence of ribs>, a: without ribs, b: ribbed. **Lemma surface:** 133, rough <where>, a: generally, b: above, c: in the middle, d: below, e: on veins, g: in lines <unrelated to veins>. 134, hairy <extent of general indumentum> a: all along, b: above <excluding apical ornament>, c: in the middle, d: below, e: at base. 135, hairy <where>, a: on back, b: on veins <recorded under margins if veins marginal>, c: between veins. 136, with <hair type>, a: simple hairs, b: tubercle-based hairs, f: capitate hairs, g: clavate hairs. **Lemma margins:** 137, <fusion>, a: free, b: connate below. 138, <vernation - if free>, a: flat, c: involute, d: convolute. 139, <coverage of palea>, a: exposing palea, b: covering most of palea, c: interlocking with palea keels. 140, <roughness>, a: without distinctive roughness, b: scaberulous, c: scabrous. 141, <hairiness>, a: eciliate <not distinct from general indumentum>, b: ciliolate, c: ciliate, e: pubescent, f: pilose. 142, hairy <extent of hairiness>, a: all along, b: above, d: below, e: at base. **Lemma apex:** 143, <incision>, a: entire, b: erose/dentate, c: lobed. 144, <number of primary teeth or lobes>/-fid, a: 2, b: ≥3. 145, <equality of lobes>, a: with simple equal lobes, b: with outer lobes longer, c: with outer lobes shorter, e: with lateral lobes bidentate, f: with irregular lobes. 146, <presence of awn>, a: muticous, b: pungent, c: mucronate <may be further described using awn characters>, d: awned. 147, <number of awns>/-awned, a: 1, b: 3, c: ≥5. **Principal lemma awn:** 148, < position>, a: apical, b: subapical, c: from a sinus, d: dorsal. 149, arising <origin of dorsal awn as fraction of lemma length>/ way up back of lemma, a: 0–0.5, b: 0.6–1. 150, <overall shape>, a: straight, b: curved, c: geniculate. 151, <flexion relative to lemma or column>, a: ascending, b: spreading, c: reflexed <through an angle>, d: recurved <with a semicircular bend> at base of limb, e: briefly coiled <or twisted> at base of limb. 152, <nature of tip>, a: attenuate, b: pungent, c: with scarcely tapering limb, d: with clavate limb. 153, <length including column in mm>, a: 0–3, b: 4–10, c: ≥11. 154, <exsertion>, a: clearly exserted from spikelet, b: not or scarcely exserted from spikelet. 155, <presence of column>, a: without a column, b: with a straight or slightly twisted column, c: with twisted column. 156, <coloured>, a: coloured, b: not coloured. **Column of lemma awn:** 157, <general indumentum>, a: glabrous <except for top>, b: hispidulous, c: puberulous, d: pubescent, f: hirtellous, g: hirsute. 158, <presence of apical hairs at junction with limb>, a: without distinct apical hairs, c: with bearded apex. 159, <flattened>, a: flattened, b: not flattened. 160, <twisted>, a: twisted, b: not twisted. **Lateral lemma awns:** 161, <presence>, a: absent, b: present. 162, arising <origin - if principal unbranched>, a: on <unlobed> apex, b: on apex of lobes, c: on inner edge of lobes, g: dorsally. **Palea:** 163, <presence>, a: present, b: absent or minute. 164, <gape>, a: embraced by lemma <tip may protrude apically>, e: gaping, f: readily splitting down midline. 165, <outline>, a: linear, b: lanceolate, c: elliptic, d: oblong, e: ovate, f: orbicular, g: obovate. 166, <fraction of lemma length>/ length of lemma, a: 0–0.6, b: ≥0.7. 167, <consistency>, a: hyaline, b: membranous, e: cartilaginous, f: coriaceous, g: indurate. 168, <number of veins, including keels>/-veined, a: 0–1, b: 2, c: ≥3. 169, <presence of keels>, a: without keels, b: 1-keeled, d: 2-keeled. **Palea keels:** 170, keels <spacing>, a: separated, b: approximate, c: contiguous above a sulcus. 171, <presence of wings>, a: wingless, b: winged. 172, <roughness>, a: smooth, b: scaberulous, c: scabrous, d: spinulose. 173, <indumentum>, a: eciliate, b: puberulous, c: pubescent, d: ciliolate, e: ciliate. **Palea surface:** 174, <indumentum - excluding keels and apex>, a: glabrous, b: puberulous, c: pubescent, d: pilose. 175, with <hair type>, a: simple hairs, c: turgid hairs. **Palea apex:** 176, <ornamentation>, a: undifferentiated, c: pubescent, d: ciliate, g: scaberulous. 177, <presence of awn>, a: muticous, b: with excurrent keel veins, c: awned. **Rhachilla extension:** 178, <indumentum>, a: glabrous, b: sparsely hairy, c: pubescent, d: pilose, e: villous. **Apical sterile florets:** 179,

<resemblance to fertile>, a: resembling fertile though underdeveloped, b: distinct from fertile <and variously modified>. **Apical sterile lemmas:** 180, <presence of awns, at least on distal floret>, a: muticous, b: mucronate, c: awned. **Lodicules:** 181, <number>, a: absent, b: 1, c: 2. **Anthers:** 182, <number>, a: 1–2, b: 3. 183, <length in mm>, a: 0–1, b: ≥1. **Stigmas:** 184, <indumentum>, a: plumose, b: sparsely hairy, c: pubescent. **Ovary:** 185, <presence of apical appendage>, a: unappendaged, b: with a fleshy appendage below style insertion, c: with a fleshy appendage above style insertion, e: beaked. 186, <indumentum>, a: glabrous, b: with a few apical hairs, c: pubescent on apex, d: pubescent all over. **Caryopsis:** 187, <length in mm>, a: 0–1.5, b: ≥1.6. **Hilum:** 188, <shape>, a: punctiform, b: elliptic, c: linear <and straight>.
