## Supplementary Appendix S3 for "Phylogeny, morphology and the role of hybridization as driving force of evolution in grass tribes Aveneae and Poeae (Poaceae)"

**Supplementary Appendix S3.** Character evolution and ancestral state reconstructions (ASR) for 74 characters in 218 taxa of Poodae (Aveneae and Poeae). Circles at the nodes are color-coded according to the tips to reflect ancestral character states (see key to codes in suppl. Appendix S2) with sizes of differently colored wedges indicating likelihood of presence of each state at that node (see text for further explanation). Differently colored wedges at the tips indicate the presence of different character states according to suppl. Fig. S2.

### (1) Plant duration

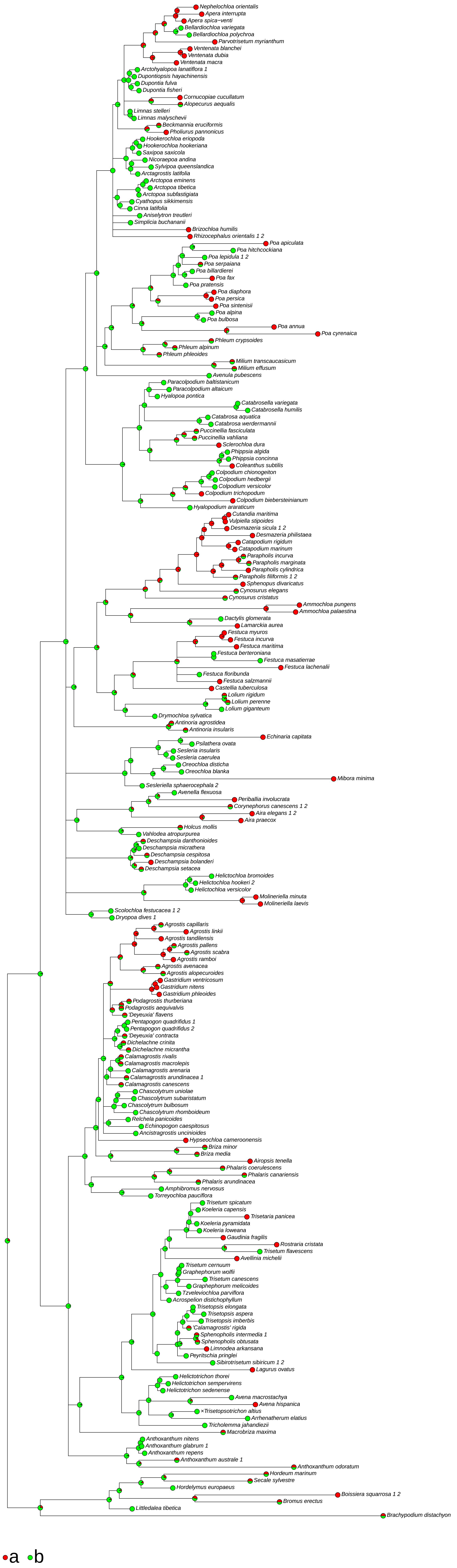

### (004) Rhizomes presence

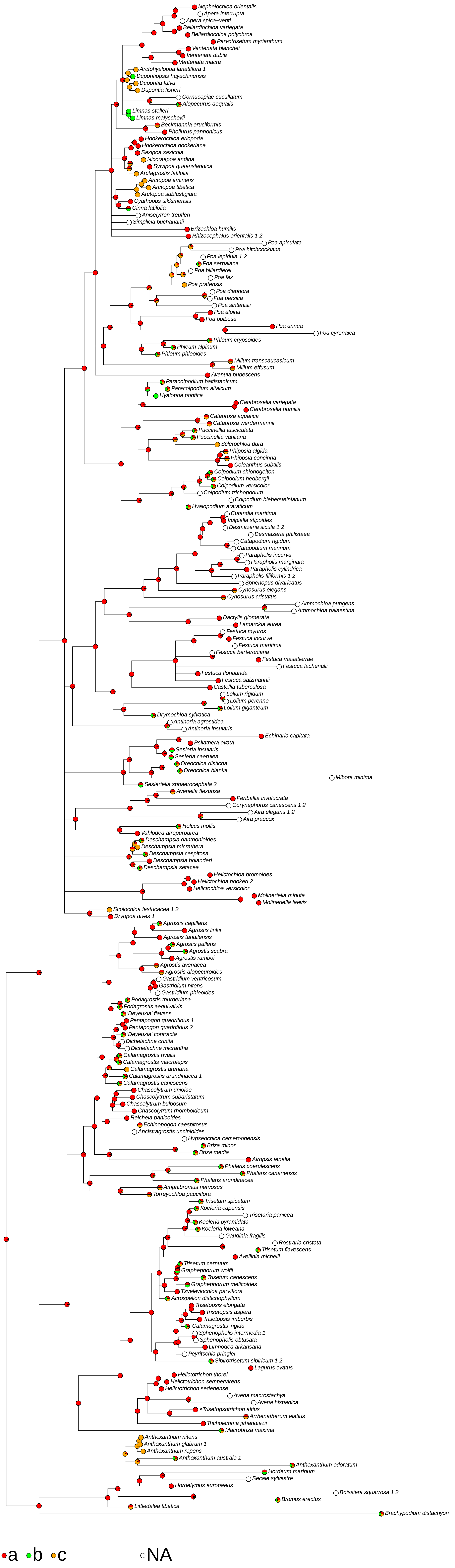

### (012) Ligule structure

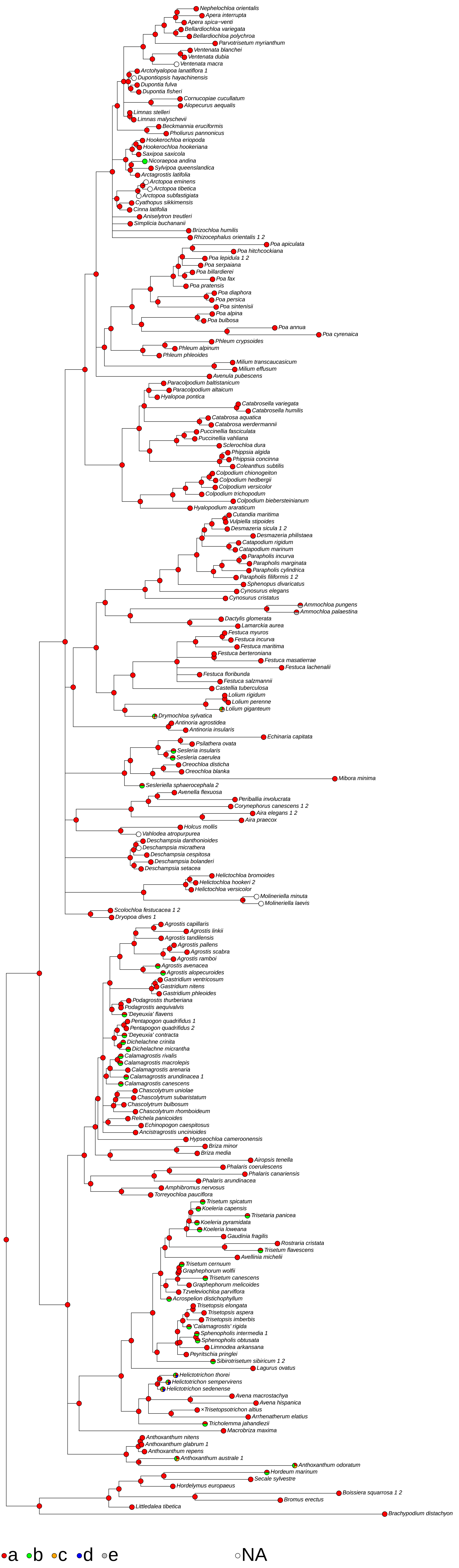

### (017) Leaf-blades <vernation>

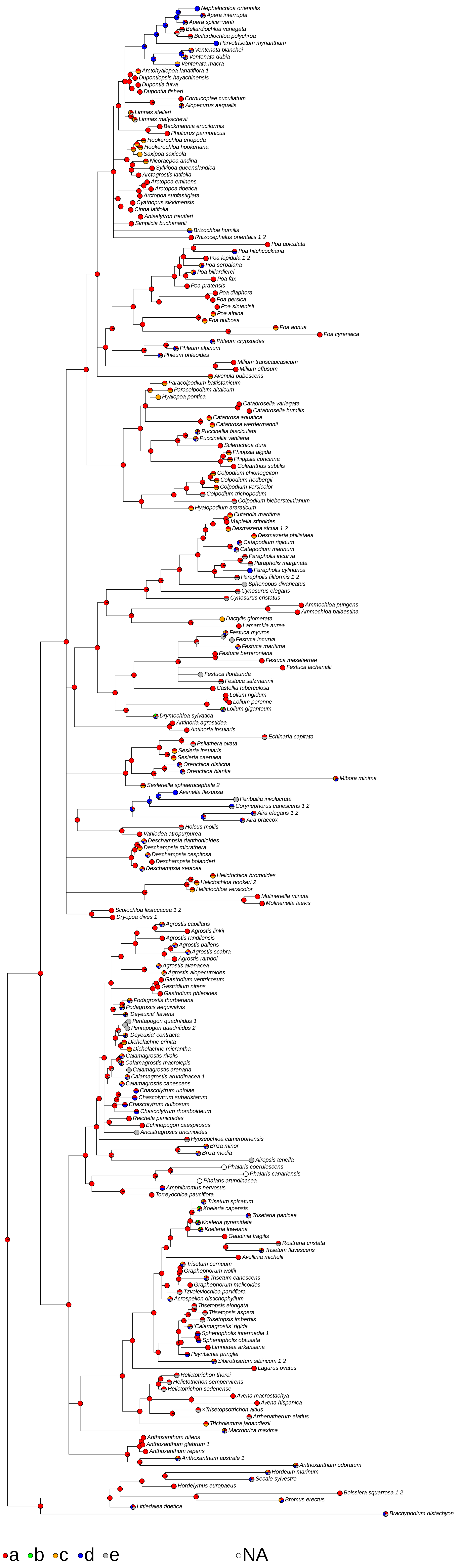

### (028) Inflorescence <type>

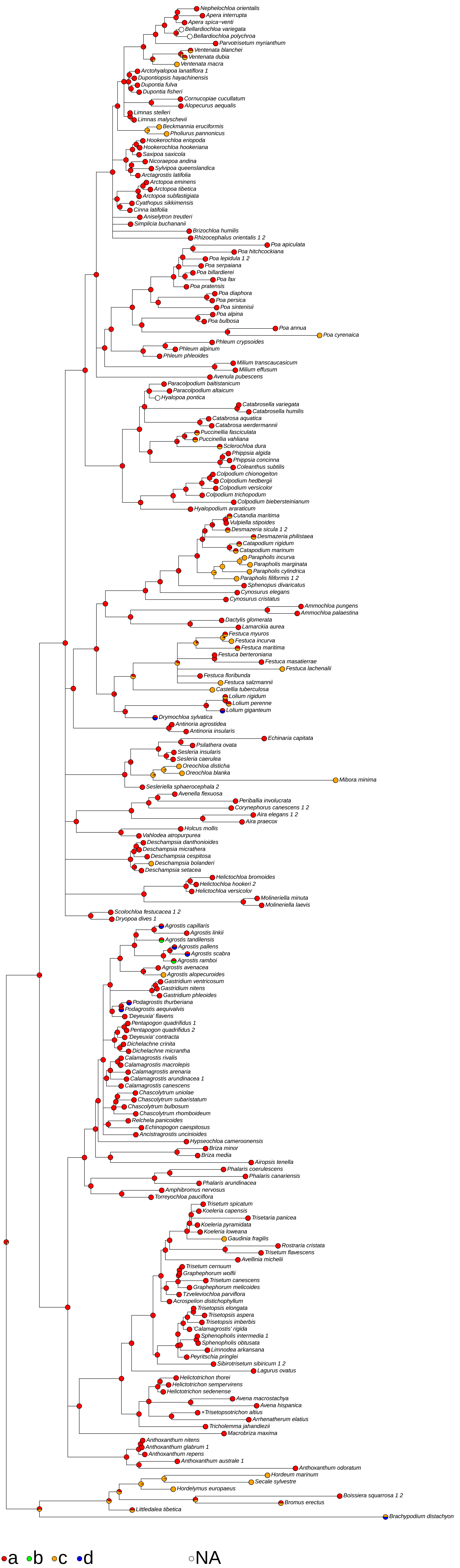

### (031) Panicle <type>

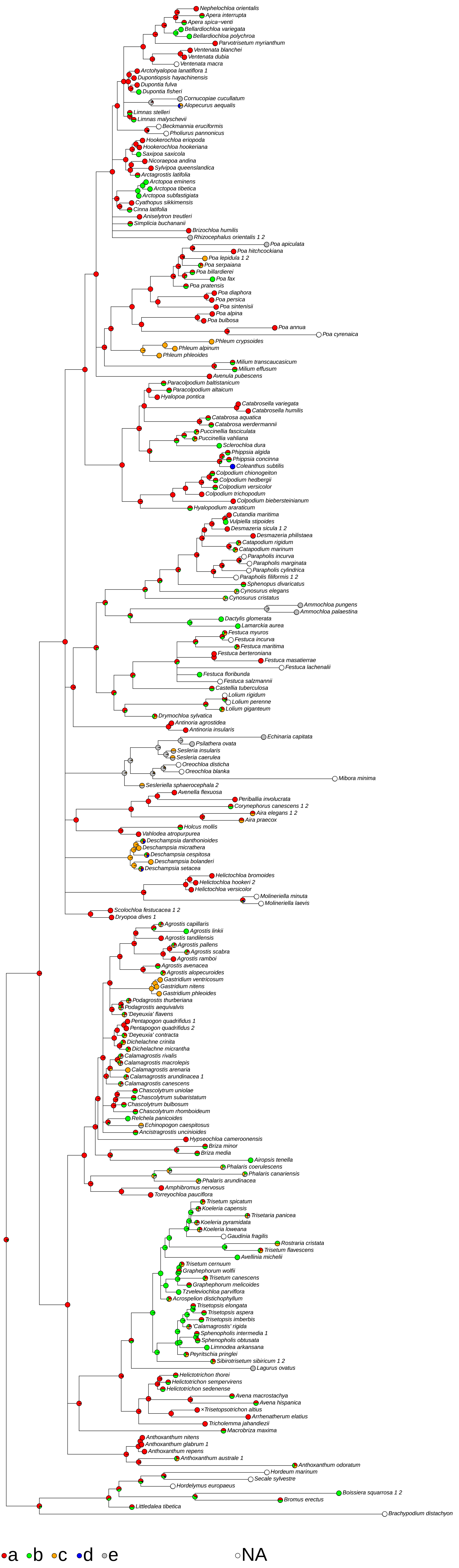

### (035) Panicle bearing sparse

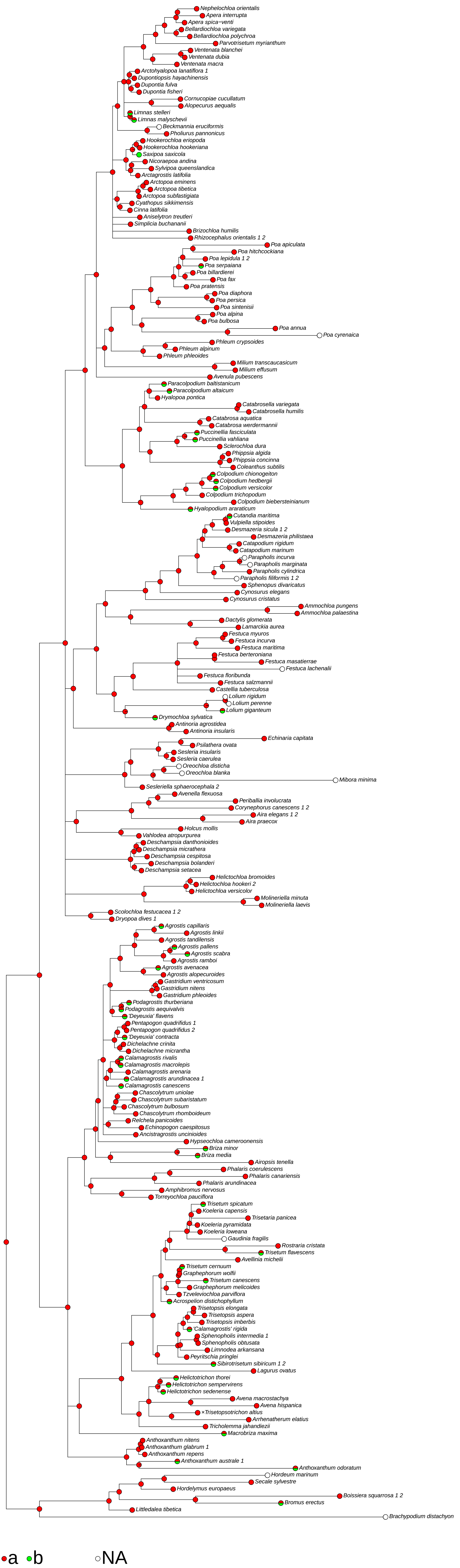

### (038) Primary panicle branches <amount of branching>

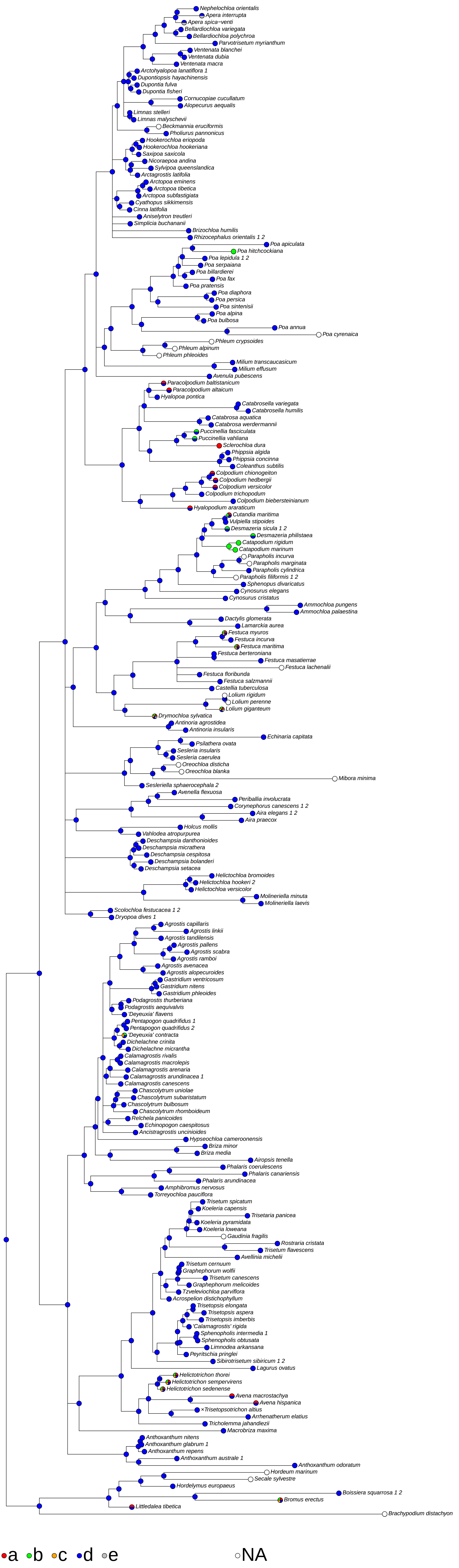

### (041) Panicle branches <stiffness>

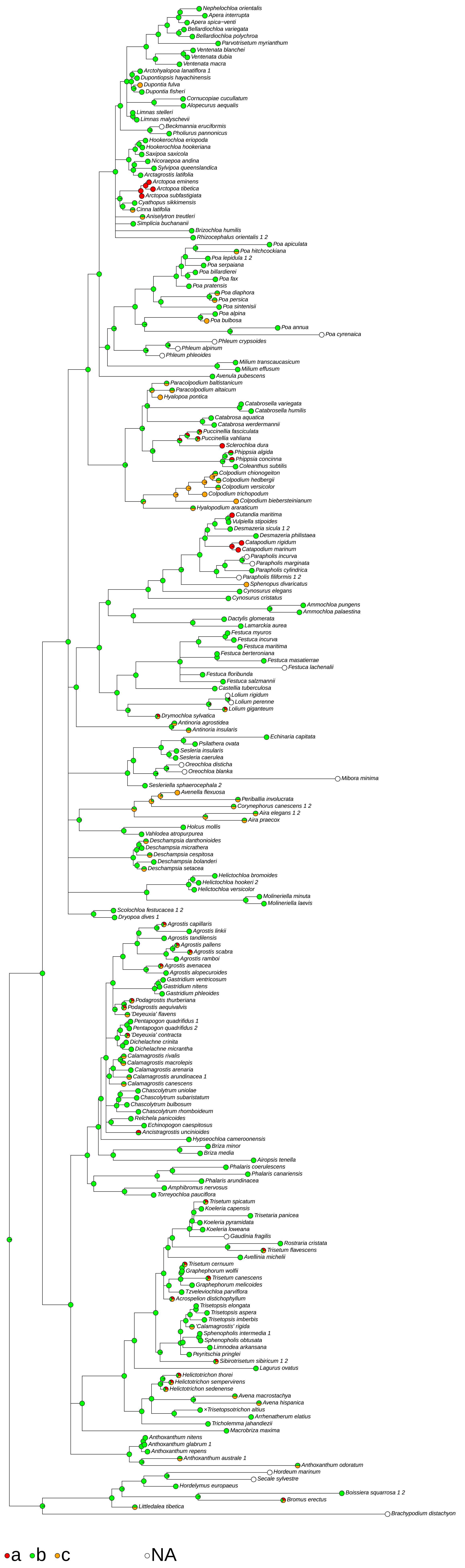

### (042) Panicle branches <roughness>

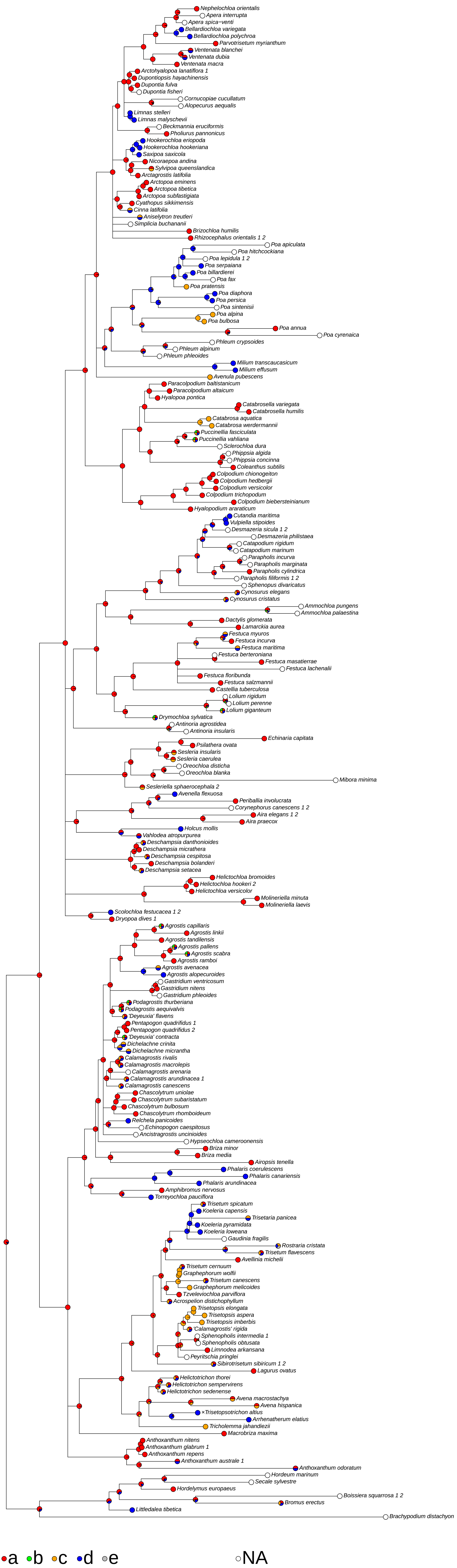

### (046) Fertile spikelets <presence of pedicels, which may be connate>

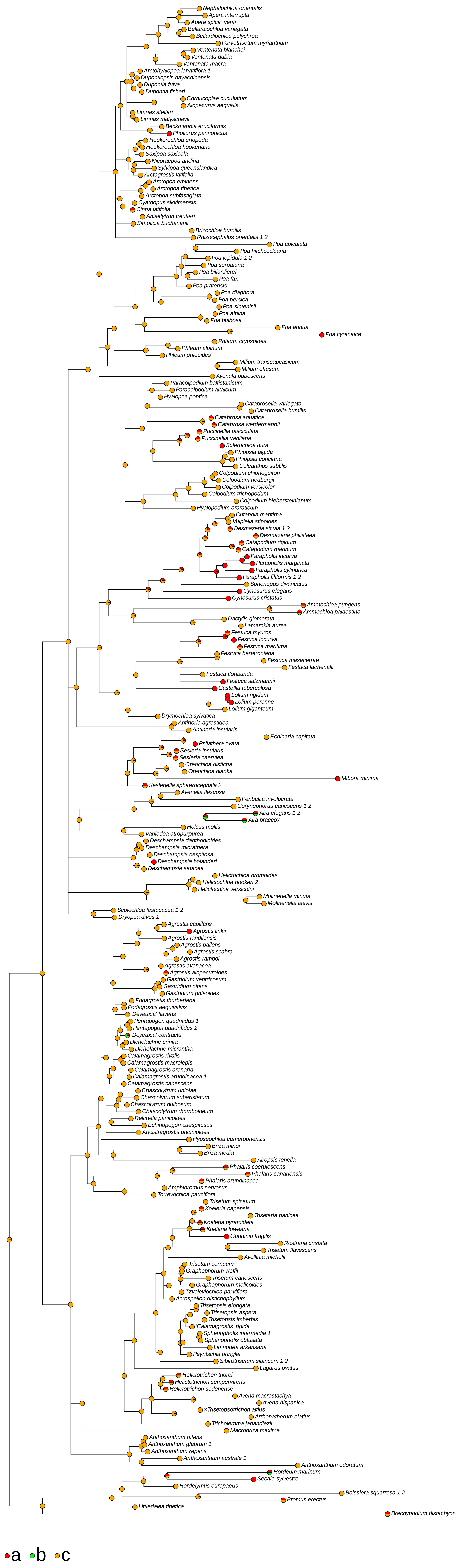

### (047) Pedicels <presence>

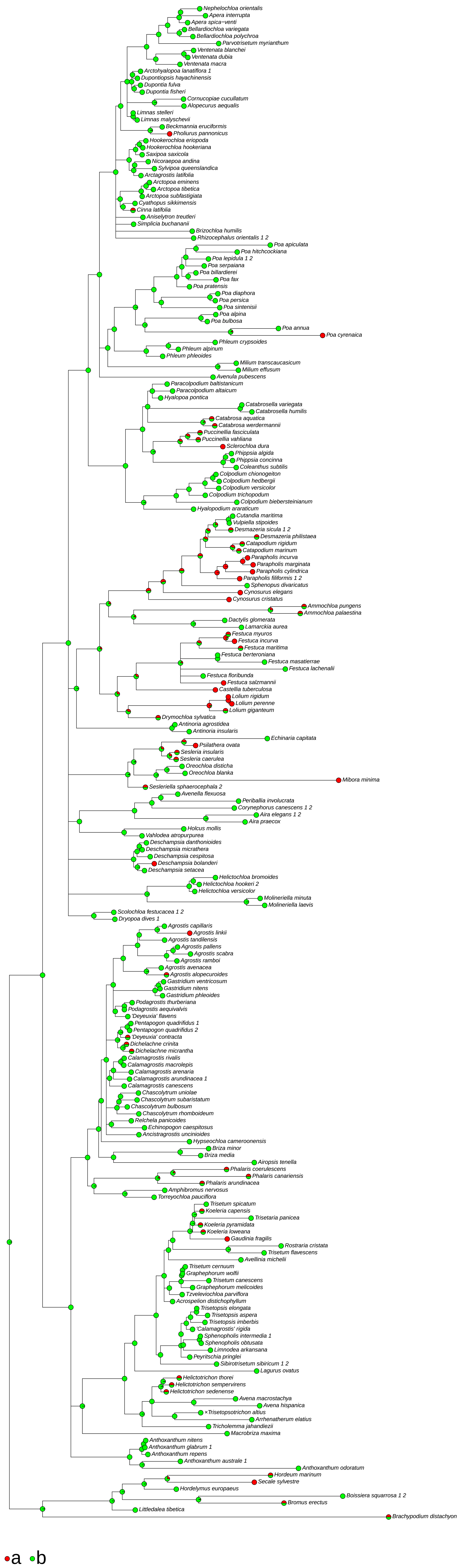

### (050) Spikelets comprising <number of fertile florets>

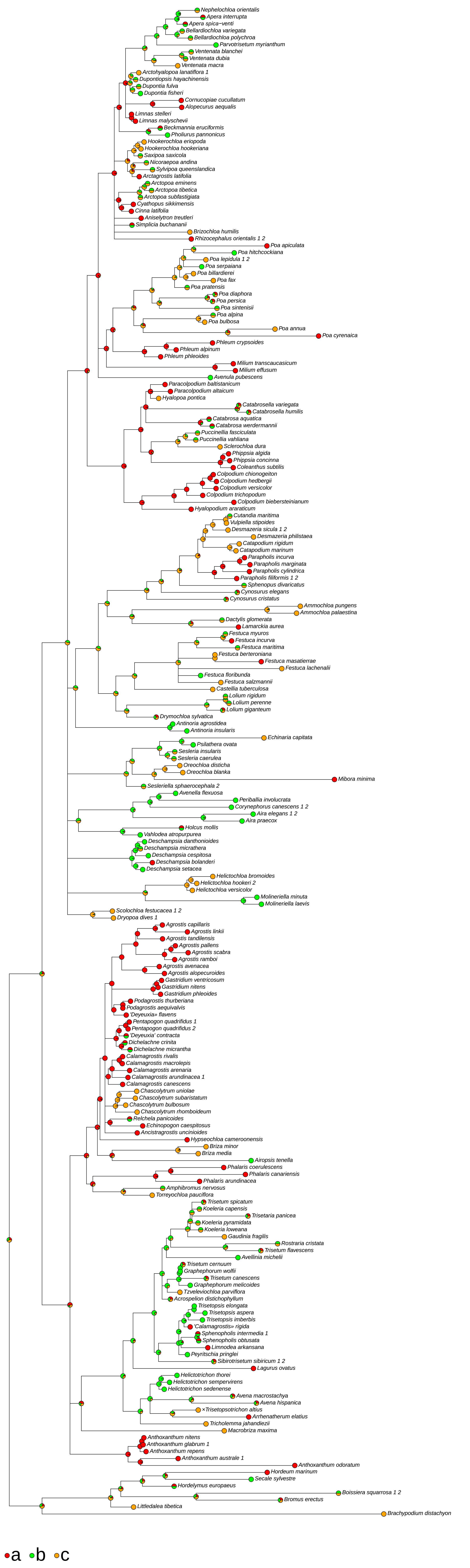

### (051) Spikelets <presence of apical florets>

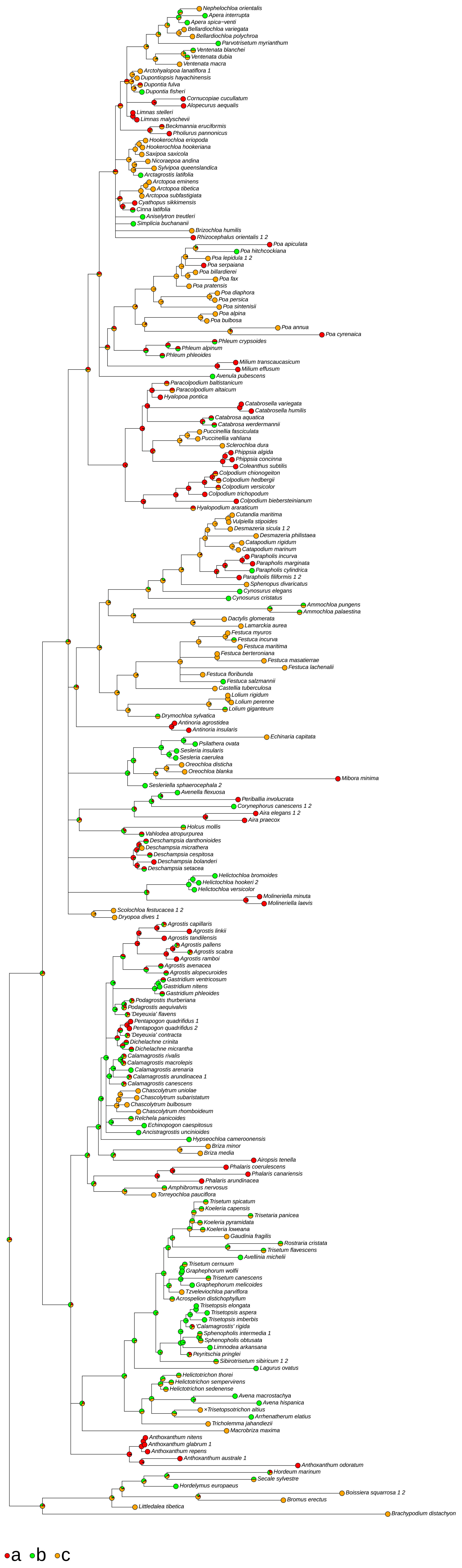

### (052) Spikelets <key couplet for 1-flowered groups>

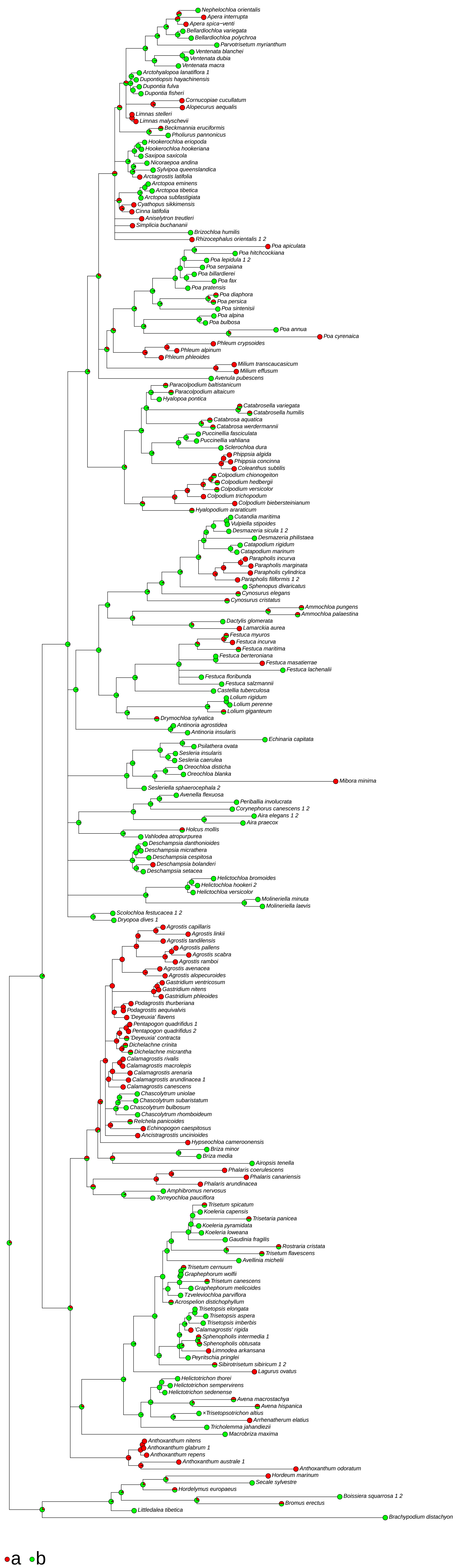

### (053) Spikelets

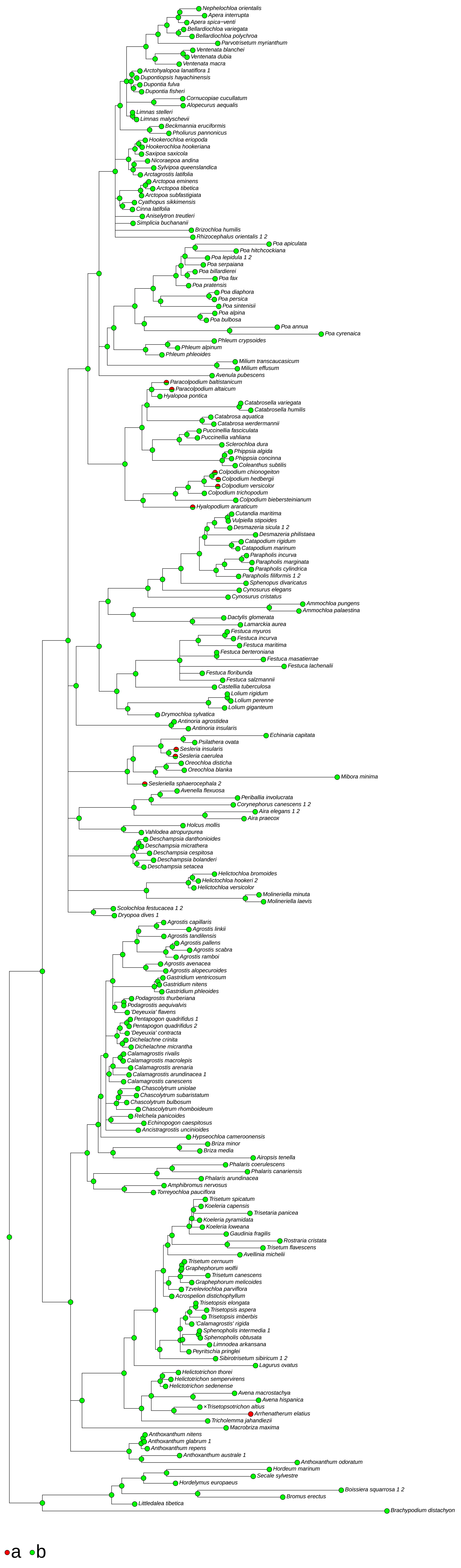

### (054) Spikelets <compression of fertile>

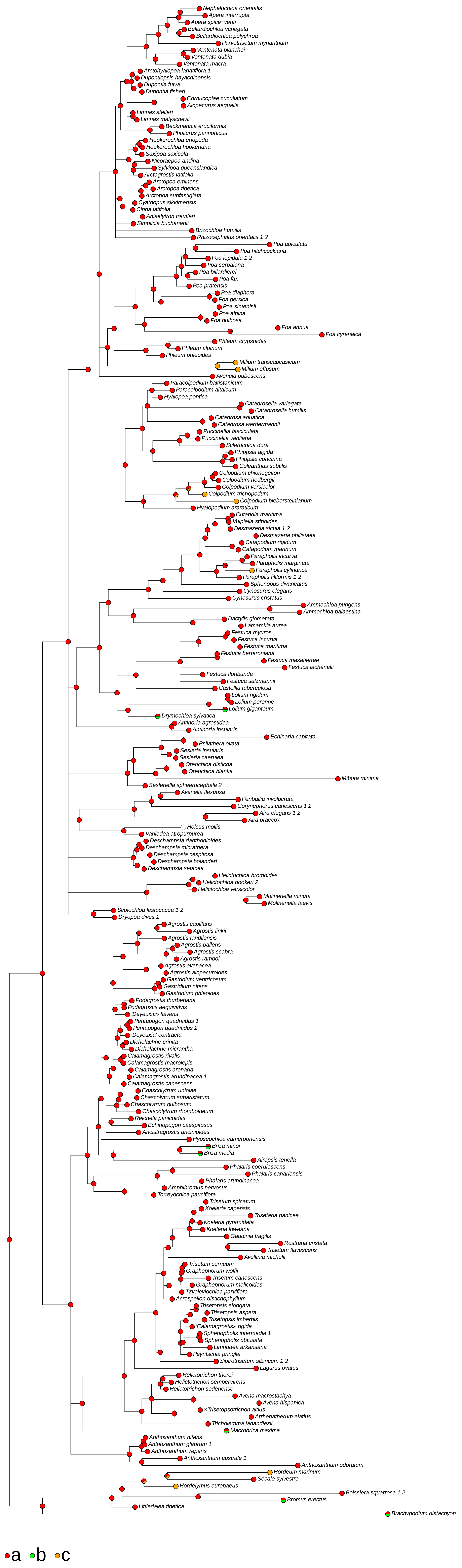

### (055) Spikelets compressed <how much>

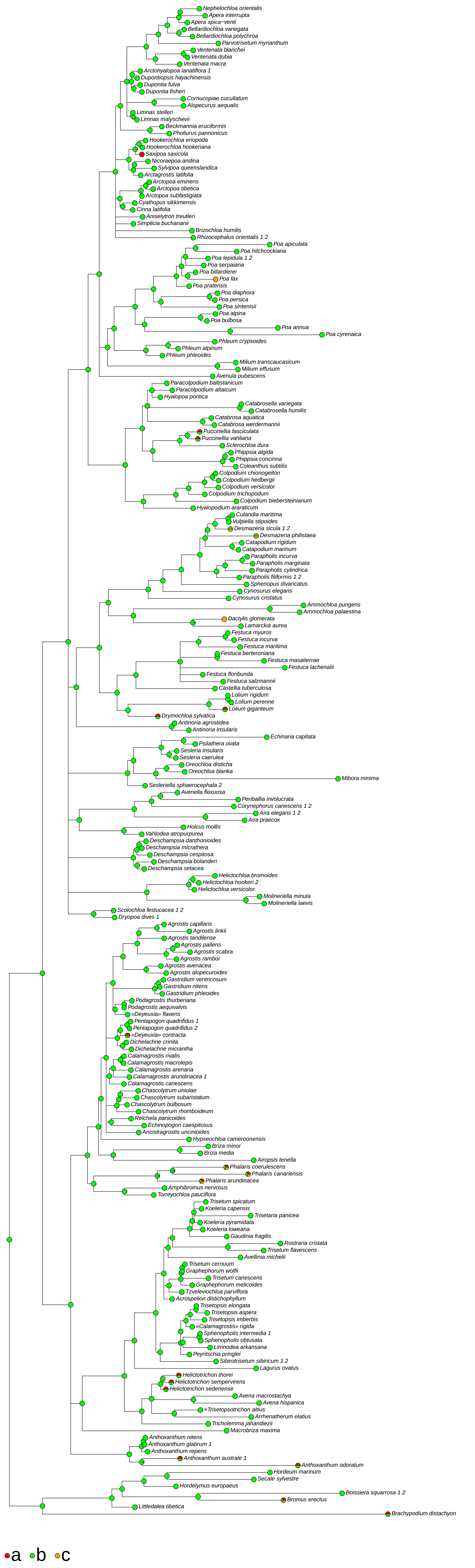

### (057) Spikelets <length in mm>

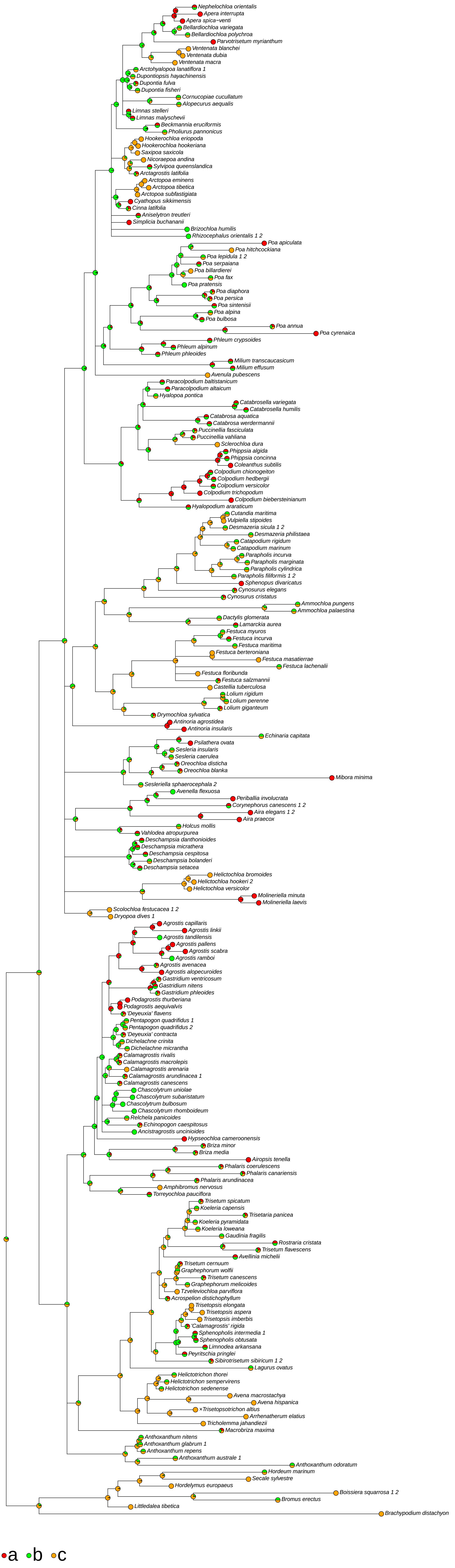

### (058) Spikelets <abscission>

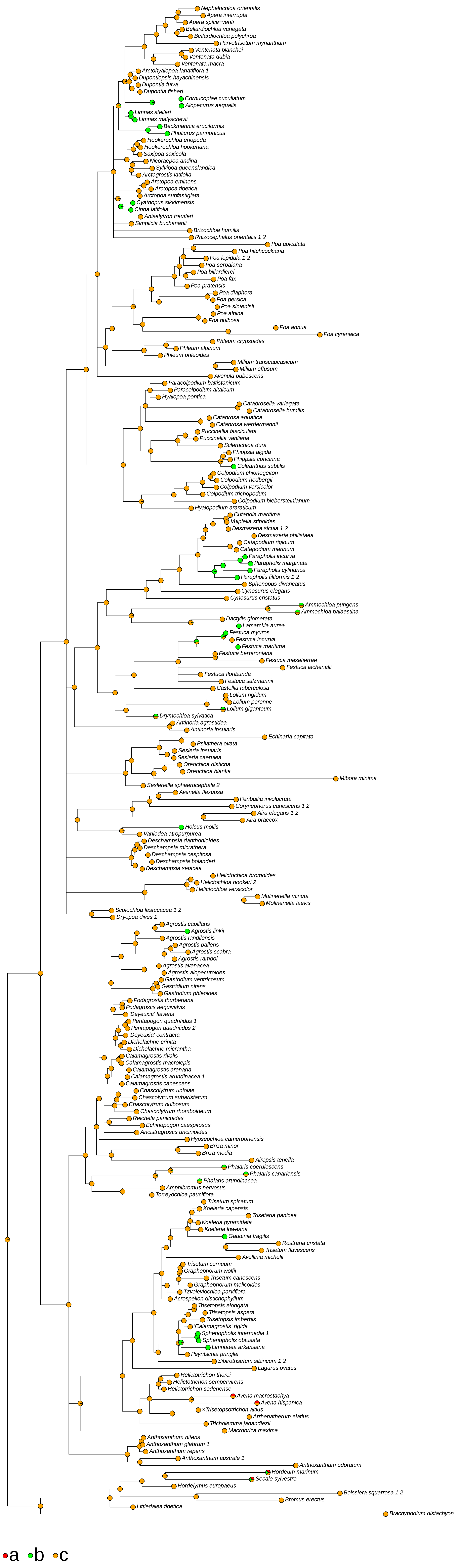

### (061) Spikelets disarticulating <where - if rhachilla deciduous>

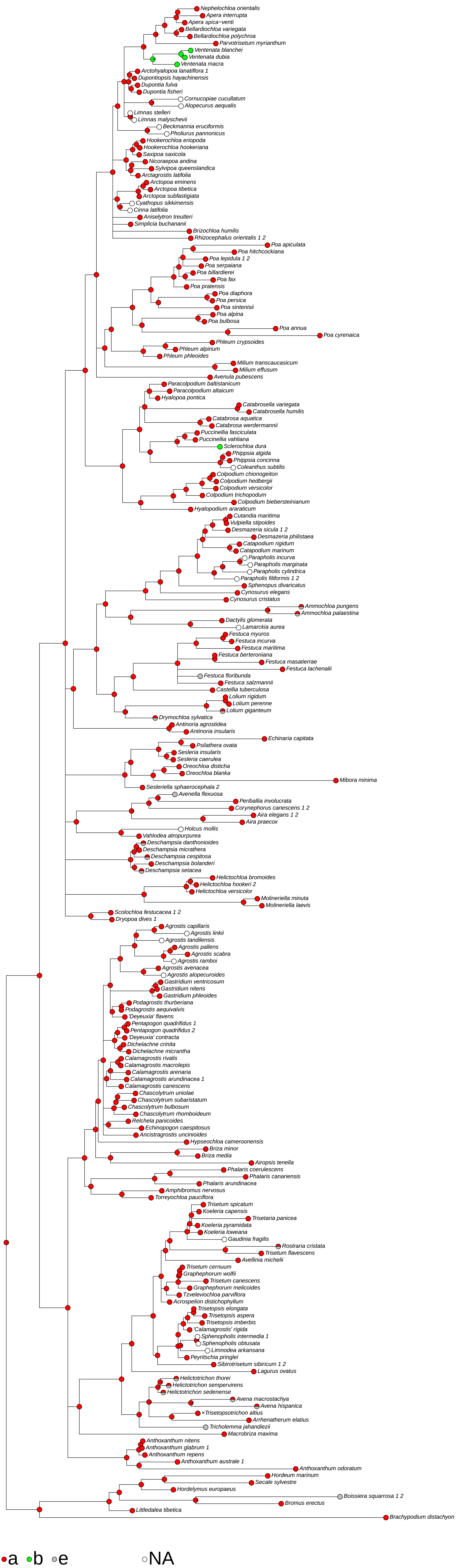

### (063) Rhachilla internodes <thickening>

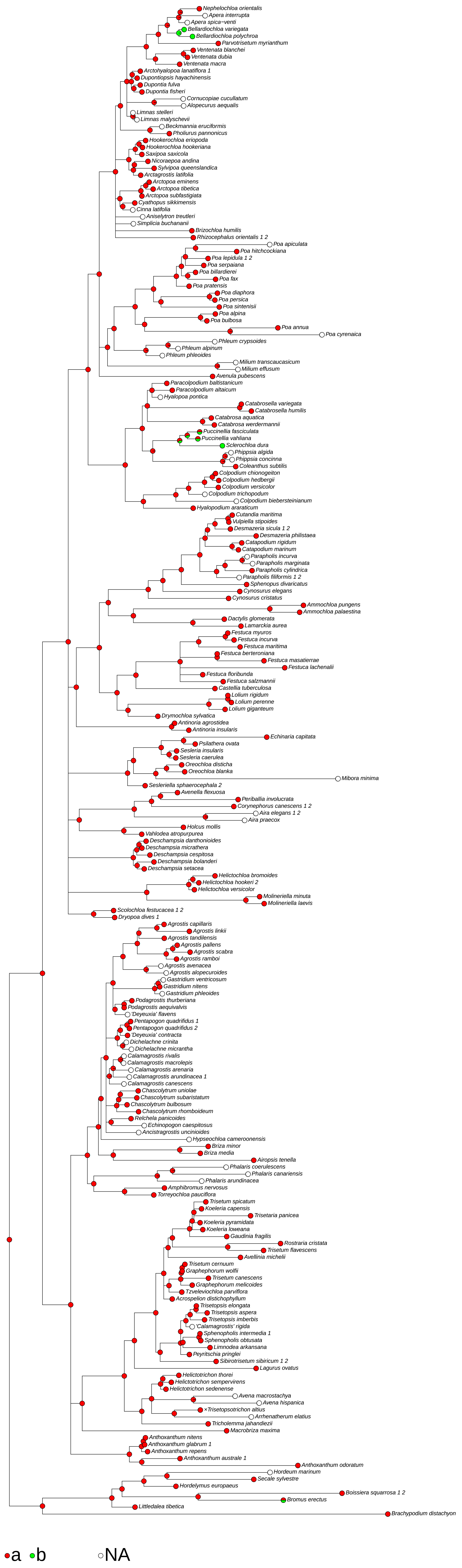

### (064) Rhachilla internodes <alignment>

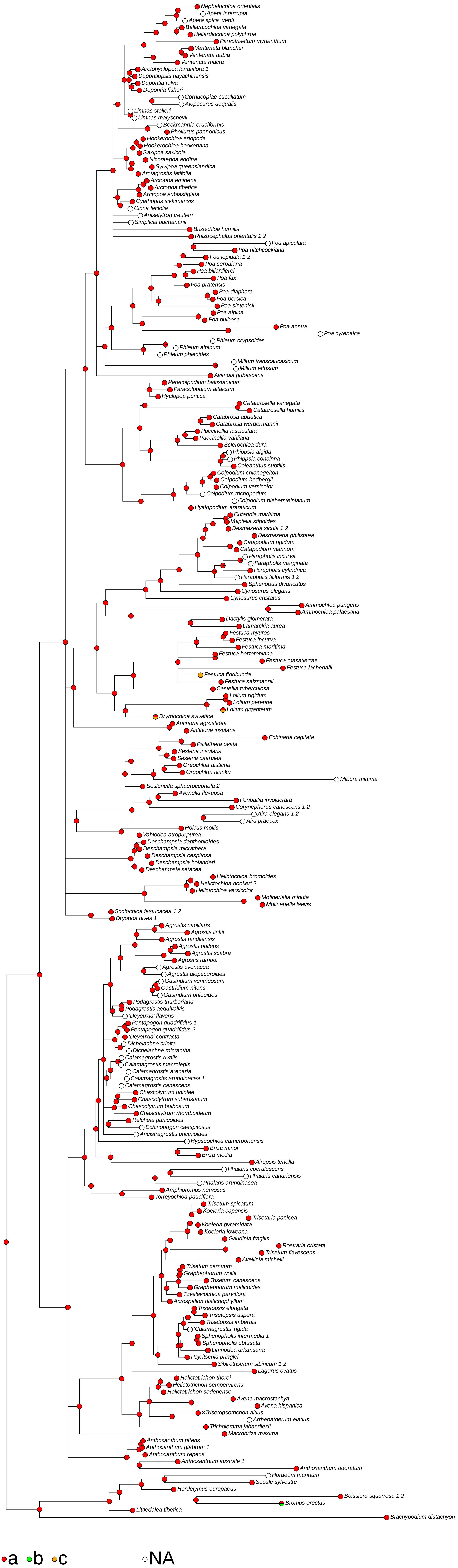

### (071) Glumes <persistence - if spikelets break up>

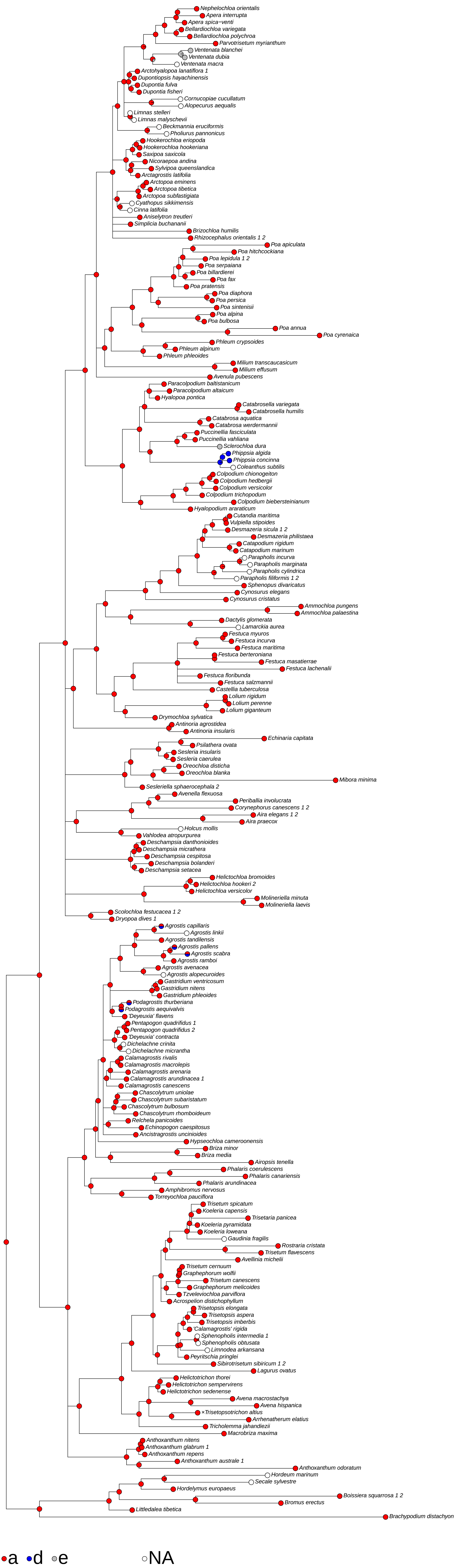

### (072) Glumes <similarity, apart from some difference in size>

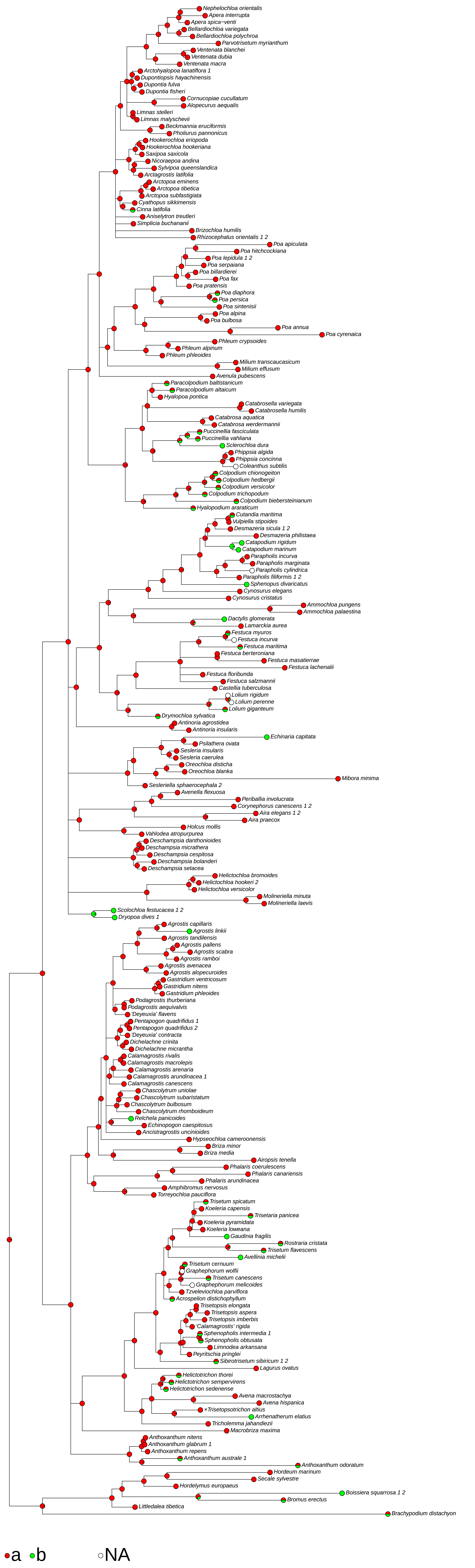

### (073) Glumes <length of longer relative to rest of spikelet, excluding awns>

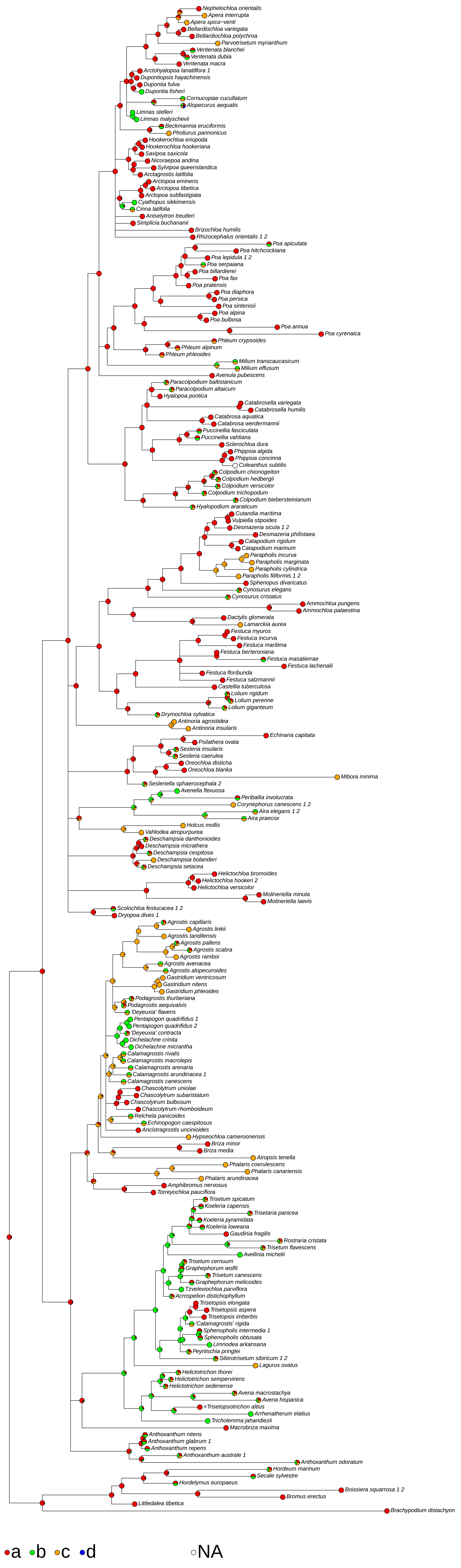

### (074) Glumes <consistency relative to fertile lemma>

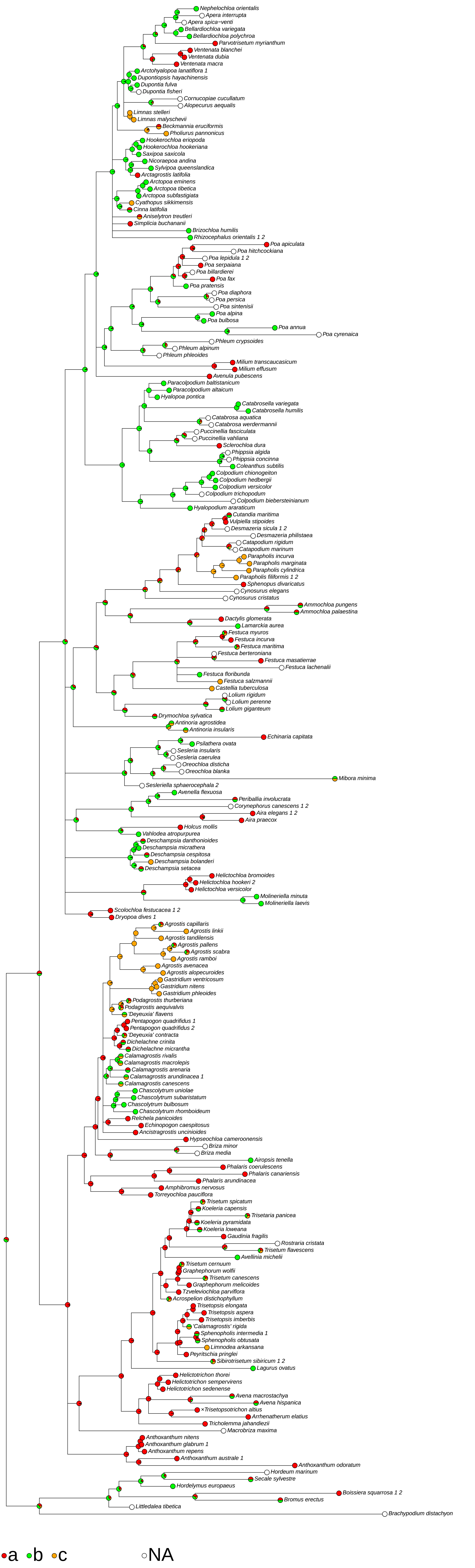

### (078) Lower glume <length in mm>

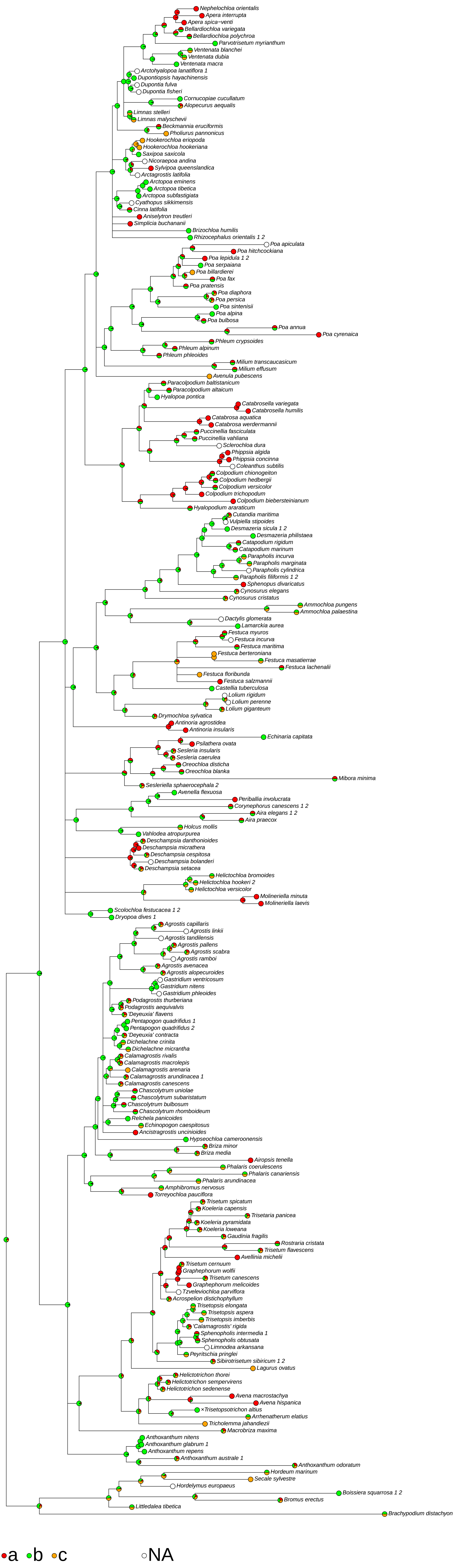

### (079) Lower glume <length as fraction of upper>

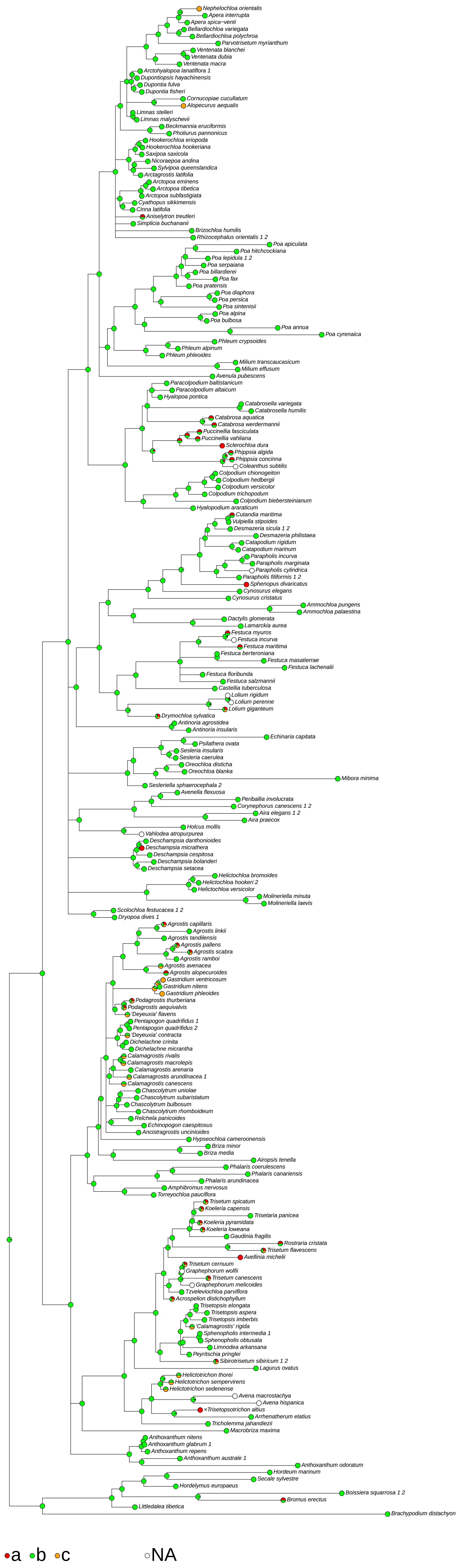

### (081) Lower glume <marginal consistency>

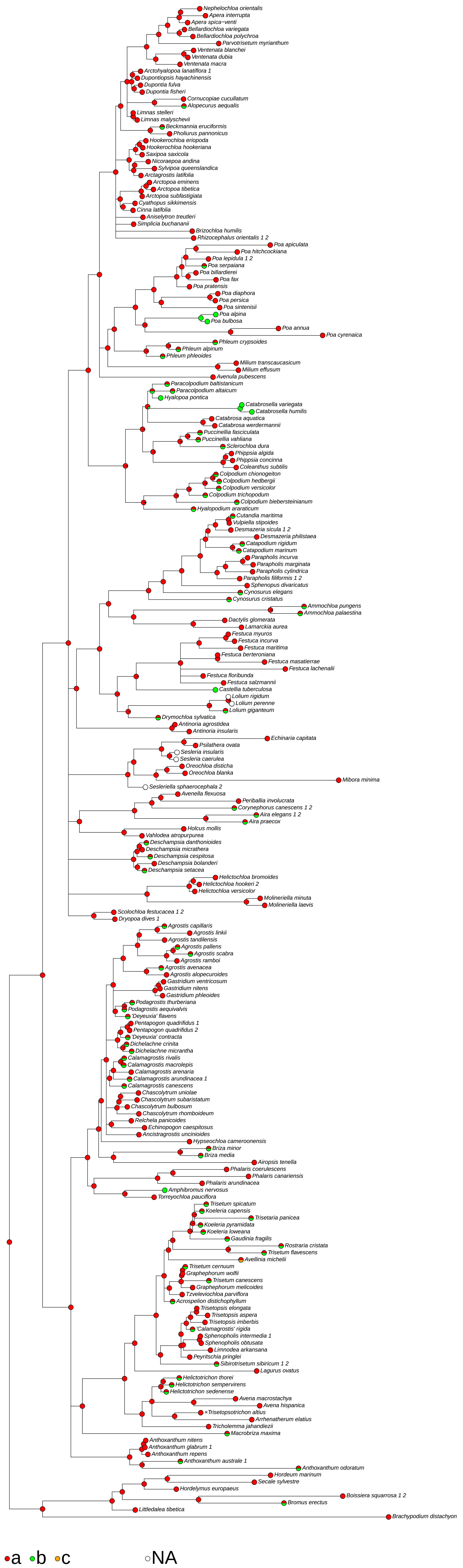

### (082) Lower glume <presence of keels>

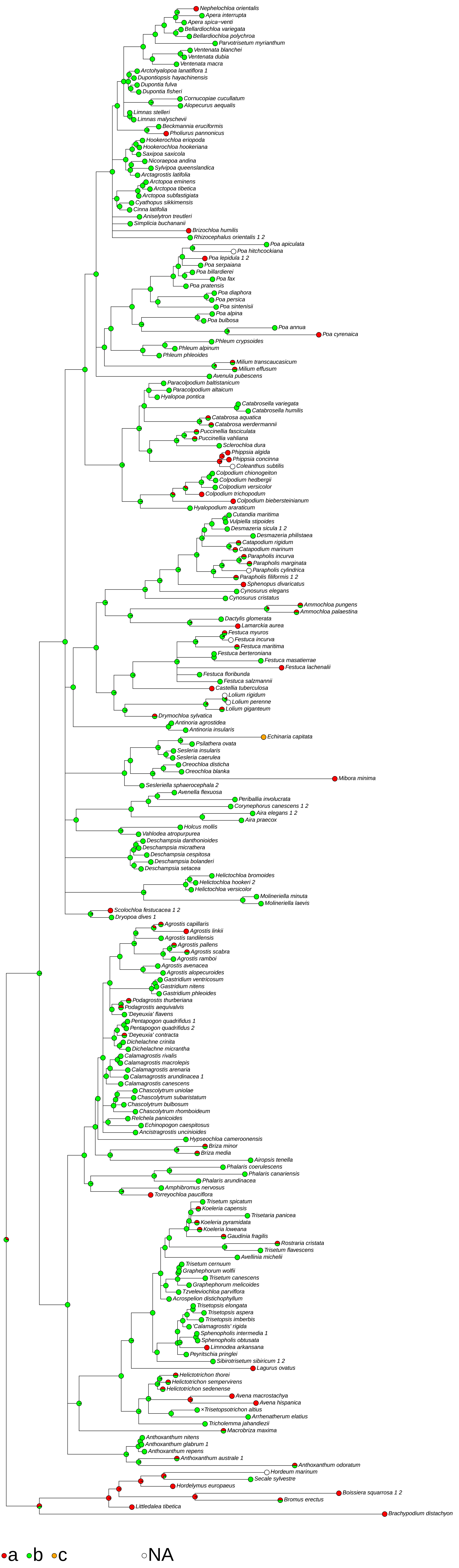

### (083) Lower glume keeled <extent>

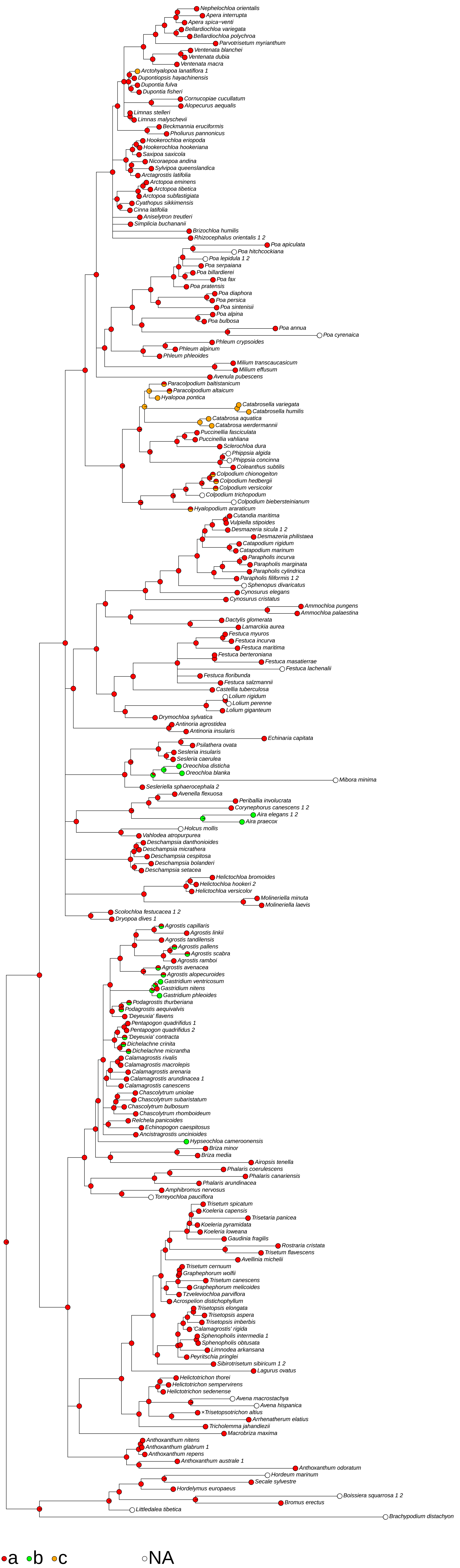

### (085) Lower glume <vein number>

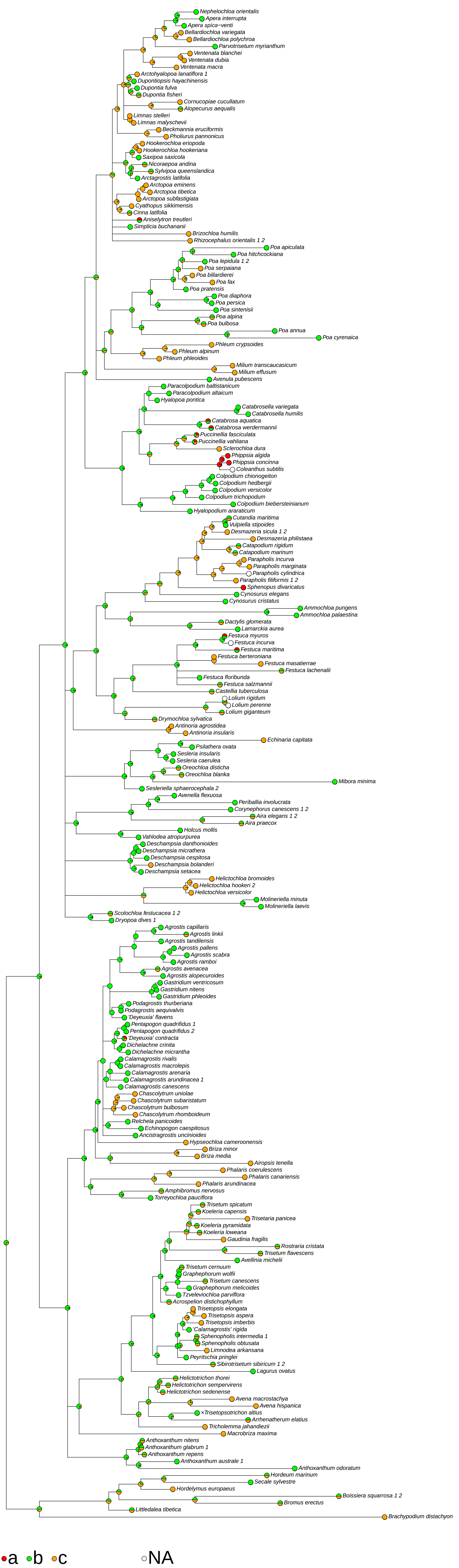

### (087) Lower glume primary vein <presence of hairs or spines>

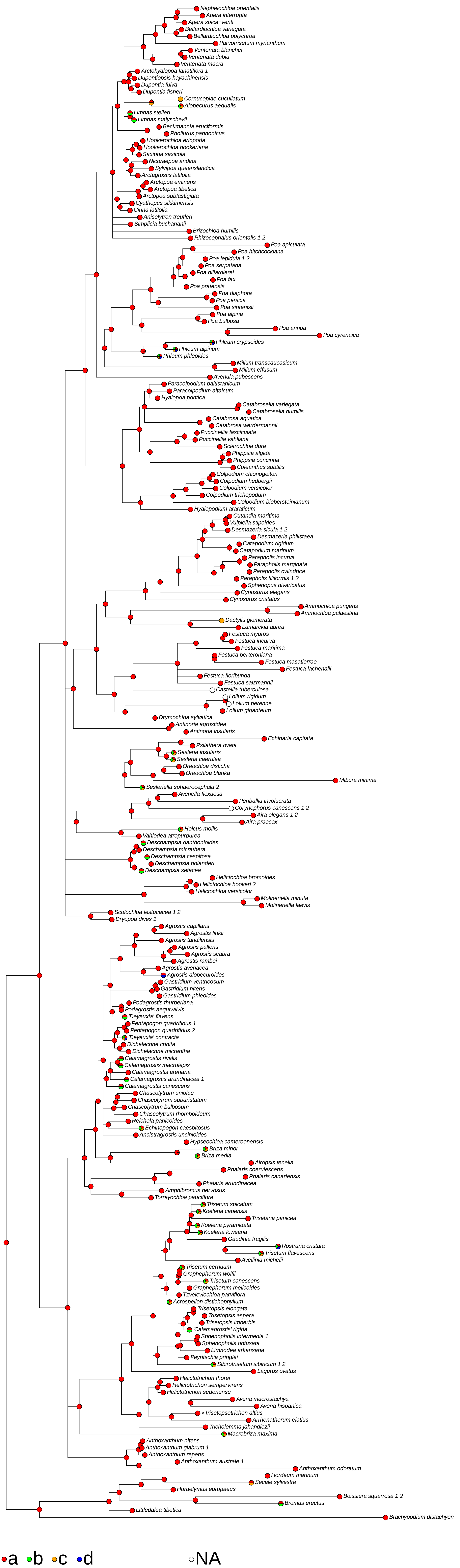

### (088) Lower glume lateral veins <clarity>

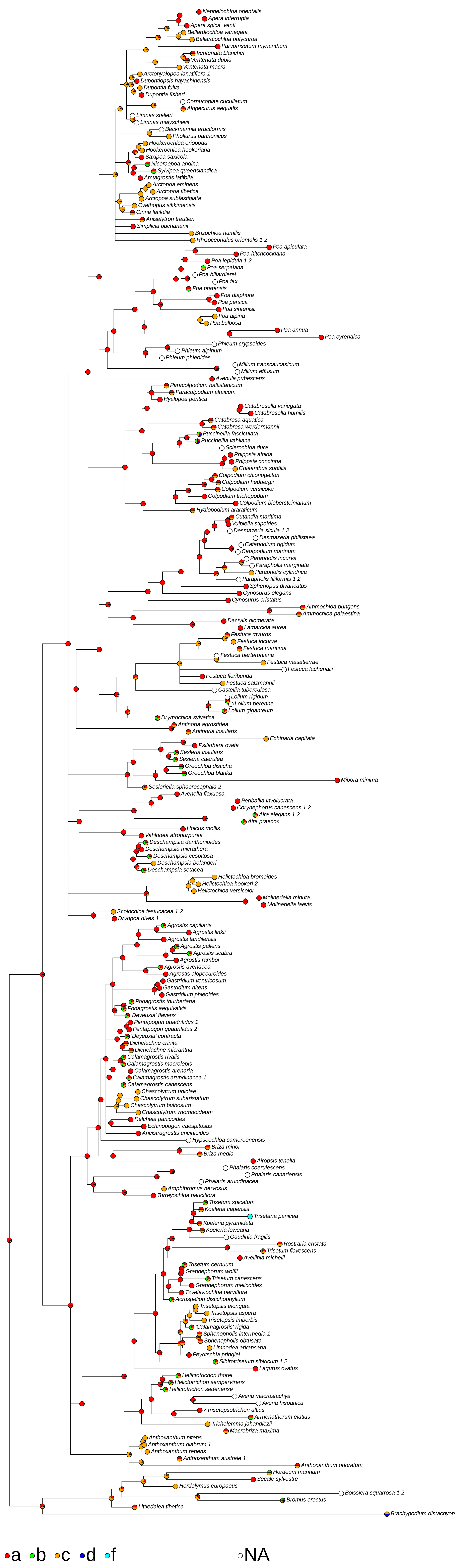

### (095) Upper glume <length in mm>

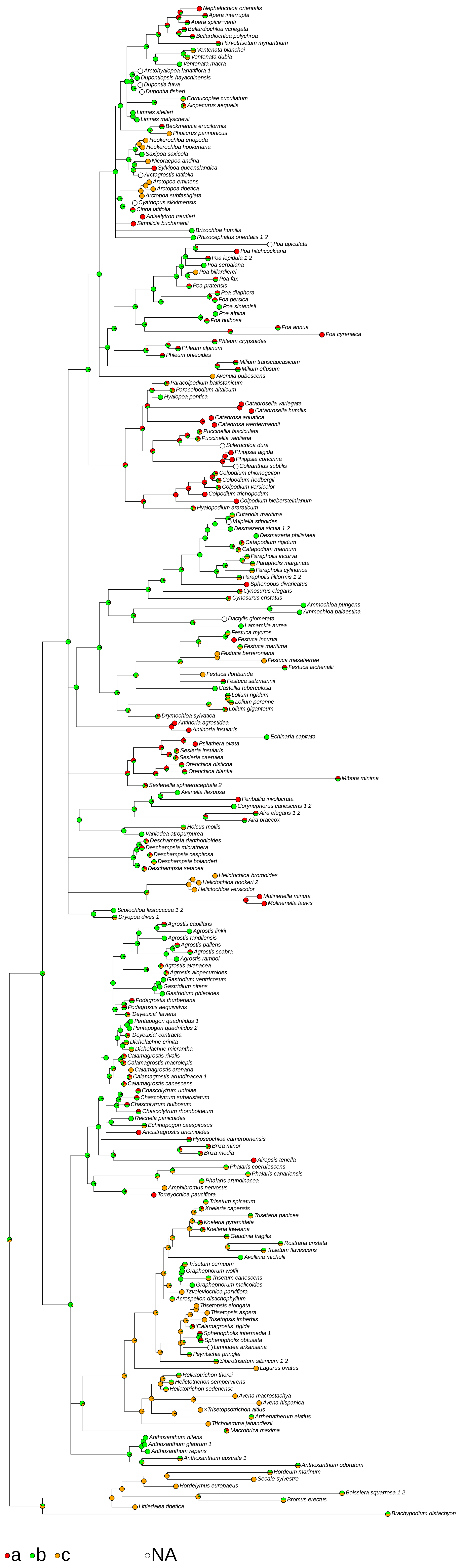

### (096) Upper glume <length as fraction of lemma>

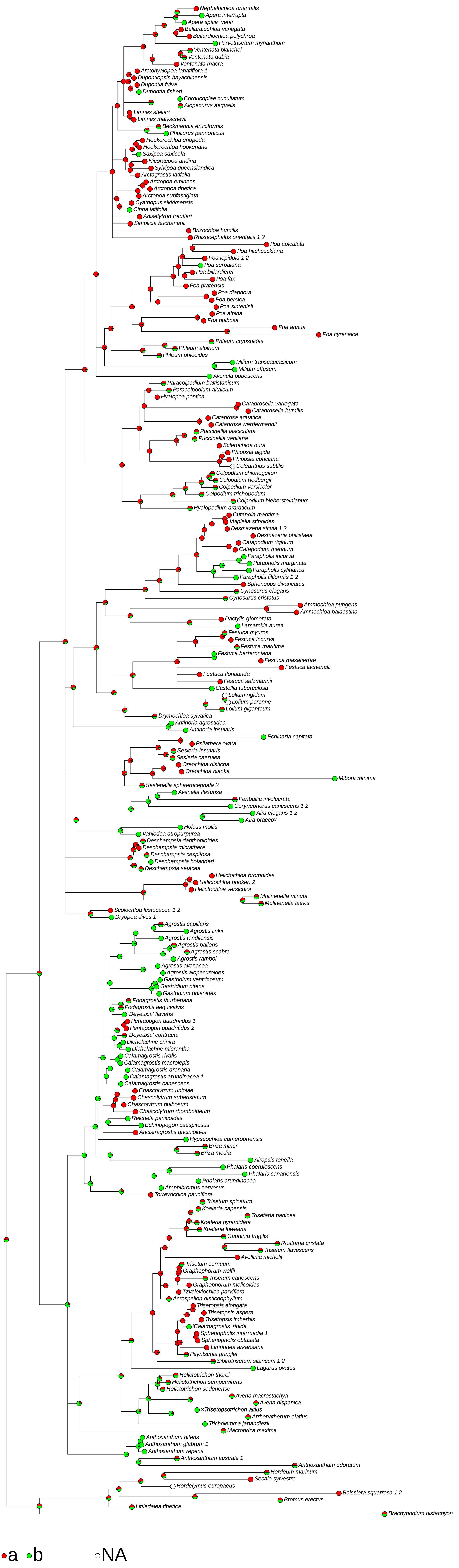

### (098) Upper glume with <marginal consistency>

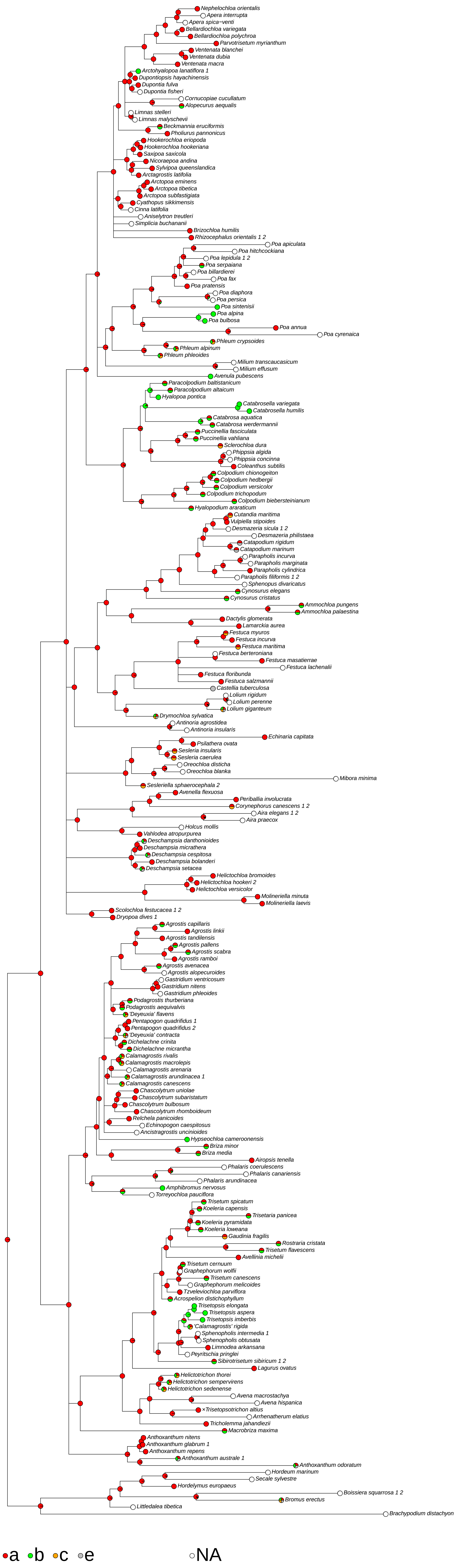

### (099) Upper glume <keels>

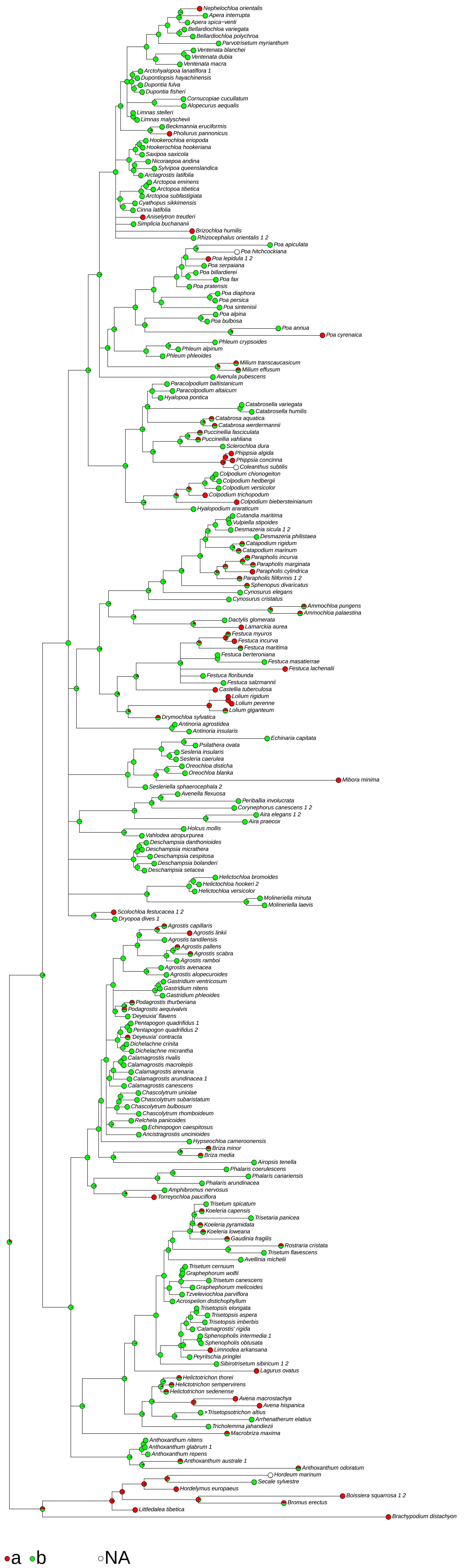

### (100) Upper glume keeled <extent>

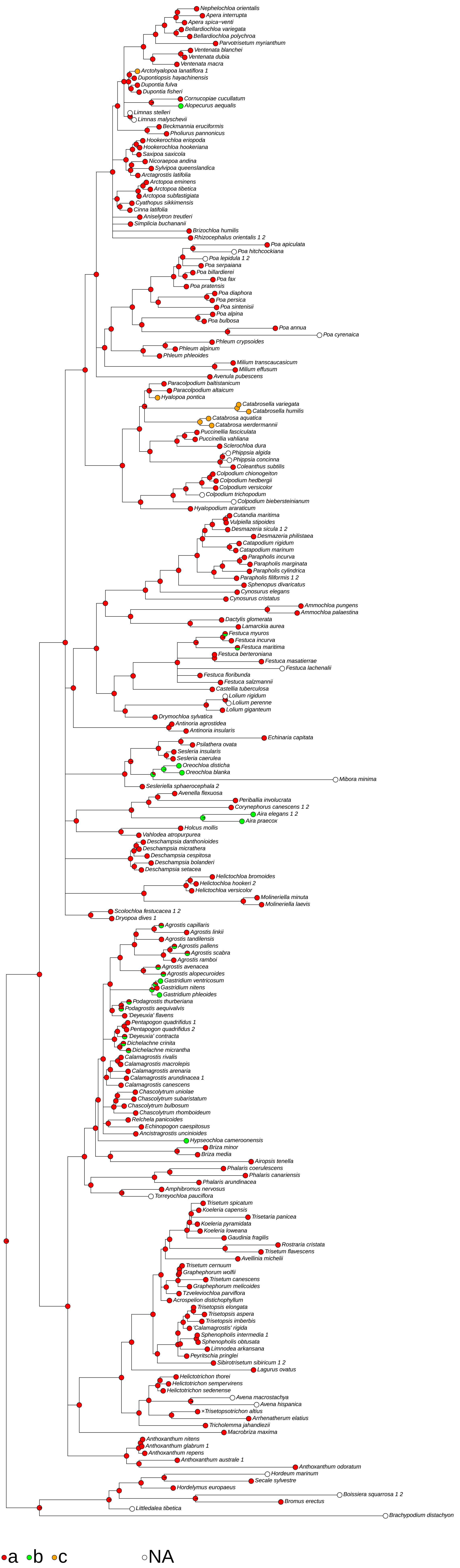

### (102) Upper glume <vein number>

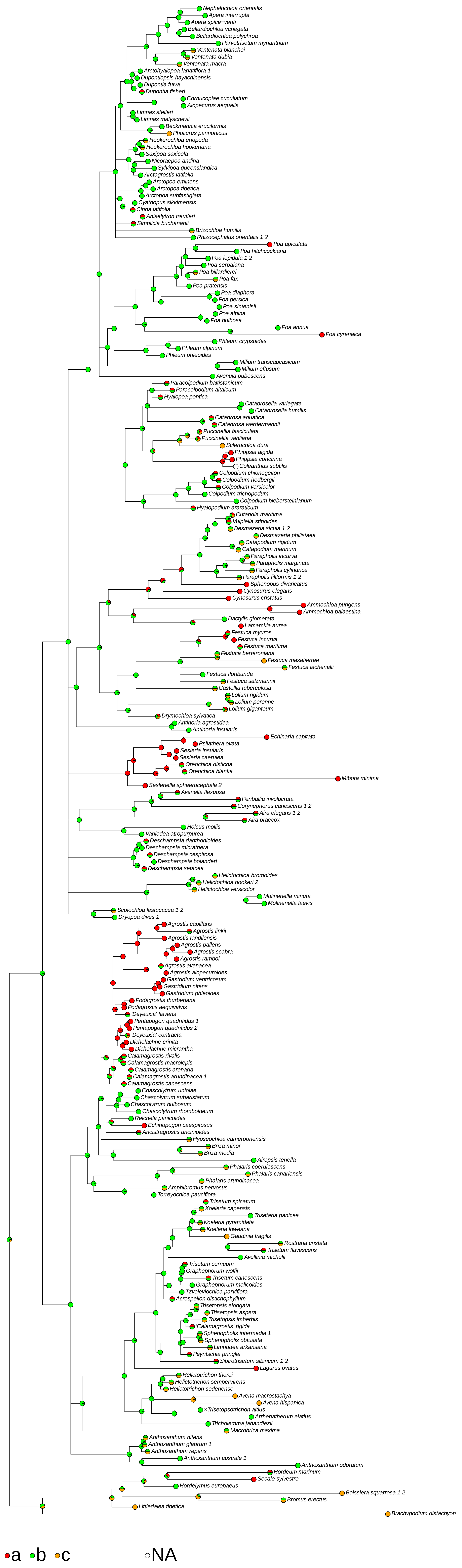

### (103) Upper glume primary vein <clarity of mid or keel vein>

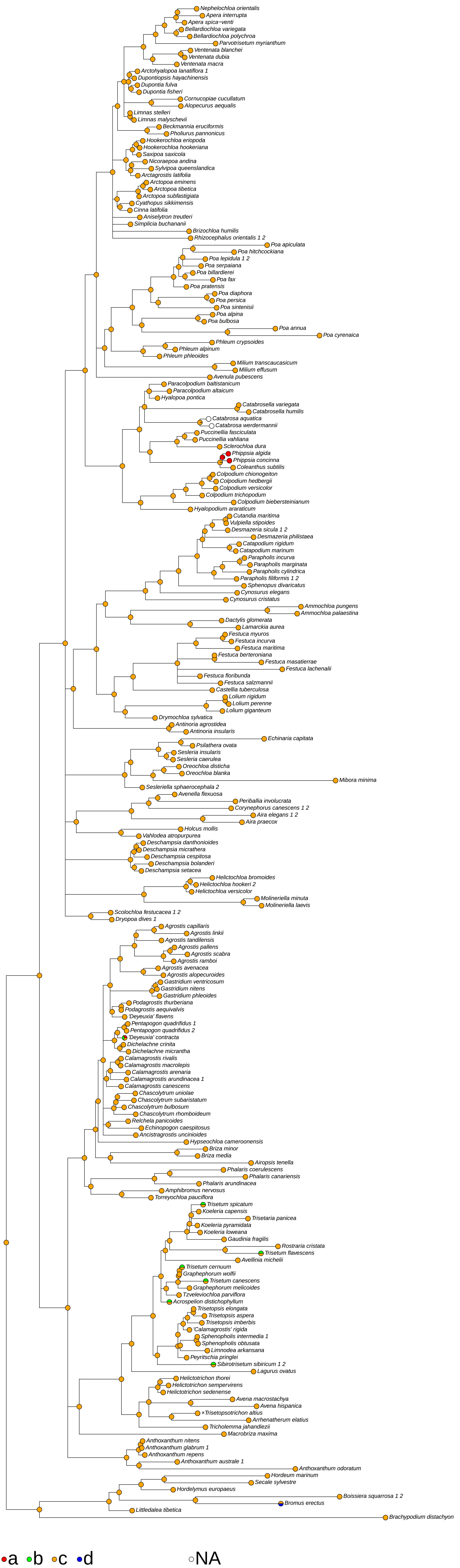

### (104) Upper glume primary vein <roughness>

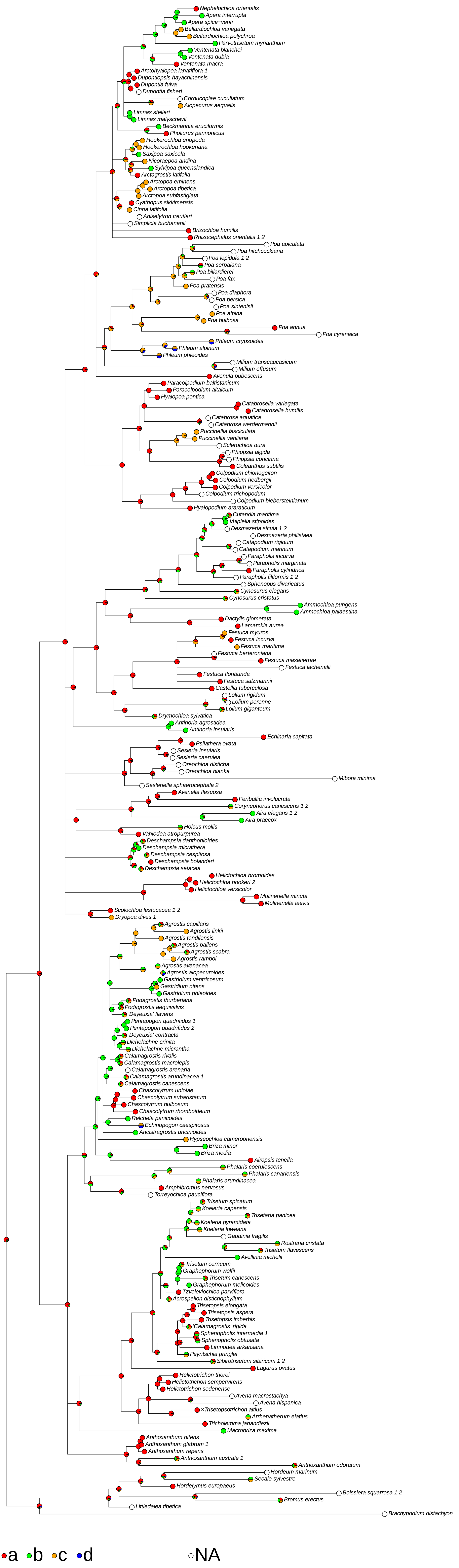

### (106) Upper glume lateral veins <clarity>

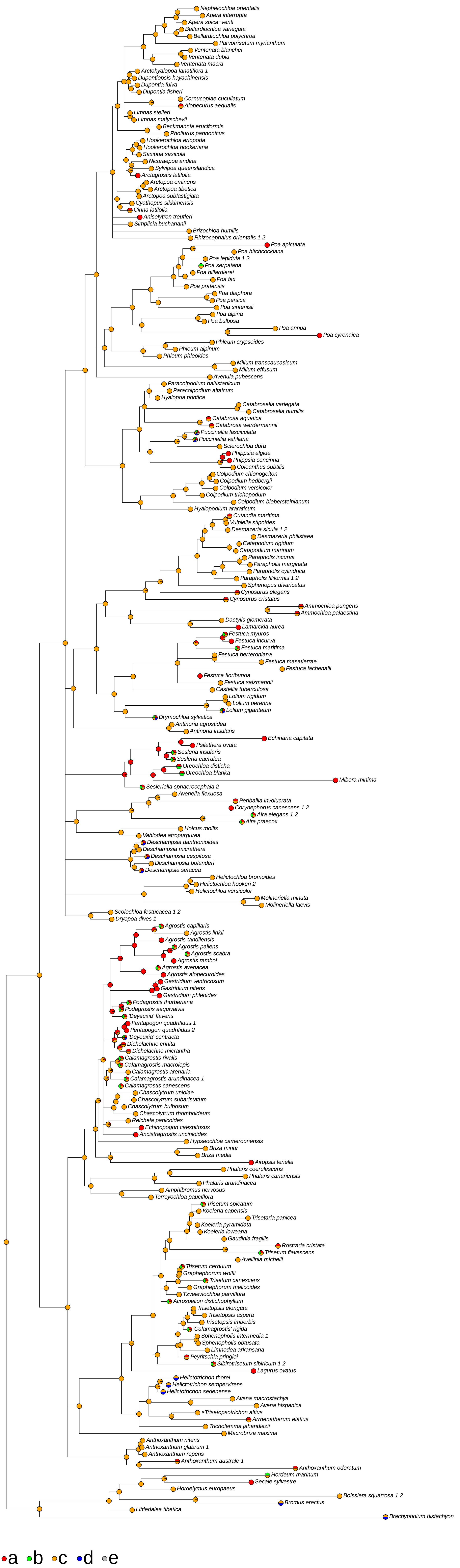

### (114) Upper glume <presence of awns>

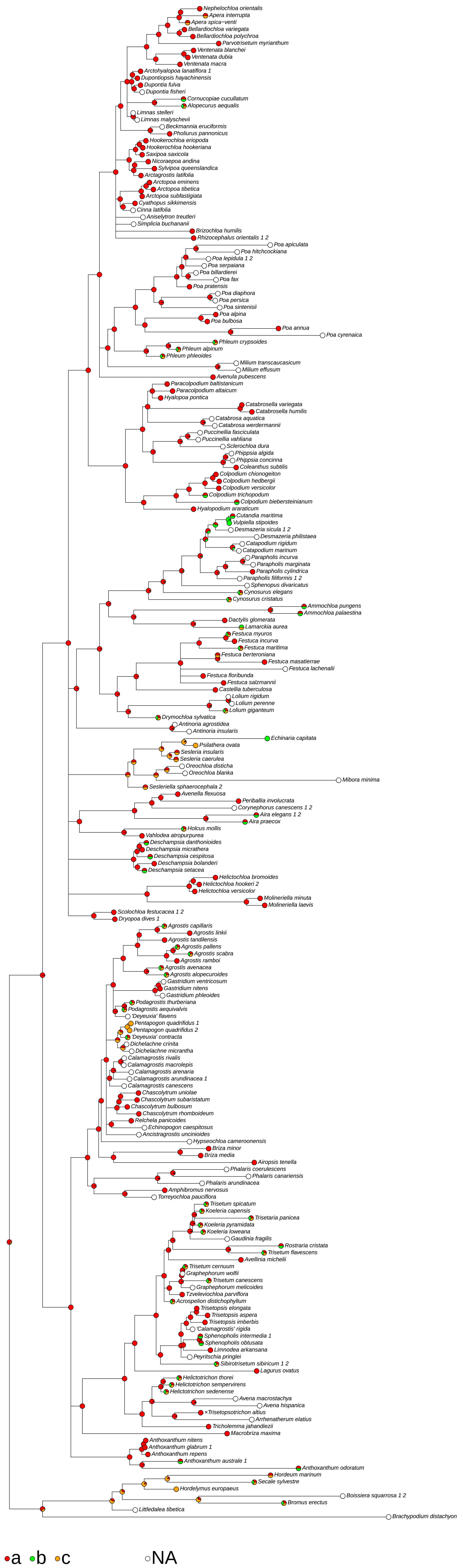

### (115) Basal sterile florets <presence, including vestiges>

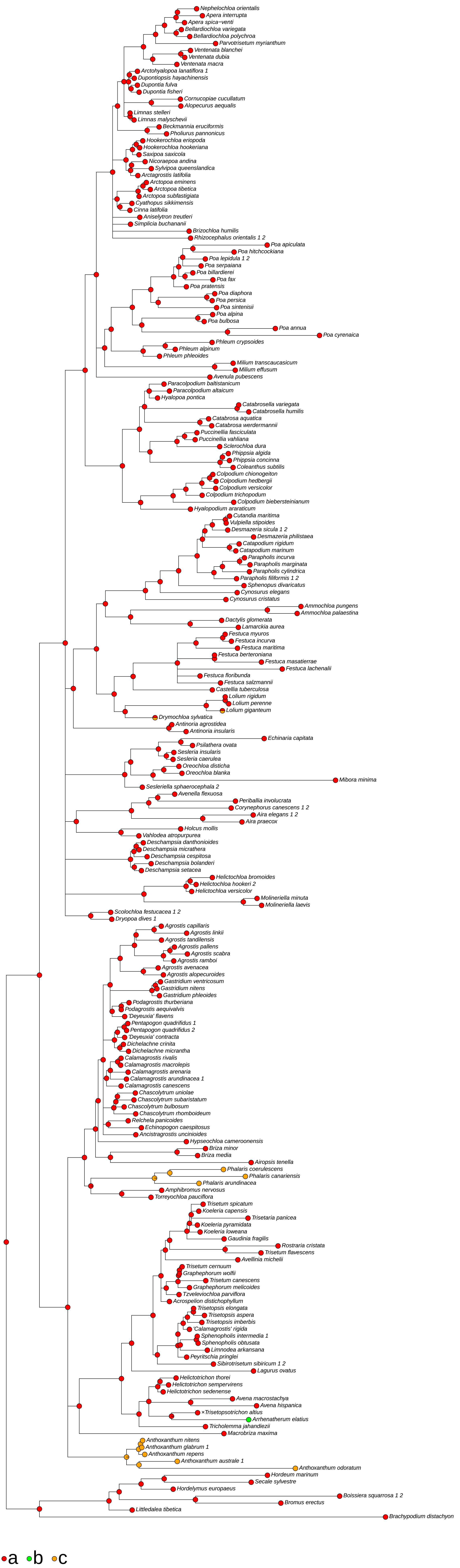

### (116) Fertile florets <similarity if more than 1>

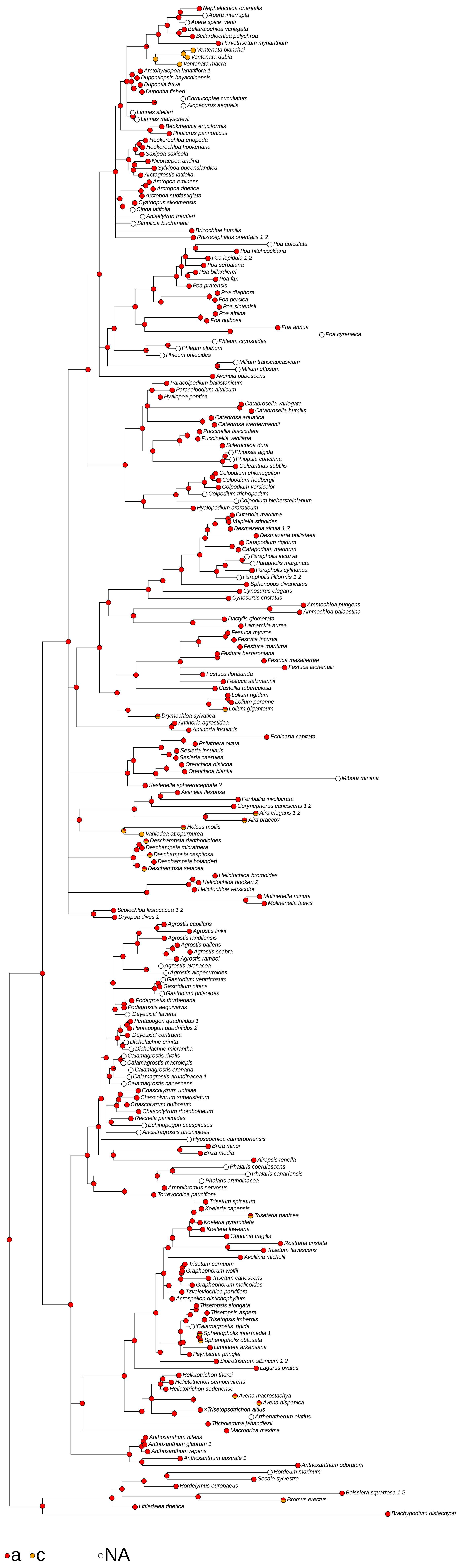

### (119) Fertile lemma <length in mm>

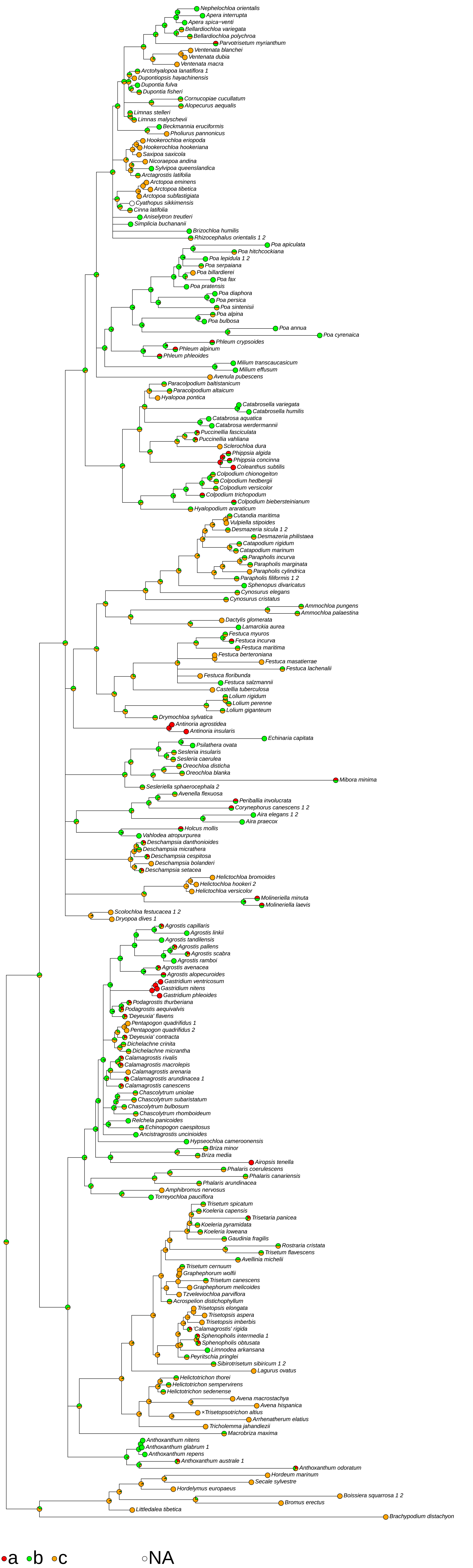

### (122) Fertile lemma <presence of keel>

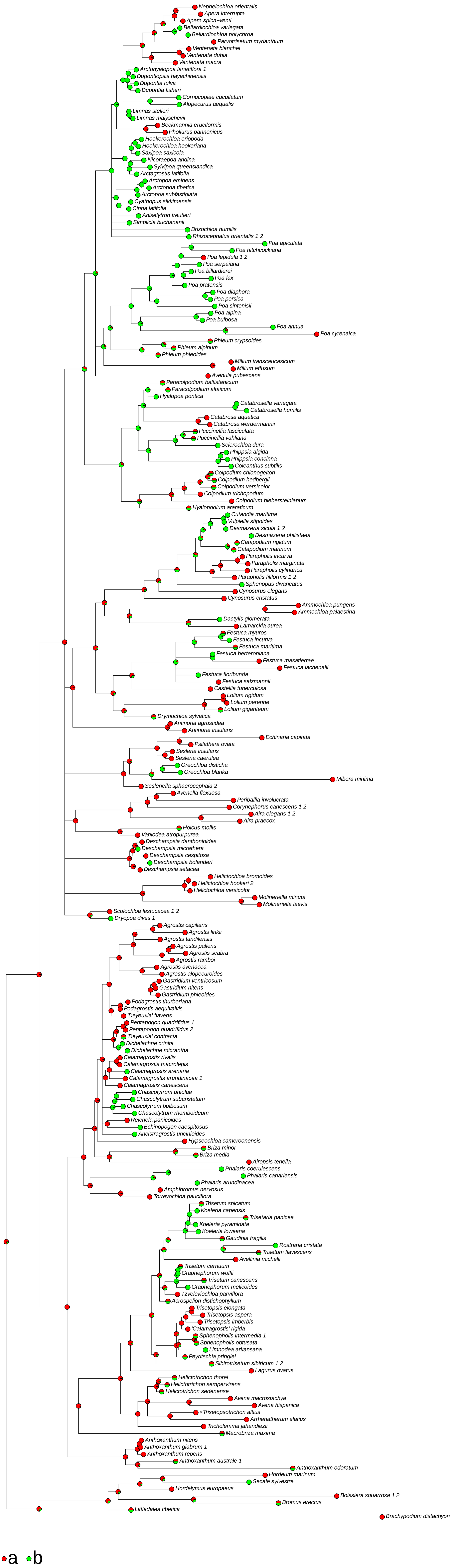

### (125) Fertile lemma <vein number>

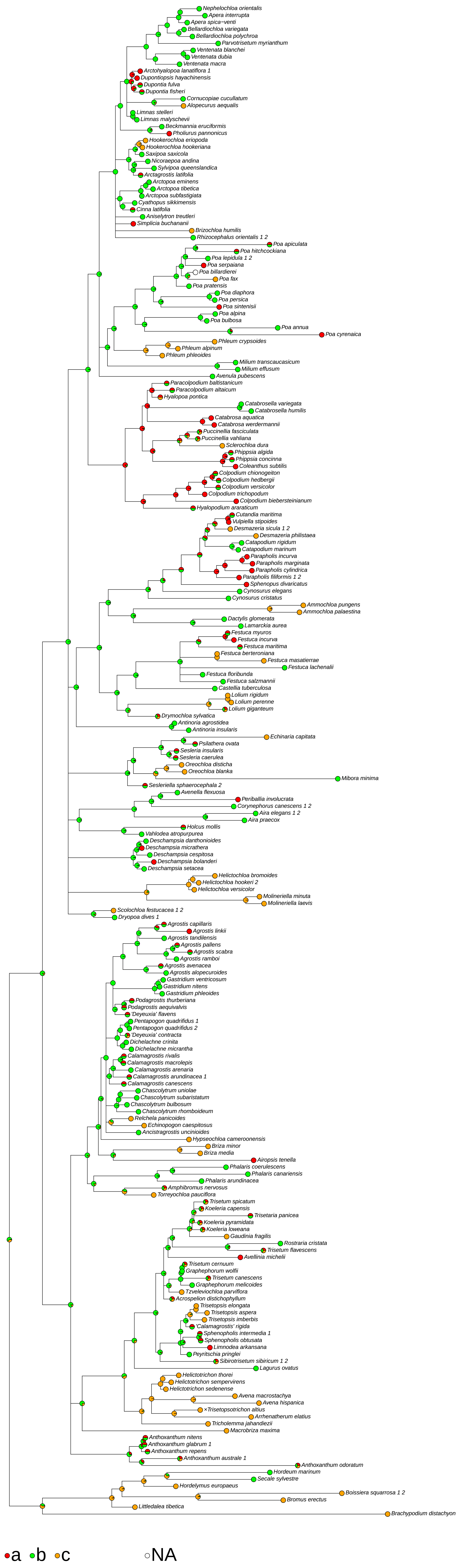

### (126) Fertile lemma

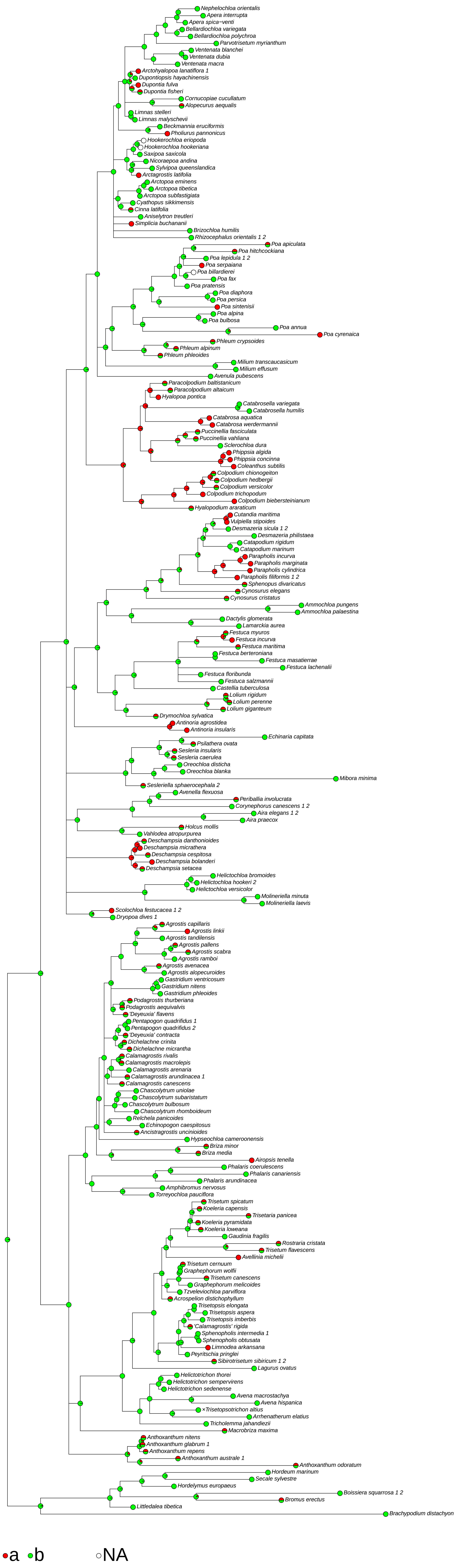

### (133) Lemma surface rough <where>

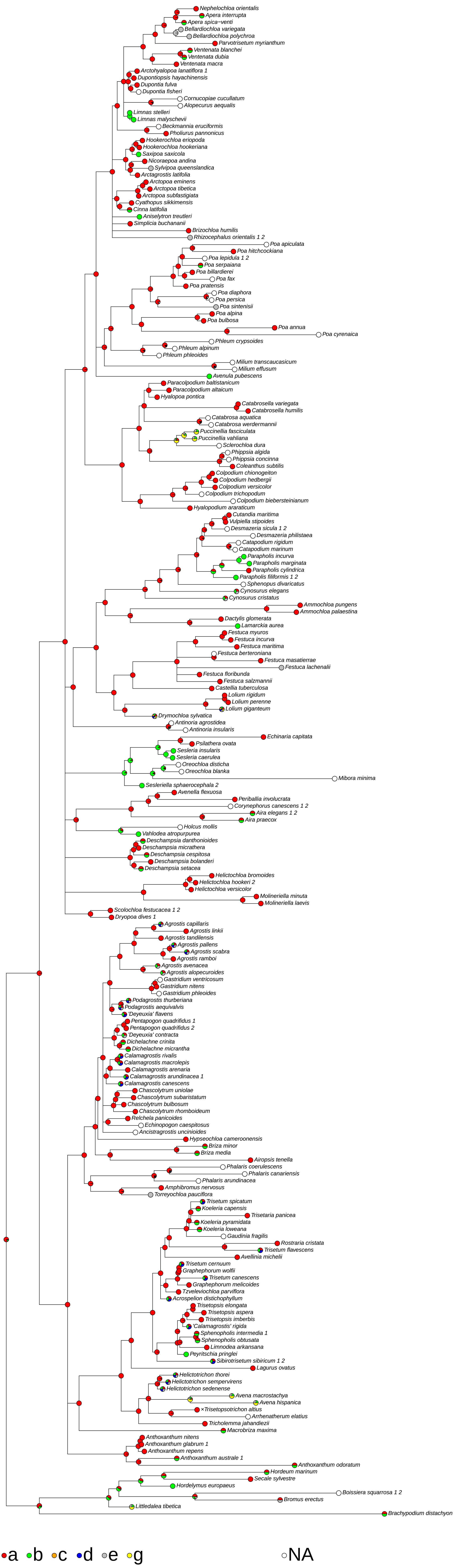

### (134) Lemma surface hairy <extent of general indumentum>

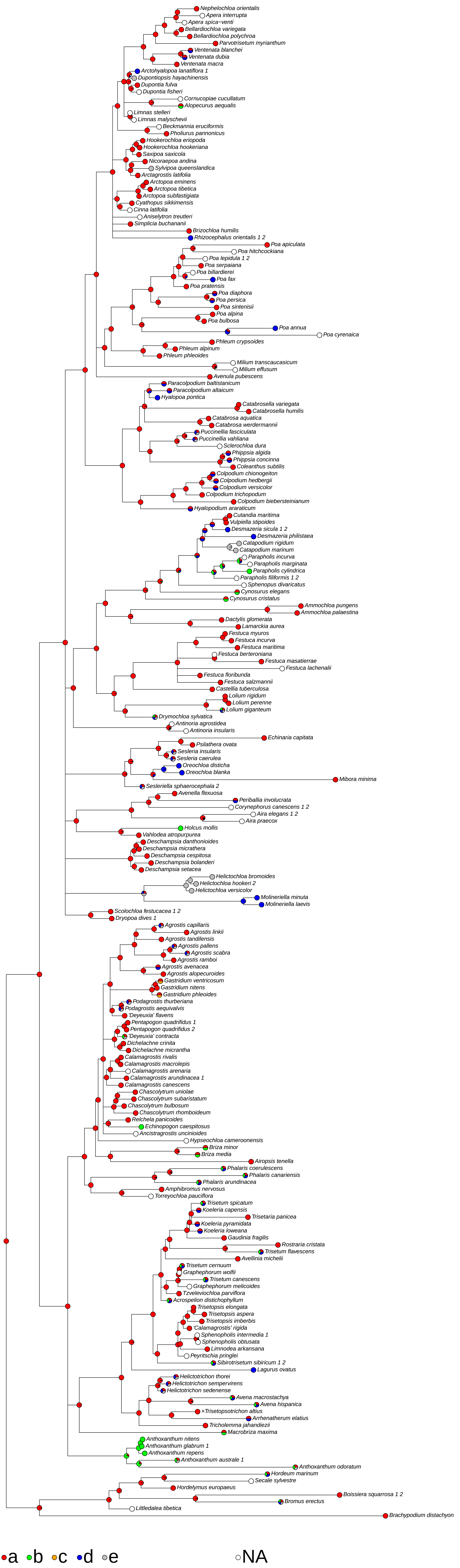

### (143) Lemma apex <incision>

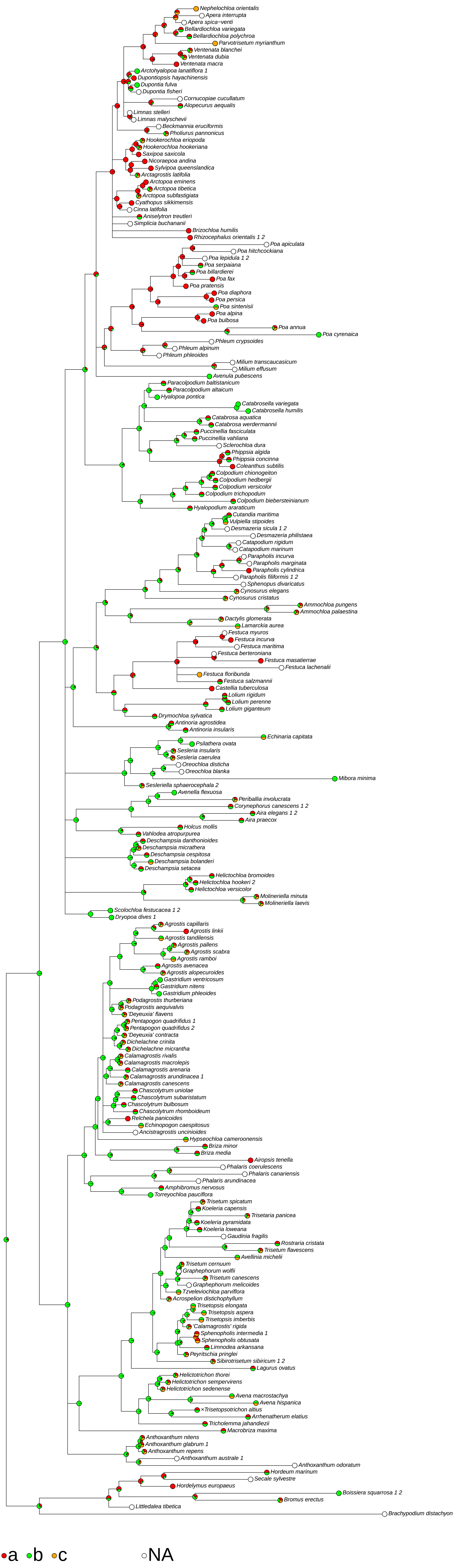

### (145) Lemma apex <equality of lobes>

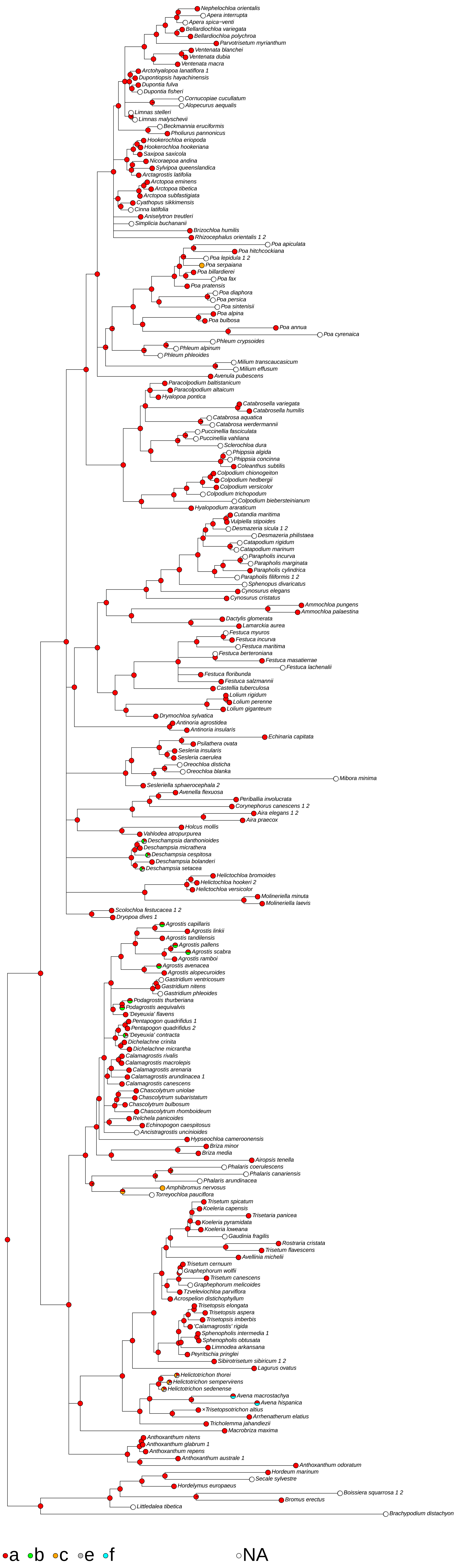

### (146) Lemma apex <presence of awn>

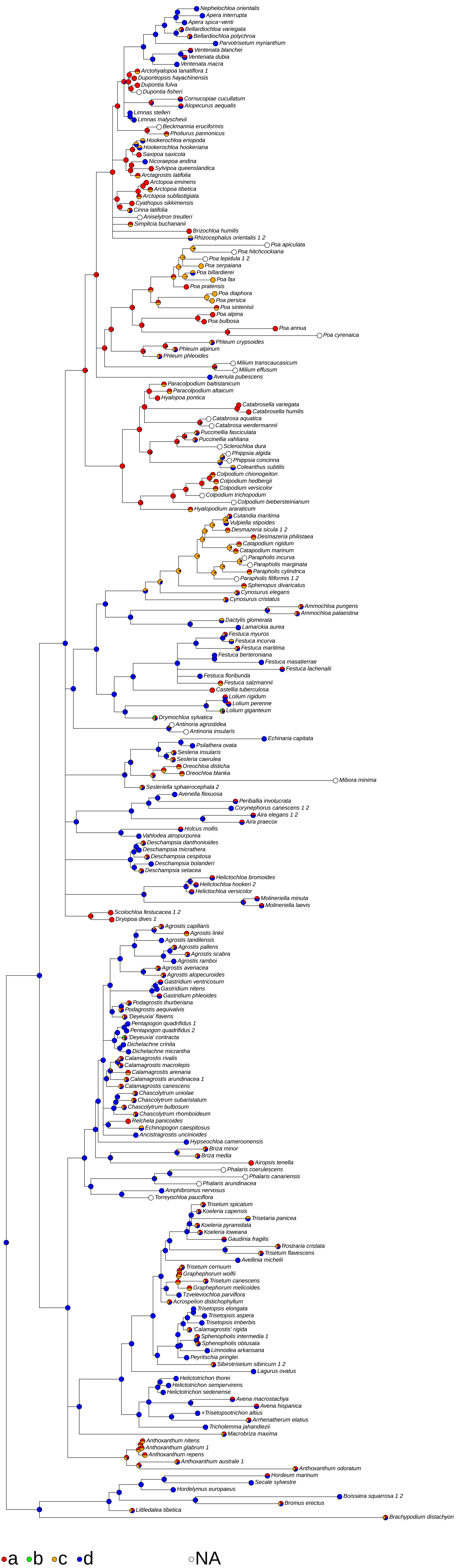

### (148) Principal lemma awn <position>

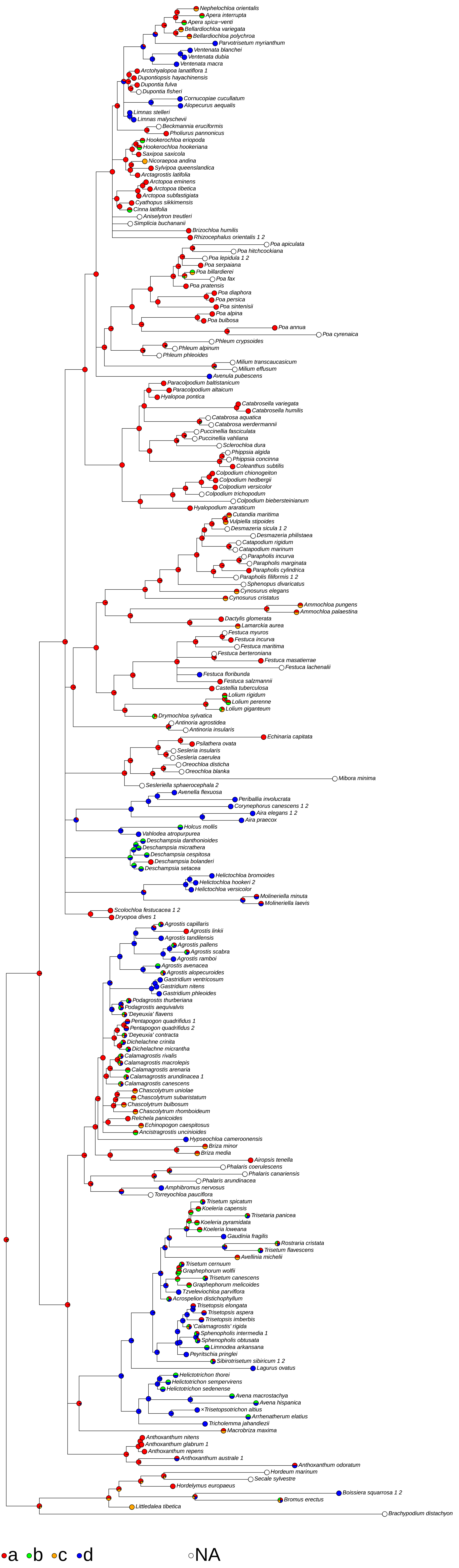

### (150) Principal lemma awn <overall shape>

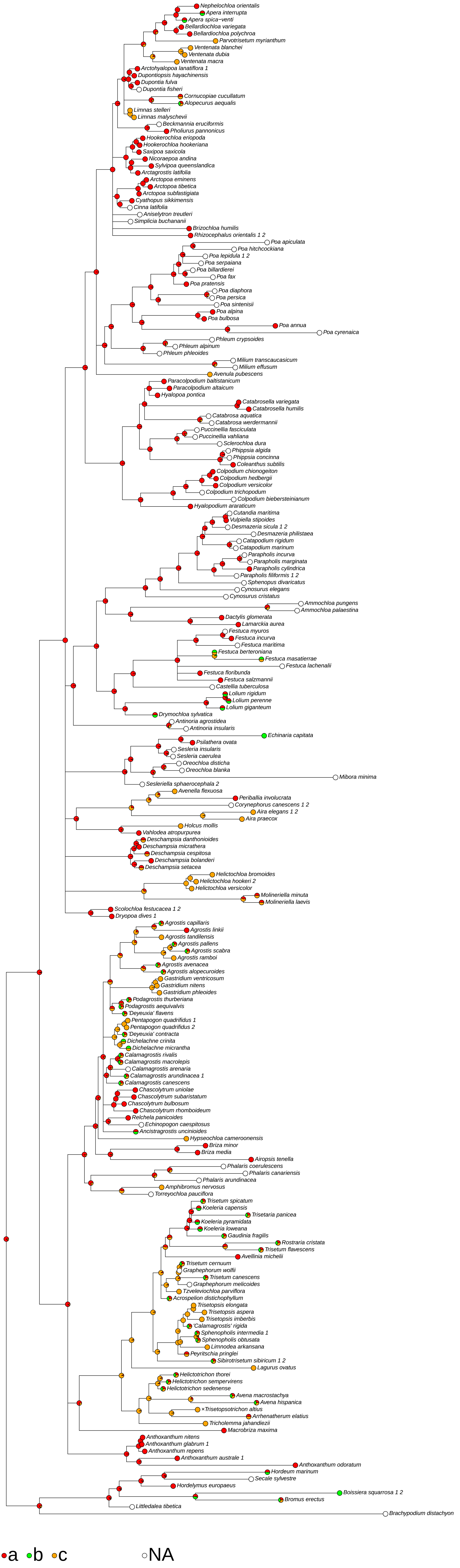

### (151) Principal lemma awn <flexion relative to lemma or column>

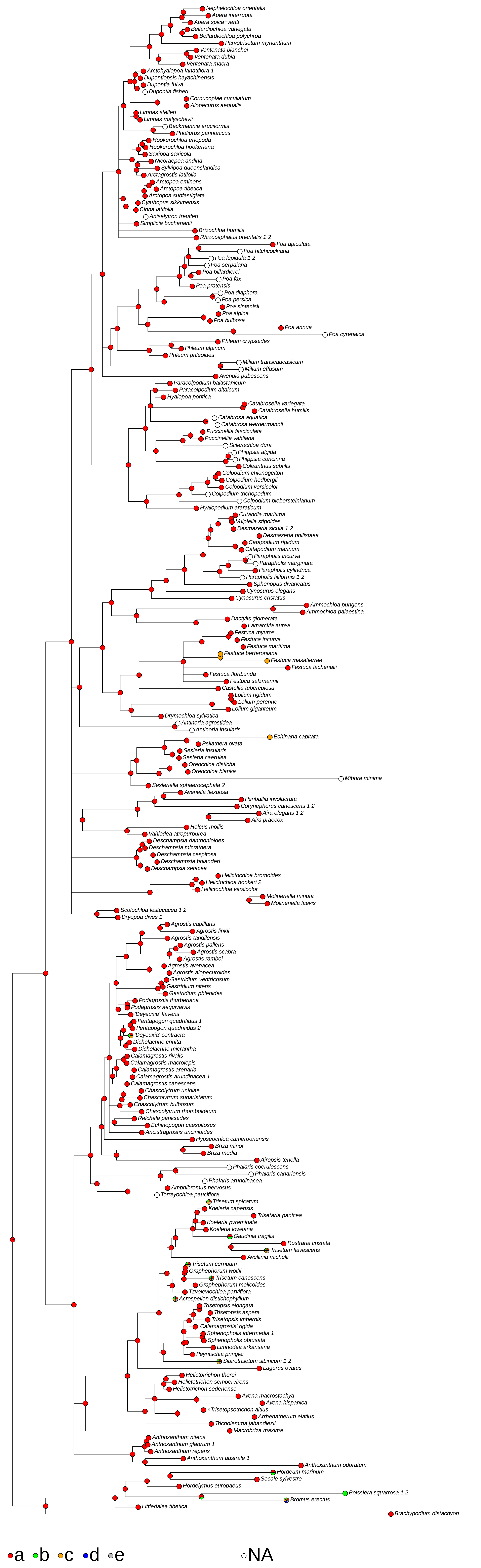

### (154) Principal lemma awn <exsertion>

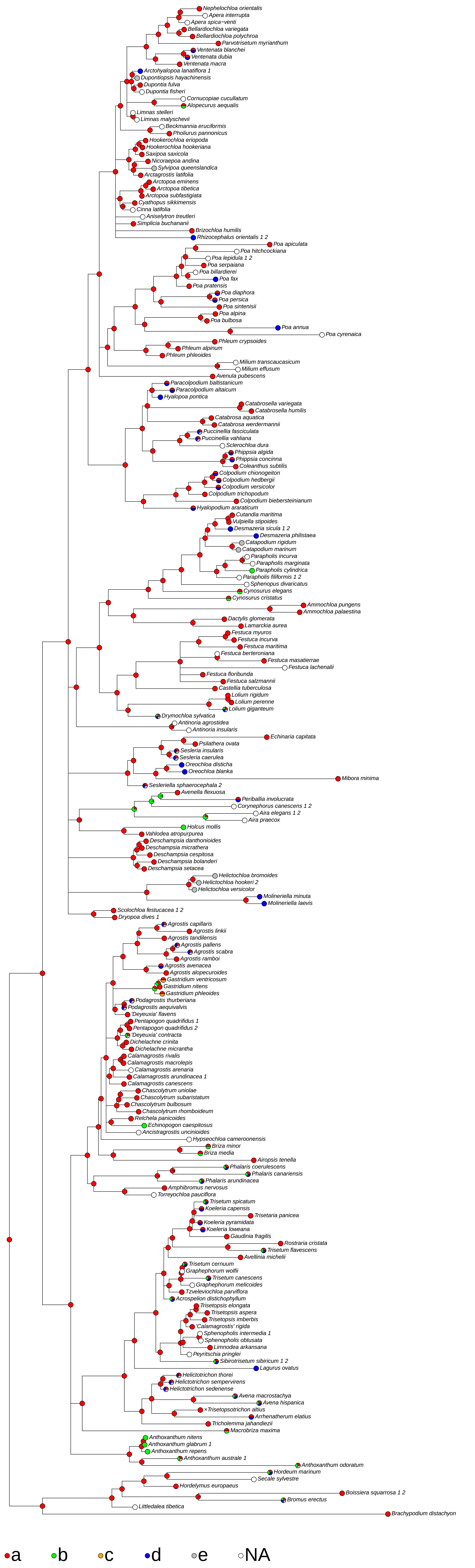

### (155) Principal lemma awn <presence of column>

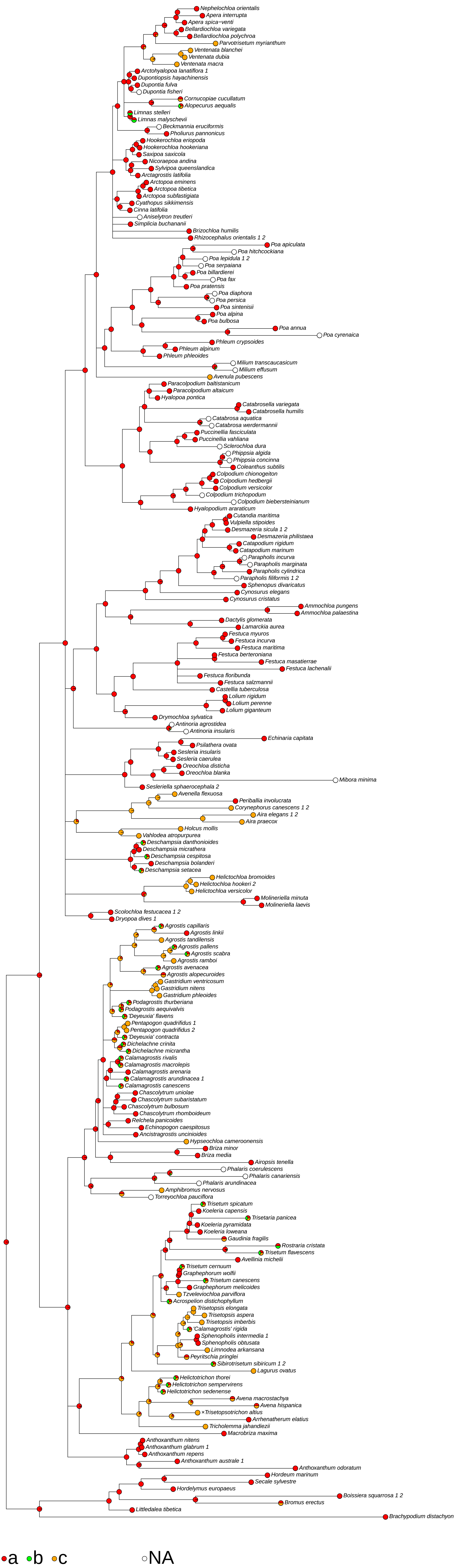

### (156) Principal lemma awn <coloured>

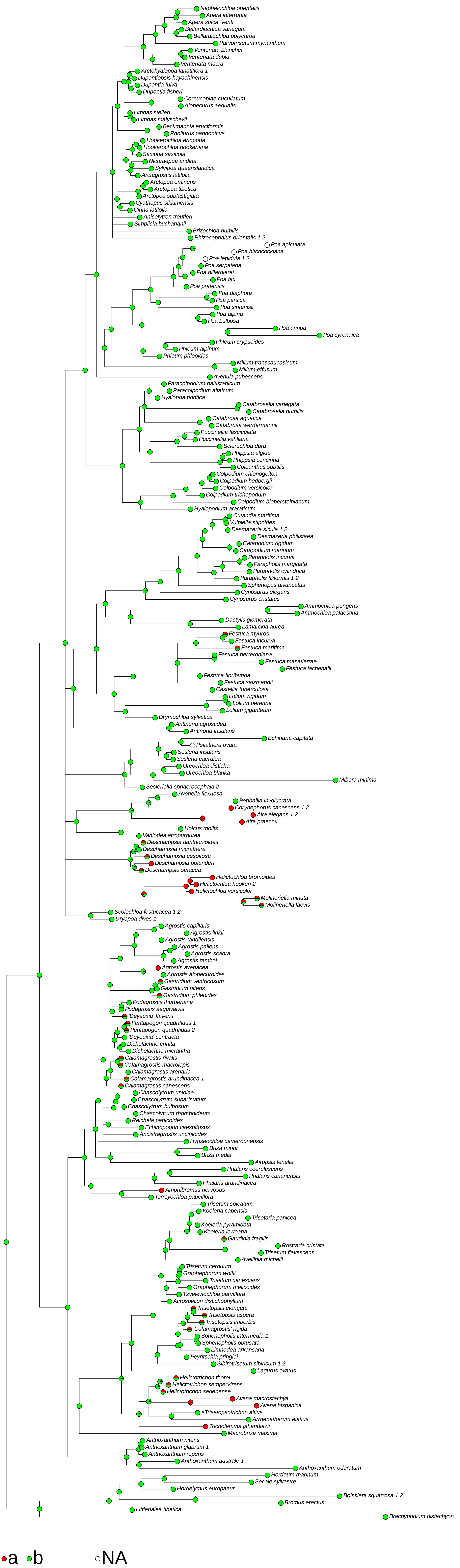

### (159) Column of lemma awn <flattened>

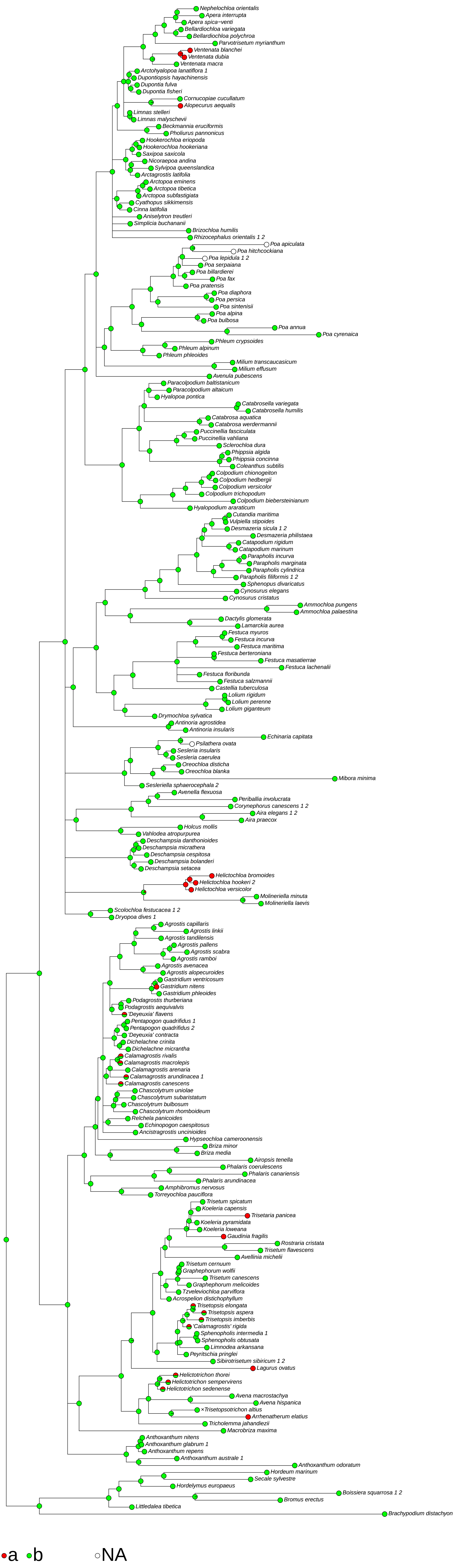

### (160) Column of lemma awn <twisted>

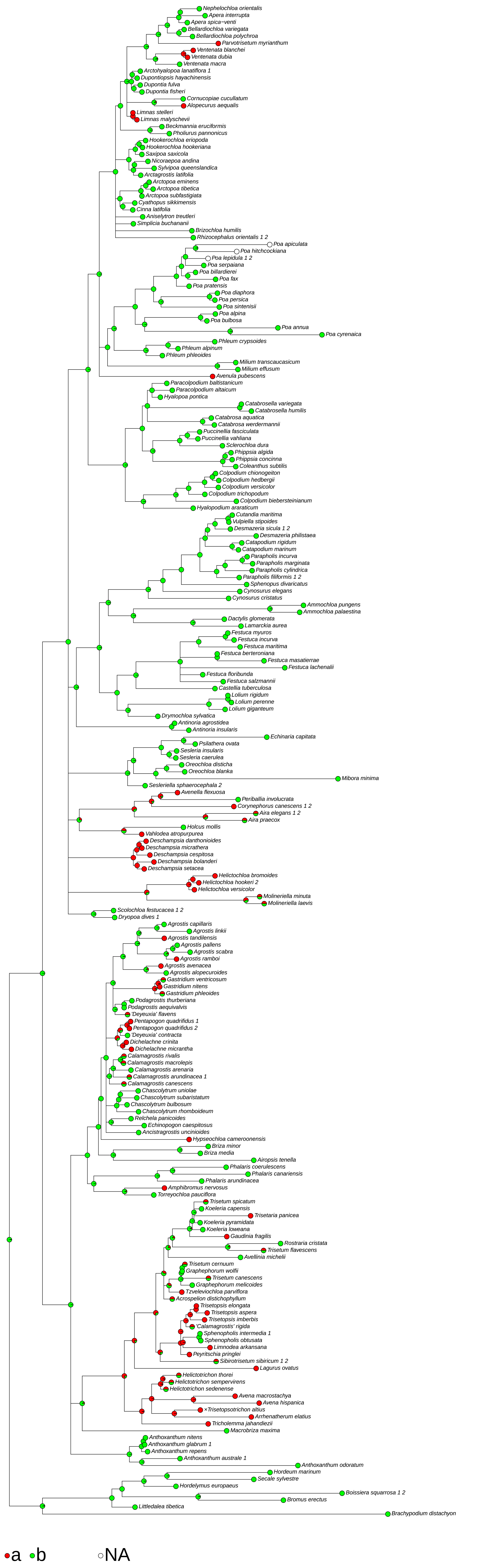

### (161) Lateral lemma awns <presence>

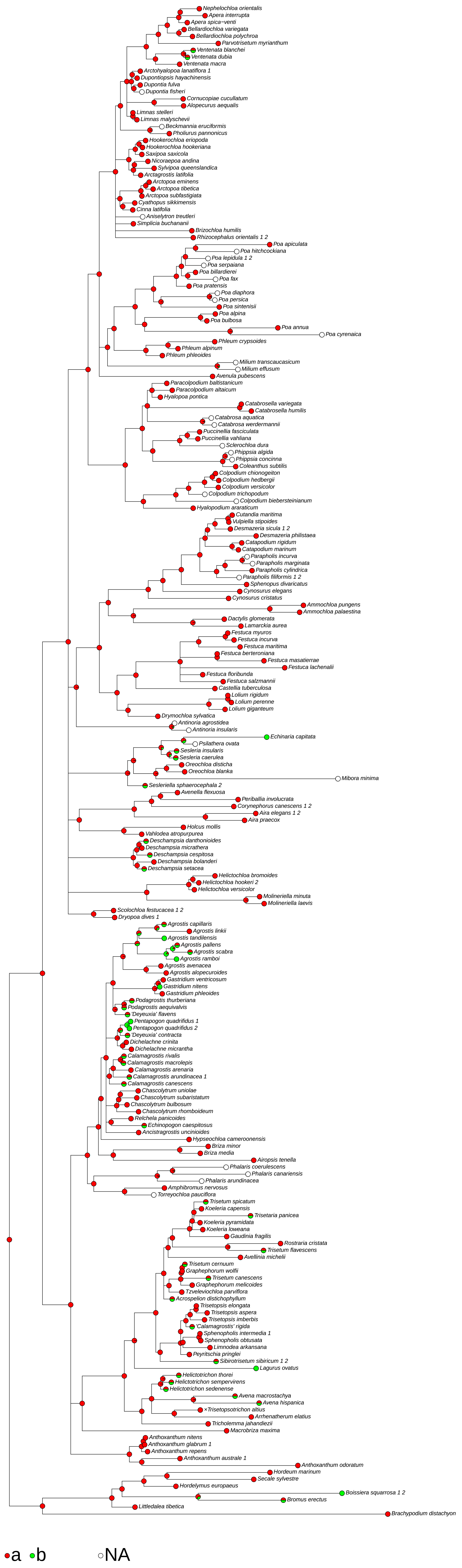

### (166) Palea <fraction of lemma length>

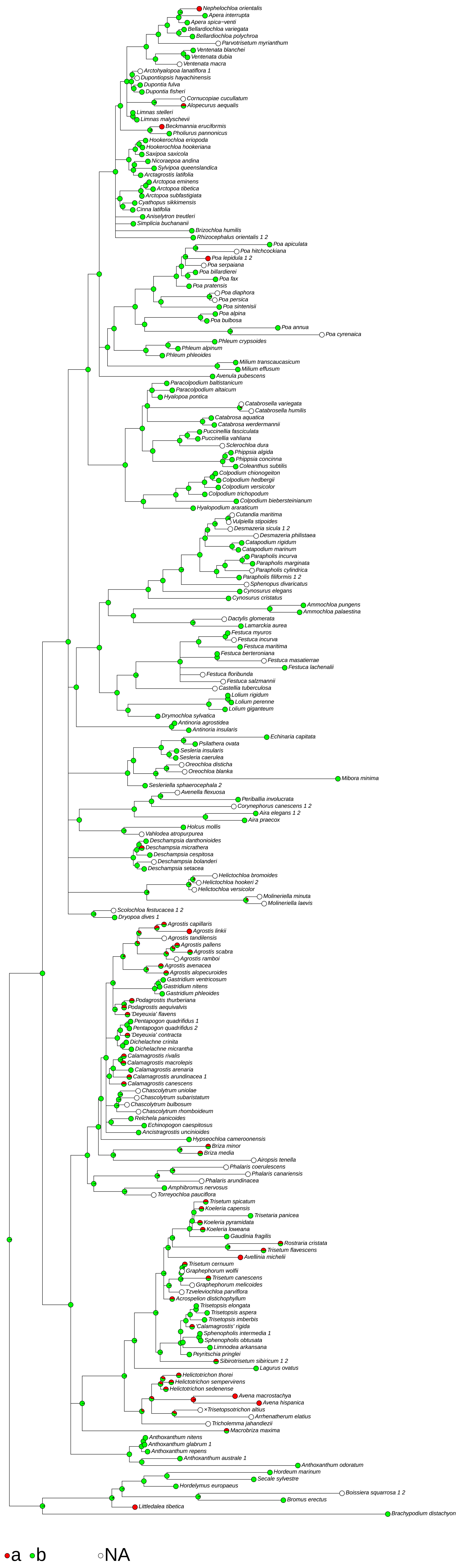

### (172) Palea keels <roughness>

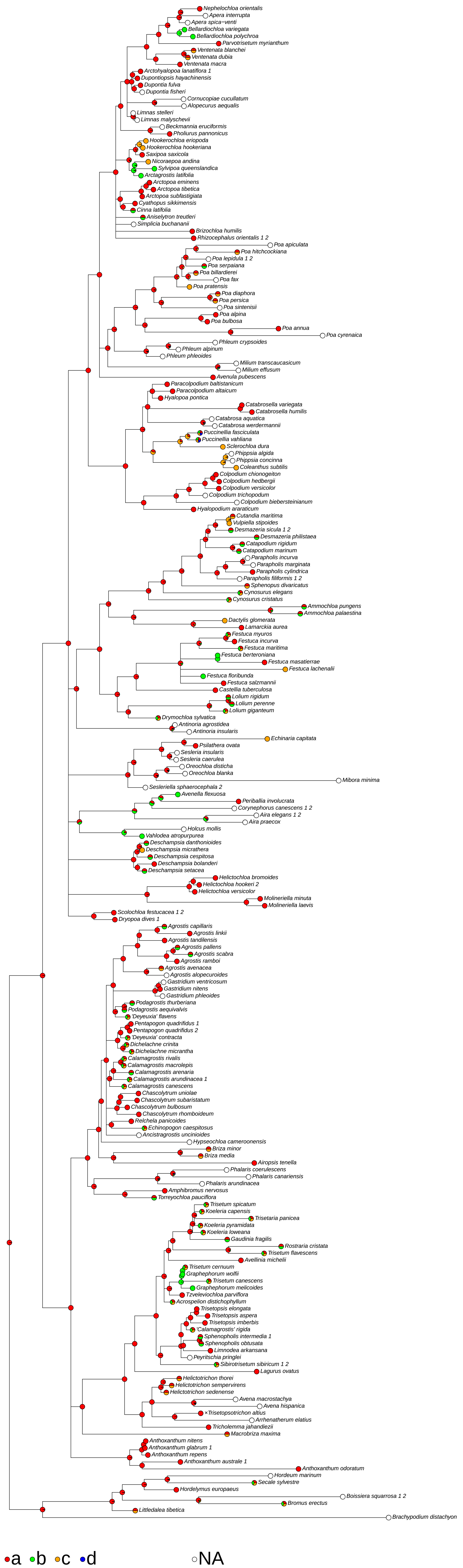

### (173) Palea keels <indumentum>

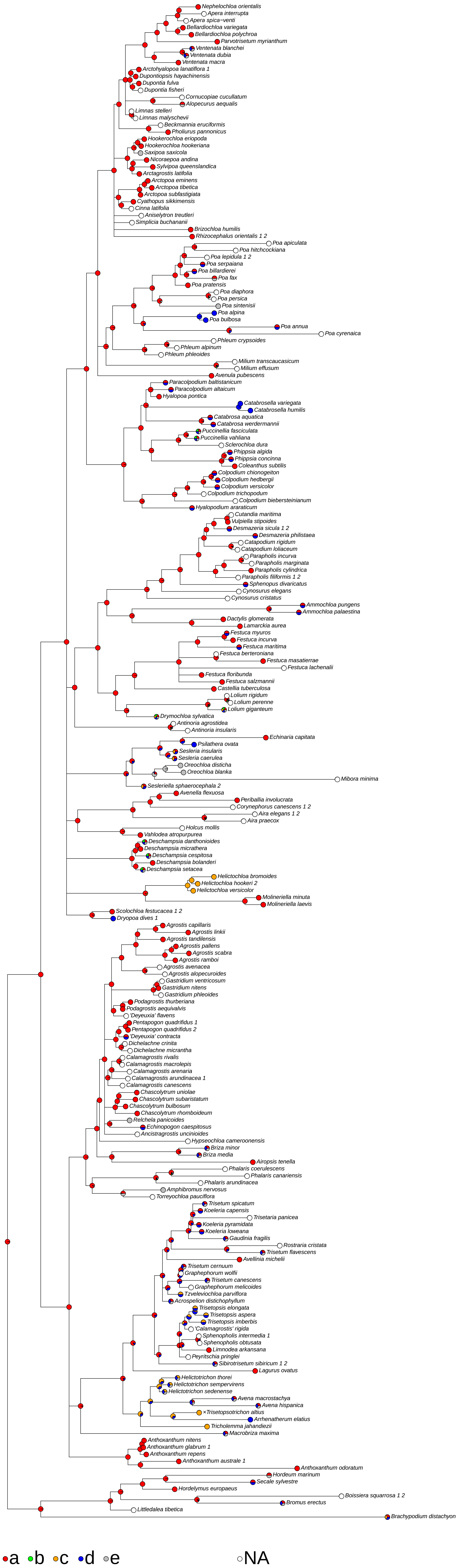

### (178) Rhachilla extension <indumentum>

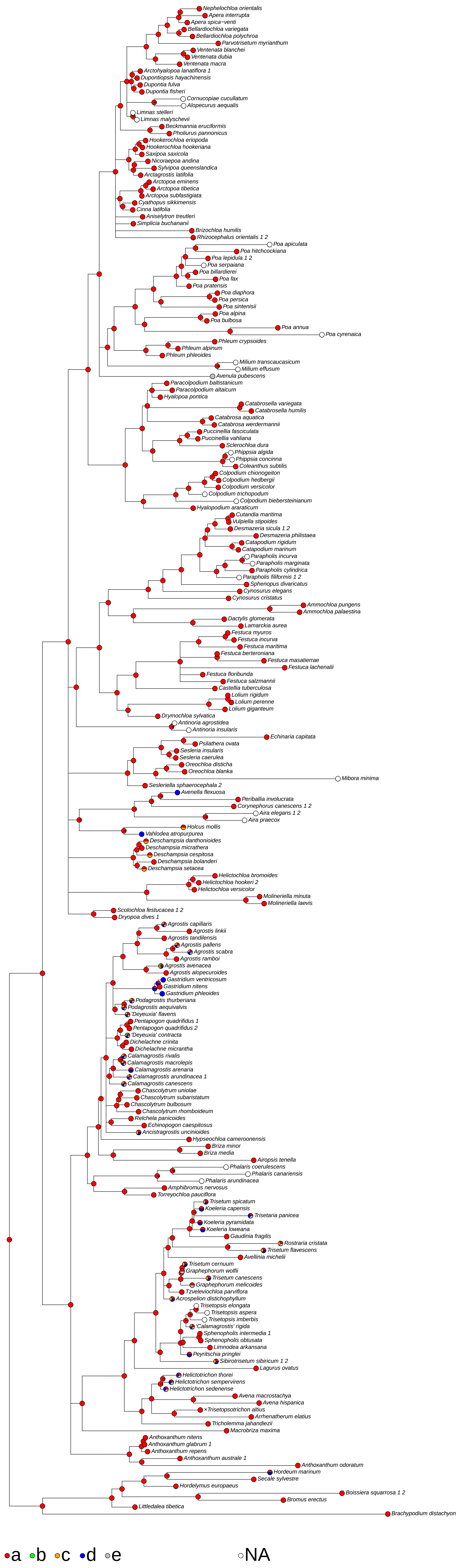

### (179) Apical sterile florets <resemblance to fertile>

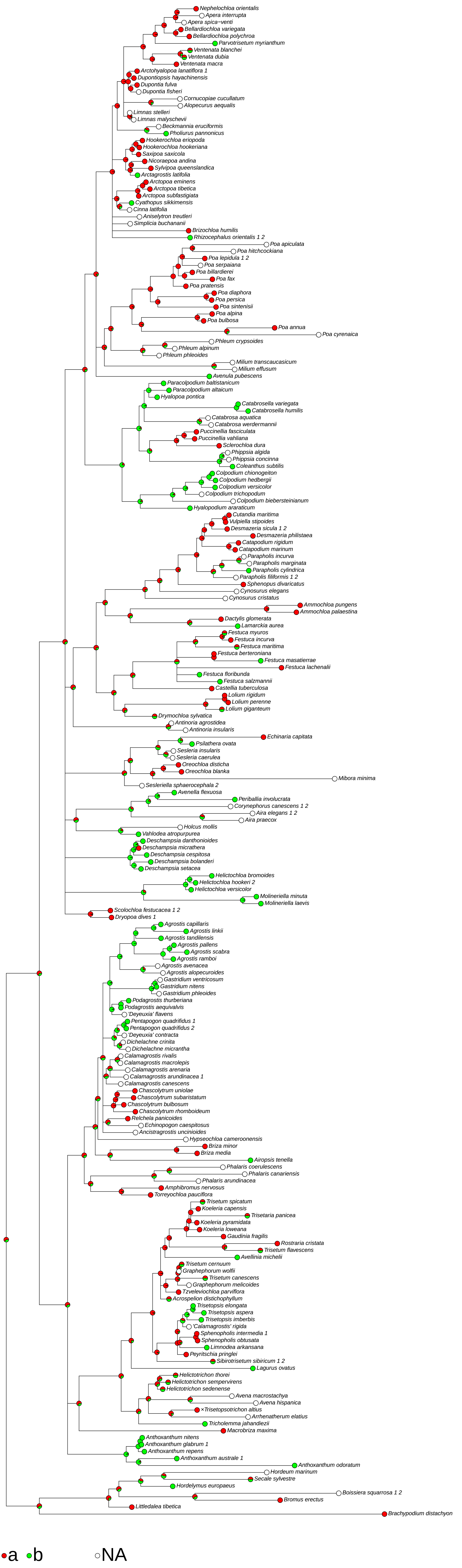

### (182) Anthers <number>

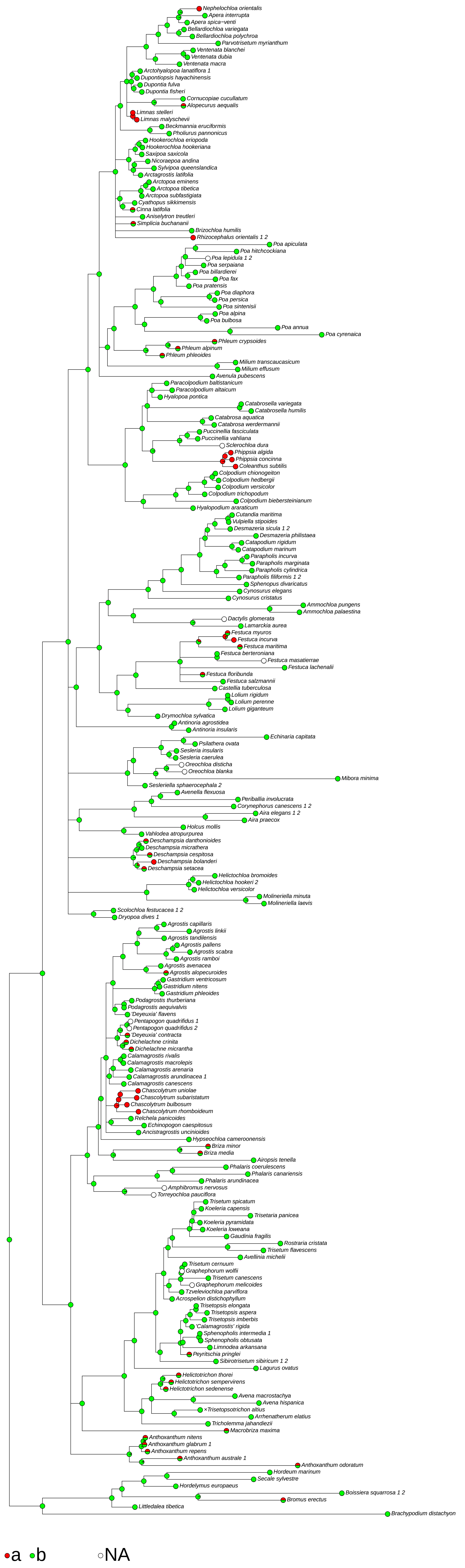

### (183) Anthers <length in mm>

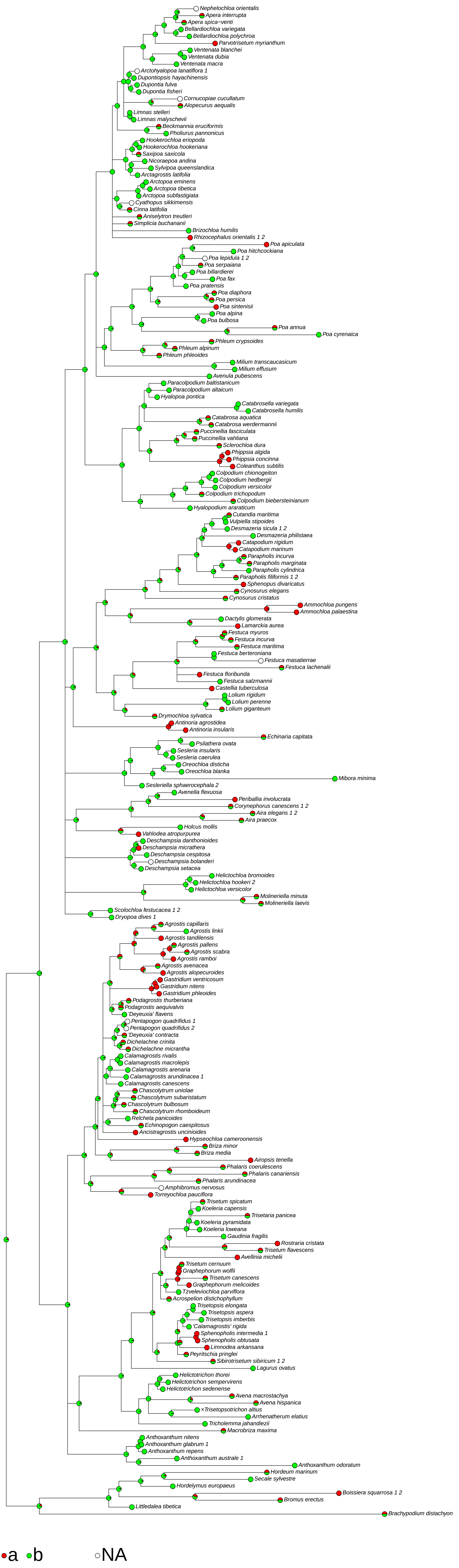

### (187) Caryopsis <length in mm>

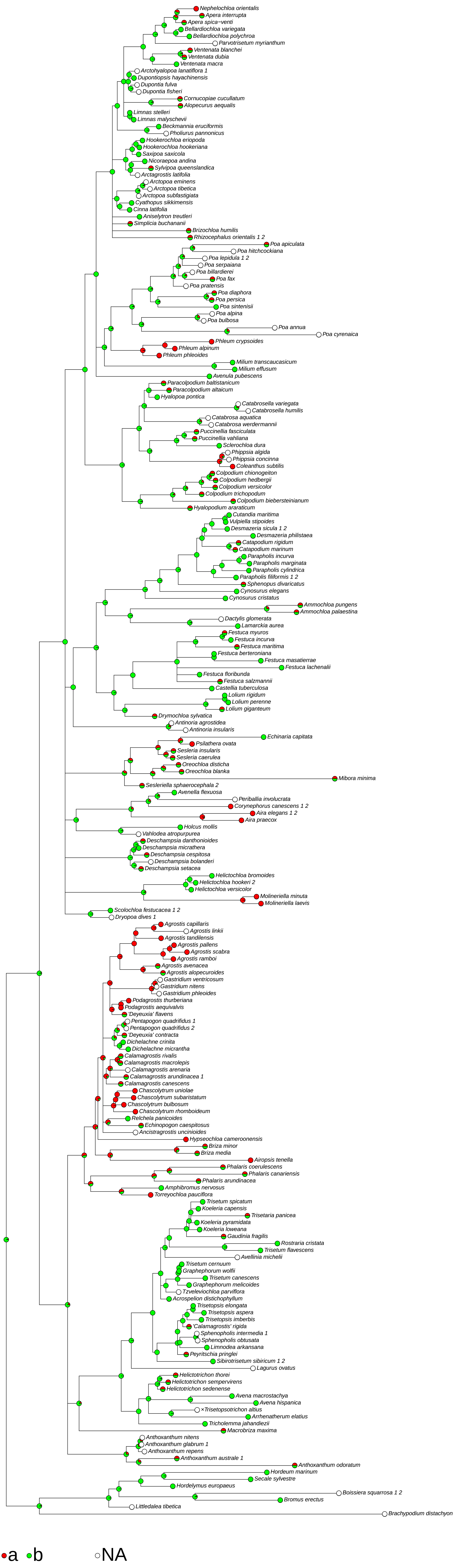

### (188) Hilum <shape>

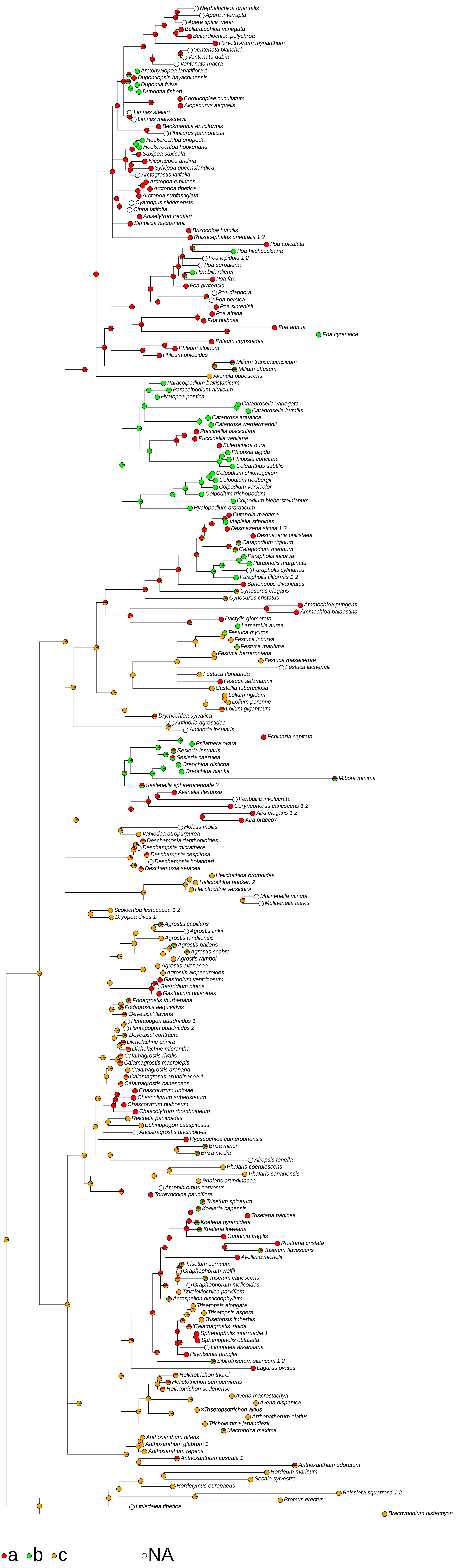
